# The tuatara genome: insights into vertebrate evolution from the sole survivor of an ancient reptilian order

**DOI:** 10.1101/867069

**Authors:** Neil J. Gemmell, Kim Rutherford, Stefan Prost, Marc Tollis, David Winter, J. Robert Macey, David L. Adelson, Alexander Suh, Terry Bertozzi, José H. Grau, Chris Organ, Paul P. Gardner, Matthieu Muffato, Mateus Patricio, Konstantinos Billis, Fergal J Martin, Paul Flicek, Bent Petersen, Lin Kang, Pawel Michalak, Thomas R. Buckley, Melissa Wilson, Yuanyuan Cheng, Hilary Miller, Ryan K. Schott, Melissa Jordan, Richard Newcomb, José Ignacio Arroyo, Nicole Valenzuela, Tim A. Hore, Jaime Renart, Valentina Peona, Claire R. Peart, Vera M. Warmuth, Lu Zeng, R. Daniel Kortschak, Joy M. Raison, Valeria Velásquez Zapata, Zhiqiang Wu, Didac Santesmasses, Marco Mariotti, Roderic Guigó, Shawn M. Rupp, Victoria G. Twort, Nicolas Dussex, Helen Taylor, Hideaki Abe, James M. Paterson, Daniel G. Mulcahy, Vanessa L. Gonzalez, Charles G. Barbieri, Dustin P. DeMeo, Stephan Pabinger, Oliver Ryder, Scott V. Edwards, Steven L. Salzberg, Lindsay Mickelson, Nicola Nelson, Clive Stone, Ngatiwai Trust Board

**Affiliations:** Allan Wilson Centre, Department of Anatomy, University of Otago, PO Box 56, Dunedin 9054, New Zealand; LOEWE-Center for Translational Biodiversity Genomics, Senckenberg Museum, 60325 Frankfurt, Germany; South African National Biodiversity Institute, National Zoological Garden, Pretoria, 0184, South Africa; School of Life Sciences, Arizona State University, 427 E Tyler Mall, Tempe, AZ 85281 USA; School of Informatics, Computing, and Cyber Systems, Northern Arizona University, 1295 S Knoles Drive, Flagstaff, AZ 86011 USA; School of Fundamental Sciences, Massey University, Private Bag 11 222, Palmerston North, 4442, New Zealand; Peralta Genomics Institute, Chancellor’s Office, Peralta Community College District, 333 East 8th Street, Oakland, CA 94606; School of Biological Sciences, The University of Adelaide, Adelaide, SA 5005, Australia; Department of Ecology and Genetics - Evolutionary Biology, Evolutionary Biology Centre (EBC), Uppsala University, Norbyvägen 18D, 75236 Uppsala, Sweden; Evolutionary Biology Unit, South Australian Museum, Adelaide, SA 5000, Australia; Amedes Genetics, Amedes Medizinische Dienstleistungen GmbH, Jägerstr. 61, 10117 Berlin, Germany; Museum für Naturkunde Berlin, Leibniz-Institut für Evolutions- und Biodiversitätsforschung an der Humboldt-Universität zu Berlin. Invalidenstraße 43, 10115. Berlin, Germany; Department of Earth Sciences, Montana State University, Bozeman, MT 59717; Department of Biochemistry, University of Otago, PO Box 56, Dunedin 9054, New Zealand; European Molecular Biology Laboratory, European Bioinformatics Institute, Wellcome Genome Campus, Hinxton, Cambridge CB10 1SD, UK; Section for Evolutionary Genomics, The GLOBE Institute, Faculty of Health and Medical Sciences, University of Copenhagen, Øster Farimagsgade 5, 1353 København K, Denmark; Edward Via College of Osteopathic Medicine, Blacksburg, VA 24060, USA; Center for One Health Research, Virginia–Maryland College of Veterinary Medicine, Blacksburg, VA 24060, USA; Institute of Evolution, University of Haifa, Haifa, 3498838, Israel; Manaaki Whenua - Landcare Research, Private Bag 92170, Auckland, New Zealand; School of Biological Sciences, The University of Auckland, Private Bag 92019, Auckland, New Zealand; School of Life and Environmental Sciences, The University of Sydney, Sydney, NSW 2006, Australia; Biomatters Ltd, Level 2, 18 Shortland Street, Auckland 1010 New Zealand; Department of Vertebrate Zoology, National Museum of Natural History, Smithsonian Institution, Washington DC 20560, USA; The New Zealand Institute for Plant & Food Research, Private Bag 92169, Auckland, New Zealand; Departamento de Ecología, Facultad de Ciencias Biológicas, Pontificia Universidad Católica de Chile, Avenida Libertador Bernardo O’Higgins 340, Santiago, Chile; Iowa State University, Department of Ecology, Evolution, and Organismal Biology, 251 Bessey Hall, Ames IA, 5011 USA; Instituto de Investigaciones Biomédicas CSIC-UAM, 28029-Madrid, Spain; Division of Evolutionary Biology, Faculty of Biology, Ludwig-Maximilian University of Munich, D-82152 Planegg-Martinsried, Germany; Centre for Genomic Regulation (CRG), The Barcelona Institute for Science and Technology, Barcelona, Catalonia, Spain, Universitat Pompeu Fabra (UPF), Barcelona, Catalonia, Spain; School of Biological Sciences, University of Canterbury, New Zealand; Global Genome Initiative, National Museum of Natural History, Smithsonian Institution, 1000 Constitution Ave., Washington, DC 20560, USA; AIT Austrian Institute of Technology, Center for Health and Bioresources, Molecular Diagnostics, Muthgasse 11, 1190 Vienna, Austria; San Diego Zoo Institute for Conservation Research, Escondido, CA 92027 USA; Department of Organismic and Evolutionary Biology and the Museum of Comparative Zoology, Harvard University, Cambridge, MA 02138 USA; Department of Biomedical Engineering, Johns Hopkins University, Baltimore, MD 21205 USA; School of Biological Sciences, Victoria University of Wellington, PO Box 600, Wellington 6140, New Zealand; Ngatiwai Trust Board, 129 Port Road, PO Box 1332, Whangarei, New Zealand

## Abstract

The tuatara (*Sphenodon punctatus*), the only living member of the archaic reptilian order Rhynchocephalia (Sphenodontia) once widespread across Gondwana, is an iconic and enigmatic terrestrial vertebrate endemic to New Zealand. A key link to the now extinct stem reptiles from which dinosaurs, modern reptiles, birds and mammals evolved, the tuatara provides exclusive insights into the ancestral amniotes. The tuatara genome, at ∼5 Gbp, is among the largest vertebrate genomes assembled. Analysis of this genome and comparisons to other vertebrates reinforces the uniqueness of the tuatara. Phylogenetic analyses indicate tuatara diverged from the snakes and lizards ∼250 MYA. This lineage also shows moderate rates of molecular evolution, with instances of punctuated evolution. Genome sequence analysis identifies expansions of protein, non-protein-coding RNA families, and repeat elements, the latter of which show an extraordinary amalgam of reptilian and mammalian features. Sequencing of this genome provides a valuable resource for deep comparative analyses of tetrapods, as well as for tuatara biology and conservation. It also provides important insights into both the technical challenges and the cultural obligations associated with genome sequencing.

---

The tuatara (*Sphenodon punctatus*) is an iconic and enigmatic terrestrial vertebrate, unique to New Zealand^1^. The only living member of the archaic reptilian order Rhynchocephalia (Sphenodontia) which last shared a common ancestor with other reptiles some 250 million years ago (Figure 1), the tuatara represents an important link to the now extinct stem reptiles from which dinosaurs, modern reptiles, birds and mammals evolved and is thus important for our understanding of amniote evolution^2^.

**Figure 1.**
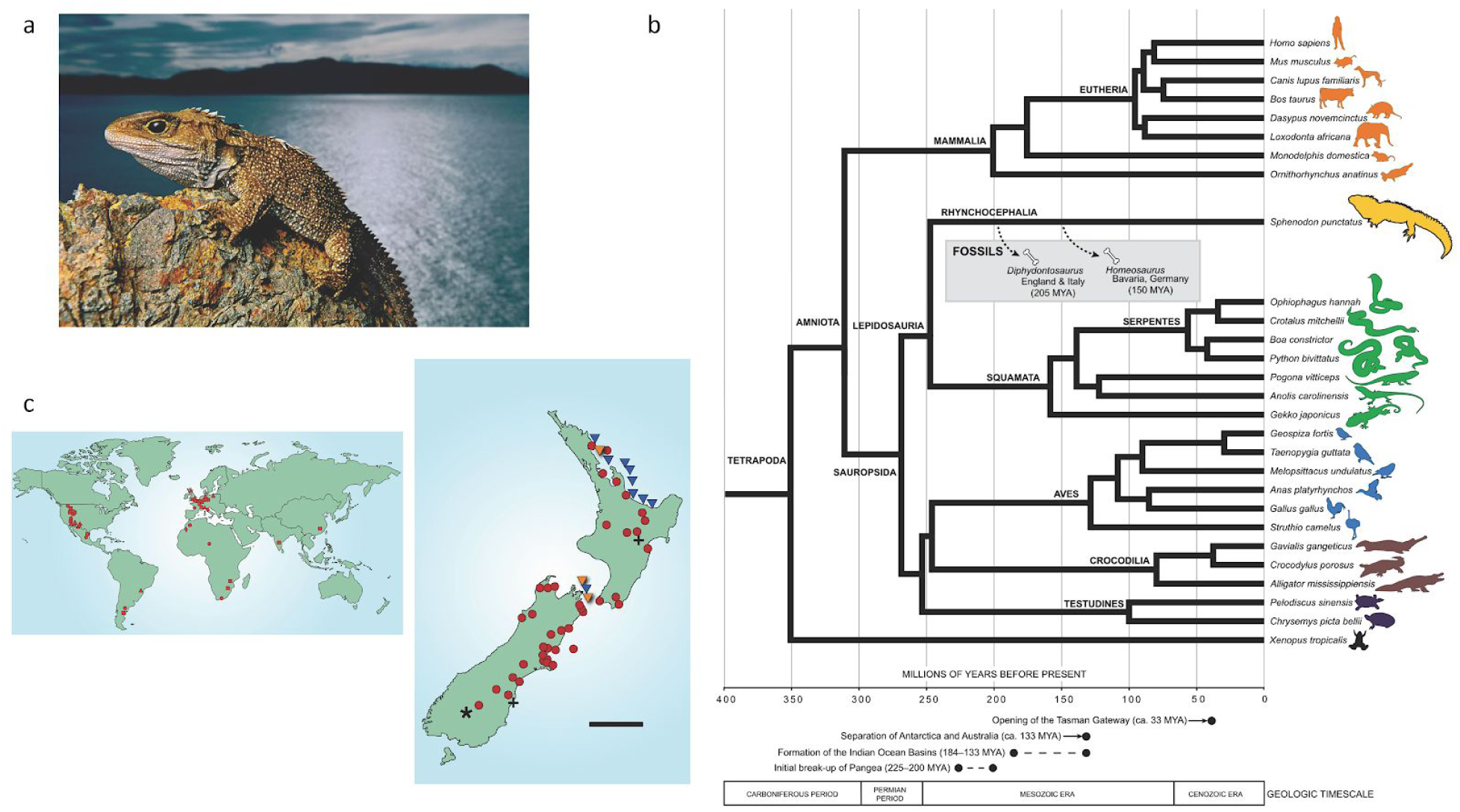
a) The tuatara, *Sphenodon punctatus*, is the sole survivor of the Order Rhynchocephalia. b) The rhynchocephalians appear to have originated in the Early Mesozoic (∼250-240 mya) and were common and globally distributed for much of that era (c). Their geographical range progressively contracted after the Early Jurassic (∼200-175 mya); the most recent fossil record outside of New Zealand is from Argentina in the Late Cretaceous (∼70 mya). c) The last bastion of the rhynchocephalians are 32 islands off the coast of New Zealand, which have recently been augmented by the establishment of ∼10 new island or mainland sanctuary populations using translocations. The current global population is estimated at ∼100,000 individuals. Rhynchocephalian fossil localities are redrawn from Jones et al. (2009). Legend: triangles, Triassic; squares, Jurassic; filled circles, Cretaceous; diamonds, Palaeocene; asterisk, Miocene; pluses, Pleistocene; open circles, Holocene; down triangles, extant populations; orange down triangles, populations investigated in this study. Scale bar represents 200 Km. Photo credit: Frans Lanting.

It is also a species of significance in other contexts. First, the tuatara is a *taonga*, or special treasure, for Māori, who hold that tuatara are the guardians of special places^1^. Second, the tuatara is internationally recognised as a critically important species vulnerable to extinction due to habitat loss, predation, disease, global warming and other factors^1^. Third, the tuatara displays a variety of morphological and physiological innovations that have puzzled scientists since the first description of the species in 1867^1,3^. These include a unique combination of features shared variously with lizards, turtles, and birds, which left its taxonomic position in doubt for many decades^1^. That taxonomic conundrum has been largely addressed using molecular approaches^4^, but the timing of the split of tuatara from the lineage that forms the modern squamata (lizards and snakes), the rate of evolution of tuatara, and the number of species of tuatara remains contentious^1^. Last, there are aspects of tuatara biology that are unique, or atypical of reptiles. These include the possession of a parietal eye whose function remains in dispute, a unique form of temperature-dependent sex determination (TSD) that sees females produced below, and males above, 22°C, extremely low basal metabolic rates, and significant longevity^1^.

To shed new insight into the biology of this extraordinary species we have sequenced the tuatara genome in partnership with Ngatiwai, the Māori iwi (tribe) who hold kaitiakitanga (guardianship) over the tuatara populations located on islands in the far North of New Zealand. This partnership, unique among the genome projects undertaken to date, had a strong practical focus on developing resources and information that will improve our understanding of tuatara and aid future conservation efforts. It is an exemplar for future genome initiatives that aspire to meet access and benefit-sharing obligations to indigenous communities required under the Convention for Biological Diversity^5^.

We find that the tuatara genome, as well as the animal, is an amalgam of ancestral and derived characteristics. Tuatara have 2n=36 chromosomes in both sexes, consisting of 14 pairs of macrochromosomes and four pairs of microchromosomes^6^. The genome size, estimated at 5 Gbp, is among the largest vertebrate genomes sequenced to date and this is predominantly explained by an extraordinary diversity of repeat elements, many of which are unique to tuatara. Given its size and complexity this genome has posed a variety of technical challenges, so we also outline how we have innovated and adapted a variety of technologies and assembly approaches to produce the *de novo* assembly of one of the largest and most complex high-quality vertebrate genomes undertaken to date.

## Sequencing, assembly, synteny and annotation

The tuatara genome assembly is 4.3 Gbps and consists of 16,536 scaffolds with an N50 scaffold length of 3 Mbp (Table S1). Genome assessment using BUSCO^7^ indicates 85.6% of sequences are present and complete based on the eukaryota gene set. Subsequent annotation identified 17,448 genes, of which 16,185 were identified as one-to-one orthologs (Supplementary Materials 2). Local gene-order conservation is high; 75% or more of the tuatara genes showed conservation with birds, turtles and crocodilians (Supplementary Figures 1 and 2). We also find that components of the genome, 15 Mbp and greater, are syntenic with other vertebrates, with the protein-coding gene order and orientation maintained between tuatara, turtle, chicken, and human (Supplementary Figures 1–3).

**Figure 2.**
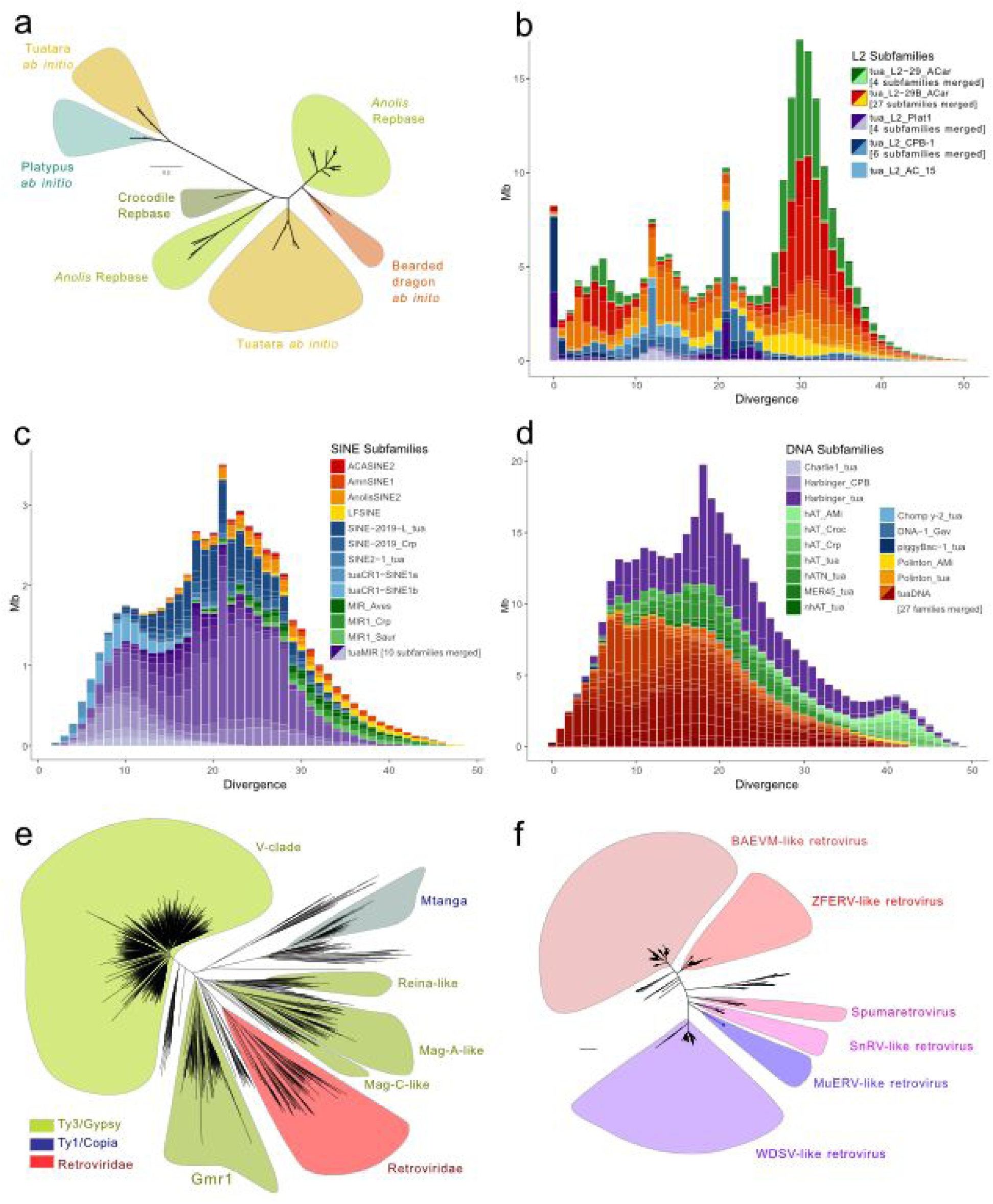
Analysis of the repeat landscape in the tuatara genome identifies unique repeat families, evidence of recent activity and a greater expansion and diversity of repeats than any other vertebrate. a) Phylogenetic analysis based on the RT domain of L2 repeats identifies two L2 subfamilies; one typical of other lepidosaurs and one that is similar to platypus L2. Phylogeny based on L2 elements >1.5 kbp long with RT domain >200 aa. b) Landscape plot of L2 retrotransposons, the dominant LINEs observed in the tuatara genome, accounting for 10% of the genome. Only LINE subfamilies that occupy more than 1,000 bp are shown. c) Landscape plot of SINE retrotransposons suggests the tuatara genome is dominated by MIR sequences most typically associated with mammals, with tuatara now the amniote genome in which the greatest MIR diversity has been observed. Only SINE subfamilies that occupy more than 1,000 bp are shown. d) Landscape plot of DNA transposons including 24 novel DNA transposon families from tuatara. e) The tuatara genome contains ∼7500 full-length long terminal repeat (LTR) retroelements including nearly 450 endogenous retroviruses (ERVs). The general spectrum of tuatara LTR retroelements is comparable to that of other sauropsids, but lacks Bel-Pao LTR retroelements which are present in anole, and has an abundance of Ty1/Copia retroelements. f) Tuatara ERVs span five major retroviral clades and include at least 37 complete spumaretroviruses previously found only in coelacanth, sloth, aye-aye and Cape golden mole. This is the first report of this ancient group of ERVs in a sauropsid.

**Figure 3.**
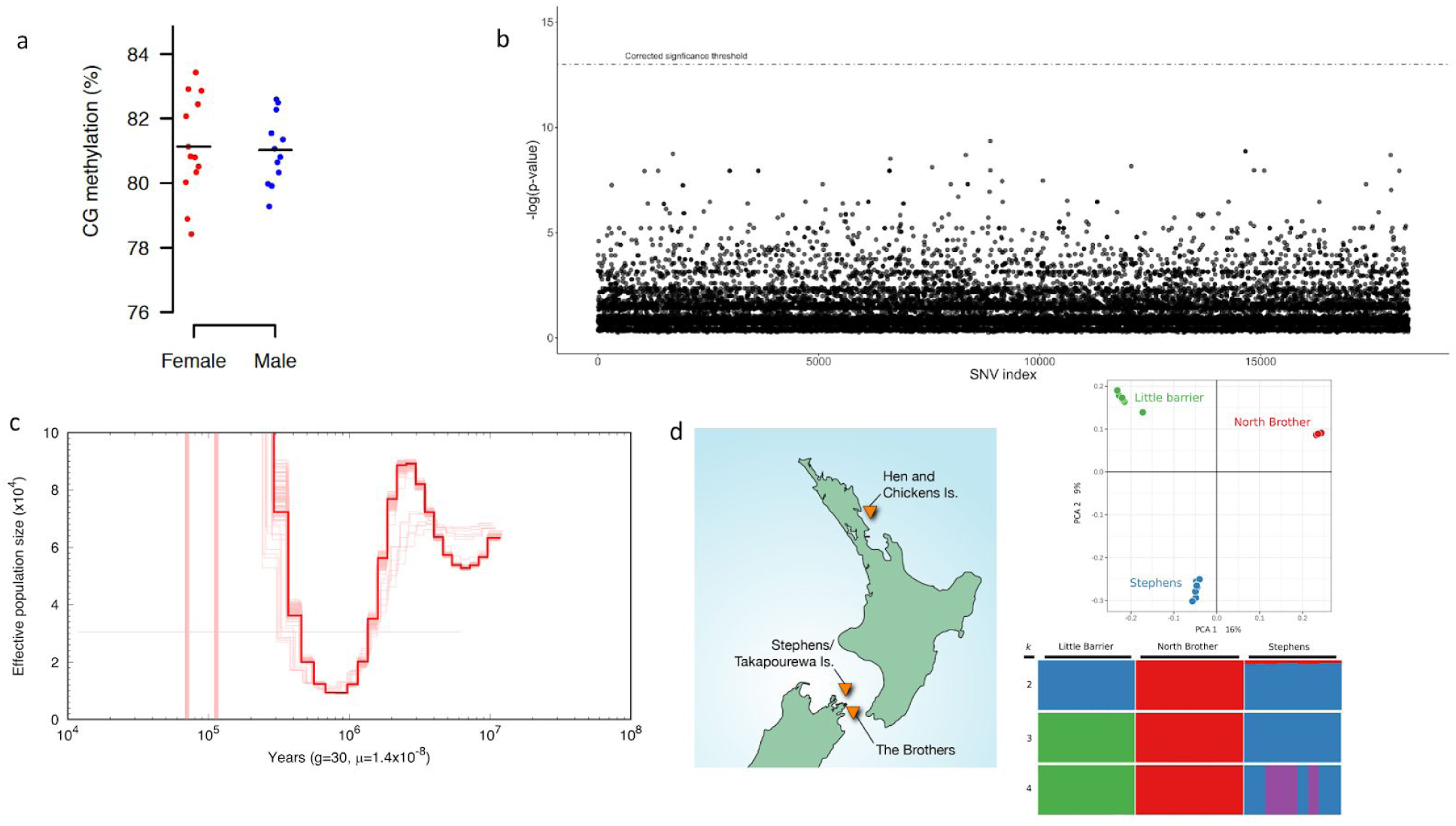
a) Methylation levels in the tuatara genome are high, mean 81%, but show no significant differences among the sexes. b) No single nucleotide variant (SNV) is significantly differentiated with respect to sex in the tuatara genome. The dashed line represents the threshold for statistical significance after accounting for multiple testing. c) PSMC plots of the demographic history of tuatara using a mutation rate of 1.4×10-8 substitutions per site per generation and a generation time of 30 years. d) We examined the three known axes of genetic diversity in tuatara: northern New Zealand (Little Barrier Is./ Hauturu), and two islands in Cook Strait (Stephens Is. / Takapourewa; North Brother Island) using genotype by sequencing methods. Principal component analysis and Structure plots demonstrate significant structure among tuatara populations and strongly support prior suggestions that tuatara on the Brothers Island are genetically distinct and warrant separate management.

## Genomic architecture

### Repetitive elements

In-depth repeat analyses revealed that at least (64%) of the tuatara genome assembly is composed of repetitive sequences, made up of transposable elements (31%) and low copy number segmental duplications (33%). While the total transposable element content is similar to other reptiles^8–11^, there are notable differences in the types of repeats found which make this genome appear more mammal-like than reptile-like. Furthermore, a number of repeat families show evidence of recent activity and greater expansion and diversity than seen in other vertebrates (Figure 2).

Among long interspersed elements (LINEs), L2 elements account for most of the LINE sequences in the tuatara genome (10% of the genome) and some may still be active, a remarkable observation given that CR1 elements are the dominant LINE in other sauropsid genomes^8–10, 12–14^ but only account for a relatively small fraction (∼4%) of the tuatara genome (Figure 2 panel a and b; Table S2; Supplementary Materials 4). There are also potentially active CR1 elements in tuatara (Supplementary Materials 4). L1 elements, which are prevalent in placental mammals account for only a tiny fraction of the genome (<1%) (Table S2). Phylogenetic analysis of L2 elements identifies an L2 subfamily present in tuatara, but absent from other lepidosaurs, that is also common in monotremes (Supplementary Materials 4, Figures 4.3 and 4.4) where these repeats account for 18–23% of the genome^15^. These data suggest that the LINE composition of the tuatara genome is more monotreme-like than reptile-like. Whether this implies an ancestral presence of these repeats or an ancient horizontal transfer event remains to be determined, but it appears that stem sauropsid ancestors had quite a different repeat composition than has been inferred from prior comparisons using birds and lizards.

Many short interspersed elements (SINEs), TEs which depend on the enzymatic machinery of LINEs, belong to copies from ancient CORE-SINEs present in all amniotes^16^. We also identified at least 16 SINE subfamilies which were recently active in the tuatara genome (Figure 2 panel c; Supplementary Materials 5). Most of these SINEs are from MIR subfamilies. While consistent with the pattern described above for L2 elements, which are required for mobilisation of these SINEs, this pattern is unusual for sauropsids but reminiscent of the repeat landscape of monotreme and marsupial mammals^17, 18^. Notably, the diversity of tuatara MIR subfamilies is higher than in marsupials^17^, the highest previously known in amniotes. In the human genome, hundreds of fossil MIR elements act as chromatin and regulatory domains^19^, which suggests that the recent activity of diverse tuatara MIR subfamilies may have influenced regulatory rewiring on rather recent evolutionary timescales.

We detected 24 novel DNA transposon families with no sequence similarity to known families or each other, suggesting frequent tuatara germline infiltration by novel DNA transposons through horizontal transfer^20^. No fewer than 30 DNA transposon subfamilies were recently active, spanning a diverse range of cut-and-paste transposons and polintons (Figure 2 panel d; Supplementary Materials 5). This diversity is higher than in anole lizards^21^ and vespertilionid bats^22–24^, which are among the few amniotes identified with recently active DNA transposons. Strikingly, we found thousands of identical DNA transposon copies, suggesting very recent, and/or ongoing, activity. It is thus possible that cut-and-paste transposition is shaping the tuatara genome in similar ways as in vespertilionid bats^22–24^.

We identified approximately 7,500 full-length long terminal repeat (LTR) retroelements including endogenous retroviruses (ERVs), which we classified into at least 12 groups using phylogenetic reconstructions of their retrotranscriptase domains (Figure 2 panel e and f; Supplementary Materials 6). We identified nearly 450 ERVs from five major retroviral clades and some 5,500 Ty3/Gypsy elements representing five major clades. Notably, a Ty1/Copia element (Mtanga-like) is abundant in the tuatara genome with more than 1,140 full-length elements. The general spectrum of tuatara LTR retroelements is comparable to that of other sauropsids, but lacks Bel/Pao LTR retroelements which are present in anole^25^. An unexpected result is the presence of at least 37 complete spumaretroviruses in the tuatara genome (Figure 2 panel f; Supplementary Materials 6). Spumaretroviruses are considered among the most ancient ERVs^26, 27^ and were previously found only in coelacanth^28^, sloth^29^, aye-aye^30^ and Cape golden mole^31^. Thus, the occurrence of several full-length endogenous spumaretroviruses in tuatara, some encoding a complete envelope gene, is the first in a sauropsid genome.

The tuatara genome further contains more than 8,000 non-coding RNA (ncRNA)-related elements. The bulk of these elements (∼6,900) are derived from recently active transposable elements. Most overlap with a novel CR1-mobilized SINE (Figure 2 panel c; Supplementary Materials 7), which contains nearly 3,900 elements with strong matches to tRNA covariance models (over 10,000 elements in total). The remaining high copy number elements include sequences closely related to ribosomal RNAs, spliceosomal RNAs and signal recognition particle RNAs.

Finally, an extraordinarily high proportion (33%) of the tuatara genome originates from low copy number segmental duplications, with 6.7% of these of recent origin based on their high level of sequence identity (>94% identity), which is more than seen in other vertebrates^15^. The tuatara genome is 2.4x larger than the anole genome and this difference appears thus to be driven disproportionately by segmental duplications.

The overall repeat architecture of tuatara is unlike anything previously reported, showing a unique amalgam of features previously viewed as characteristic of either reptilian or mammalian lineages. This truly reflects the phylogenetic position of this astonishing evolutionary relic – a combination of ancient amniote features as well as a surprisingly dynamic and diverse repertoire of lineage-specific TEs.

### Epigenetic features

Low-coverage sequencing using post-bisulfite adaptor tagging (PBAT)^32^ shows ∼81% of CpG sites are methylated in tuatara (Figure 3a). When PBAT is used to compare against other vertebrate species, DNA methylation levels are the highest reported for an amniote — markedly different methylation to those observed in mouse, human (∼70%) and chicken (∼50%) and more similar to that seen in *Xenopus* (82%) and zebrafish (78%). One possible explanation for this high rate of DNA methylation is the extraordinary level of repetitive elements found in this genome, many of which appear recently active and might be regulated via DNA methylation (see above).

Examination of the CpG distribution in tuatara, sheds new light on the commonalities and diversity of nCpG distribution across vertebrates, a trait that is linked to DNA methylation and may consequently impact gene expression (Supplementary Materials 8). Tuatara display a bimodal nCpG distribution in all genomic regions examined (gene promoters, exons, introns, and intergenic sequences), similar to recent reports in painted turtles^33^ and alligator, both of which have tempertaure dependent sex determination (TSD, Supplementary Figure 4). The bimodal nCpG pattern seen in introns of these TSD reptiles may mark a divide between TSD taxa and the genotypic sex determination (GSD) vertebrates previously reported to display a unimodal nCpG distribution in introns^33^. However, testing explicitly for the existence of bimodality in all genomic regions examined using mixed models, we find a bimodal pattern of nCpG that appears to be ancestral and conserved across all vertebrates irrespective of their sex-determining mechanism. This finding agrees with the recent detection of bimodal DNA methylation in gene bodies in human, chicken, fish, and tunicates^34^.

### The mitochondrial genome

The mitochondrial (mt) genome in the tuatara reference animal is 18,078 bp, containing 13 protein coding, 2 rRNA, and 22 tRNA genes, standard among animals (Supplementary Figure 5). This contradicts previous reports^35, 36^ that the tuatara mt-genome lacks 3 genes: ND5, tRNA^Thr^, tRNA^His^. These three genes are found, with an additional copy of tRNA^Leu(CUN)^ and an additional non-coding block (NC2) in a single segment of the mt-genome. This Illumina assembly result is confirmed with 40 Oxford Nanopore mitochondrial DNA reads, of which one covers the entire molecule. Three non-coding areas with Control Region (heavy strand replication origin, O_H_) features (NC1–3), and two copies of tRNA^Leu(CUN)^ adjacent to NC1 and NC2 possess identical or near identical sequence, possibly as a result of concerted evolution. A stem-and-loop structure is observed in regions encoding tRNA^Asn^ and tRNA^Cys^, which may supplement for the missing structure normally observed for the origin of light strand replication (O_L_) in this location^37^. The tRNA^Lys^ gene is duplicated with the first copy possibly a pseudo-gene^35^. The tRNA^Cys^ gene encodes a tRNA with a D-arm replacement loop^38^.

## The unique biology of the tuatara is reflected in its genome

Befitting its taxonomic distinctiveness, we find that the tuatara genome displays multiple innovations in genes associated with immunity, odour reception, thermal regulation and selenium metabolism.

### MHC

Genes of the major histocompatibility complex (MHC) play an important role in disease resistance^39^, mate choice and kin recognition^40^ and are among the most polymorphic in the vertebrate genome. Prior work^41^ mapped the core MHC genes to tuatara chromosome 13q, although a small number of class I genes map to chromosome 4p. Here, annotation of tuatara MHC regions and comparisons to the gene organization to six other representative species (Supplementary Materials 9, Supplementary Figure 6), identified 56 genes spanning 13 scaffolds, including six class I genes, six class II genes, 19 class III genes, 18 framework genes, and seven extended class II genes. Four class I and four class II genes were not linked to other MHC related genes and were excluded from the comparative analysis as they may be located outside the core MHC region on chromosome 13^41^.

The genomic organization of tuatara MHC is most similar to that of the green anole, which we interpret as being typical of the Lepidosauria (Supplementary Figure 6). Unlike the chicken MHC, which was proposed to represent the minimal essential MHC gene cluster^42^, tuatara, anole, alligator, and turtle show a high gene content and complexity that is more similar to the MHC regions of amphibians and mammals than birds. While the majority of genes annotated in the tuatara MHC are well conserved as one-to-one orthologs, extensive genomic rearrangements were observed among these distant lineages (Supplementary Figure 6). The most noteworthy rearrangement lies in the relative locations of genes in the class I and class III regions (as defined by their locations in the human MHC), with these two regions inverted between the mammalian and frog MHC, and the genes interspersed in the tuatara and anole genomes. This finding raises questions over the traditional categorization of these genes as class I or class III region genes, as evolutionarily there is no explicable difference between the two sets of genes in terms of gene clustering, organization, or function.

### Vision and smell

The tuatara is a highly visual predator that is able to capture prey under extremely low light conditions^1^. Despite this extreme nocturnal visual adaptation, the tuatara shows strong morphological evidence of an ancestrally diurnal visual system including a fovea^43^ and photoreceptors that while superficially rod-like appear to be evolutionary derived from cones^44^, similar to the situation in snakes and geckos^45^. The tuatara genome provides an opportunity to uncover the molecular basis of these morphological adaptations. A search for visual genes identified all five vertebrate opsin genes in the tuatara genome, and a comparative analysis revealed one of the lowest rates of visual gene loss for any amniote (Supplementary Materials 10). This low gene loss contrasted sharply with high rates of gene loss in ancestrally nocturnal lineages (Supplementary Figure 7).

Analysis of the selective constraint operating upon a subset of visual genes (those involved in phototransduction) showed strong negative selection and no concerted evidence of long-term shifts in selective pressures found in other groups that have evolutionary modified photoreceptors i.e., rod-like cones and cone-like rods^46^ (Supplementary Materials 10). Collectively, these results suggest a unique path to nocturnal adaptation in tuatara from a diurnal ancestor. Furthermore, the retention of five visual opsins and conserved nature of the visual system strongly suggests tuatara possess robust colour vision, potentially at low light levels where most animals are colour-blind. This surprisingly broad visual repertoire may be explained by the dichotomy in tuatara life-history: juvenile tuatara, often take up a diurnal and arboreal lifestyle to avoid the terrestrial, nocturnal, adults that may predate them^1^.

Odorant receptors (ORs), expressed in the dendritic membranes of olfactory receptor neurons, are responsible for the detection of odours. Species that depend strongly on their sense of smell to interact with their environment, find prey, identify kin and avoid predators may be expected to have a large number of ORs. Four hundred and seventy two genes encoding predicted ORs were identified from the genome of tuatara (Supplementary Materials 11). Of these, 341 sequences were predicted to be intact ORs, encoding proteins 303–345 amino acids in length (<u>10.5281/zenodo.2592798</u>). The remainder consisted of 9 full length sequences missing the initial start codon, 60 partial genes, 22 genes containing frameshifts, and 40 presumed pseudogenes. Many ORs were found as tandem arrays, with up to 26 found on a single scaffold. Tuatara OR genes do not contain introns, following the conventional vertebrate OR gene structure^47, 48^.

Odorant receptor gene number and diversity varies greatly amongst the Sauropsida. Birds possess from 182–688 genes, the green anole lizard has 156 genes, while crocodilian and testudine repertoires are much larger containing 1000–2000 genes^47–51^. Tuatara has a similar number of ORs to birds, but many more than the genome of the single lizard species published to date. Interestingly the tuatara genome contains a high percentage of intact OR genes, 85%, compared to other published OR sets from sauropsid genomes. This may reflect a strong reliance on olfaction by tuatara and therefore pressure to maintain a substantial repertoire of ORs (Supplementary Figure 8). There is some evidence that olfaction may play a role in identifying prey^52, 53^ and that cloacal secretions may potentially act as pheromones^54, 55^.

### Thermoregulation

The tuatara, a behavioural thermoregulator, is notable for having the lowest optimal body temperature of any reptile (16–21 °C). They can remain active at temperatures as low as 5 °C, while temperatures over 28 °C are generally fatal^1^. Transient Receptor Potential (TRP) ion channel genes play an important role in thermoregulation, participating in thermosensation and cardiovascular physiology^56 57^. Inhibition of cold- and heat-sensitive TRP channels in the saltwater crocodile produces alterations in thermoregulation^58^ leading us to hypothesise that TRP genes may be linked to the extraordinary thermal tolerance of tuatara. We therefore undertook comparative genomic analysis of TRP genes in tuatara and six other amniotes (lizard, snake, alligator, chicken, turtle, and human) to look for evidence of genomic adaptations related to thermoregulation in tuatara (Supplementary Materials 12). We identified 37 TRP-like sequences, spanning all seven described subfamilies of TRPs in the tuatara genome (Supplementary Figure 9) — an unusually large repertoire of TRP genes.

Among this suite of genes we identified new thermo- and non-thermosensitive TRP genes that appear to result from gene duplication, and have been differentially retained in tuatara. For example, tuatara is unusual in possessing an additional copy of a thermosensitive TRPV-like gene (TRPV1/2/3) classically linked to the detection of moderate to extreme heat^56^ — a feature it shares with turtles (Supplementary Figure 9). A strong signature of positive selection among heat-sensitive TRP genes (TRPA1, TRPM and TRPV) was also observed.

In general these results show a high rate of differential retention and positive selection in genes for which function in heat sensation is well established^56 57^. Given the role of these genes, it seems probable that these genomic changes in TRP genes are associated with the evolution of thermoregulation in this species.

### Selenoproteins and tRNASec genes

Tuatara are, barring tortoises, the longest-lived reptiles, likely exceeding 100 years^1^. This enhanced life-span may be linked to genes that afford protection against reactive oxygen species either directly as antioxidants or by regulating redox pathways^59^. One class of genes known to afford such protection are the selenoproteins, proteins that incorporate selenium in the form of selenocysteine (Sec), the 21st amino acid. The human genome encodes 25 selenoproteins whose roles include antioxidation, redox regulation, thyroid hormone synthesis, calcium signal transduction and others^60^.

We identified 26 selenoprotein genes in the tuatara genome, along with genes encoding the machinery for selenoprotein synthesis (Supplementary Materials 13). The number of selenoproteins, and protein families to which they belong, are similar to other vertebrate genomes^61^, but for the presence of two selenoproteins from the SELENOW (previously SelW) family. SELENOW is a short protein with unknown function that has a thioredoxin domain (Rdx family, PF10262). Two SELENOW selenoproteins, SELENOW1 and SELENOW2, were predicted to be present in the last common ancestor of vertebrates, but most genomes have only one of the two genes^61^. SELENOW2 was predicted to have been lost in the amniote stem (sauropsids and mammals). The presence of SELENOW2 in tuatara suggests that this loss must have occurred later in evolution, after the split of sauria and mammals.

The selenocysteine-specific tRNA (tRNA-Sec) plays a central role in both the biosynthesis and incorporation of Sec^60^ and is usually present as a single functional copy, as is observed in the squamates. In tuatara we identified four eukaryotic tRNA-Sec genes in the same scaffold within a ∼5 kbp region of the genome, each of which appears to be functional (Supplementary Materials 13).

While further work is needed, the unusual selenoprotein gene and tRNA composition of tuatara may be linked to the extraordinary longevity of tuatara, or might arise as a response to the low levels of selenium and other trace elements in New Zealand’s terrestrial systems.

### Sex determination genes and markers

Tuatara have a unique mode of temperature dependent sex determination (TSD) where higher egg incubation temperatures result in males^1^. Thus we were interested to determine which genes previously implicated in vertebrate sex determination were conserved in tuatara (Table S3). We found orthologs for many genes known to act antagonistically in masculinising (e.g., *sf1*, *sox9*) and feminising (e.g., *rspo1, wnt4*) gene networks to, respectively, promote testicular or ovarian development^62^. We also found orthologs of several genes recently implicated in TSD, including *cirbp*^62^. Orthologs for some notable vertebrate sex determining genes (e.g., *dmrt1* and *amhI*) were absent in our multiple species alignments. We suspect these are not genuinely absent in tuatara and result from alignment failure due to high sequence divergence, because of the prior identification of *dmrt1* in our genome data^63^ and its earlier localisation in the tuatara genome by gene mapping^6^.

Tuatara have no obviously differentiable sex chromosomes^6^, but we hypothesized that sex specific markers might be identifiable through comparison of known sex males and females at a genomic and epigenetic level. We found no significant sex specific differences in global CG methylation between males and females (Figure 3a). Likewise, a search for sex specific SNPs^64^ using population genomic resources (detailed below) identified no sex specific SNPs (Figure 3b).

## Phylogeny, evolutionary rates and evidence for punctuated evolution

Albert Günther discovered that the “remarkable saurian” tuatara, described in the 18th century as “a monstrous animal of the lizard kind” is not a lizard but rather a unique reptile classified in a separate taxonomic order, Rhynchocephalia^3^. We placed the tuatara within the context of vertebrate evolution by carrying out multiple phylogenomic analyses that incorporated both whole genome alignments and clusters of single-copy orthologs (Supplementary Materials 14 and 15), recapitulating many patterns observed from the fossil record corroborated during the genomic era (Figure 1). After their appearance ∼312 million years ago^65^, amniote vertebrates diversified into two groups: the synapsids, which include all mammals, and the sauropsids, which include all reptiles and birds. By analysing 818,968 fourfold degenerate sites obtained from 27 vertebrates in a maximum likelihood framework, we obtained full phylogenomic support for a monophyletic Lepidosauria, marked by the divergence of the tuatara lineage from all squamates (lizards and snakes) during the early part of the Triassic Period ∼246 million years ago as estimated using a penalized likelihood method (Supplemental Materials 14). The phylogenetic placement and age of divergence for the tuatara was similarly recovered by analysing 245 single-copy gene orthologs using concatenated, species tree, and molecular clock methods (Figure 1; Supplementary Materials 15 and 16).

The rate of molecular evolution in the tuatara has been suggested to be paradoxically high in contrast to the apparently slow rate of morphological evolution^66, 67^. However, we find that the actual divergence in terms of DNA substitutions per site per million years at fourfold degenerate sites has been relatively low, particularly with respect to lizards and snakes, making it the slowest-evolving lepidosaurian reptile yet analysed (Supplementary Figure 10). We also find that amniote evolution in general can be described by a model of punctuated evolution, where the amount of genomic change is related to the degree of species diversification within clades^68, 69^. The tuatara falls well below this trend, suggesting this “living fossil” has accumulated substitutions at a rate expected given the lack of rhynchocephalian diversity (Supplementary Figure 11; Supplementary Materials 16). This suggests that rates of phenotypic and molecular evolution were not decoupled throughout amniote evolution^70^.

## Patterns of selection

In two different sets of analyses, we find that most genes exhibit a pattern of molecular evolution that suggests the tuatara branch evolves at a different rate than the rest of the tree (Supplementary Materials 17, Table S4). Approximately 659 of 4,284 orthologs tested had significantly different ω values (ratios of non-synonymous to synonymous substitutions or dN/dS) on the tuatara branch relative to other birds and reptiles tested. Although none of these orthologs had ω values suggestive of strong positive selection (i.e. > 1), the results do indicate that shifts in patterns of selection are affecting many genes and functional categories of genes across the tuatara genome, including genes involved in RNA regulation, metabolic pathways and general metabolism.

For genes involved in sex determination, we find that the entire length of the tree is shorter and that dN and dN/dS values are significantly lower than for typical genes in the genome (Supplementary Materials 17). Thus, while we find that most genes available for these analyses show strong levels of purifying selection, genes involved in sex determination are even more conserved, highlighting their critical function for reproduction and survival^62^.

## Population genomics

Once widespread across the supercontinent of Gondwana, Rhynchocephalia are now represented by a single species, the tuatara, found on a few offshore islands in New Zealand (Figure 1c). Tuatara have declined because of introduced pests and habitat loss^1^. They remain at risk of extinction due to their highly restricted distribution and the threats imposed by disease and climate-change-induced changes in sex ratios, which could profoundly impact their survival^71^. Earlier population genetic studies have shown populations in northern New Zealand are genetically distinct from those in the Cook Strait, and that the population on North Brother Island in the Cook Strait might be a distinct species^72^. Although subsequent studies have not found evidence for species-status^73^, this population is still managed as a separate conservation unit.

We took advantage of the tuatara reference genome to perform the first ancestral demographic and population genomic analyses of this species. First, we investigated genome wide signals for demographic change using PSMC (Supplementary Materials 18). The reconstructed demography (Figure 3c) reveals an increase in effective population size (N_e_) detectable around 10 Mya, a drastic decrease in N_e_ about 1–3 Mya, and a rapid increase in N_e_ between 500 kya and 1 Mya. These events correlate well with the known geological history of New Zealand^74^, reflecting an increase in available landmass following the Oligocene drowning, a period of significant climatic cooling that likely reduced tuatara habitat, and the formation of land bridges that facilitated population expansion.

Next, we obtained population samples spanning the three axes of genetic diversity in tuatara^73, 75^: northern New Zealand (Little Barrier Is./ Hauturu), and two islands in Cook Strait (Stephens Island / Takapourewa; North Brother Island) (Figure 3d), sampling 10 individuals each. We used a genotype-by-sequencing approach to generate 22 Gb of data from these individuals (Supplementary Materials 19). After applying filters and thinning the data to analyse only unlinked loci, we obtained 52,171 single nucleotide variants (SNVs) for population genomic analysis.

Analysis of the SNV dataset reveals that modern tuatara populations are highly differentiated (Figure 3d). Our genome-wide estimate of the fixation index (F_ST_) is 0.54, and more than two thirds of variable sites have an allele which is restricted to a single island. All of the island populations have relatively low genetic diversity (nucleotide diversity ranges from 8 × 10^-4^ for North Brother to 1.1 × 10^-3^ for Little Barrier). The low within-population diversity and marked population structure we observe in tuatara suggests modern island populations were isolated from each other sometime during the last glacial maximum at ∼18 kya.

Our results also support the distinctiveness of the Brothers Island tuatara, which has variously been described as *Sphenodon punctatus* or *Sphenodon guntheri*^72^. Recently the two species have been again collapsed to one^73^, but we find that the Brothers Island population forms a distinct group in all clustering analyses (Figure 3d). This population is highly inbred and shows evidence of a severe bottleneck, most likely reflecting a founder event around the last glaciation^75^. It is not clear if the distinctiveness we observe is due to allele frequency changes brought about by this bottleneck, or is reflective of a deeper split in the population history of tuatara. Regardless of the historical reasons for the North Brother population’s genetic distinctiveness, it is an important source of genetic diversity in tuatara, containing 8,480 private alleles in our dataset, and warrants ongoing conservation as an independent unit.

## A cultural dimension

The tuatara is a taonga, or special treasure, for many Māori — notably Ngātiwai and Ngāti Koata who are the kaitiaki (guardians) of tuatara. The tuatara genome project was undertaken in partnership with Ngātiwai iwi; our common goal being to increase knowledge and understanding of tuatara to aid the long term conservation of this species. Ngātiwai were involved in all decision-making regarding the use of the genome data by potential project partners; for each new project proposed we discussed the benefits that might accrue from this work and how these could be shared. The need to engage with, and protect the rights of, indigenous communities in such a transparent way has seldom been considered in the genomic projects published to date, even those related to human genomics, but is now a mandated consideration under the Nagoya Protocol^76^. Our partnership, which required significant time and investment from both parties since its initiation in 2011, is a first step towards an inclusive model of genomic science that we hope others will adopt and improve upon. While each partnership is unique, we provide a template genome research agreement (Supplementary Materials 20) that we hope will be useful to others.

## Discussion

The tuatara genome is among the highest quality genomes assembled for any reptile, and one of the largest amniote genomes thus far assembled. The tuatara has a genomic architecture unlike anything previously reported. Tuatara possess a unique amalgam of features that were previously viewed as characteristic of either the mammalian or reptilian lineages. Notable among these features are unusually high levels of repetitive sequences classically considered mammalian, many of which appear to have been recently active, and a level of genome methylation that is the highest of any amniote thus far investigated. We also found a mitochondrial genome (mtDNA) gene content at odds with published reports for this species that mistakenly omitted the ND5 gene^35^; this gene is indeed present and nested within a repeat rich region of the mtDNA.

Phylogenetic studies reveal new insights into the timing and speed of amniote evolution, including new evidence of punctuated genome evolution across this phylogeny. Contrary to past claims^66^ we find that the evolutionary rate for tuatara is not exceptionally fast, rather it is the slowest-evolving lepidosaurian reptile yet analysed.

Investigations of genomic innovations identified genetic candidates that may explain tuatara’s ultra-low active body temperature, extraordinary longevity, and apparent resistance to infectious disease. Further functional exploration will refine our understanding of these unusual facets of tuatara biology, while the tuatara genome itself will enable many future studies to explore the evolution of complex systems across the vertebrates in a more complete way than has previously been possible.

Our population genomic work supports the taxonomic distinctiveness of the Brothers Island tuatara, which has variously been described as *Sphenodon punctatus* or *Sphenodon guntheri*^72^ over the past three decades, but has recently been resynonomised^73^. Our findings show significant genetic differences among the populations, leading us to suggest that a separate conservation plan be re-established for each population.

Finally, this landmark genome will greatly aid future work on population differentiation, inbreeding, and local adaptation in this global icon, the last remaining species of the once globally dominant reptilian order Rhynchocephalia (Sphenodontia).

## Methods

### General methods

A full description of the Methods can be found in the Supplementary Materials.

### Sampling and sequencing

A blood sample was obtained from a large male tuatara from Lady Alice Island (35°53’24.4“S 174°43’38.2”E), New Zealand, with appropriate ethical and iwi permissions. Total genomic DNA and RNA were extracted and sequenced using the Illumina HiSeq 2000 and MiSeq sequencing platforms (Illumina, San Diego, CA, USA) supported by New Zealand Genomics Ltd (Supplementary Materials 1).

### Genome, transcriptome and epigenome

#### Genome assembly and scaffolding

Raw reads were *de novo* assembled using Allpaths-LG (version 49856)^77^. With a total input data of 5,741,034,516 reads for the paired-end libraries and 2,320,886,248 reads of the mate pair libraries, several setups were tested for the optimal proportion of input data. Based on assembly statistics an optimal setup was found using 85% of the fragment libraries and 100% of the jumping libraries. Gaps were closed using GapCloser^78^ and the assembly scaffolded with the transcriptome (see below) using L_RNA_Scaffolder^79^. We further scaffolded the assembly using Chicago libraries and HiRise^80^.

We assembled a *de novo* transcriptome as a reference for read mapping using total RNA derived from the blood sample of our reference male using a standard RNA-Seq pipeline^81^, and a collection of prior transcriptomic data collected from early stage embryos^82^. In total we had 131,580,633 new 100bp read pairs and 60,637,100 prior 50 bp read pairs^82^. These were assembled using Trinity v2.2.0^83, 84^ running default parameters. To reduce redundancy in the assembled transcripts, we used CD-HIT-EST v4.6.6^85^ and then assessed assembly quality and completeness before and after redundancy removal using BUSCO v1.22^7^ and Transrate v1.0.3^86^ (Supplementary Materials, section 1.4).

Low-coverage BS-seq was undertaken using a modified post-bisulfite adapter tagging (PBAT) method^87^ to explore global patterns of methylation in the tuatara genome for representatives of both sexes (12 males, 13 females) (Figure 1c, 3d). Briefly, purified DNA was subjected to bisulfite conversion using the EZ Methylation Direct Mag Prep kit (Zymo, D5044). Converted DNA was then subjected to single-ended sequencing (1x 150 bp) on an Illumina MiSeq. Raw reads were trimmed in two steps using TrimGalore v0.4.0 and trimmed reads were mapped using Bismark v0.14.3 with the --pbat option^87^. Global methylation levels for each species were calculated from Bismark reports by dividing methylated cytosine calls by total cytosine calls in both CG and non-CG contexts.

### Repeat and gene annotation

We used a combination of *ab initio* repeat identification in CARP/RepeatModeler/LTRharvest^15, 88, 89^, manual curation of specific newly identified repeats, and homology to repeat databases to investigate the repeat content of the tuatara genome (Supplementary Materials 1.6). From these three complementary repeat identification approaches, the CARP results were in-depth annotated for LINEs and segmental duplications (Supplementary Materials 4), the RepeatModeler results were in-depth annotated for SINEs and DNA transposons (Supplementary Materials 5), and the LTRharvest results were in-depth annotated for LTR retrotransposons (Supplementary Materials 6).

For the gene annotation we used RepeatMasker (v4.0.3) along with our partially curated RepeatModeler library plus Repbase sauropsid repeat database to mask transposable elements in the genome sequence prior to the gene annotation. We did not mask simple repeats at this point to allow for more efficient mapping during the homology based step in the annotation process. Simple repeats were later soft-masked and protein-coding genes predicted using MAKER2^90^. We used anole lizard (*A. carolinensis*, version AnoCar2.0), python (*P. bivittatu*s, version bivittatus-5.0.2,) and RefSeq (www.ncbi.nlm.nih.gov/refseq) as protein homology evidence, which we integrated with *ab initio* gene prediction methods including BLASTX, SNAP^91^ and Augustus^92^. Non-coding RNAs were annotated using Rfam covariance models (v13.0)^93^ (Supplementary Materials 7).

### Ortholog calling

We performed a phylogenetic analysis to infer orthology relationships between the tuatara and 25 other species (Supplementary Materials Tables 2.1 and 2.2) using the Ensembl GeneTree method^94^. Multiple-sequence alignments, phylogenetic trees and homology relationships were extracted in various formats (<u>10.5281/zenodo.2542570</u>). We also calculated the Gene Order Conservation (GOC) score^95^, which uses local synteny information around a pair of orthologous genes to compute how much the gene order is conserved. For each of these species, we chose the paralogue with the best GOC score and sequence similarity, which resulted in a total set of 3,168 clusters of orthologs (Supplementary Materials Table 2.3).

### Gene tree reconstructions and substitution rate estimation

We constructed phylogenies using just fourfold degenerate site data derived from whole genome alignments for 27 tetrapods, analysed as a single partition in RAxML v8.2.3^96^. Using the topology and branch lengths obtained from the best maximum likelihood phylogeny, we estimated absolute rates of molecular evolution in terms of substitution per site per million years and estimated the divergence times of amniotes via the semiparametric penalized likelihood (PL) method^97^ with the program r8s v1.8^98^ (Supplementary Materials 14.5).

We also generated gene trees based on 245 single-copy orthologs found between all species using a maximum likelihood based multi-gene approach (Supplementary Materials 15). Sequences were aligned using the codon-based aligner Prank^99^. The FASTA format alignments were then converted to Phylip using the catfasta2phyml.pl script (https://github.com/nylander/catfasta2phyml). Next, we used the individual exon Phylip files for gene tree reconstruction using RaxML^96^ using a GTR + G model^96^. Subsequently, we binned all gene trees to reconstruct a species tree and carried out bootstrapping using Astral^100^. Astral applies binning of gene trees with similar topologies, based on an incompatibility graph between gene trees. Subsequently, it chooses the most likely species tree under the multi-species coalescent model (Supplementary Materials Figure 15.1).

### Divergence times and tests of punctuated evolution

We inferred time-calibrated phylogenies with BEAST v2.4.8^101^ using the CIPRES Science Gateway^102^ to explore divergence times (Supplementary Materials 16.1). Subsequently, we used Bayesian phylogenetic generalized least squares to regress the total phylogenetic path length (of four-fold degenerate sites) on the net number of speciation events (nodes in a phylogenetic tree) as a test for punctuated evolution^103^. (Supplementary Materials 16.2).

### Analysis of genomic innovations

We explored the genomic organisation and sequence evolution of genes associated with immunity, vision, smell, thermoregulation, longevity and sex determination (Supplementary Materials 8–13). Tests of selection were undertaken across multiple genes, including those linked to metabolism, vision and sex determination (Supplementary Materials 17) using multispecies alignments and PAML^104^.

### Population genomics

Demographic history was inferred from the diploid sequence of our tuatara genome using a pairwise sequential Markovian coalescent (PSMC) method^105^ as detailed in Supplementary Materials 18. We also sampled 10 animals from each of three populations that span the main axes of genetic diversity in tuatara (Supplementary Materials Table 19.1) and used a modified genotype-by-sequencing approach^106^ to obtain 52,171 single nucleotide variants (SNVs) that we used for population genomic analysis, investigations of loci associated with sexual phenotype and estimates of genetic load (Supplementary Materials 19).

### Reporting summary

Further information on experimental design is available in the Nature Research Reporting Summary linked to this article.

### Permits and ethics

Samples were collected under Victoria University of Wellington Animal Ethics approvals 2006R12; 2009R12; 2012R33; 22347 and held and used under permits 45462-DOA and 32037-RES 32037-RES issued by the New Zealand Department of Conservation.

### Data availability

The Tuatara Genome Consortium Project Whole Genome Shotgun and genome assembly are registered under the umbrella BioProject PRJNA418887 and BioSample SAMN08038466. Transcriptome read data are submitted under SRR7084910 (whole blood) together with prior data SRR485948. The transcriptome assembly is submitted to GenBank with ID GGNQ00000000.1. Illumina short-read and nanopore long read sequence are in SRAs associated with PRJNA445603. The assembly (GCA_003113815.1) described in this paper is version QEPC00000000.1 and consists of sequences QEPC01000001-QEPC01016536. Maker gene predictions are available from Zenodo, DOI: <u>10.5281/zenodo.1489353</u>. The repeat library database developed for tuatara is available from Zenodo, DOI: <u>10.5281/zenodo.2585367</u>

## Acknowledgements

N.J.G. acknowledges the support of Ngatiwai iwi, Allan Wilson Centre, University of Otago, New Zealand Department of Conservation, New Zealand Genomics Limited, Illumina. J.I.A. was supported by CONICYT National Doctoral Scholarship N°21130515. Ensembl annotation was supported by the Wellcome Trust (WT108749/Z/15/Z) and the European Molecular Biology Laboratory.

We also acknowledge Ngāti Kaota, Te Ātiawa and Ngāti Manuhiri for granting permission to reuse tuatara samples obtained from Stephens Island (Takapourewa), Little Barrier Island (Hauturu) and North Brother Island, respectively. We gratefully acknowledge all the people involved in obtaining and curating the samples held in the Victoria University tuatara collection.

Finally we thank Aleksey Zimin, Daniela Puiu, Guillaume Marcais, James Yorke and Ross Crowhurst for help and discussions about genome assembly, Ian Fiddes, Joel Armstrong and Benedict Paten for help with comparative genome alignments and annotation, the National eScience Infrastructure (NeSI) and Swedish National Infrastructure for Computing (SNIC) through the Uppsala Multidisciplinary Center for Advanced Computational Science (UPPMAX) for computational support, Robbie McPhee for help with figures, and Tamsin Braisher for manuscript coordination and editing.

## Contributions

N.J.G. designed the original concept and scientific objectives and oversaw the project and analyses. N.J.G., L.M., N.N., H.T., O.R., S.V.E., C.S. contributed samples or assisted in sample preparation and permitting. N.J.G., K.M.R., S.P., M.T., D.W., R.M., D.L.A., A.S., T.B., J.H.G., C.O., P.P.G., M.M. M.P., K.B., F.J.M., P.F., B.P., L.K., P.M., T.R.B., M.W., Y.C., H.M., R.K.S., M.J., R.N., J.I.A., N.V., T.A.H., J.R., V.P., C.R.P., V.M.W., L.Z., R.D.K., J.M.R., V.V.Z., Z.W., D.S., M.M., R.G., s.M.R., V.G.T., N.D., H.A., J.M.P., D.G.M., V.L.G., C.G.B., D.P.D., S.P., S.L.S. planned and carried out experiments or analyses. N.J.G., K.M.R., S.P., M.T., D.W., R.M., D.L.A., A.S., T.B., J.H.G., C.O., P.P.G., M.M. M.P., K.B., F.J.M., B.P., L.K., P.M., T.R.B., M.W., Y.C., H.M., R.K.S., M.J., R.N., J.I.A., N.V., T.A.H., J.R., V.P., C.R.P., V.M.W., L.Z., R.D.K., J.M.R., V.V.Z., Z.W., D.S., M.M., R.G., S.M.R., V.G.T., N.D., H.T., H.A., J.M.P., D.G.M., V.L.G., C.G.B., D.P.D., S.P., S.L.S. contributed to the interpretation and presentation of results in the main manuscript and supplementary documents. N.J.G. wrote the first draft of the manuscript with input from all other authors.

---

**Supplementary Figure 1.**
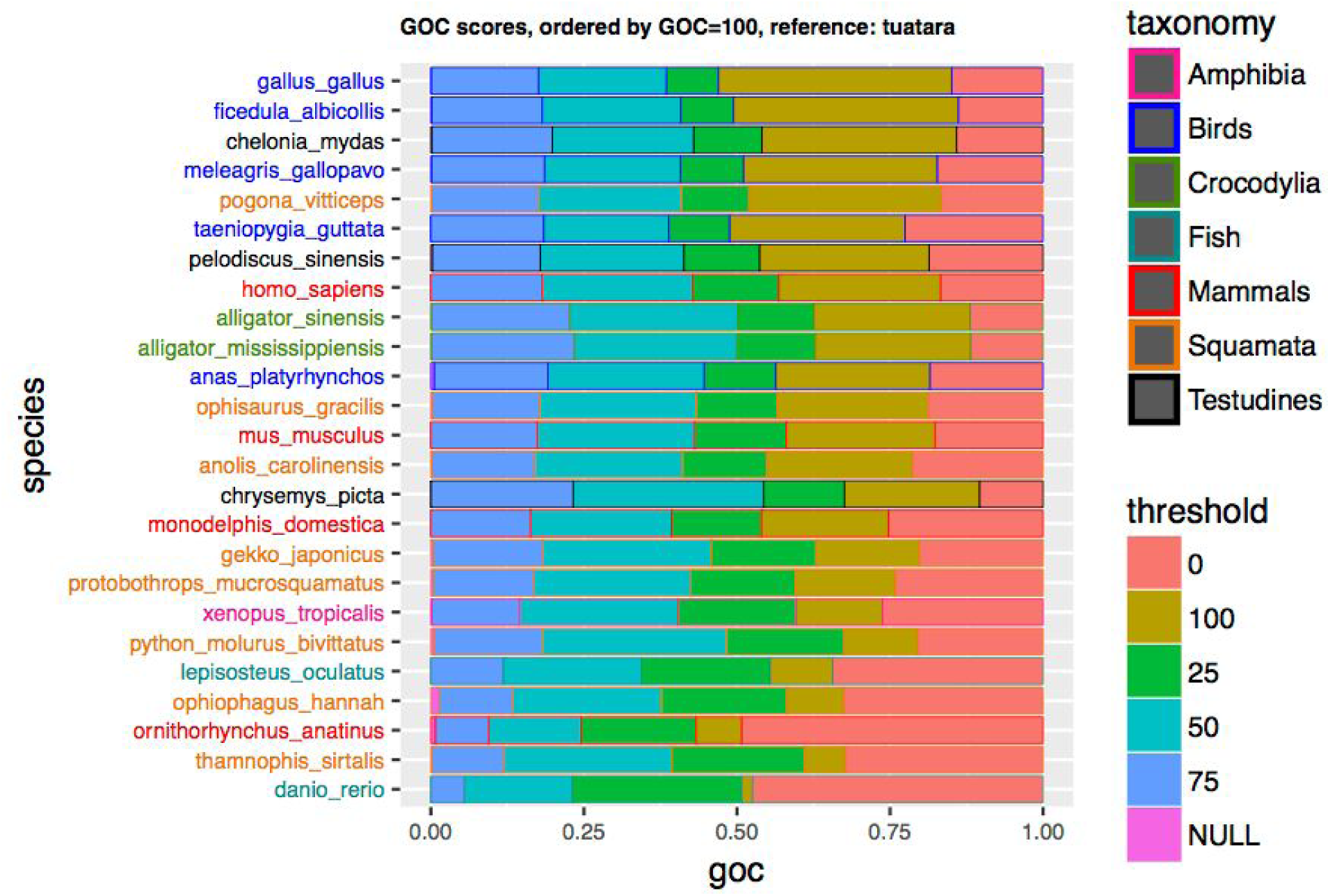
Gene Order Conservation score distribution using the tuatara as a reference. Species are ordered by the proportion of top-scoring orthologs.

**Supplementary Figure 2.**
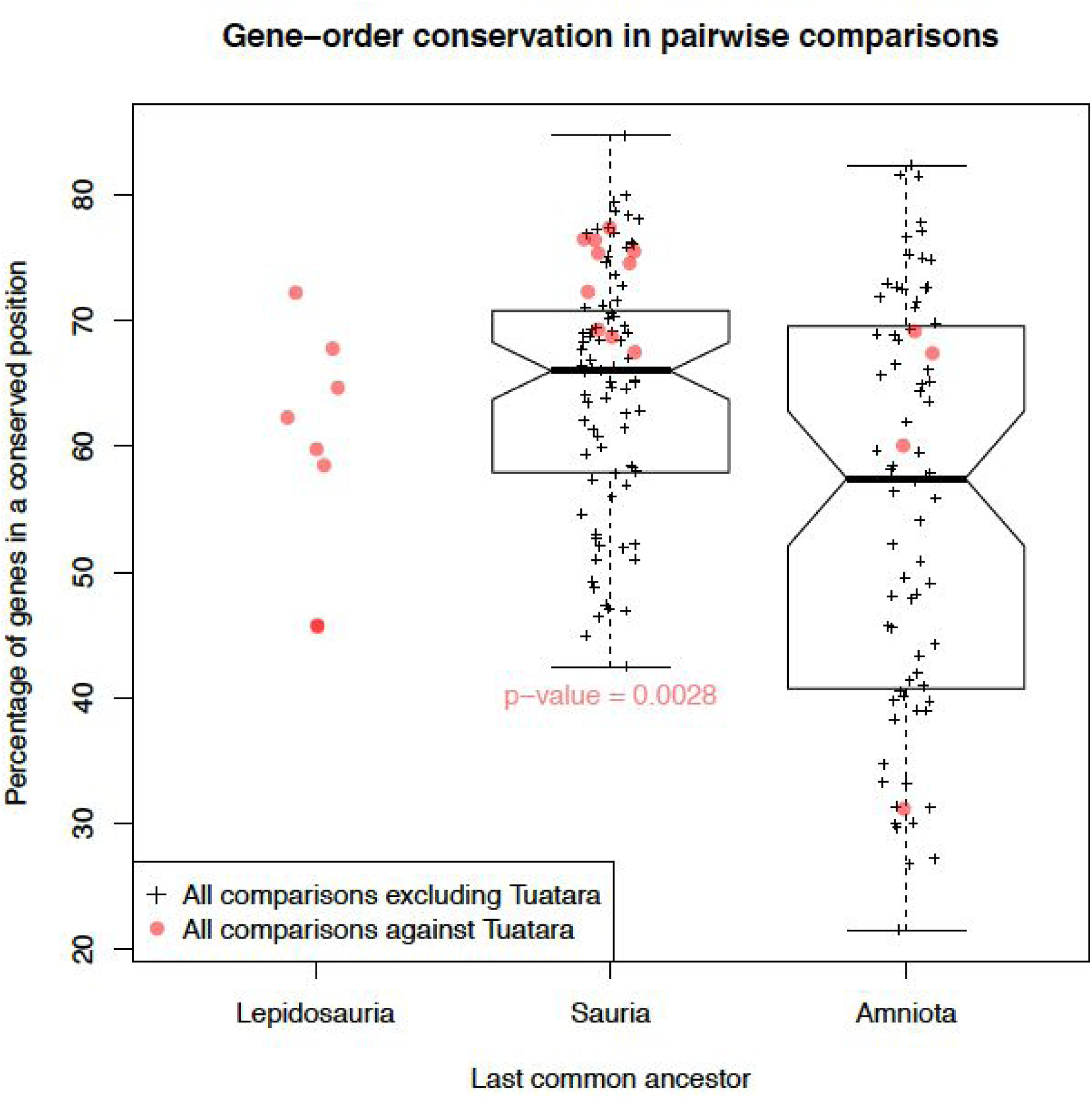
Gene-order conservation versus divergence time. For each taxonomic grouping, we analyzed the percentage of genes that are found in a conserved position across all pairs of genomes. Pairwise comparisons involving Tuatara are shown in plain red circles, whilst comparisons that do not involve Tuatara are black (boxplot and “+” signs). The conservation of gene-order between Tuatara, birds and turtles is significantly higher (p-value = 2.8e-3) than that observed between squamates birds and turtles. As Tuatara is the only remaining rhynchocephalian, there is no control distribution for the Lepidosauria ancestor.

**Supplementary Figure 3.**
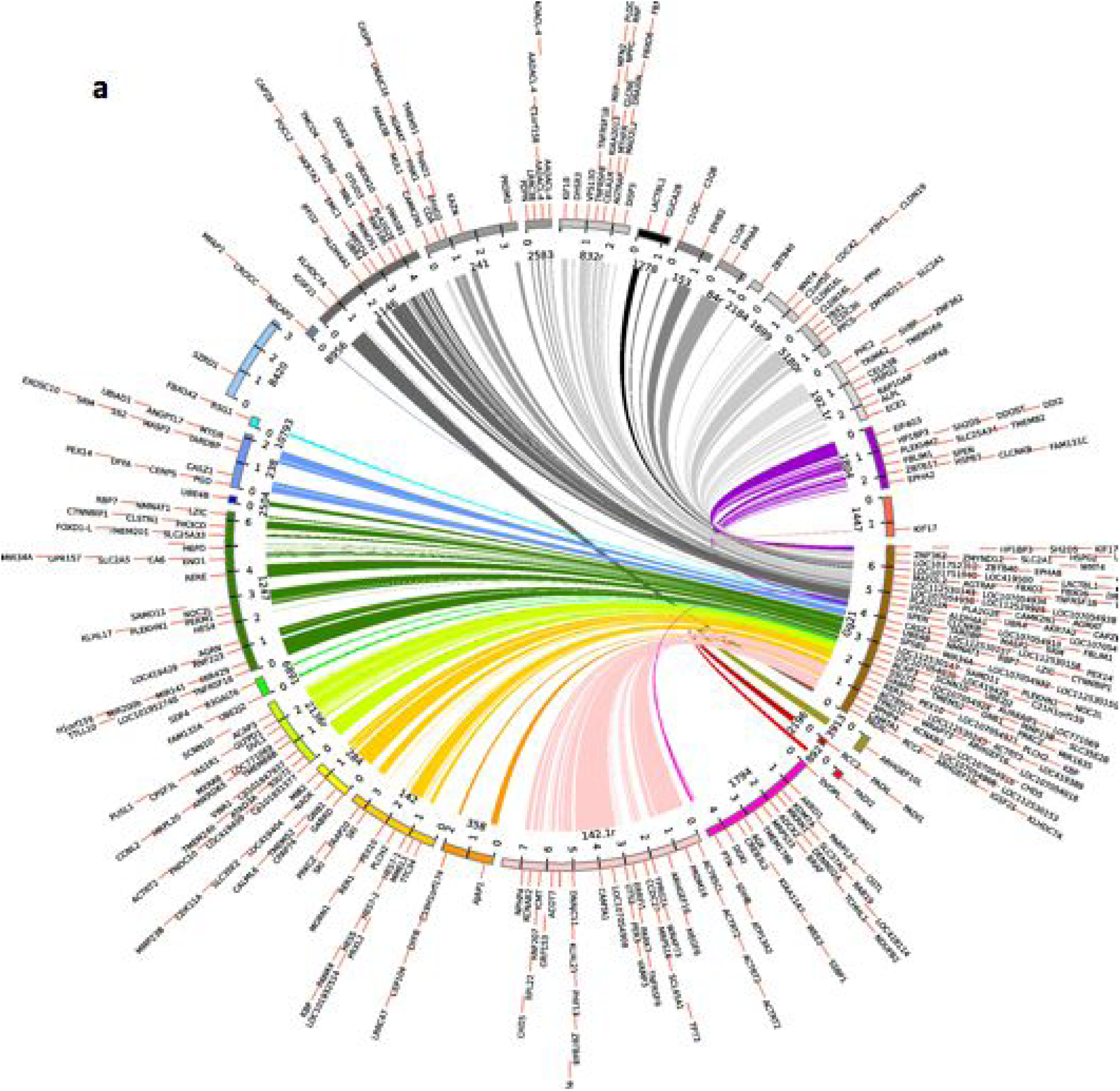
Synteny between *G. gallus* chromosome 21 and tuatara contigs. Circos plot highlighting the synteny between chicken chromosome 21 (assembly GRCg6a) and multiple tuatara contigs. Gene names shown derive from the chicken assembly.

**Supplementary Figure 4.**
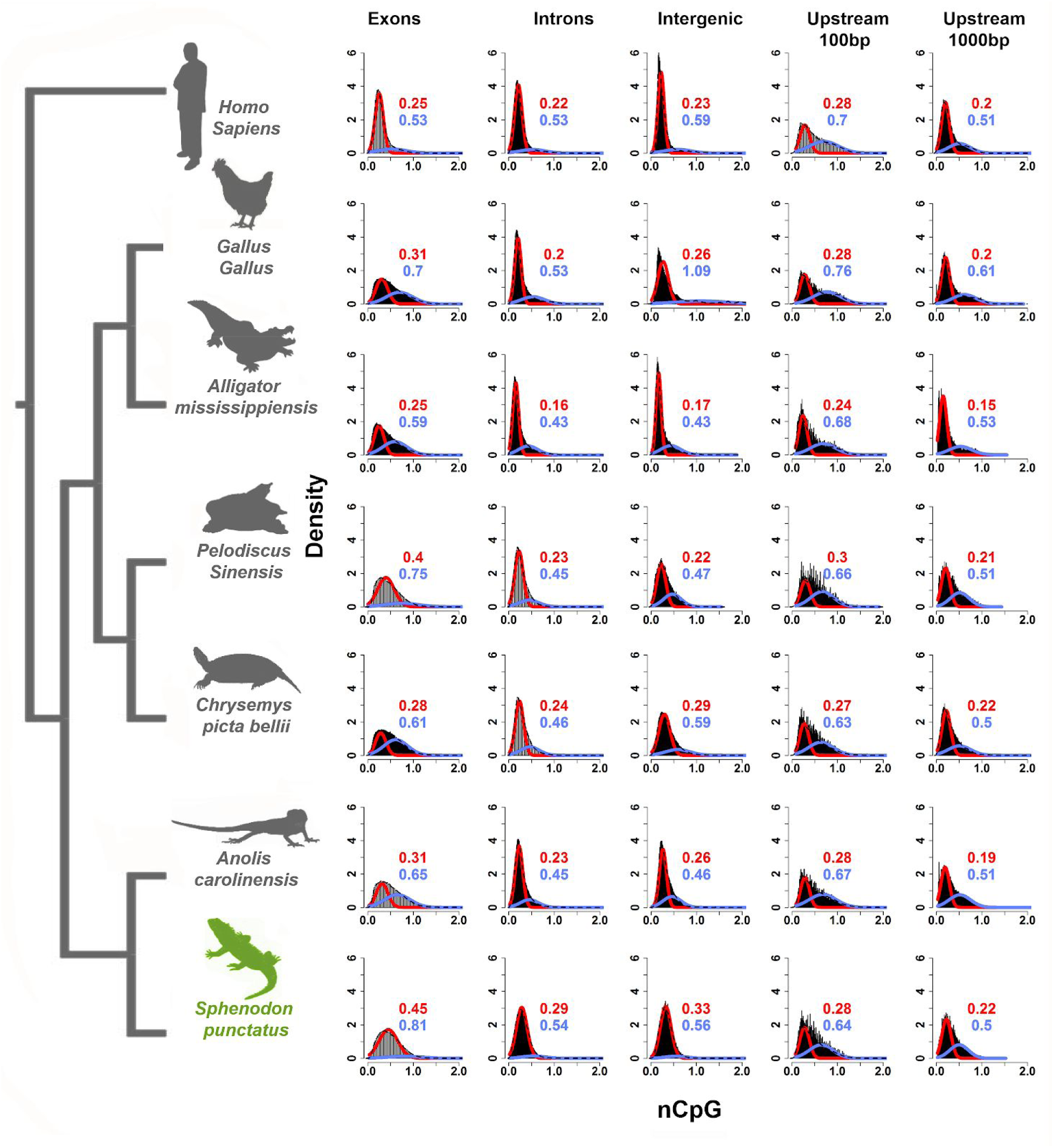
Normalized CpG distributions (nCpG) for tuatara and other vertebrates.

**Supplementary Figure 5.**
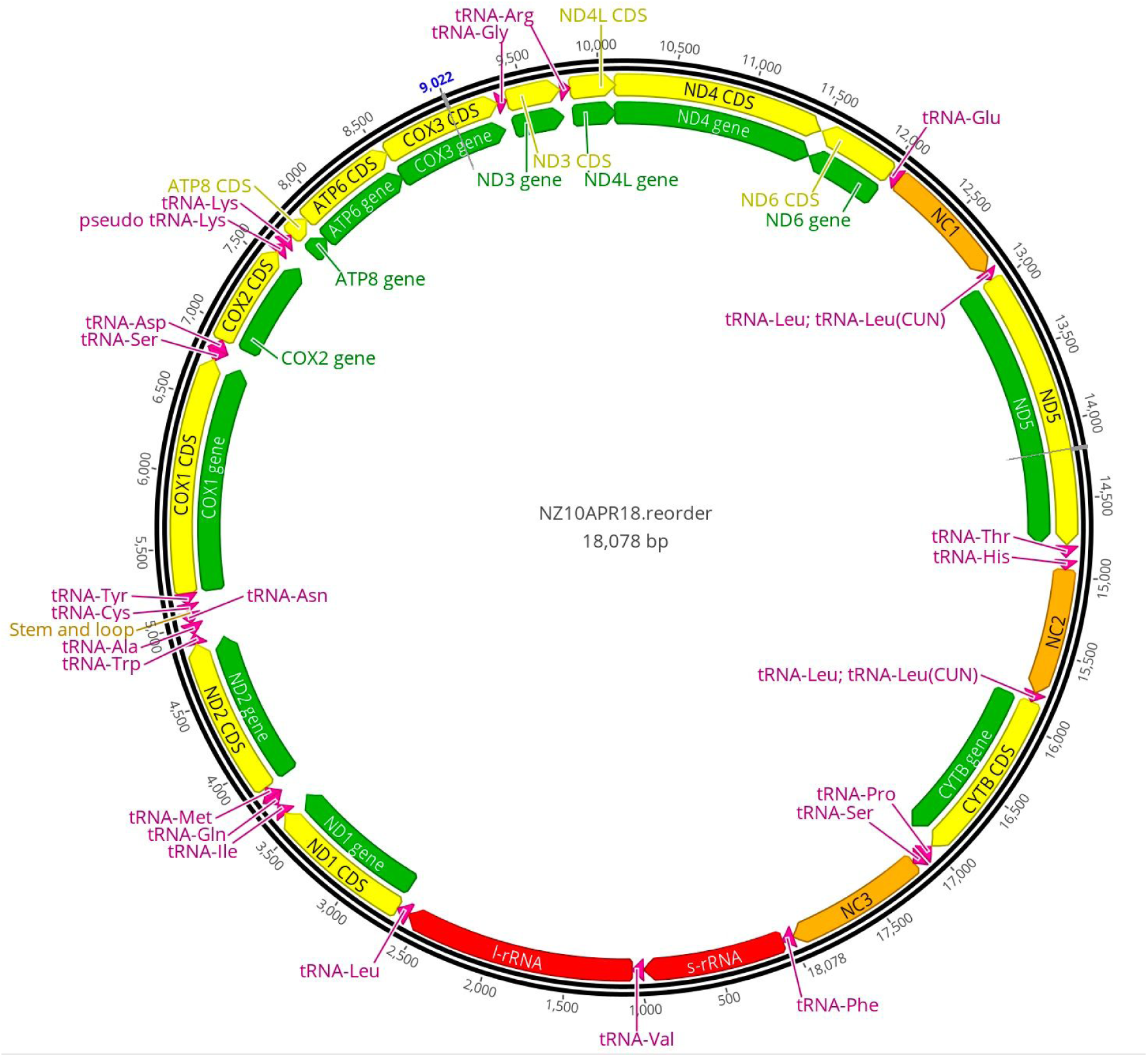
The mitochondrial (mt) genome in the Lady Alice Island reference animal is 18,078 bp, containing 13 protein coding, 2 rRNA, and 22 tRNA genes, standard among animals contradicting prior reports that 3 genes: ND5, tRNA^Thr^, tRNA^His^ were absent. Three non-coding areas with Control Region (heavy strand replication origin, O_H_) features (NC1–3), and two copies of tRNA^Leu(CUN)^ adjacent to NC1 and NC2 possess identical or near identical sequence, possibly as a result of concerted evolution. The gene and structure order is: tRNA^Phe^, 12S rRNA, tRNA^Val^, 16S rRNA, tRNA^Leu(UUR)^, ND1, tRNA^Ile^, tRNA^Gln^, tRNA^Met^, ND2, tRNA^Trp^, tRNA^Ala^, tRNA^Asn^, O_L_-like structure, tRNA^Cys^, tRNA^Tyr^, COI, tRNA^Ser(UCN)^, tRNA^Asp^, COII, pseudo-tRNA^Lys^, tRNA^Lys^, ATP8, ATP6, COIII, tRNA^Gly^, ND3, tRNA^Arg^, ND4L, ND4, ND6, tRNA^Glu^, NC1, tRNA^Leu(CUN)^ copy one, ND5, tRNA^Thr^, tRNA^His^, NC2, tRNA^Leu(CUN)^ copy two, Cytb, tRNA^Pro^, tRNA^Ser(AGY)^, and NC3.

**Supplementary Figure 6.**
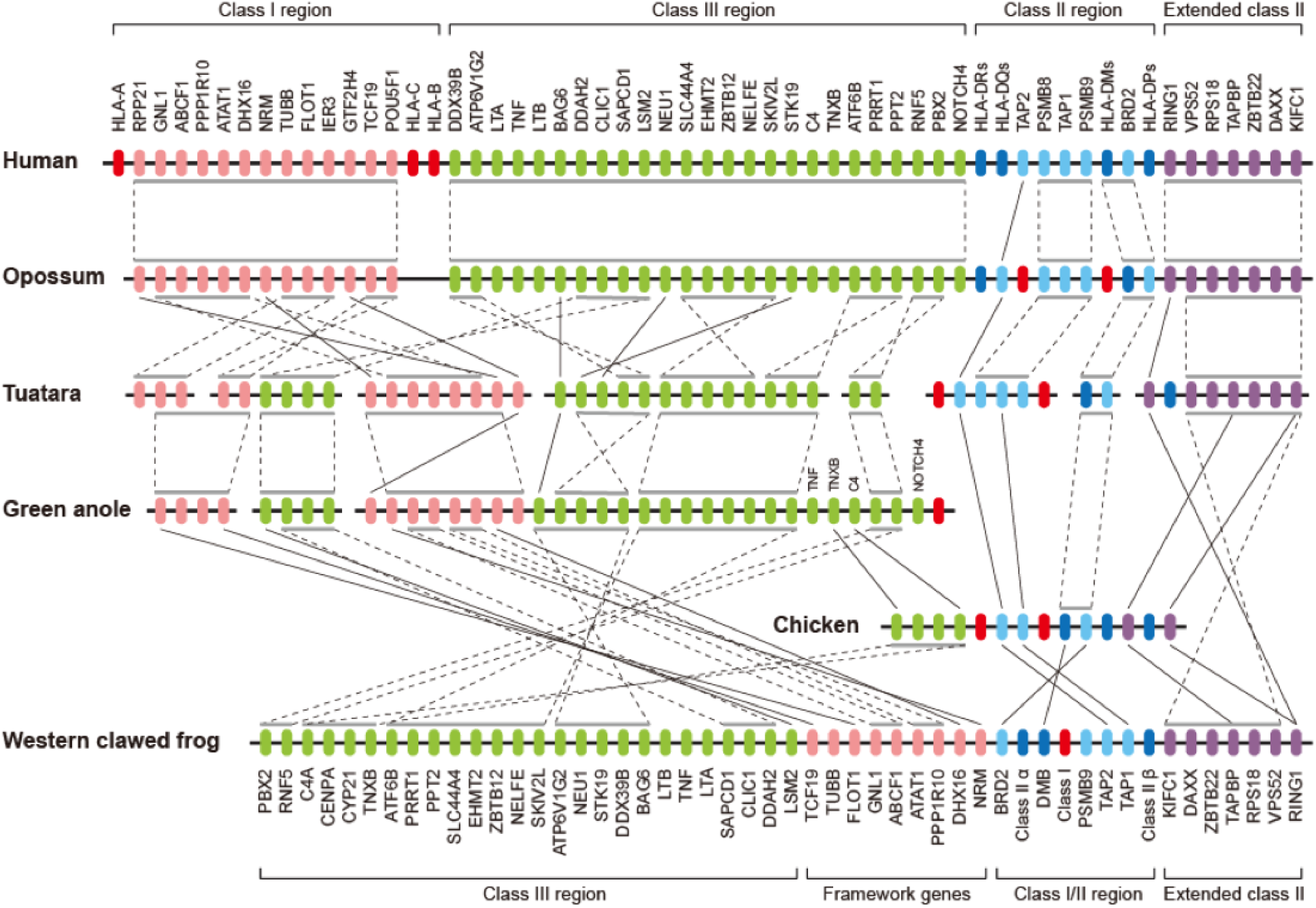
Comparative analysis of the MHC core region. Only genes that were annotated in the tuatara genome were included in the analysis. Orthologs between species are connected by a solid line; the grey bars above/below genes indicate syntenic blocks and are linked by dashed lines between species. Anolis class I/II and extended class II regions are not shown due to the high degree of genome assembly fragmentation in these regions. Color legend: red – class I genes, pink – class I region framework genes, green – class III genes, dark blue – class II genes, light blue – class II region framework genes, purple – extended class II genes.

**Supplementary Figure 7.**
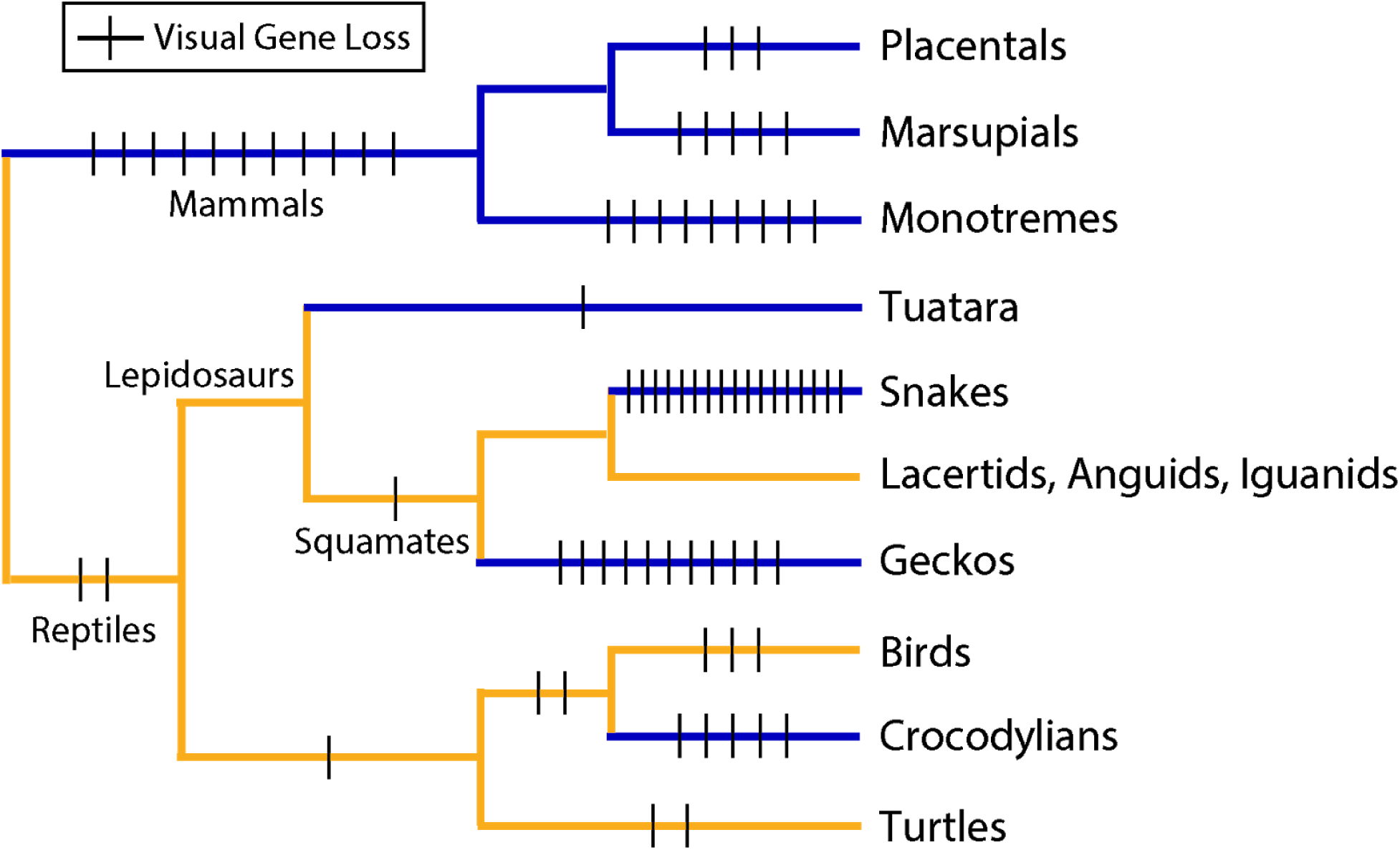
Phylogenetic tree of reptiles depicting inferred visual gene losses. Lineages are coloured based on a rough approximation of their ancestral activity pattern (blue, nocturnal; yellow, diurnal). Note that that tuatara lineage has experienced some of the lowest rates of gene loss despite a nocturnal ancestry, which in other lineages is associated with increased gene loss.

**Supplementary Figure 8.**
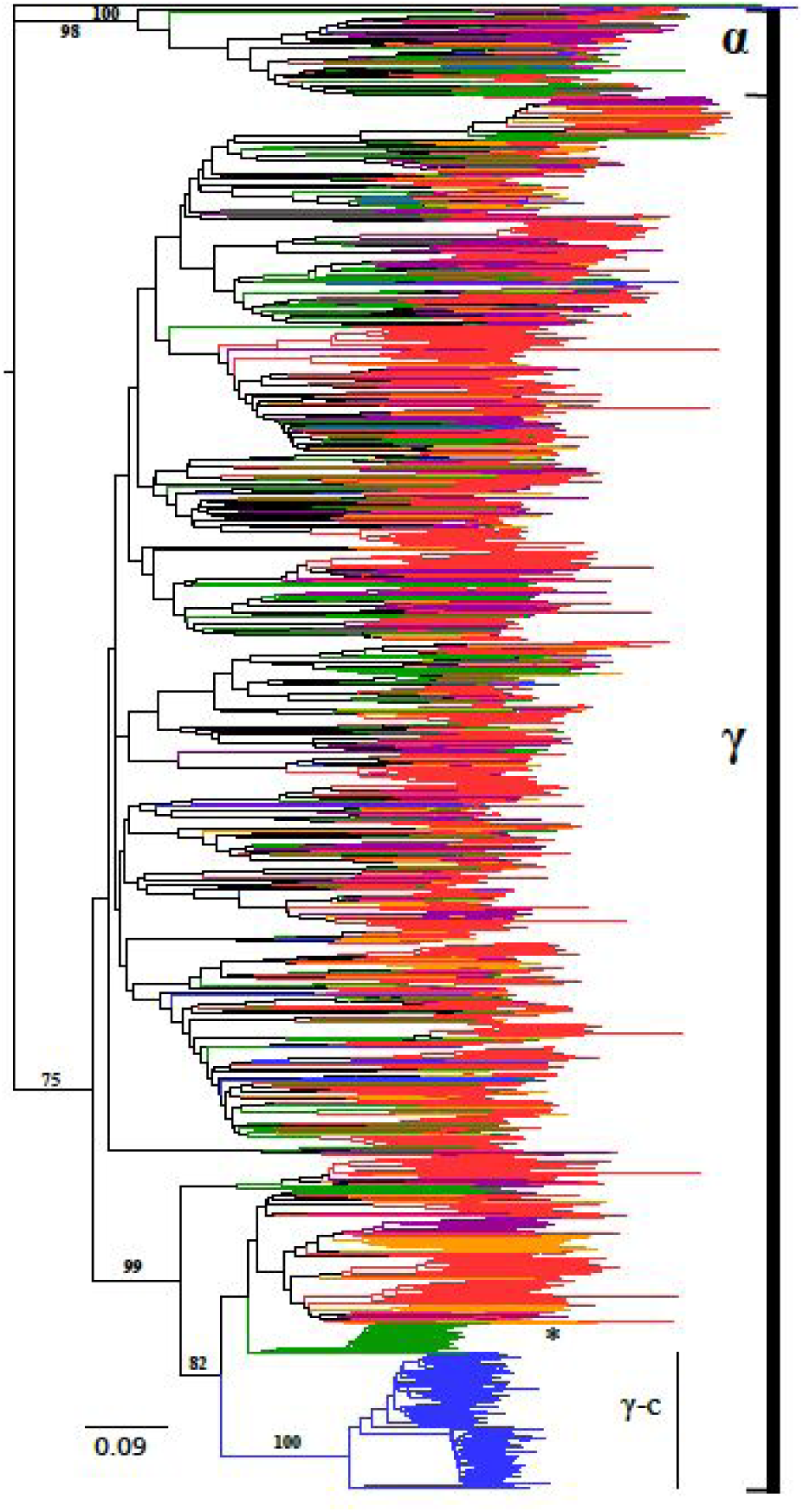
The evolutionary history of terrestrial Sauropsid ORs was inferred using the Neighbor-joining method. The unrooted tree contains 3213 amino acid sequences. Branches are coloured according to the following categories: Green – tuatara, Blue – birds (*Gallus gallus*, *Taeniopygia guttata*), Red – snakes (*Notechis scutatus*, *Ophiophagus Hannah, Protobothrops mucrosquamatus*, *Pseudonaja textilis*, *Python bivittatus* and *Thamnophis sirtalis*), Orange – lizards (*Anolis carolinesis, Pogona vitticeps*) and Purple – gecko (*Gekko japonicas*). Bootstrap support values above 75% (1000 replicates) are indicated for major branch splits relating to the different OR groups and branches leading to the species-specific OR expansions in birds (group γ-c) and tuatara (*).

**Supplementary Figure 9.**
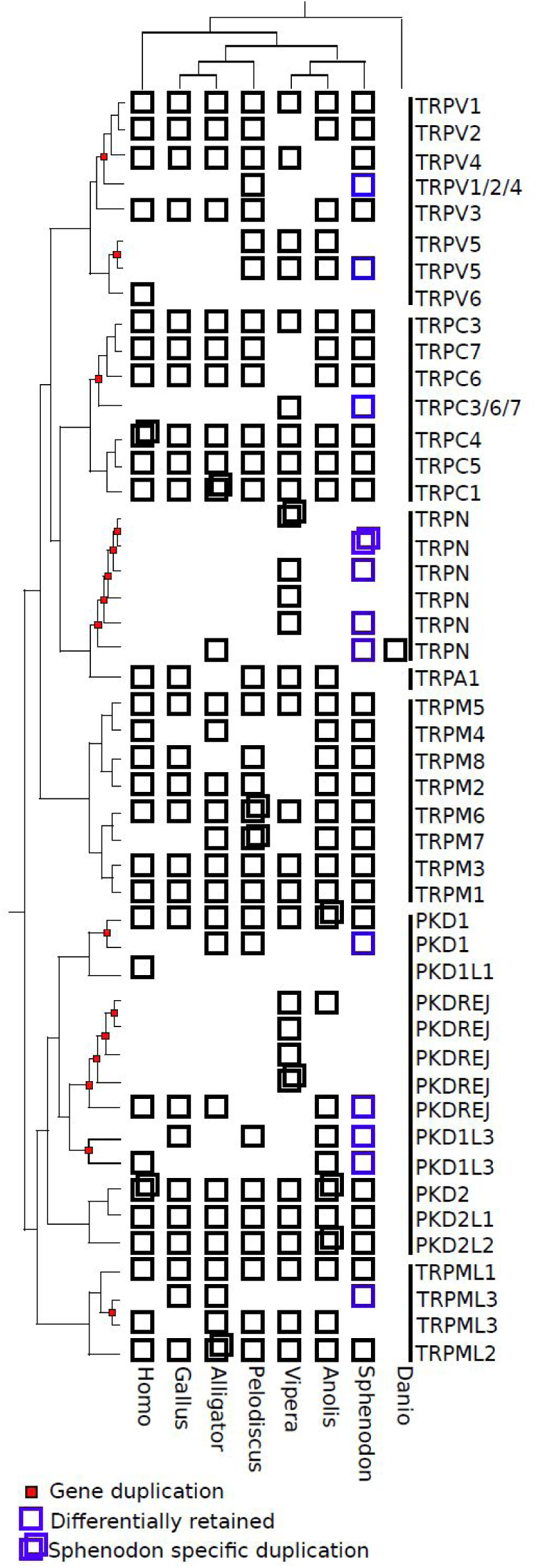
The repertoire of TRP genes identified in tuatara (*Sphenodon punctatus)* and other six vertebrate species: lizard (*Anolis carolinensis*), viper (*Vipera berus*), turtle (*Pelodiscus sinensis*), alligator (*Alligator mississippiensis*), chicken (*Gallus gallus*), and human (*Homo sapiens*). Small red squares on nodes indicate gene duplications, blue boxes indicate gene differential retentions and duplicated boxes indicate species-specific recent duplicates. Empty spaces are differential losses.

**Supplementary Figure 10.**
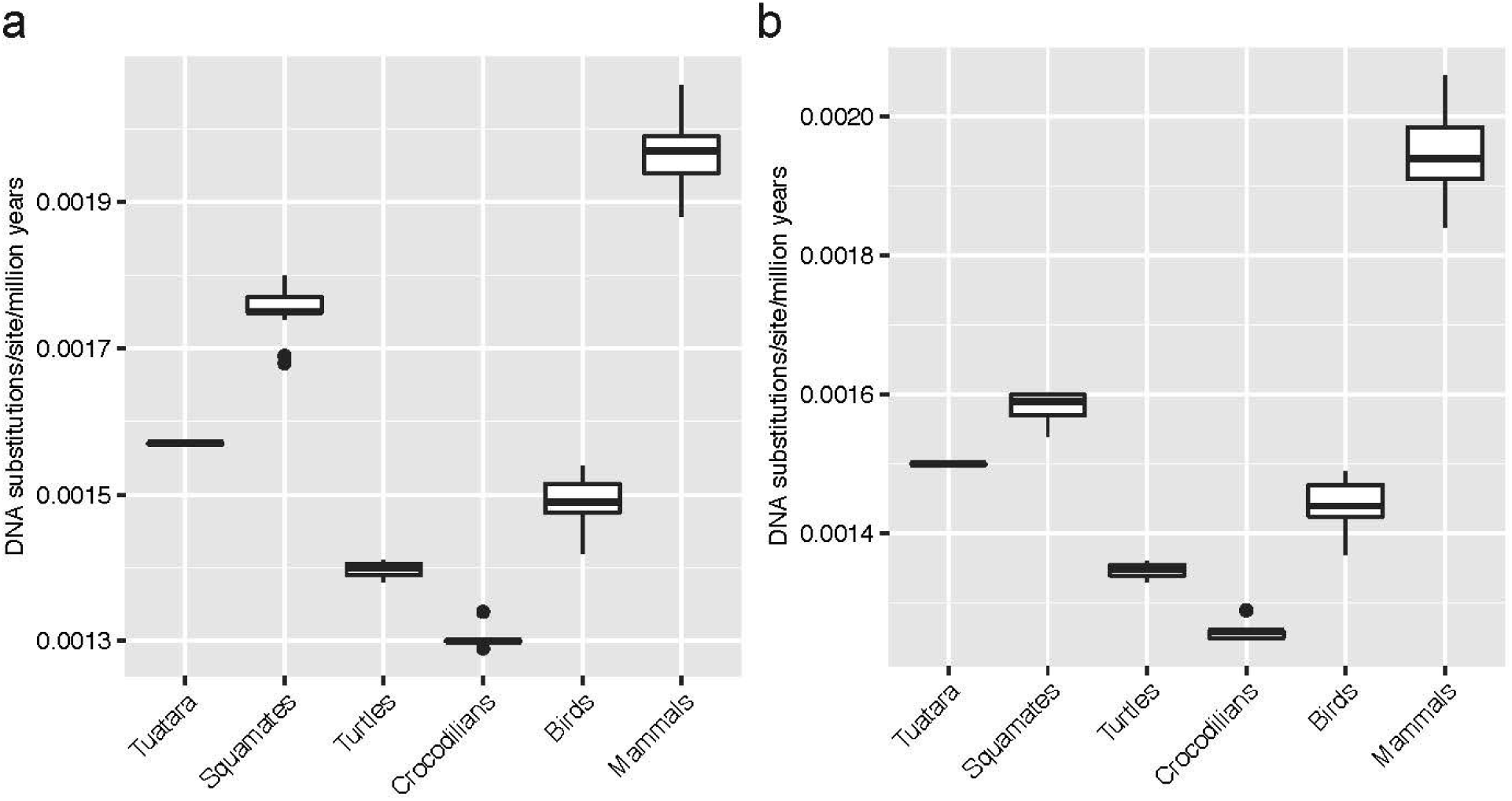
Estimated DNA substitution rates of amniote clades based on fourfold degenerate sites. Boxplots showing the distribution of estimated substitution rates by clade using semiparametric penalized likelihood in r8s with (a) fossil constraints from Benton et al. (Benton et al. 2015) and Head (J. J. Head 2015) and (b) median TMRCA estimates from timetree.org.

**Supplementary Figure 11.**
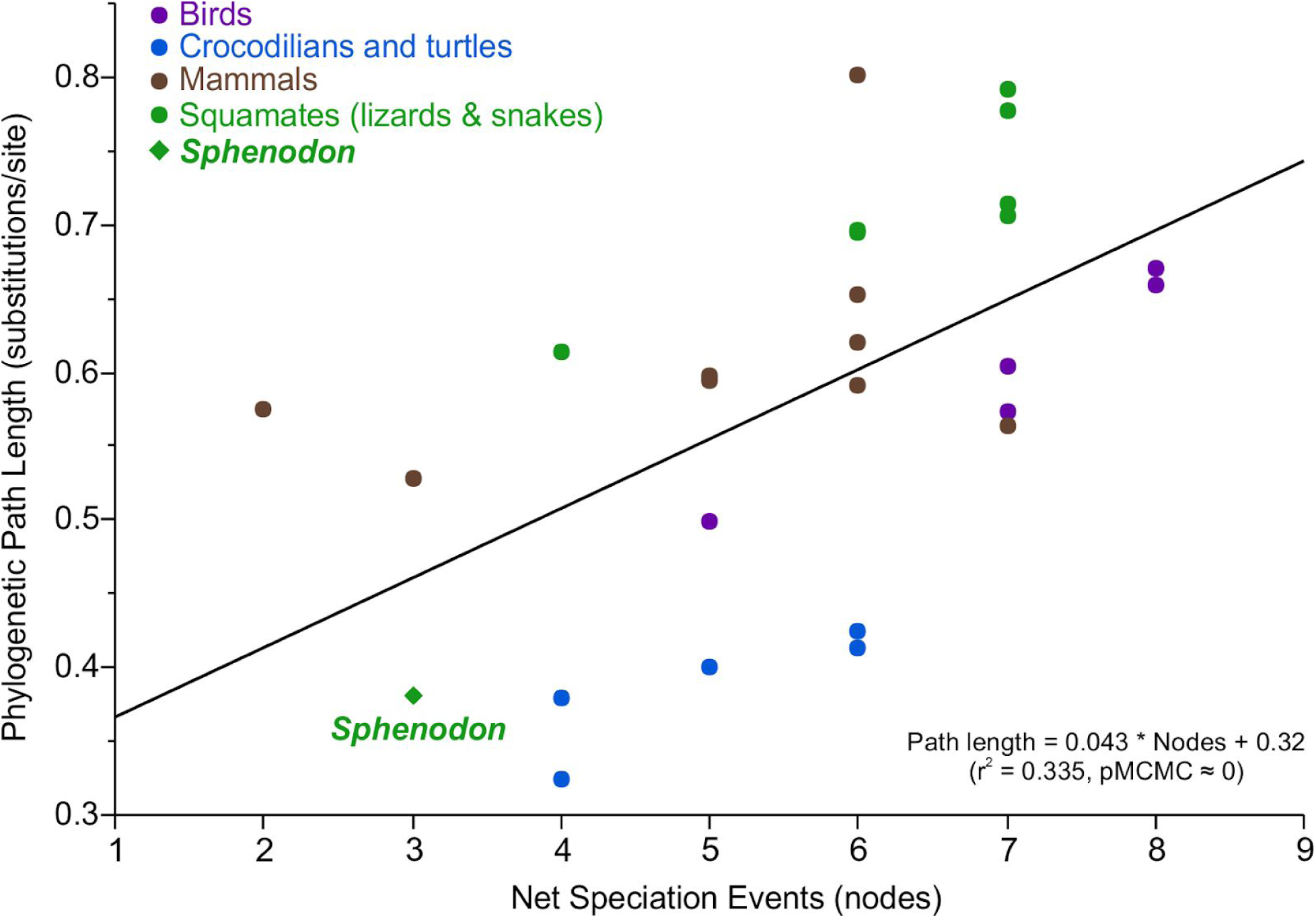
Testing for evidence of punctuated evolution. The process of punctuated genome evolution predicts that the amount of evolution in a given species’ genome should correlate with the net number of speciation events. We used Bayesian phylogenetic generalized least squares to regress the total phylogenetic path length (of four-fold degenerate sites) on the net number of speciation events (nodes in a phylogenetic tree). We find strong evidence for punctuated evolution, which accounts for 33.5% (r^2^; 95% credible interval = 0.34 to 0.38) of deviation from the molecular clock at four-fold degenerate sites.

**Supplementary Table 1.**
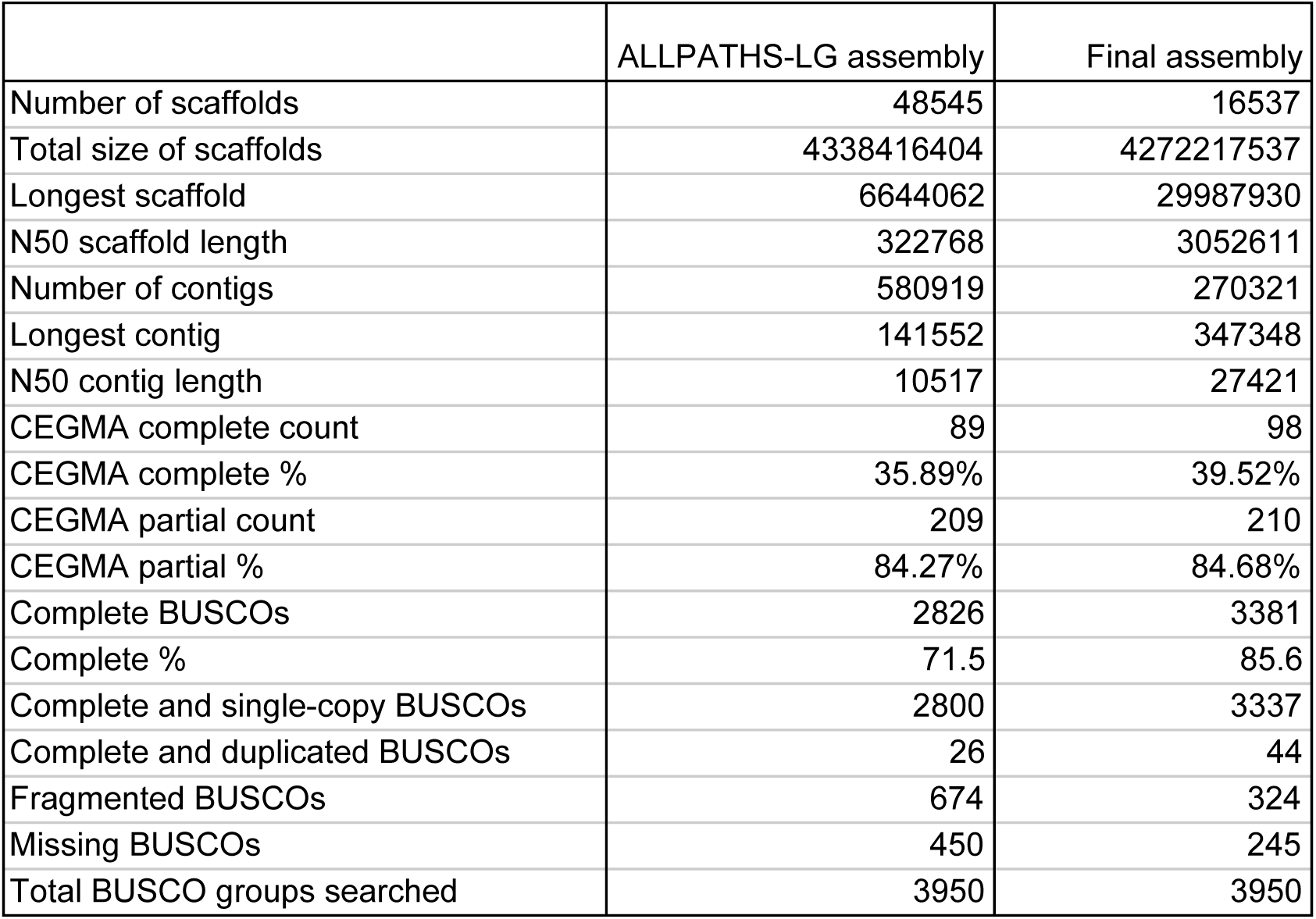
Assembly statistics and quality metrics for the tuatara genome.

|  | ALLPATHS-LG assembly | Final assembly |
| --- | --- | --- |
| Number of scaffolds | 48545 | 16537 |
| Total size of scaffolds | 4338416404 | 4272217537 |
| Longest scaffold | 6644062 | 29987930 |
| N50 scaffold length | 322768 | 3052611 |
| Number of contigs | 580919 | 270321 |
| Longest contig | 141552 | 347348 |
| N50 contig length | 10517 | 27421 |
| CEGMA complete count | 89 | 98 |
| CEGMA complete % | 35.89% | 39.52% |
| CEGMA partial count | 209 | 210 |
| CEGMA partial % | 84.27% | 84.68% |
| Complete BUSCOs | 2826 | 3381 |
| Complete % | 71.5 | 85.6 |
| Complete and single-copy BUSCOs | 2800 | 3337 |
| Complete and duplicated BUSCOs | 26 | 44 |
| Fragmented BUSCOs | 674 | 324 |
| Missing BUSCOs | 450 | 245 |
| Total BUSCO groups searched | 3950 | 3950 |

**Supplementary Table 2:**
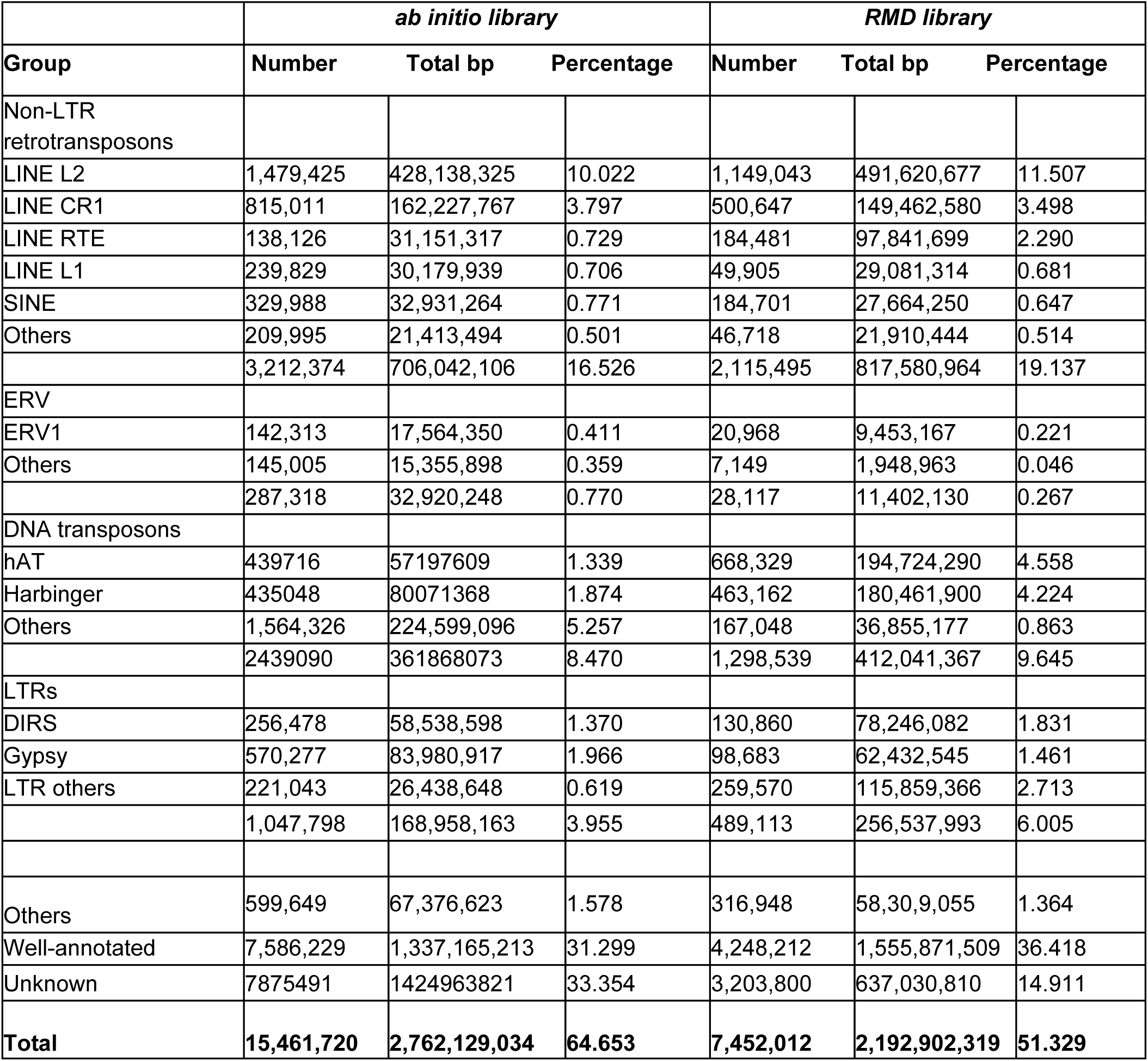
Copy number and fraction of tuatara genome covered by.

**Supplementary Table 3.**
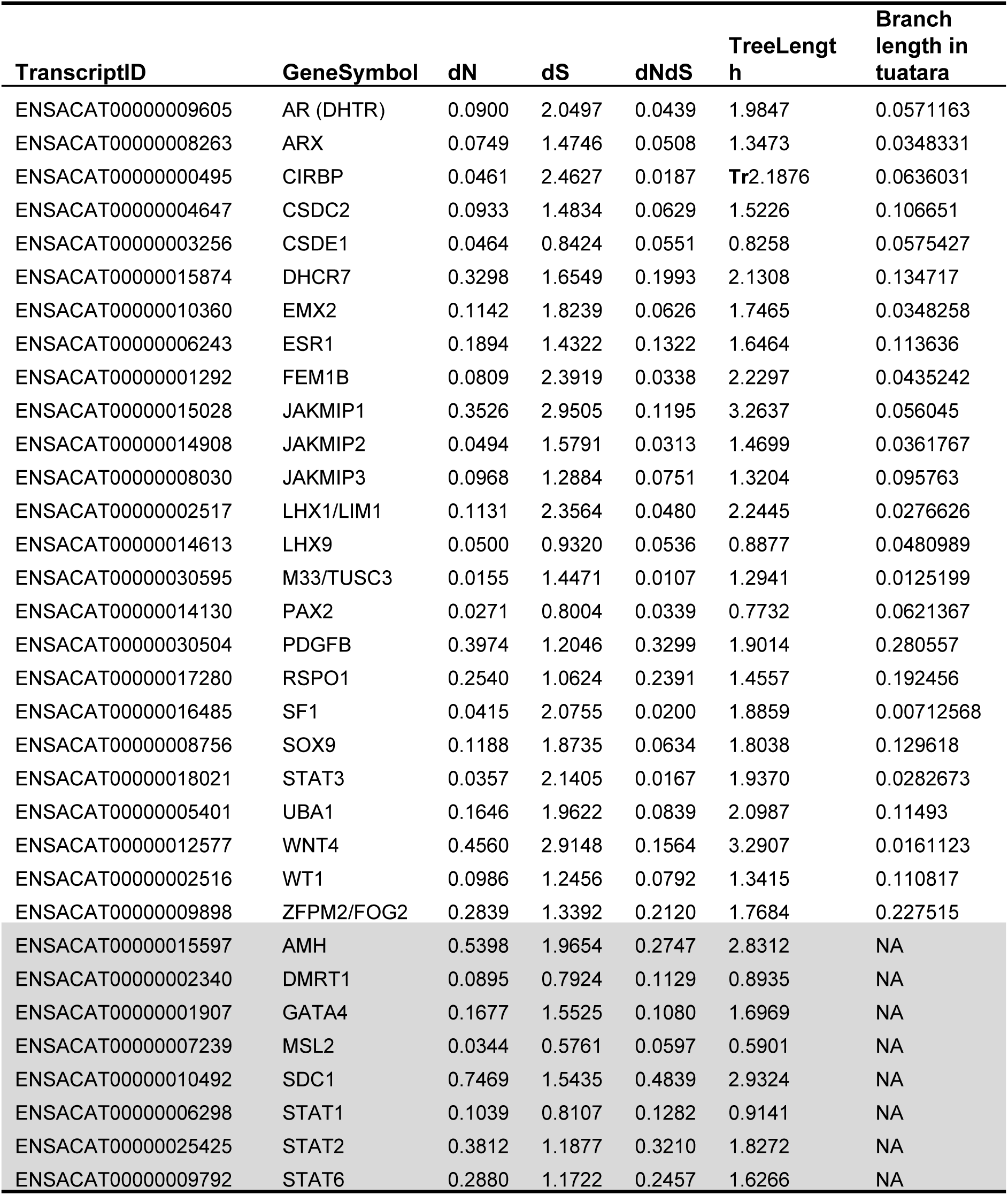
Vertebrate sex-determining genes found in the tuatara genome and their substitution rates. We computed substitution rates, d_N_, d_S_, and d_N_/d_S_ for genes previously identified as playing a primary role in sex determination (Supplementary materials 16). For eight sex-determining genes we did not identify homologous sequence in the tuatara in the multiple species alignment (gray). This does not mean that they don’t exist in the tuatara genome, but that they were not found in the current version of the multiple alignments, perhaps because of high divergence.

**Supplementary Table 4.**
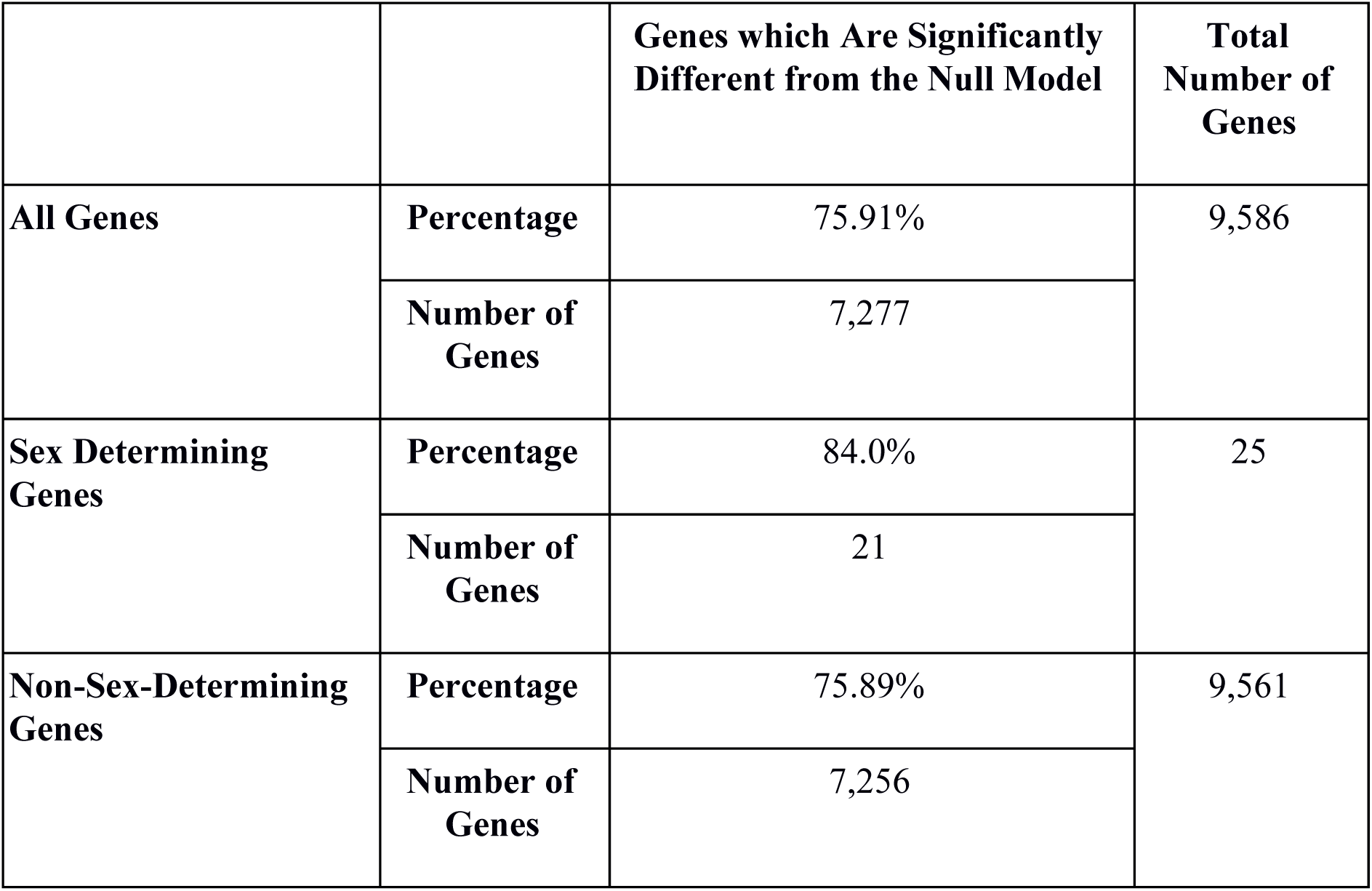
Tuatara–squamate gene tree evolution compared between all genes and sex determining genes. A log-likelihood test comparing the likelihood of a model where the rate of evolution was the same on all branches compared to the likelihood of a model where there was a different rate of evolution on the tuatara branch was conducted for all genes present in both the tuatara and anole (since it was the reference for the alignment) and were present in a minimum of three species.

