## Supplementary Materials for "The tuatara genome: insights into vertebrate evolution from the sole survivor of an ancient reptilian order"

### Table of contents

**1 TUATARA GENOME SEQUENCING, ASSEMBLY AND ANNOTATION** **8**

1.1 Sampling and sequencing 8

1.2 Assembly using Allpaths-LG 8

1.3 Scaffolding the assembly with HiRise 9

1.4 Transcriptome 9

1.5 Epigenome 9

1.6 Repeat annotation 10

1.7 Gene annotation 10

**2 ORTHOLOG CALLING** **12**

2.1 Summary 12

2.2 Methods 12

2.3 Tables 13

2.4 Figures 16

2.5 Download links 17

2.5.1 Source code 17

2.6 Command-lines 18

**3 ENSEMBL ANNOTATION** **19**

3.1 Summary 19

3.2 Methods 19

3.2.1 Ensembl gene prediction 19

3.3 Figures 21

3.4 Tables 22

**4 AB INITIO REPETITIVE DNA ANNOTATION OF THE TUATARA GENOME** **23**

4.1 Introduction 23

4.2 Results 23

4.2.1 Repeat coverage in Sphenodon punctatus 23
4.2.2 Classification of L2 elements in the tuatara genome 26
4.2.3 Tuatara L2 do not cluster with chicken CR1 27
4.2.4 Phylogenetic analysis of tuatara L2 compared to other vertebrates 28
4.2.5 Tuatara L2 may still be active 30
4.2.6 Possible horizontal transfer of L2 between monotreme and tuatara 33
4.2.7 Phylogenetic analysis of tuatara CR1 compared to other vertebrates 35
4.2.8 Potential active CR1 elements in the tuatara genome 37
4.2.9 Divergence rate of CR1 elements in the tuatara genome 38
4.2.10 Unclassified (un-annotated) consensus sequences are probably segmental duplications 39
4.3 Discussion 44
4.3.1 Segmental duplications in the tuatara genome 44
4.3.2 Significance of the LINE retrotransposons in the tuatara genome 45
4.4 Materials and methods 46
4.4.1 Reference genomes: repeat identification, annotation 46
4.4.1.1 Ab initio repeats identification and annotation 46
4.4.2 Analysis of LINE elements in the tuatara genome 47
4.4.3 Resolving L2 classification 47
4.4.4 Dendrogram construction from LINE nucleotide sequence alignments 48
4.4.5 Phylogenetic analysis of L2 elements using RT domain sequences 48
4.4.6 Potential horizontal transfer of L2 elements between tuatara and platypus 49
4.4.7 Divergence rate of CR1 elements in the tuatara genome 49
4.4.8 Identification of novel repeat sequences from the tuatara genome 49
4.4.9 Problems with RMD derived consensus sequences: 50
**5 REPEAT ANNOTATION: SINES AND DNA TRANSPOSONS 52**
5.1 Methods 52
5.2 Results and discussion: DNA transposons 53

53

5.3 Results and discussion: SINE retrotransposons 55

**6 REPORT ON THE DIVERSITY OF LTR RETROELEMENTS IN THE GENOME OF SPHENODON PUNCTATUS 65**

6.1 Introduction 65

6.2 Results 66

6.3 Discussion and conclusions 67

6.4 Methods 67

**7 TUATARA GENOME: NON-CODING RNA ANNOTATION AND ANALYSIS 69**

7.1 Introduction 69

7.2 Methods 69

7.3 Figures 70

7.4 Tables 71

**8 THE nCpG DISTRIBUTION OF THE TUATARA GENOME PERMITS REVISITING THE PATTERNS AND EVOLUTION OF CpG CONTENT IN VERTEBRATES 73**

8.1 Introduction 73

8.2 Methods 74

8.2.1 Data source 74

8.2.2 Data processing 74

8.3 Results and discussion 74

8.4 Figures 76

8.5 Tables 77

**9 EVOLUTION OF GENOMIC ORGANIZATION OF THE MHC 78**

**10 MOLECULAR EVOLUTION OF TUATARA VISUAL SYSTEM 80**

10.1 Introduction 80

10.2 Methods 80

10.3 Results and discussion 81

10.3.1 Visual gene loss in tuatara and amniotes 81

10.3.2 Visual pigments and photoreceptors of tuatara 82

10.3.2.1 Selective pressures of tuatara phototransduction genes 83

10.3.2.2 Evolution of the tuatara visual system 83

10.4 Figures 85

10.5 Tables 88

**11 ODORANT RECEPTORS 90**

11.1 Methods 90

11.2 Results 90

**12 TRANSIENT RECEPTOR POTENTIAL (TRP) ION CHANNELS GENES IN SPHENODON PUNCTATUS 93**

12.1 Background 93

12.2 Methods 94

12.3 Results and discussion 94

12.4 Figures 97

12.5 Tables 101

**13 SELENOPROTEINS 102**

13.1 The tuatara selenoproteome 102

13.2 Multiple tRNASec gene copies 102

**14 THE TUATARA GENOME AND A COMPARISON OF DNA SUBSTITUTION RATES ACROSS AMNIOTES 107**

14.1 Introduction 107

14.2 Whole genome alignments 107

14.3 Data extraction 108

14.4 Phylogenetic analysis 108

14.5 Substitution rate estimation 108

14.6 Results 109

**15 PHYLOGENETIC ANALYSIS OF VERTEBRATE SINGLE-COPY ORTHOLOGS 116**

**16 AN ANALYSIS OF DIVERGENCE TIMES AND TEST FOR PUNCTUATED EVOLUTION118**

16.1 Divergence times 118

16.2 Punctuated evolution 118

**17 PATTERNS OF SELECTION ON TUATARA ORTHOLOGS 121**

17.1 Methods 121

17.1.1 Positive selection on the tuatara lineage 121

17.1.2 Patterns of molecular evolution at sex determining genes 121

17.2 Results 122

17.2.1 Positive selection on the tuatara lineage 122

17.2.2 Patterns of molecular evolution at sex determining genes 123

17.3 Discussion 127

**18 RECONSTRUCTION OF THE DEMOGRAPHIC HISTORY OF THE TUATARA 128**

18.1 Methods 128

18.2 Results 128

18.3 Discussion 128

**19 POPULATION GENOMICS ANALYSES 130**

19.1 Methods 130

19.1.1 Sample collection and library preparation 130

19.1.2 Read mapping and genotyping 130

19.1.3 Functional annotation of SNVs 131

19.1.4 Population genomic analysis 131

19.1.5 Differentiation with respect to sex 131

19.2 Results 132

19.2.1 Both SNV datasets support strong population structure 132

19.2.2 No loci are differentiated by sex 132

19.2.3 No evidence for excess genetic load in any population 132

**20 GENOME RESEARCH AGREEMENT TEMPLATE 138**

**21 REFERENCES 144**

### 1 TUATARA GENOME SEQUENCING, ASSEMBLY AND ANNOTATION

#### 1.1 Sampling and sequencing

A blood sample was obtained from a large male tuatara from Lady Alice Island (35°53'24.4"S 174°43'38.2"E), New Zealand, with appropriate ethical and iwi permissions under permit NO-32037-RES issued by the New Zealand Department of Conservation. Total genomic DNA was extracted using proteinase K digestion and phenol-chloroform extraction, while RNA was prepared using Trizol extraction. Sequencing was undertaken using the Illumina HiSeq 2000 and MiSeq sequencing platforms (Illumina, San Diego, CA, USA). The sequencing libraries consisted of paired end (PE) libraries with insert sizes of 140 bp, 375 bp, 500 and 575 bp and three mate paired (MP) libraries with insert sizes of 2.3 kbp, 4.75 kbp and 8 kbp. The paired end libraries were prepared using the Illumina TruSeq PCR-Free DNA library kit, while the mate pair libraries were prepared using the Illumina TruSeq DNA library kit as per manufacturer's instructions. These libraries were normalised and pooled across 32 lanes on an Illumina HiSeq2000 using 2 x 100 bp paired end sequencing at New Zealand Genomics Ltd., Dunedin. We further supplemented these data with additional Illumina TruSeq and Kappa DNA libraries, with insert sizes of 400 bp and 480 bp, respectively. These libraries were normalised and pooled cross five Illumina MiSeq 2 x 250 bp runs by New Zealand Genomics Ltd., Dunedin, New Zealand.

#### 1.2 Assembly using Allpaths-LG

Raw reads were de novo assembled using Allpaths-LG version 49856[1]). Allpaths-LG was run with several libraries with different insert sizes; overlapping paired-end libraries (PE) of 100 bp with an insert size of 180 bp, paired-end reads with insert size of 350 bp and 550 bp, respectively and several mate pair libraries (MP) with insert sizes of 5 kb, 8 kb and 9 kb. The overlapping paired-end reads were added as fragment libraries, and the 350 bp and 550 nt insert sized libraries and all MP libraries (>180 nucleotide insert size) were added as jumping libraries. Libraries with an insert size up to 550 nt were set with an inward read orientation; all others were set to be oriented away from each other in an outward direction. With a total input data of 5,741,034,516 reads for the paired-end libraries and 2,320,886,248 reads of the mate pair libraries several setups were tested for the optimal proportion of input data. Based on assembly statistics an optimal setup was found using 85% of the fragment libraries and 100% of the jumping libraries. PrepareAllPathsInputs.pl was therefore run with FRAG_COVERAGE=85. On the emergement assembly we used GapCloser[2] and then scaffolded the assembly using our transcriptome (see below) using L_RNA_Scaffolder[3].

#### 1.3 Scaffolding the assembly with HiRise

We scaffolded the Allpaths-LG de novo assembly using HiRise, a software pipeline designed specifically for using proximity ligation data to scaffold genome assemblies[4]. We supplemented our shotgun libraries generated with 44x coverage of Chicago library reads (based on 1–50 kb pairs). Shotgun and Chicago library sequences were aligned to the draft input assembly using a modified SNAP read mapper (http://snap.cs.berkeley.edu). The separations of Chicago read pairs mapped within draft scaffolds were analyzed by HiRise to produce a likelihood model for genomic distance between read pairs, and the model was used to identify and break putative misjoins, to score prospective joins, and make joins above a threshold. After scaffolding, shotgun sequences were used to close gaps between contigs.

#### 1.4 Transcriptome

A de novo transcriptome was assembled as a reference for read mapping and expression quantification using total RNA derived from the blood sample our reference male using a standard RNA-Seq pipeline[5], and a collection of prior transcriptomic data collected from early stage embryos[6]. In total we had 146,739,908 read pairs comprising 131,580,633 new 100 bp read pairs and 15,159,275 prior 50 bp read pairs[6]. These were assembled using Trinity v2.2.0[7,8] running default parameters. To reduce redundancy in the assembled transcripts, CD-HIT-EST v4.6.6[9] was used to collapse contigs with at least 98% identity (e.g., representing alternative alleles from the same locus).

#### 1.5 Epigenome

Low-coverage BS-seq was undertaken using a modified post-bisulfite adapter tagging (PBAT) method[10]. Briefly, 100 ng of purified DNA was subjected to bisulfite conversion using the EZ Methylation Direct Mag Prep kit (Zymo, D5044). Converted DNA underwent first strand synthesis with a biotin labelled adapter sequence possessing seven random nucleotides at its 3' end (BioP5N7, biotin- ACACTCTTTCCCTACACGACGCTCTTCCGATCTNNNNNNN). The product of first strand synthesis was captured using streptavidin coated Dynabeads (Thermo, 11205D) and magnetic immobilisation. Double stranded DNA was created using the immobilized first-strand as a template and an additional adapter also possessing seven random nucleotides at its 3' end (P7N7, GTGACTGGAGTTCAGACGTGTGCTCTTCCGATCTNNNNNNN). Unique molecular barcodes and sequences necessary for binding to Illumina flow-cells were added to libraries by

PCR using 1X HiFi HotStart Uracil+ Mix (KAPA, KK2801 and 10 µM indexed Truseq-type oligos, with thermal cycling as follows: 13*(94°C, 80 sec; 65°C, 30 sec; 72°C, 30 sec).

Successive rounds of paired-end 150 bp sequencing was performed on an Illumina MiSeq. Raw reads were subsequently trimmed in two steps using TrimGalore v0.4.0 (http://www.bioinformatics.babraham.ac.uk/projects/trim_galore) with stringency set to 2. First, adaptor sequences were removed and 10 bp was trimmed from the 5' end to remove nucleotide bias associated with production oligos; reads that became shorter than 105 bp were discarded to avoid bias. In the second step low-quality base calls (Phred score <20) were removed.

Trimmed reads were mapped using Bismark v0.14.3[11] with the --pbat option. Global methylation levels were calculated from Bismark reports by dividing methylated cytosine calls by total cytosine calls in both CG and non-CG contexts.

#### 1.6 Repeat annotation

We initially annotated repeats in the assembly using RepeatModeler to generate a de novo repeat library http://repeatmasker.org/RepeatModeler.html[12]. Next, we used a combination of *ab initio* repeat identification in CARP/RepeatModeler/LTRharvest[12–14], manual curation of specific newly identified repeats, and homology to repeat databases to investigate the repeat content of the tuatara genome. From these three complementary repeat identification approaches, the CARP results were in-depth annotated for LINEs and segmental duplications (Supplementary Materials 4), the RepeatModeler results were in-depth annotated for SINEs and DNA transposons (Supplementary Materials 5), and the LTRharvest results were in-depth annotated for LTR retrotransposons (Supplementary Materials 6). Our manually curated repeat library (https://doi.org/10.5281/zenodo.2585367) was subsequently used to mask repeats in the assembly using RepeatMasker v4.0.3 (http://www.repeatmasker.org/RepeatMasker.html)[15].

#### 1.7 Gene annotation

We used RepeatMasker[15] along with our manually curated de novo repeat database (above) to mask tandem elements in the genome sequence, but did not mask simple repeats at this point to allow for more efficient mapping during gene annotation. Simple repeats were later soft-masked during the gene annotation using MAKER2[16]. Protein-coding genes were predicted using MAKER2[16], using anole lizard (*A. carolinensis*, version AnoCar2.0), python (*P. bivittatus*, version bivittatus-5.0.2,) and RefSeq (www.ncbi.nlm.nih.gov/refseq) as protein homology evidence, which we integrated with ab-initio gene prediction methods including BLASTX, SNAP[17] and Augustus[18]. The SNAP HMM file was generated by training the anole lizard gene sequences. An Augustus model file was generated by training the tool on the OrthoDB set of 3,026 core genes of vertebrates from the genome completeness assessment tool BUSCO[19].

Predicted genes were subsequently used as query sequences in a blastx database search of the NR database (the non-redundant database, http://www.ncbi.nlm.nih.gov/). Blastx alignments with e-value greater than 1e-10 were discarded, and the top hit was used to annotate the query genes. Non-coding RNAs were annotated using Rfam covariance models (v13.0)[20] with Infernal (v1.1)[21] and tRNAscan-SE (v1.3.1)[22], with default score thresholds and parameters.

### 2 ORTHOLOG CALLING

Matthieu Muffato, Mateus Patricio

#### 2.1 Summary

We used the Ensembl method[23] to infer orthology relationships between the tuatara genome and another 25 species. Using sequence similarity and the conservation of local gene order, we could extract 3,168 groups of orthologs with exactly one representative gene in each species.

#### 2.2 Methods

We performed a phylogenetic analysis to infer orthology relationships between the tuatara and another 25 species (see Tables 2.1 and 2.2) using the Ensembl method[23], whose infrastructure had to be expanded to cater for externally annotated genomes (e.g. RefSeq annotations, see Table 2.2). The method is composed of three main steps: (1) a clustering stage, where proteins are clustered into families using hcluster_sg based on NCBI BLAST+ e-values, (2) an alignment stage, where we compute a multiple sequence alignment of each family with M-Coffee or Mafft, and (3) a phylogenetic-tree inference and reconciliation stage using TreeBeST (the reconciliation with a species tree allows calling duplication and speciation events). The species-tree employed in this analysis was the NCBI taxonomy[24]. The computation took in total more than 900 CPU-days on the EMBL-EBI compute cluster and was orchestrated by the Ensembl Hive workflow management system[25].

We then computed an independent evidence score named “Gene Order Conservation” (GOC) score. The GOC score uses local synteny information around a pair of orthologous genes to compute how much the gene order is conserved. Figure 2.1 is the distribution of the GOC scores (using tuatara as a reference) on all species, coloured according to the taxonomy and ordered by the proportion of top-scoring orthologs. It indicates that birds and turtles are the closest species (synteny-wise) to tuatara. Then, tuatara is closer to alligators and squamates, with the exception of the central bearded dragon *Pogona vitticeps* which exhibits a very high level of synteny. The position of the four snakes reminds us that synteny is not necessarily correlated to the evolutionary distance, but rather is a reflection of the rearrangement rate. Figure 2.2 shows the pairwise conservation of synteny, highlighting the relative closeness of some clades (birds and testudines for instance) and the divergence of others (snakes).

An initial phylogenetic analysis was carried on clusters of orthologs that have exactly one copy in all genomes, but these proved to be scarce (128 ortholog groups). This is due to the incompleteness of each assembly and genome used in the analysis, and the discrepancy between the phylogenetic history of some gene families and the species tree. In order to circumvent that, we looked at the 3,040 groups that span all species but have paralogues in at least one of them. Assuming that rearrangements are more likely to happen to a group of contiguous genes, rather than genes in isolation, we decided to use the GOC score as a selector for high-confidence orthologs[26]. For each of those species, we chose the paralogue with the best GOC score and sequence similarity. This way, we could use in total 3,168 clusters of orthologs with exactly 1 representative gene per species (see Table 2.3). Data are available at: 10.5281/zenodo.2542570.

#### 2.3 Tables

**Table 2.1 List of the genomes found in Ensembl**

| **Species name** | **Assembly name** | **Ensembl gene-set version** |
|---|---|---|
| *Gallus gallus* | Galgal4 (GCA_000002315.2) | 2011-12-Ensembl |
| *Meleagris gallopavo* | UMD2 (GCA_000146605.1) | 2010-09-Ensembl |
| *Anas platyrhynchos* | BGI_duck_1.0 (GCA_000355885.1) | 2009-09-Ensembl |
| *Taeniopygia guttata* | taeGut3.2.4 | 2008-08-Ensembl |
| *Ficedula albicollis* | FicAlb_1.4 (GCA_000247815.1) | 2012-05-Ensembl |
| *Pelodiscus sinensis* | PelSin_1.0 (GCA_000230535.1) | 2011-11-Ensembl |
| *Anolis carolinensis* | AnoCar2.0 (GCA_000090745.1) | 2010-09-Ensembl |
| *Monodelphis domestica* | BROADO5 (GCF_000002295.2) | 2007-02-Ensembl |
| *Homo sapiens* | GRCh37 (GCA_000001405.14) | 2010-07-Ensembl |
| *Ornithorhynchus anatinus* | OANA5 (GCF_000002275.2) | 2007-01-Ensembl |

| | | |
|---|---|---|
| *Danio rerio* | GRCz10 (GCA_000002035.3) | 2014-09-Ensembl |
| *Lepisosteus oculatus* | LepOcu1 (GCA_000242695.1) | 2012-01-Ensembl |
| *Xenopus tropicalis* | JGI_4.2 (GCA_000004195.1) | 2010-09-Ensembl |
| *Mus musculus* | GRCm38 (GCA_000001635.5) | 2012-01-Ensembl |

**Table 2.2. List of non-Ensembl genomes**

| **Species name** | **Annotation source** | **Download link (GFFs and FASTAs)** |
|---|---|---|
| *Sphenodon punctatus* | augustus_maker | http://doi.org/10.5281/zenodo.1489354 |
| *Alligator mississippiensis* | refseq | ftp://ftp.crocgenomes.org/pub/ICGWG/Genome_drafts/alligator.current/ |
| *Alligator sinensis* | refseq | ftp://ftp.ncbi.nlm.nih.gov/genomes/all/GCF/000/455/745/GCF_000455745.1_ASM45574v1 |
| *Chelonia mydas* | refseq | ftp://ftp.ncbi.nlm.nih.gov/genomes/all/GCF/000/344/595/GCF_000344595.1_CheMyd_1.0 |
| *Chrysemys picta* | refseq | ftp://ftp.ncbi.nlm.nih.gov/genomes/all/GCF/000/241/765/GCF_000241765.3_Chrysemys_picta_bellii-3.0.3 |
| *Gekko japonicus* | refseq | ftp://ftp.ncbi.nlm.nih.gov/genomes/all/GCF/001/447/785/GCF_001447785.1_Gekko_japonicus_V1.1 |
| *Protobothrops mucrosquamatus* | refseq | ftp://ftp.ncbi.nlm.nih.gov/genomes/all/GCF/001/527/695/GCF_001527695.2_P.Mucros_1.0 |
| *Python molurus* | refseq | ftp://ftp.ncbi.nlm.nih.gov/genomes/all/GCF/000/186/305/GCF_000186305.1_Python_molurus_bivittatus |

| | | |
|---|---|---|
| *bivittatus* | | -5.0.2 |
| *Thamnophis sirtalis* | refseq | ftp://ftp.ncbi.nlm.nih.gov/genomes/all/GCF/001/077/635/GCF_001077635.1_Thamnophis_sirtalis-6.0 |
| *Ophiophagus hannah* | refseq | ftp://ftp.ncbi.nlm.nih.gov/genomes/all/GCA/000/516/915/GCA_000516915.1_OphHan1.0 |
| *Ophisaurus gracilis* | gigascience | http://gigadb.org/dataset/100119 |
| *Pogona vitticeps* | gigascience | http://gigadb.org/dataset/100166 |

**Table 2.3 Statistics on the clusters of orthologs**

<table>
<tr><td colspan="2">Total number of ortholog clusters</td><td>23,111</td></tr>
<tr><td colspan="2">Clusters with less than 26 species</td><td>19,943</td></tr>
<tr><td rowspan="3">Clusters with all 26 species present</td><td>Total</td><td>3,168</td></tr>
<tr><td>Clusters with exactly 1 gene per species</td><td>128</td></tr>
<tr><td>Clusters with paralogues</td><td>3,040</td></tr>
</table>

#### 2.4 Figures

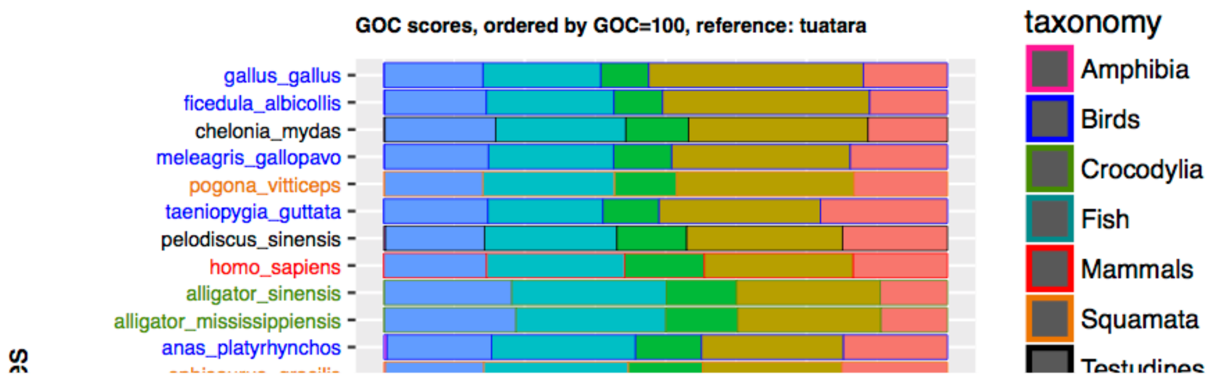

**Figure 2.1 Gene Order Conservation score distribution using tuatara as reference (species ordered by the proportion of top-scoring orthologs)**

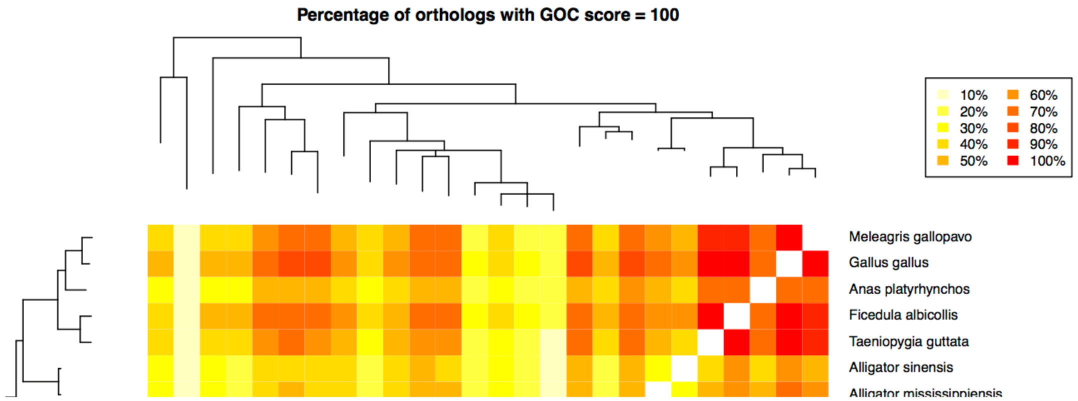

**Figure 2.2 Heat map representing the proportion of top-scoring orthologs between each pair of species**

### 2.5 Download links

#### 2.5.1 Source code

- Ensembl pipeline to compute trees and orthologs: https://github.com/Ensembl/ensembl and https://github.com/Ensembl/ensembl-compara branch "release/87"

- Ensembl Hive workflow management system https://github.com/Ensembl/ensembl-hive branch "version/2.3"
- Plotting script: https://github.com/Ensembl/ensembl-compara/blob/release/89/scripts/homology/plotGocData.r

#### 2.6 Command-lines

- Ensembl pipeline to compute trees. After setting up paths and server locations in `ensembl-compara/modules/Bio/EnsEMBL/Compara/PipeConfig/Example/TuataraProteinTrees_conf.pm`
  ```
  $ init_pipeline.pl
  Bio::EnsEMBL::Compara::PipeConfig::Example::TuataraProteinTrees_conf
  ```
  And then run the pipeline with the *beekeeper.pl* command-line that is advertised.
- Ensembl pipeline to extract Fasta, XML, TSV and EMF files from the gene-tree database.
  ```
  $ init_pipeline.pl
  Bio::EnsEMBL::Compara::PipeConfig::DumpTrees_conf
  -member_type protein -clusterset_id default
  ```
- Extraction of clusters of orthologs
  ```
  $ perl
  ensembl-compara/scripts/examples/homologyForPhylogeny.pl
  -compara_url mysql://user@server:port/database_name
  -species_set_file /path/to/file_with_all_species_names
  -outdir /path/to/output_dir/ -species_threshold 26 -debug 1
  > /path/to/debug_output_file
  ```
- Generation of the plots, reference species should be defined on reference_species.dat file, one species per line.
  ```
  $ Rscript ensembl-compara/scripts/homology/plotGocData.r
  /your_source_dir/tuatara.tree /your_source_dir/
  reference_species.dat
  ```

### 3 ENSEMBL ANNOTATION

Konstantinos Billis, Fergal J Martin

#### 3.1 Summary

Ensembl annotation, undertaken independent of the gene set annotated using MAKER2 (Supplementary Section 1.7), identified 17628 protein-coding genes, 767 pseudogenes and 910 noncoding RNAs.

#### 3.2 Methods

##### 3.2.1 Ensembl gene prediction

The tuatara genome assembly (version ASM311381v1) was annotated with the Ensembl gene annotation system[27]. Protein-coding gene models were annotated by combining alignments of UniProt[28] and RNA-Seq models generated from the publicly available blood sample (SRR7084910). The genome was repeat-masked with RepeatMasker, using the RepBase library (parameters: -species vertebrates) and using a custom library generated with RepeatModeler, and Dust[29]. Protein-coding models were generated by aligning reptiles, birds, mammal and other vertebrate protein sequences with experimental evidence from UniProt protein existence levels 1 (evidence at the protein level) and 2 (evidence at the transcript level). These alignments were made to the repeat-masked genome in a splice aware manner using GenBlast[30]. Our in-house RNA-Seq pipeline[31] also generated protein-coding models. Reads from blood sample were aligned to the unmasked genome using BWA[32]. The alignments were processed by collapsing the transcribed regions into a set of rough exons. Partially aligned reads were re-mapped using Exonerate[33] and this step identified spliced reads or introns. These introns together with the set of transcribed exons were combined to produce transcript models. The longest open reading frame in each of these models were BLASTed[34] against the set of UniProt vertebrate proteins with experimental evidence. For all ORFs with a BLAST match, the top hit was then aligned directly to the translated ORF using MUSCLE[35]. The alignment coverage and percent identity were calculated and stored, this was used later to help select well-supported transcripts and filter out fragmented ORFs.

Data from the above two pipelines were filtered to remove poorly supported models. Untranslated regions were added to the coding models using RNA-Seq data. The preliminary sets of coding models were combined, prioritizing well-supported models built from UniProt proteins and the RNA-Seq data at each locus. Redundant transcript structures were filtered out and the remaining models were collapsed into an initial set of protein coding genes.

The set of protein coding gene models was screened for pseudogenes. Cases where there was obvious evidence of repeated frame shifting, in-frame stop codons or genes that were completely covered in repeats were flagged as pseudogenes. Single exon protein coding genes were also examined for evidence of a multi-exon counterpart elsewhere in the genome as such cases implied likely retrotransposition events. These were flagged as processed pseudogenes.

Potential small non-coding RNA genes were initially identified by a BLAST of RFAM and miRbase[36] sequences against the genome. The resulting BLAST hits were then further filtered using RNAfold for miRNAs and Infernal[36] for other small non-coding gene types. The remaining models were screened for redundancy and the remainder formed the small non-coding gene set.

The Ensembl gene annotation for tuatara is released in Ensembl version 95, including orthologs, gene trees, and whole-genome alignments other species. Also included are tracks for the RNA-Seq transcript models, the BAM alignment coverage and the complete set of splice junctions identified by our pipeline. An indexed BAM file for the blood sample alignments will be available for download. Additionally, the MAKER annotation generated by the tuatara consortium will be available in Ensembl as a supplementary gene track.

Data for tuatara can be accessed in Ensembl in a variety of ways including the genome browser, the Perl and REST APIs, the ftp site, BioMart and direct access to our MySQL databases.

#### 3.3 Figures

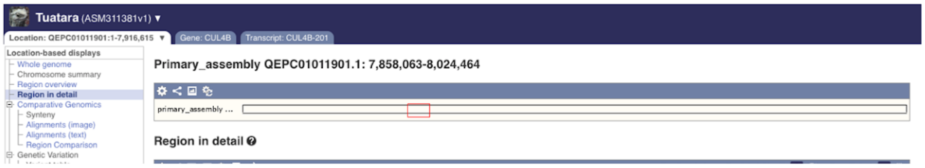

**Figure 3.1 Tuatara genome annotation as seen in the Ensembl browser.** The Location tab allows the visualisation of genes, sequence conservation, and other annotation aligned to the genome. The lower panel, called the Region in Detail view, displays the following tracks: aligned proteins from the Homology pipeline, RNAseq models that built based on RNAseq pipeline, RNAseq coverage of the region, introns that were reported by RNAseq pipeline and the two available gene sets from Ensembl and the Tuatara Consortium.

#### 3.4 Tables

**Table 3.1 Gene and transcript counts for tuatara.**

| **Biotype** | **Gene counts** | **Transcript counts** |
| --- | --- | --- |
| *Protein coding* | 17628 | 25240 |
| *Pseudogene* | 767 | 767 |
| *ncRNA* | 910 | 910 |
| *IG genes* | 20 | 20 |

### 4 *AB INITIO* REPETITIVE DNA ANNOTATION OF THE TUATARA GENOME

Lu Zeng, Terry Bertozzi, R. Daniel Kortschak, Joy M. Raison, David L. Adelson*

#### 4.1 Introduction

In this section, we use an *ab initio* repeat identification and annotation method[13] to investigate the repeat content of the tuatara genome. In order to evaluate the performance of our *ab initio* method, we have compared the repeat libraries generated from our method and from RepeatModeler[12] (RMD). Our analysis revealed that most repeats in the tuatara genome are non-LTR LINE L2 retrotransposons. We also investigated the evolutionary relationships between tuatara L2 sequences and those from other vertebrates using both the full nucleotide sequences and the Reverse Transcriptase (RT) domains of these L2 sequences. We found two main sub-families of L2 in the tuatara, one similar to lizard L2s and the other similar to platypus L2s. The latter provide potential evidence for a horizontal transfer event of L2s between lizards and monotremes. Finally, there were a large number of repeated sequences that remain unclassified from our *ab initio* method and these are likely segmental duplications in the tuatara genome. However, some highly repeated sub-sequences in these duplications are potential novel non-autonomous transposons.

#### 4.2 Results

##### 4.2.1 Repeat coverage in *Sphenodon punctatus*

Using the *ab initio* method, around 64% of the reference tuatara genome was annotated as repetitive (Table 4.1), and 31% of the genome was comprised of known repeats. This indicated that the tuatara genome was significantly enriched with repeats compared to other reptile genomes. One interesting observation was that the fraction of non-LTR retrotransposons in the tuatara genome was much higher (16.5%) than in placental mammals, with L2 being the dominant LINE element. We estimated that there are 1.4 million L2 sequences (full length and fragments) in the tuatara genome, and the longest L2 sequences we found were approximately 4 kb in length. The second most abundant repeat type was CR1 (chicken repeat 1), which comprised 3% of the tuatara genome.

In contrast to our method, RepeatModeler (RMD) identified approximately 51% of the tuatara genome as repetitive, and 36% of this was annotated based on Repbase data. L2 and CR1 were also the two dominant non-LTR retrotransposons. RMD identified 9.6% of the tuatara genome as DNA transposons, mainly from the DNA/hAT (hobo/AC/Tam3) superfamily (4.5%) and the Harbinger superfamily (4.2%). However, further analysis of the RMD consensus sequences casts significant doubt on the accuracy of these findings and this identification needs to be validated (see Supplementary Materials 4.4.9 Problems with RMD derived consensus sequences).

**Table 4.1: Copy number and percentage of tuatara genome covered by interspersed repeats.**

| | *ab initio* library | | | RMD library | | |
|---|---|---|---|---|---|---|
| Group | Number | Total bp | Percentage | Number | Total bp | Percentage |
| Non-LTR retrotransposons | | | | | | |
| LINE L2 | 1,479,425 | 428,138,325 | 10.022 | 1,149,043 | 491,620,677 | 11.507 |
| LINE CR1 | 815,011 | 162,227,767 | 3.797 | 500,647 | 149,462,580 | 3.498 |
| LINE RTE | 138,126 | 31,151,317 | 0.729 | 184,481 | 97,841,699 | 2.290 |
| LINE L1 | 239,829 | 30,179,939 | 0.706 | 49,905 | 29,081,314 | 0.681 |
| SINE | 329,988 | 32,931,264 | 0.771 | 184,701 | 27,664,250 | 0.647 |
| Others | 209,995 | 21,413,494 | 0.501 | 46,718 | 21,910,444 | 0.514 |
| | 3,212,374 | 706,042,106 | 16.526 | 2,115,495 | 817,580,964 | 19.137 |
| ERV | | | | | | |
| ERV1 | 142,313 | 17,564,350 | 0.411 | 20,968 | 9,453,167 | 0.221 |
| Others | 145,005 | 15,355,898 | 0.359 | 7,149 | 1,948,963 | 0.046 |
| | 287,318 | 32,920,248 | 0.770 | 28,117 | 11,402,130 | 0.267 |
| DNA transposons | | | | | | |
| hAT | 439716 | 57197609 | 1.339 | 668,329 | 194,724,290 | 4.558 |
| Harbinger | 435048 | 80071368 | 1.874 | 463,162 | 180,461,900 | 4.224 |
| Others | 1,564,326 | 224,599,096 | 5.257 | 167,048 | 36,855,177 | 0.863 |
| | 2439090 | 361868073 | 8.470 | 1,298,539 | 412,041,367 | 9.645 |
| LTRs | | | | | | |
| DIRS | 256,478 | 58,538,598 | 1.370 | 130,860 | 78,246,082 | 1.831 |
| Gypsy | 570,277 | 83,980,917 | 1.966 | 98,683 | 62,432,545 | 1.461 |
| LTR others | 221,043 | 26,438,648 | 0.619 | 259,570 | 115,859,366 | 2.713 |
| | 1,047,798 | 168,958,163 | 3.955 | 489,113 | 256,537,993 | 6.005 |
| Others | 599,649 | 67,376,623 | 1.578 | 316,948 | 58,30,9,055 | 1.364 |
| Well-annotated | 7,586,229 | 1,337,165,213 | 31.299 | 4,248,212 | 1,555,871,509 | 36.418 |
| Unknown | 7875491 | 1424963821 | 33.354 | 3,203,800 | 637,030,810 | 14.911 |
| Total | 15,461,720 | 2,762,129,034 | 64.653 | 7,452,012 | 2,192,902,319 | 51.329 |

#### 4.2.2 Classification of L2 elements in the tuatara genome

L2 elements were found to be the most abundant repeat type in the tuatara genome using both repeat annotation methods. In the RMD repeat library, 29 L2 sequences were longer than 2 kb, with 9 of them approximately 3 kb long. In our *ab initio* library, 27 consensus sequences were longer than 2 kb with one over 3 kb long. Figure 4.1 shows the sequence similarity of L2 elements (2–4 kb) from these two methods. RMD L2 sequences were found to cluster with our L2 consensus sequences, and were most similar to turtle, platypus L2s, and anole L2 based on CENSOR annotation. In addition, L2 consensus sequences from our method that were annotated as most similar to platypus L2, were also found always clustered with turtle L2 based on CENSOR annotation. However, the Repbase library classified platypus L2 as CR1. This is one example of the confusing state of repeat annotations and nomenclature. The remaining L2 consensus sequences (including the longest L2) were found to be similar to anole L2 sequences based on Repbase annotation.

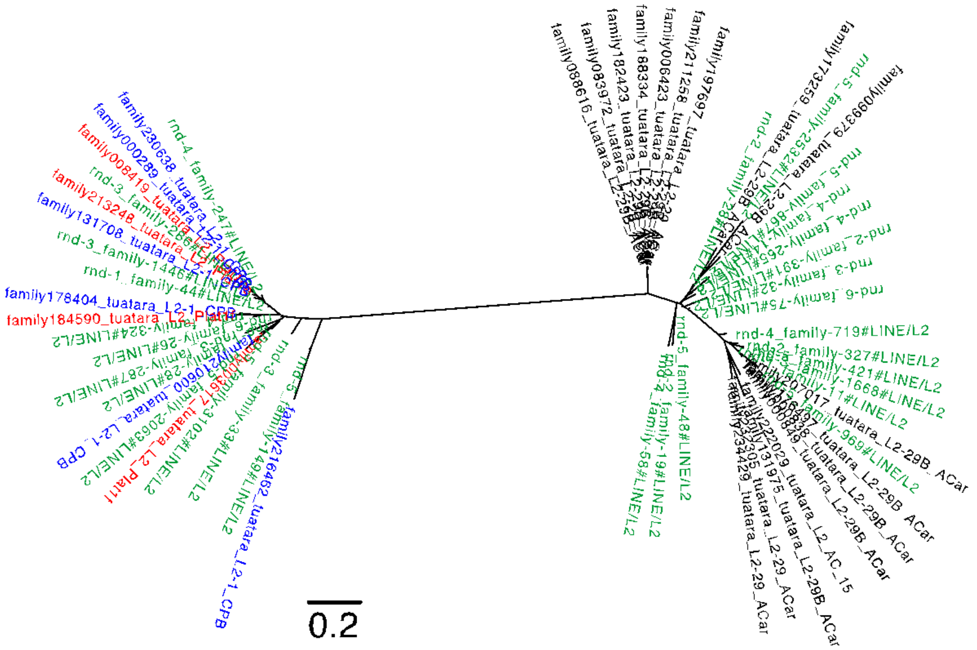

**Figure 4.1: Sequence similarity analysis of RMD L2 and *ab initio* L2 consensus sequences.** Shown is a Maximum likelihood dendrogram inferred using FastTree using a global alignment of tuatara L2 nucleotide sequences generated from both our *ab initio* method and RMD. Green labels identify RMD tuatara sequences, while other colours denote tuatara L2 sequences similar to turtle (blue), platypus (red), and anole (black).

##### 4.2.3 Tuatara L2 do not cluster with chicken CR1

As mentioned above, platypus L2 sequences are categorised as belonging to the CR1 clade according to Repbase. Therefore, we used full-length chicken CR1 sequences to analyse the overall clustering pattern of tuatara L2 sequences (Figure 4.2). With the inclusion of chicken CR1 sequences, tuatara L2 sequences split into 4 groups; one group contains repeats similar to platypus L2, turtle L2 and RMD L2, the other three groups are made up of L2 sequences that are most similar to anole. The full-length chicken CR1 elements did not cluster with the L2 consensus sequences, indicating that all of the L2 sequences we have identified are not CR1 like, in spite of the Repbase annotation.

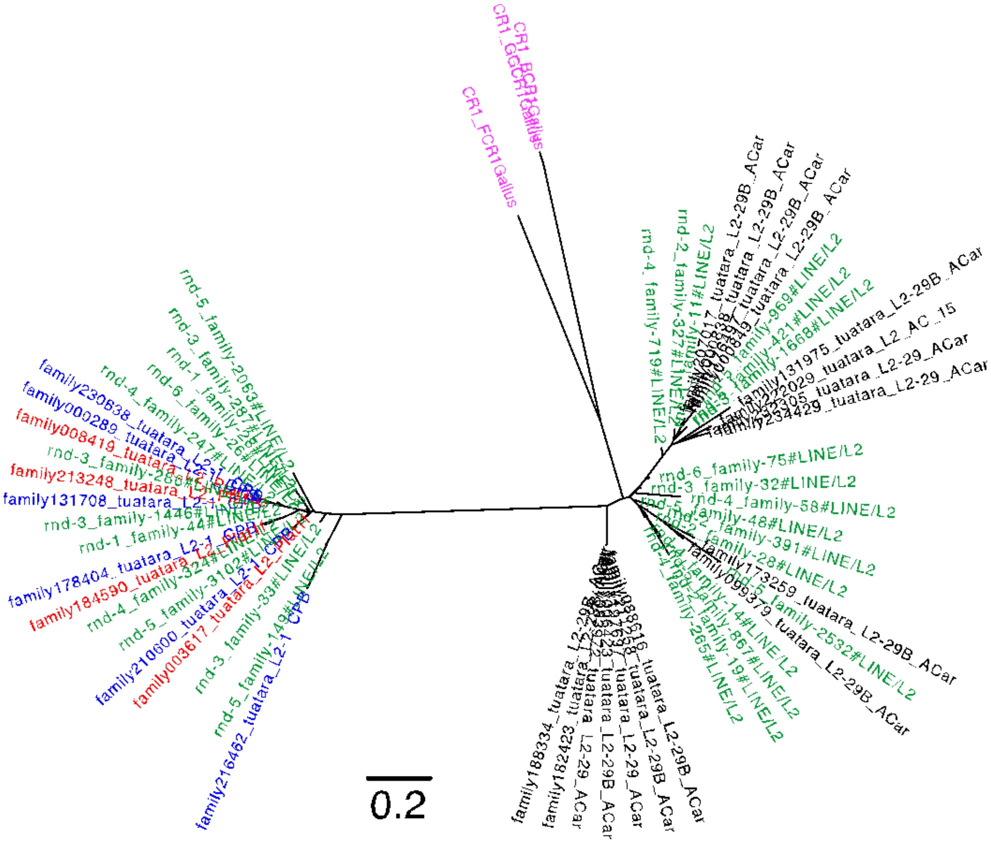

**Figure 4.2: Tuatara L2 and chicken CR1 do not cluster together.** Maximum likelihood dendrogram inferred using FastTree based on global alignment of tuatara L2 nucleotide sequences and chicken CR1. All sequences were generated from our *ab initio* method, except green labelled sequences, which are from RepeatModeler. Dark blue labels are tuatara L2 sequences similar to turtle, red labels are tuatara L2 similar to

platypus, black labels are L2 sequences similar to anole, green labels are RMD tuatara sequences, and pink labels are chicken CR1 from Repbase.

##### 4.2.4 Phylogenetic analysis of tuatara L2 compared to other vertebrates

We then globally aligned the L2 tuatara sequences with L2 sequences from other vertebrates. The resulting maximum likelihood based phylogenetic analysis clearly showed that tuatara L2 sequences are divided into two groups (Figure 4.3), one group is more similar to bearded dragon and anole L2 (L2 from RMD and our *ab initio* method), while the other group clustered with the turtle and platypus L2 (L2 from RMD and our *ab initio* method). Platypus like L2 consensus sequences identified from our *ab initio* method consistently clustered with platypus L2 from Repbase. The presence of two lineages of L2 (reptile vs monotreme) in tuatara is not characteristic of other reptiles and may be the result of incomplete lineage sorting of L2 in tuatara or raises the possibility of horizontal transfer of L2 between monotremes and the tuatara lineage.

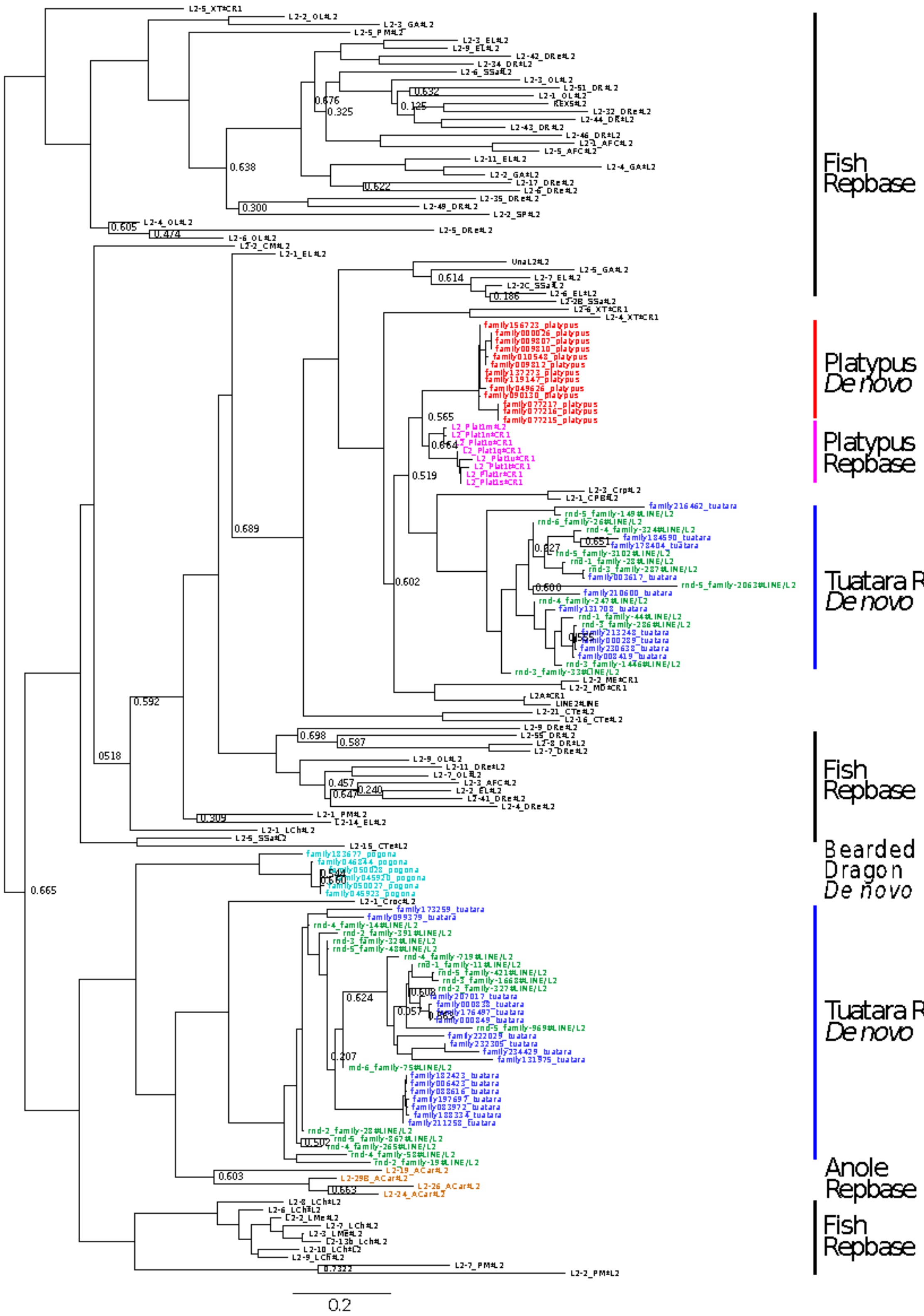
Fish Repbase
Platypus De novo
Platypus Repbase
Tuatara RMK/ De novo
Fish Repbase
Bearded Dragon De novo
Tuatara RMK/ De novo
Anole Repbase
Fish Repbase
0.2

**Figure 4.3: Phylogenetic analysis of vertebrate L2.** Maximum likelihood dendrogram inferred using FastTree based on global alignment of L2 nucleotide sequences. L2 sequences were either extracted from full genome assemblies (tuatara, platypus and pogona) or sourced from Repbase. Sequences were aligned with MUSCLE, and visualised with FigTree. Branches are coloured to indicate the original L2 source. Dark blue labels are tuatara L2 sequences generated using our *ab initio* method, dark green labels are tuatara L2 sequences from RepeatModeler, pink labels are platypus sequences from Repbase, red labels are platypus sequences from our *ab initio* method, light green labels are pogona *ab initio* sequences and orange labels are anole sequences from Repbase.

##### 4.2.5 Tuatara L2 may still be active

We carried out open reading frame (ORF) analysis of the long tuatara L2 consensus sequences (2–4 kb) and their corresponding genomic sequences to identify potentially active L2 elements. This also reduced the confounding effects of length in Figures 4.1 and 4.2, which only looked at sequence similarity within the tuatara L2 sequences we identified.

With respect to our *ab initio* consensus sequences, we found that 27 of the tuatara consensus sequences were longer than 2 kb, with 1 of them longer than 3 kb. We also found that 13 (48%) of these consensus sequences appeared to contain intact open reading frame 2 (ORF2p) (longer than 500 amino acids and contained a RVT_1 reverse transcriptase motif). While in tuatara genomic sequences we found that 639 of the tuatara genome fragments were longer than 2 kb, with four of them longer than 3 kb. Significantly, we found that 530 (83%) of these genomic sequences appeared to contain intact ORF2p, see Table 4.2. This strongly suggests that L2 elements may still be active in the tuatara genome.

**Table 4.2:** Copy number and fraction of the tuatara genome covered by interspersed repeats.

| | 3~4 kb | | 2~3 kb | |
|---|---|---|---|---|
| | Number full length | % active L2 | Number full length | % active L2 |
| RMD L2 consensus | 9 | 44 | 20 | 30 |
| *ab initio* L2 consensus | 1 | 100 | 26 | 46 |
| *ab initio* L2 genome | 4 | 100 | 635 | 83 |

We also carried out global alignments of >500aa long ORF2p sequences from tuatara and Repbase consensus sequences that had RT domains >200aa in length (see Methods). The phylogenetic analysis of RT families (Figure 4.4) clearly illustrated differences between L2 groups. This figure shows that tuatara L2 RT domains differed from anole and split into two clusters, one cluster was closer to platypus, while the other cluster was closer to bearded dragon. The L2 RT domain from crocodile diverged from all other species. Although the L2 nucleotide phylogenetic tree showed that tuatara L2 shared high similarity with anole L2, when comparing sequences based on the RT domain, they were quite different, with anole RT domains clearly separated from other species. This result is consistent with either incomplete lineage sorting of two L2 families or horizontal transfer of L2 sequences to, or from, a monotreme. Both of these families may still be active.

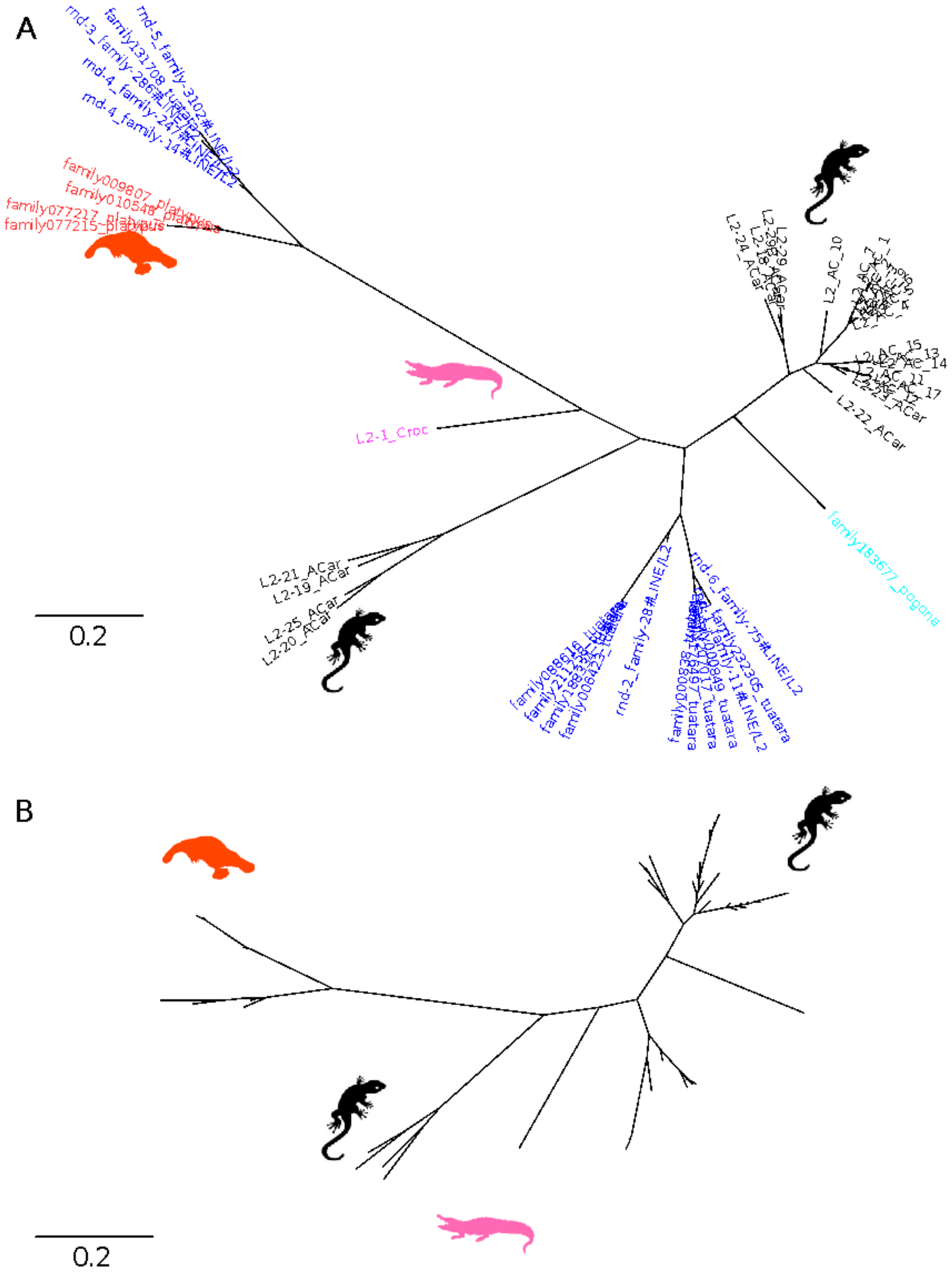

**Figure 4.4: Phylogenetic analysis based on the RT domain.** A) Maximum likelihood dendrogram inferred using FastTree from L2 RT domain multiple alignment. Branches are coloured to indicate the original L2 source: dark blue labels are tuatara L2 RT domain generated from our *ab initio* method and RepeatModeler, red branches are

platypus L2 RT domains, light blue branches are L2 RT domains from bearded dragon (pogona) and black branches were L2 RT domains from anole. B) Bayesian dendrogram inferred using MrBayes from L2 RT domain multiple alignment. Branch labelling same as panel A.

##### 4.2.6 Possible horizontal transfer of L2 between monotreme and tuatara

Based on both nucleotide and RT domain phylogenies, L2 elements consistently split into two groups, one most similar to lizard L2 (33 consensus sequences), the other most similar to platypus L2 (22 consensus sequences), indicating a possible horizontal transfer event between monotreme and tuatara. We used these sequences to generate custom L2 libraries for repeat annotation (see below).

We developed a method based on reciprocal best hits for identifying horizontally transferred sequences. First, CENSOR was used to find hits that annotated as platypus-like tuatara L2 or non-platypus like tuatara L2 in five reptile genomes (anole, alligator, crocodile, turtle and bearded dragon) and one monotreme genome (platypus) using the custom L2 libraries. Second, the CENSOR hit sequences were extracted and used as BLASTN queries to find reciprocal best hits in the custom libraries. This allowed us to determine the reliability of the original annotation of L2 hits and determine the L2 family they are most likely to belong to.

Based on the initial CENSOR output (Table 4.3), platypus-like tuatara L2 elements were found to be enriched in the platypus genome, while non-platypus tuatara L2 elements were abundant in both the anole and bearded dragon genomes. Validation of these hits using reciprocal BLASTN (Table 4.4) shows that although many fragments in reptiles were annotated as platypus-like tuatara L2 by CENSOR, they did not validate as reciprocal hits. This is most likely a product of different parameter settings for BLASTN used by CENSOR and the more stringent default settings used in the reciprocal BLASTN search. However, 16/22 platypus-like L2 tuatara consensus sequences were validated as being most similar to platypus L2 based on their reciprocal best hits. Similarly, 21/33 reptile-like tuatara L2 consensus sequences were validated as non-platypus like based on the reciprocal best-hit results.

**Table 4.3: CENSOR output.** Summary of non-redundant CENSOR hit genome intervals used as queries for reciprocal BLASTN alignment. Number of consensus sequences in custom library (in parentheses). PT L2 = Platypus-like tuatara L2; NPT L2 = Non-platypus like tuatara L2; int = Intervals; con = Consensus.

| CENSOR library | Anole | Alligator | Crocodile | Bearded dragon | Turtle | Platypus |
|---|---|---|---|---|---|---|

| | int | (con) | int | (con) | int | (con) | int | (con) | int | (con) | int | (con) |
|---|---|---|---|---|---|---|---|---|---|---|---|---|
| PT L2 (22) | 26,954 | (22) | 37,502 | (22) | 34,842 | (22) | 27,445 | (22) | 103,729 | (22) | 406,615 | (22) |
| NPT L2 (33) | 213,764 | (33) | 20,584 | (33) | 19,717 | (33) | 127,972 | (33) | 46,802 | (33) | 258,675 | (33) |

**Table 4.4: BLASTN output.** Summary of the BLASTN outputs for the reciprocal alignments of the intervals from Table 3 against platypus-like tuatara L2 and non-platypus like tuatara L2 consensus sequences. Number of validated consensus sequences (in parentheses).

| L2 Type | Anole | | Alligator | | Crocodile | | Bearded dragon | | Turtle | | Platypus | |
|---|---|---|---|---|---|---|---|---|---|---|---|---|
| | int | (con) | int | (con) | int | (con) | int | (con) | int | (con) | int | (con) |
| PT L2 (22) | 0 | (0) | 0 | (0) | 0 | (0) | 0 | (0) | 1 | (1) | 2,281 | (16) |
| NPT L2 (33) | 761 | (21) | 21 | (6) | 25 | (6) | 759 | (22) | 5 | (2) | 0 | (0) |

Furthermore, in order to estimate the relative ages of platypus-like tuatara L2 and non-platypus like tuatara L2, super consensus were built for each of L2 types (see Section 4.4), RepeatMasker[15] was then used to calculate the divergence rate between super consensuses and their corresponding tuatara L2 elements (e.g. platypus-like tuatara L2 super consensus against platypus-like tuatara L2 elements). Figure 4.5 shows that compared to non-platypus like tuatara L2, platypus-like tuatara L2 has a lower sequence substitution level, which indicates that the platypus-like tuatara L2 sequences are of more recent origin, and may have resulted from horizontal transfer from platypus, or some intermediate vector, into the tuatara genome.

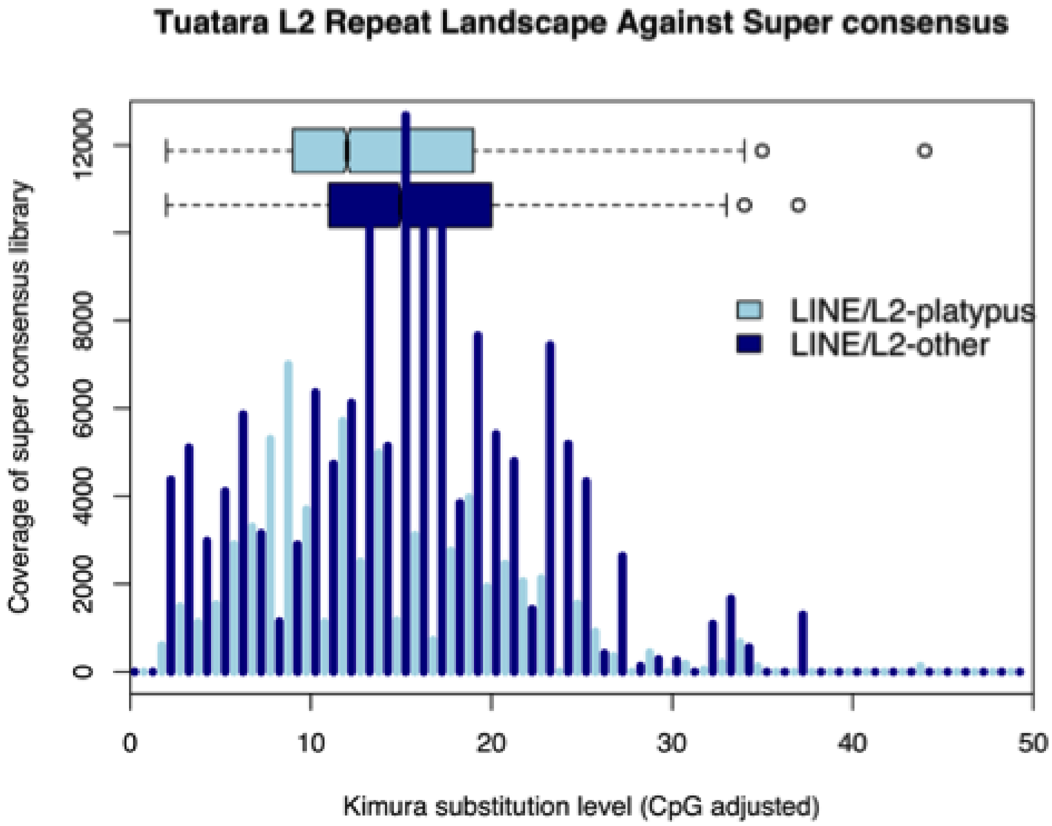

**Figure 4.5: Kimura substitution level of tuatara L2 elements.** The divergence rate of platypus-like tuatara L2 and non-platypus like tuatara L2 was calculated using the Kimura 2-Parameter divergence metric, and adjusted for 'GC' content. Dark blue bars show the sequence substitution rate of non-platypus like tuatara L2 against non-platypus like L2 super consensus, while light blue bars show the sequence substitution rate of platypus-like tuatara L2 against platypus-like L2 super consensus. Each pair of bars shares the same substitution level. Boxplot shows the upper and lower quartiles and the mean value of each tuatara L2 type with respect to their divergence rate.

##### 4.2.7 Phylogenetic analysis of tuatara CR1 compared to other vertebrates

Based on both repeat annotation methods (Table 4.1), CR1 elements were the second most abundant LINE in the tuatara genome , making up ~3% of the genome. Thirty-nine of these CR1 consensus sequences were found to be longer than 2.5 kb, potentially with an intact ORF2. The longest CR1 sequence found in the tuatara genome is 4,605 bp. We globally aligned these CR1 tuatara sequences with CR1 sequences from other vertebrates (231 in total). The resulting maximum likelihood based phylogenetic analysis showed that tuatara CR1 sequences were distinct from those found in other species, including the anole lizard (Figure 4.6).

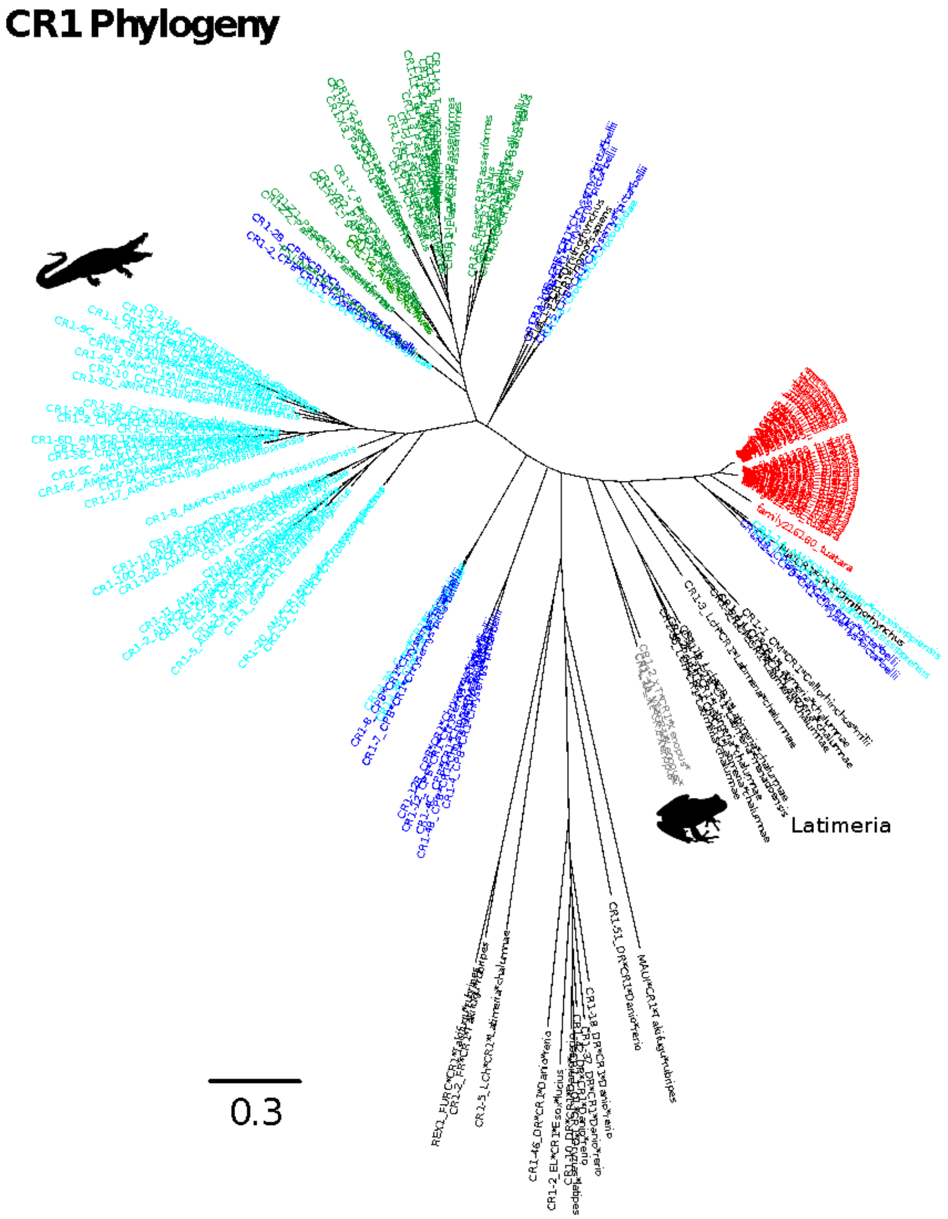

**Figure 4.6: Phylogenetic analysis of vertebrate CR1.** Maximum likelihood dendrogram inferred using FastTree based on global alignment of CR1 nucleotide sequences. CR1 sequences were extracted from species full genome assemblies (tuatara and anole lizard) using CARP [13] or sourced from RepBase. Sequences were aligned with MUSCLE and visualised with Archaeopteryx. Branch colours indicate the original CR1 source. Red labels are tuatara, dark blue labels are turtle, dark green labels are anole lizard, light blue labels are crocodilian sequences, pink labels are platypus sequences, black labels are bird CR1 sequences, and light grey labels are frog CR1 sequences.

##### 4.2.8 Potential active CR1 elements in the tuatara genome

We then further analysed the 39 CR1 consensus sequences that were longer than 2.5 kb, 13 of which contained an intact RT domain. Tuatara CR1 were split into two groups, which may represent two classes of CR1 elements in the tuatara genome. (Figure 4.7).

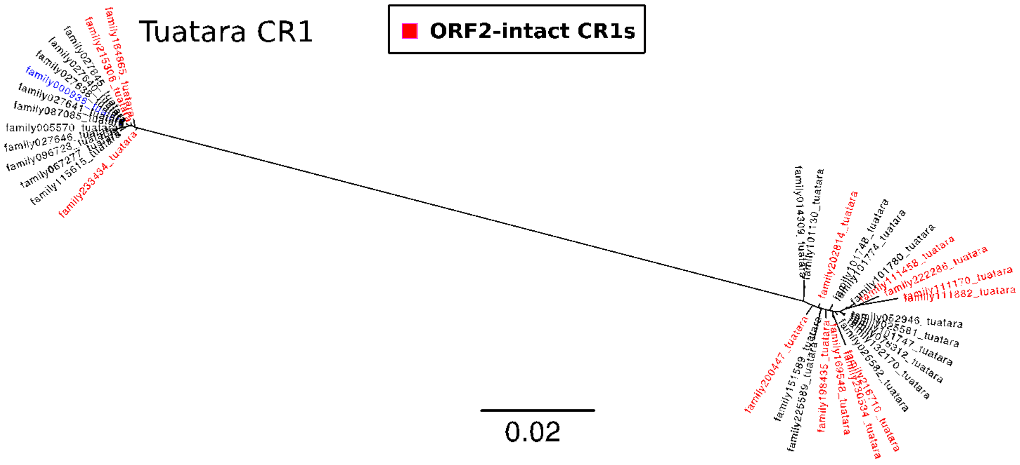

**Figure 4.7: Phylogenetic analysis of CR1 elements in the platypus genome.** Dendrogram of full-length CR1 elements in the tuatara genome. Sequences were aligned with MUSCLE, trees inferred with FastTree and visualized with Archaeopteryx. ORF2-intact CR1s are shown with a red label the tip of the branch, and the longest CR1 is labelled with dark blue.

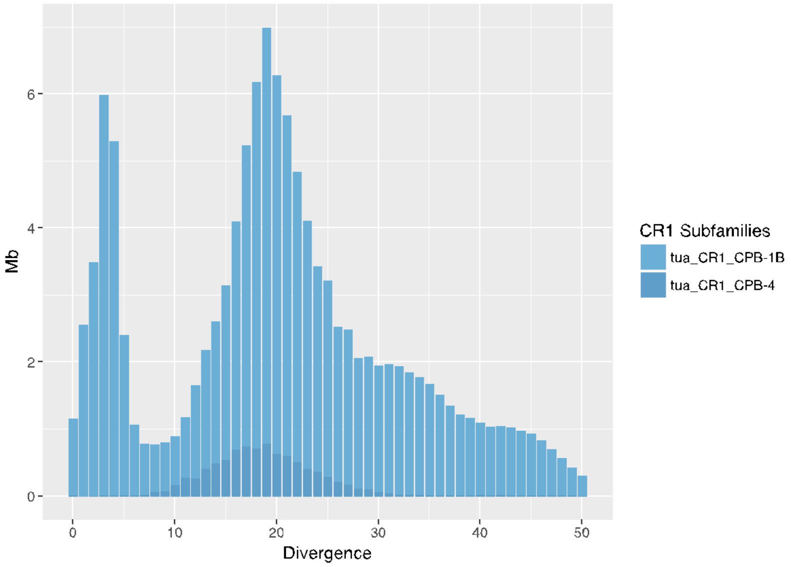

**Figure 4.8: The landscape of full-length CR1 in the tuatara genome.** Long CR1 subfamilies with a total length greater than 1,000 bp and with distances to consensus <50% are shown.

We also calculated the divergence rate of these potentially active CR1 elements, Figure 4.8 shows that the two classes of CR1 elements in the tuatara genome appear to have different CR1 activity profiles over time.

##### 4.2.9 Divergence rate of CR1 elements in the tuatara genome

Considering all CR1 elements in the tuatara genome, we identified 14 CR1 subfamilies (Figure 4.9); some of which had low divergence compared to consensus, indicating a recent wave of CR1 insertion.

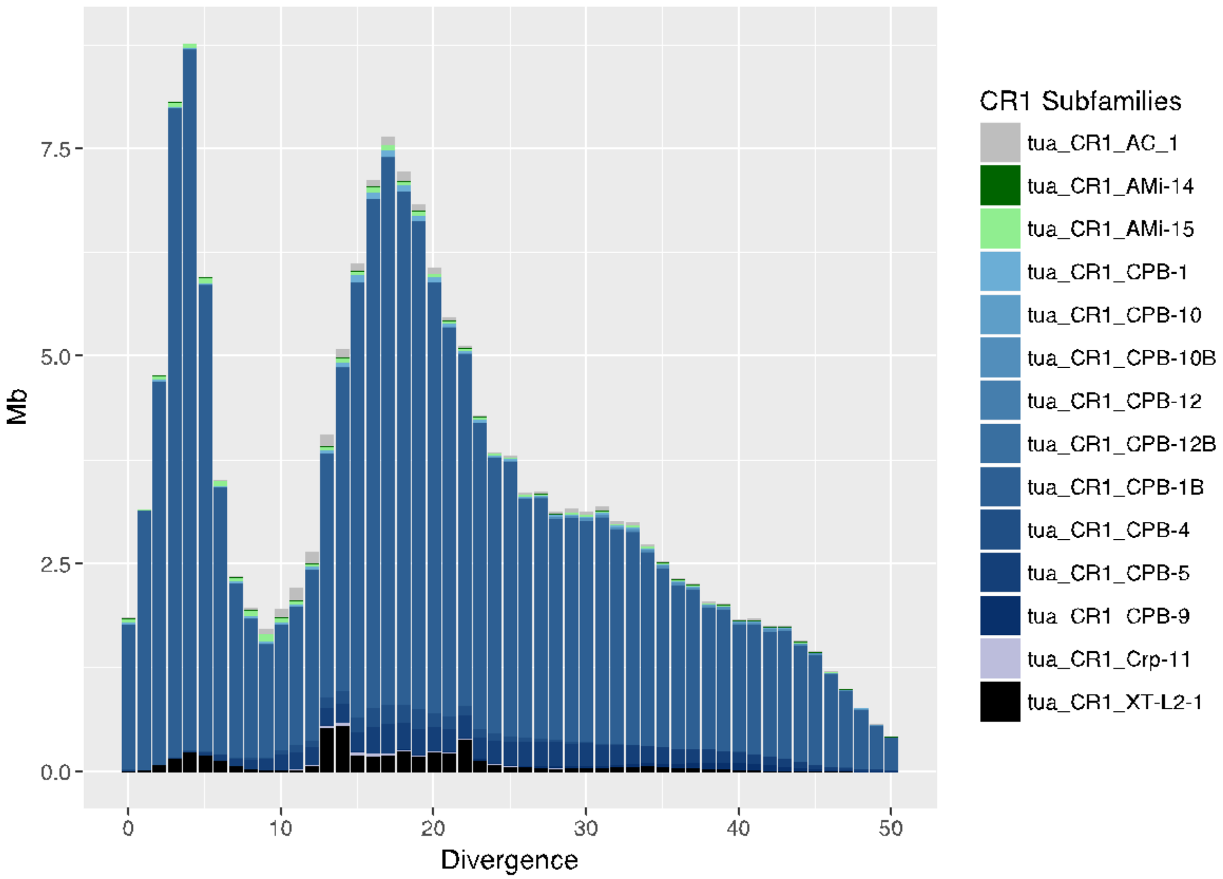

**Figure 4.9: CR1 retrotransposon landscape of the tuatara genome.** CR1 subfamilies with a total length greater than 1,000 bp and with distances to consensus <50% are shown.

##### 4.2.10 Unclassified (un-annotated) consensus sequences are probably segmental duplications

Since 216,591 of our *ab initio* consensus sequences were classified as unclassified repeats (unable to be annotated as transposable elements or gene families) and 33% of the tuatara genome was annotated using these unclassified repeats, we examined the characteristics of these un-annotated consensus sequences. Specifically, we looked at their length distribution and their copy number in the tuatara genome (Figure 4.10). As a reference, we plotted the contig lengths of the tuatara genome. Figure 4.10a shows that most of the tuatara contigs are shorter than 5 kb; the shortest contig is 800 bp and the longest 29,987,930 bp.

Figure 4.10 also shows that 92% of unclassified repeat consensus sequences were shorter than 2 kb, and the longest unclassified consensus sequence was 31,536 bp. In terms of copy number, only 0.2% of the unclassified consensus sequences were present in more than 1,000 copies. Only

12 of them were present at more than 3,000 copies, they are around 1–8 kb in length. Because most of the unclassified consensus sequences are short and present at low copy number, they are most likely the product of segmental duplications.

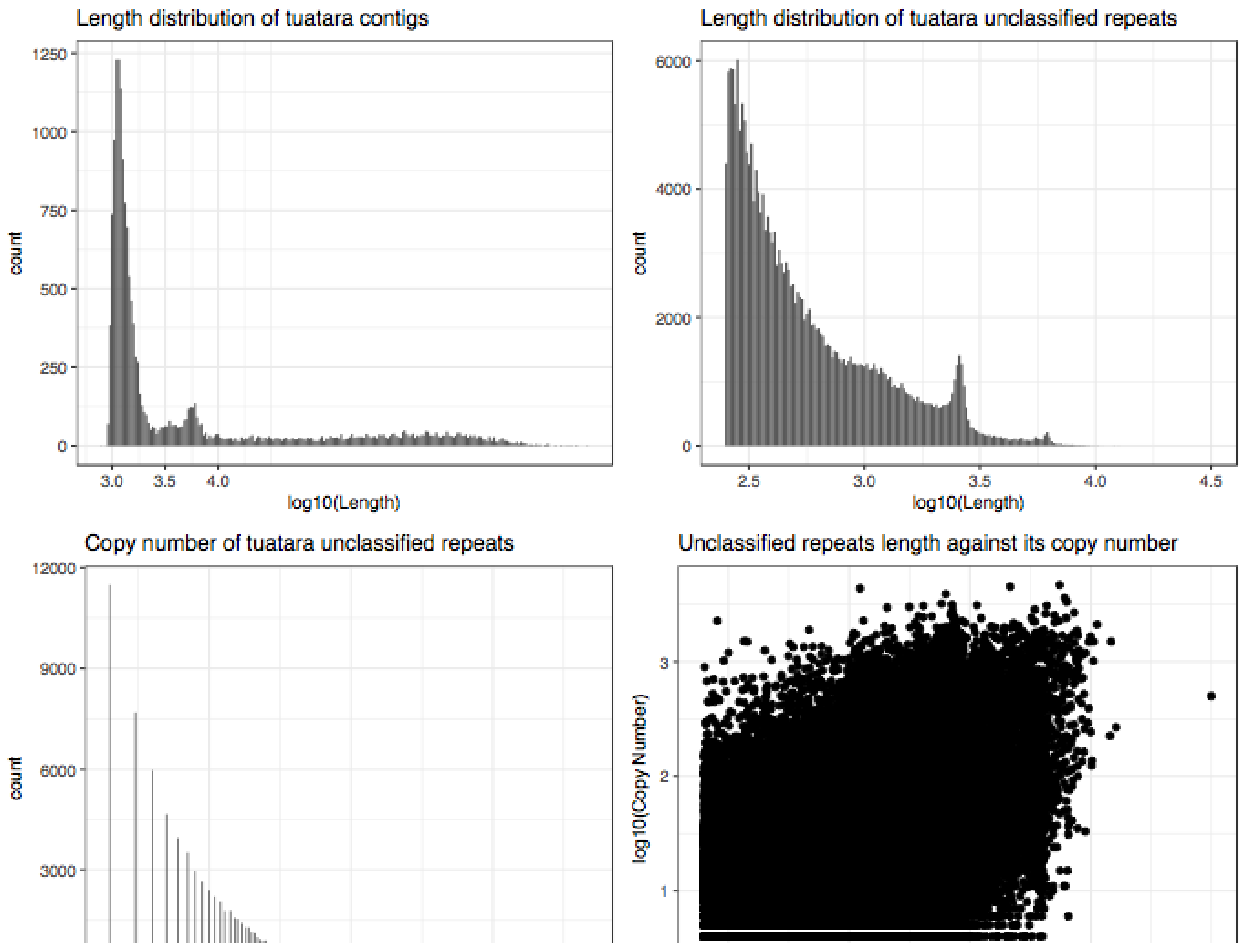

**Figure 4.10: Characteristics of unclassified consensus sequences in the tuatara genome.** a) Length distribution of tuatara contigs, transformed length with log10; b) Length distribution of tuatara unclassified repeat consensus sequences generated from

our *ab initio* method, transformed length with log10; c) Copy number distribution of unclassified repeat consensus sequences, transformed copy number with log10; d) Scatter plot of length and copy number of unclassified repeat consensus sequences.

In order to further analyse the 12 unclassified repeats with high copy number in the tuatara genome (>3,000), a coverage plot was used to visualise the copy number of genomic sequences similar to the high copy consensus (Figure 4.11). Figure 4.11 clearly shows that for most of these unclassified repeat families, significant copy number enrichment was observed in a 300–1500 bp region (>2,000 copy number), while in family 206141 (7,798 bp) and family 051465 (7,944 bp), we observed multiple peak regions with copy numbers less than 700.

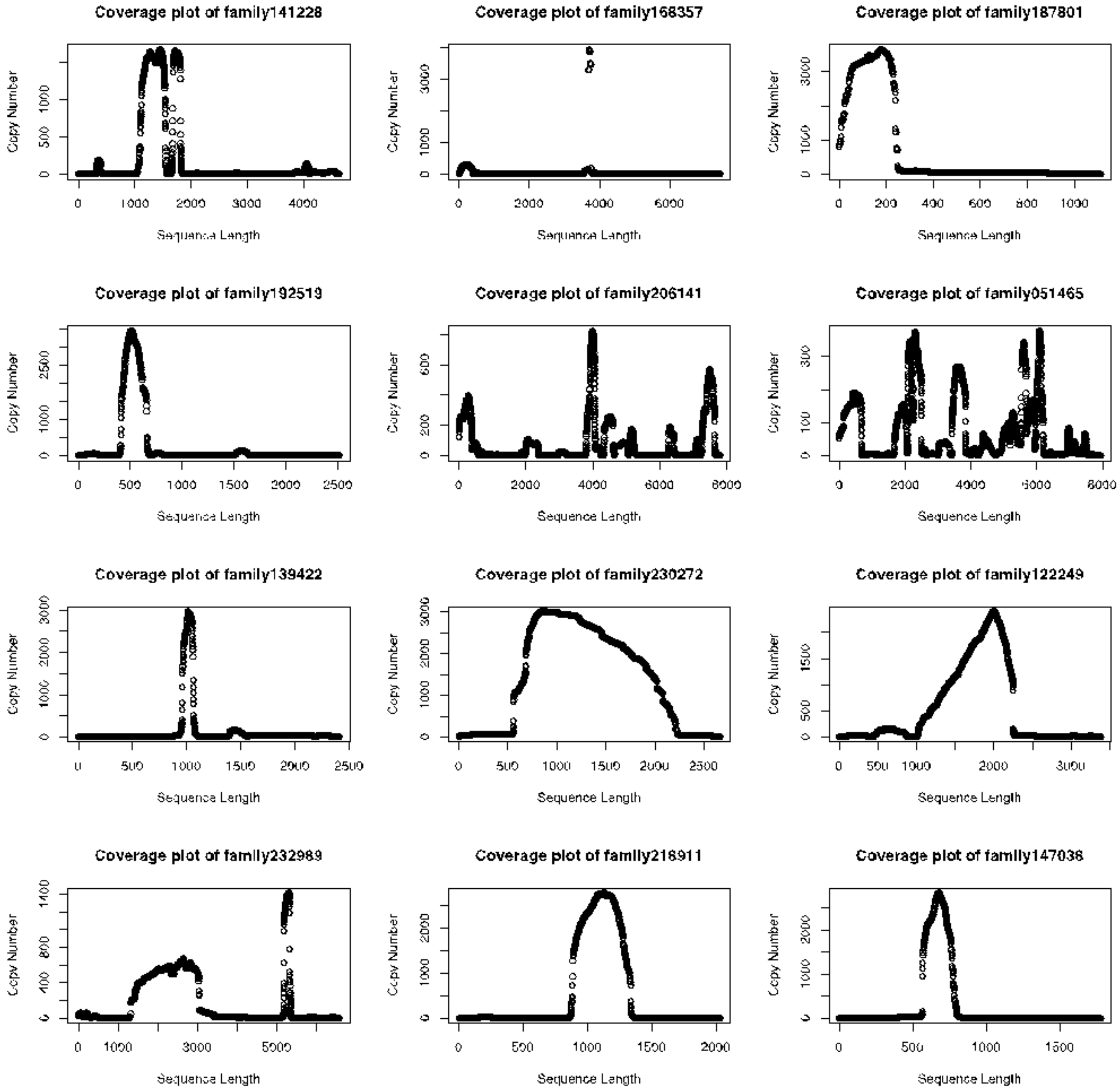
Coverage plot of family141228
Coverage plot of family168357
Coverage plot of family187801
Coverage plot of family192519
Coverage plot of family206141
Coverage plot of family051465
Coverage plot of family139422
Coverage plot of family230272
Coverage plot of family122249
Coverage plot of family232989
Coverage plot of family218911
Coverage plot of family147036
Copy Number
Sequence Length

**Figure 4.11: Coverage plots of high copy number unclassified repeats in the tuatara genome**. Overall coverage plots of 12 high copy number unclassified sequences.

Because the consensus sequences for these three families were obtained using single linkage clustering of pairwise alignments, the low baseline coverage level across the consensus indicates the approximate very low copy number of the genomic sequences used to construct the consensus. The high copy number peaks indicate the presence of highly repetitive subsequences in the consensus.

CENSOR and BLASTN were used to further characterise the consensus sub-sequences from each peak. Figure 4.12 contains outputs from both CENSOR and BLASTN that were used to annotate the peaks. In family 139422, CENSOR annotated it as a combination of different DNA transposons (Harbinger-N24 CPB, Harbinger-N33 CPB and Mariner-N5 PBa), which may indicate a potential novel DNA transposon. BLASTN annotation of the 3'end peak region was weakly similar to *Gavialis gangeticus* FAM129A non-coding regions. However, this annotation may result from a bias in the BLAST database, as there are only 2660 tuatara entries in the NCBI nr database. In family 122249, CENSOR annotated the 5′end of the coverage peak as the 5′end of hAT-N92 DR, and the 3'end of the coverage peak as the 5'end of Gyspy116-LTR DR. However, we could not find any evidence of a promoter in these two peak regions. For the rest of the unclassified sequences, we observed what seemed to be a random combination of different repeat classes. As a final test to determine if any of the high copy sub-sequences might be SINEs, we compared all peak sub-sequences with manually curated tuatara SINEs (see Supplementary Materials 5 Repeat annotation: SINEs and DNA transposons) using BLASTN (-word_size=9, -outfmt=6), but did not detect any significant similarities.

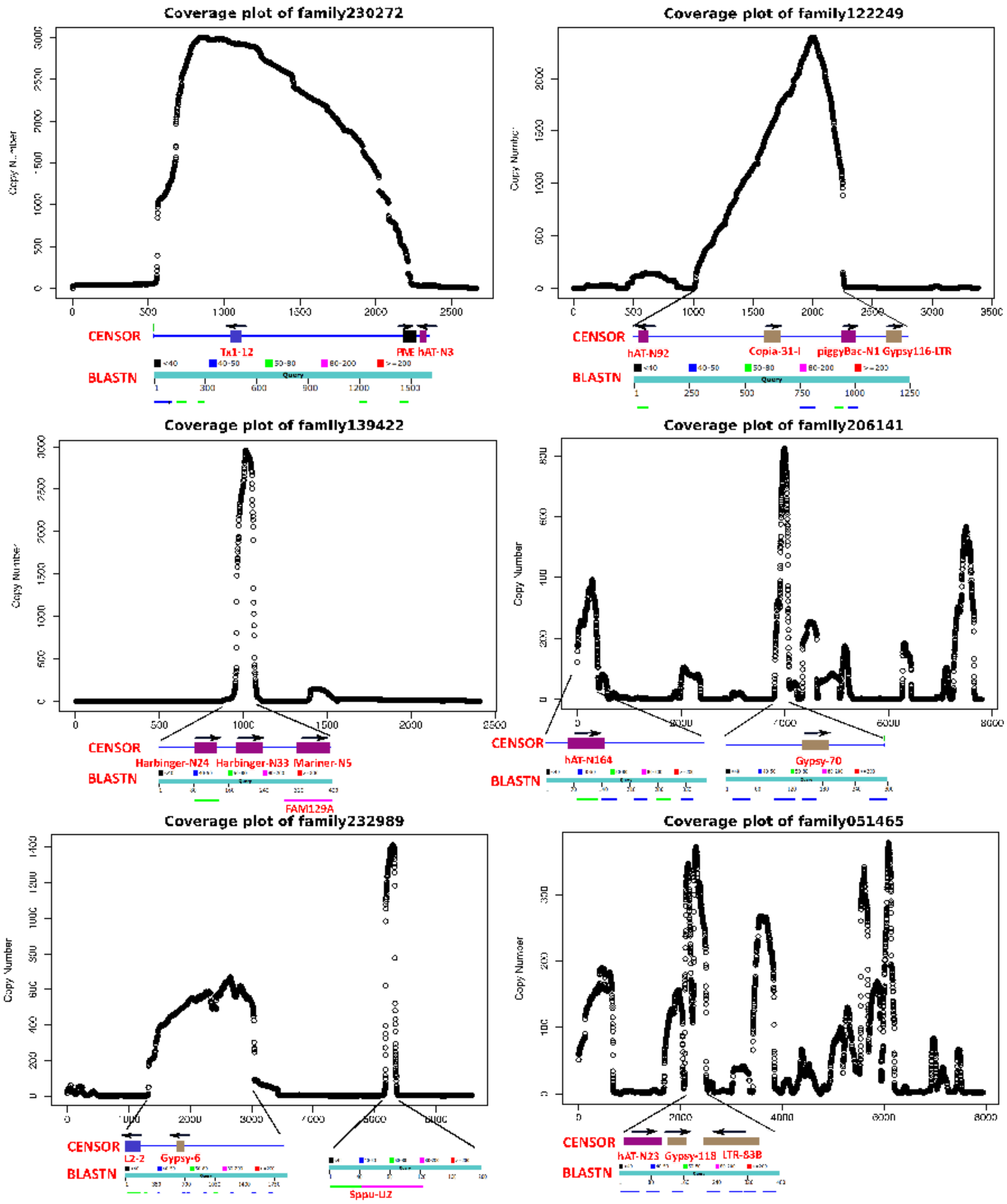
Coverage plot of family230272
Coverage plot of family122249
Coverage plot of family139422
Coverage plot of family206141
Coverage plot of family232989
Coverage plot of family051465
Copy Number
CENSOR
BLASTN
Tx1-12
PME hAT-N3
hAT-N92
Copia-31-I
piggyBac-N1 Gypsy116-LTR
Harbinger-N24 Harbinger-N33 Mariner-N5
FAM129A
hAT-N164
Gypsy-70
L2-2
Gypsy-6
Spu-U2
hAT-N23 Gypsy-118 LTR-83B
Query

**Figure 4.12: Coverage plots of the 6 high copy number unclassified repeats in the tuatara genome.** CENSOR and BLASTN annotation of the peak coverage region in unclassified family 230272, family 122249, family 139422, family 206141, family 232989 and family 051465. Black arrows represent the strand orientation of sequences based on CENSOR annotation.

In the case of unclassified family 141228 (4,531 bp), which has the highest copy number (5,497) (Figure 4.13), a promoter region was found in the middle of the peak with a score of 0.625 using the Promoter 2.0 (PolII promoter prediction tool). MEME (Multiple Em for Motif Elicitation) suite also detected a 40 bp motif around the promoter region (p-value: 1.30e-23). While both CENSOR and BLASTN has no significant hits against this peak.

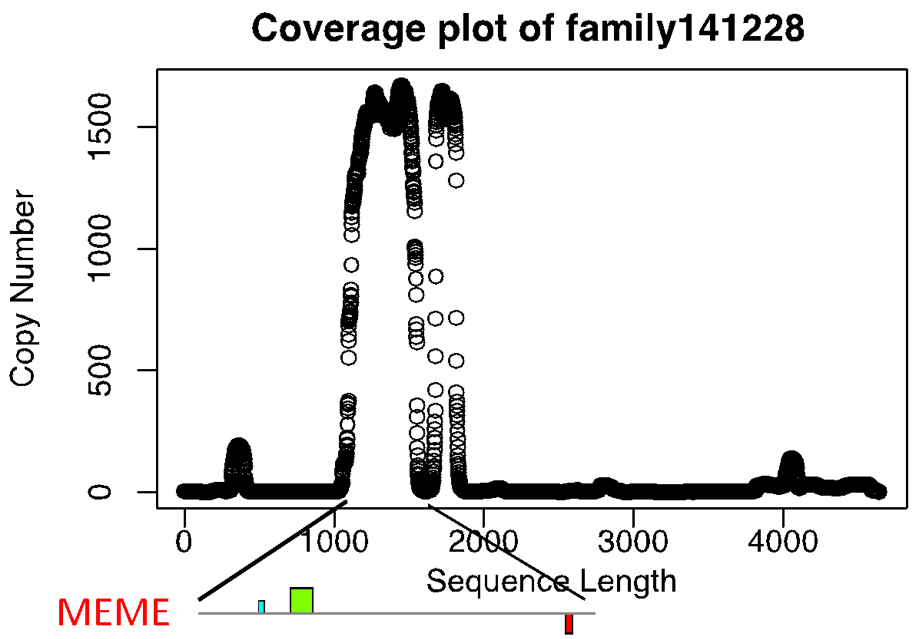

**Figure 4.13: Coverage plots of the highest copy number unclassified repeats in the tuatara genome.** MEME annotation of the peak coverage regions in unclassified family 141228.

#### 4.3 Discussion

##### 4.3.1 Segmental duplications in the tuatara genome

Compared to RepeatModeler, our method was able to classify more than one hundred times as many repetitive consensus sequences, with a broad length distribution, from 250 bp to 31,536 bp (Table 4.5). Furthermore, our *ab initio* method allowed us to identify probable segmental duplications, as 99.8% the unclassified repeat sequences from our method were present at fewer than 1,000 copies in the tuatara genome (Figure 4.10). It is worth noting that because our method identifies similar sequences with low divergence that many of the presumptive segmental duplications we found are of relatively recent origin.

Segmental duplications (SD) are usually defined as being >1 kb long and >90%[37] identical and have proven to be relevant to disease[38] and integral to studies on genome evolution[39,40]. According to previous research, segmental duplication accounts for 2.1%[41], 4.9%[42], 3.7%[43] and 6.5%[44] respectively of the Chinese alligator, green anole lizard, chicken and zebra finch genomes. While we have not characterised segmental duplications using the commonly used parameters that identify recent duplications, we found that 6.7% of the tuatara genome is present as duplicated sequences >250 bp and >94% identical. This indicates that tuatara has a greater prevalence of SD compared to other vertebrates. Furthermore, the low copy number un-annotated duplications represented 33.3% of the tuatara genome based on CENSOR analysis, indicating the presence of many older SD. It is worthwhile noting that a comparable analysis of the anole lizard genome identified a total of 12% of the genome as SD. The presence of so many SD in a vertebrate is unusual and may reflect differing drivers of genome expansion/turnover in tuatara (Figure 4.10–13).

##### 4.3.2 Significance of the LINE retrotransposons in the tuatara genome

The repeat content of reptiles and birds is lower than in mammals. The 31.3% repeat content observed in tuatara is within the bounds of previous studies where repetitive elements accounted for 23.4%[41], 27.2%[45], 10%[46], 30.4%[42], 8.5%[43] and 9.8%[44] of genome sequence in alligator, crocodile, western painted turtle, green anole lizard, chicken and zebra finch, respectively. While CR1 elements are the dominant LINEs in these genomes, L2 non-LTR LINEs are the dominant autonomous retrotransposon class in tuatara, accounting for 10% of the genome (Table 4.1). L2 are regarded as ancient repeats and most individual L2 sequences tend to be non-autonomous and truncated[47]. Less than 3% of the human genome is annotated as L2 (L2 are molecular fossils in eutheria)[48], but in platypus, L2 elements are still active and they account for 19% of the genome[49]. Because our method identifies weakly divergent repetitive elements (newly inserted elements), and because most L2 elements we found in the tuatara genome could not be identified using the Repbase library, the L2 sequences we found are most likely tuatara-specific TE of recent origin and are probably still active (Table 4.2).

Horizontal transfer (HT) is the non parent-to-offspring transmission of genetic material between individuals, a phenomenon primarily considered in a prokaryotic context. However, with assistance from a suitable vector (e.g. parasites, viruses), retrotransposons have the ability to jump between species as they do within a genome[50]. Relatively few studies have demonstrated HT of retrotransposons, including CR1s, RTEs and L1s[51–53], with no studies reporting HT for L2s. Our phylogenetic trees demonstrate that a particular class of tuatara L2 elements may have arisen from a recent transfer of platypus-like L2s into tuatara (Figures 4.1, 4.3 and 4.4), while this same class of L2 is absent from other lepidosaurs (Table 4.3 and Table 4.4).

According to the geographical distribution of fossils, monotremes are believed to have evolved in a region of Gondwanaland corresponding to Australia and Western New Guinea[54] and diverged from therian mammals about 163 to 186 million years ago[55]. In contrast, Rhynchocephalia diverged from Squamata about 220 million years ago[56] when *Sphenodon* had a very wide geographic distribution[57] before being restricted to New Zealand[58]. Because Gondwanaland broke apart around 215 to 175 million years ago, it is possible that monotremes and *Sphenodon* could have co-existed. Therefore, the platypus-like L2 elements found in tuatara are consistent with an HT event between a monotreme and sphenodon ancestor (Figure 4.5).

In conclusion, while tuatara has a transposable element content similar to other reptiles it is dominated by L2 instead of CR1 LINEs and shows evidence of recent HT and expansion of platypus-like L2 elements. The CR1 elements in the tuatara genome also showed evidence of a recent expansion of CR1 similar to those found in alligator and turtle (Figures 4.6–4.9). Tuatara also has a far higher segmental duplication content compared to other vertebrate genomes. The unusual transposable element composition and segmental duplication prevalence in the tuatara genome may indicate that drivers of genome size and complexity in tuatara differ compared to mammals, birds and other reptiles.

#### 4.4 Materials and methods

##### 4.4.1 Reference genomes: repeat identification, annotation

###### 4.4.1.1 *Ab initio* repeats identification and annotation

The tuatara draft genome assembly was used for repeat identification. All tuatara genome scaffolds were pairwise aligned using Krishna (https://github.com/biogo/examples/krishna)[59,60]

with parameters set for 94% sequence identity (-dpid) and a minimum length (-dplen) of 250 bp. The resulting alignment intervals were then used as input for igor[59,60] to define families of repeat sequences using the default parameters. Igor output was used as input for seqer[59,60] in order to generate repeat consensus sequences for each cluster/family based on MUSCLE (v3.8.31) alignments[35]. Only family members within 95% of the length of the longest family member were aligned and to avoid consensus sequence expansion due to indels in the global alignment, a maximum of 100 randomly chosen sequences/family were included in the alignment.

Identifiable repeat consensus sequences were annotated using CENSOR[61] with the Repbase 'Vertebrate' library (downloaded on 1st March, 2016, includes 41,908 sequences). Further annotation of consensus sequences was based on WU-BLAST alignment against a comprehensive retroviral and retrotransposon protein database assembled from the National Centre for Biotechnology Information[62], and against swissprot to identify known protein-coding genes from large gene families inappropriately included in the repeat set. Consensus sequences identified as either simple sequence repeats (SSRs) or protein-coding sequences, but not similar to retrotransposon or endogenous retrovirus protein-coding sequences, were removed from the consensus set. A total of 3,585 consensus sets were removed, including 3,550 that were identified as protein-coding sequences, and 35 that were identified as SSRs. After annotation of these tuatara repeat consensus sequences, CENSOR was used to map these sequences back to the tuatara genome in combination with the Repbase 'Vertebrate' library. Table 4.5 shows the comparison of repeat libraries generated using RepeatModeler[12] and our *ab initio* method[13].

**Table 4.5: Summary of *Sphenodon punctatus* repeat library metrics**

| | RepeatModeler Library | *Ab initio* Library |
|---|---|---|
| No. of consensus sequences | 2,322 | 232,111 |
| Total length (MB) | 1.3 | 171 |
| Well-annotated Consensus sequences | 901 | 15,520 |
| Unclassified Consensus sequences | 1,421 | 216,591 |
| Min./Max. Length (bp) | 36/6,350 | 250/31,536 |

#### 4.4.2 Analysis of LINE elements in the tuatara genome

In order to investigate the similarity of tuatara L2 and CR1 sequences generated from our *ab initio* method and RepeatModeler (http://www.repeatmasker.org/RepeatModeler.html), we extracted long L2 and CR1 consensus sequences (2–4 kb for L2, 2.5–4 kb for CR1) from both libraries. MUSCLE was used to carry out global alignments between the two repeat sequence sets. FastTree (v2.1.8)[63] was used to infer a maximum likelihood phylogeny from the alignment output. FigTree (v1.4.2)[64] was used to visualise and annotate the tree using repeat class labels.

##### 4.4.3 Resolving L2 classification

Many of the tuatara L2 consensus sequences were annotated as being most similar to platypus L2. This introduced an annotation problem because, according to Repbase, almost all platypus L2 consensus sequences were annotated as belonging to the CR1 clade, rather than L2 clade. In order to resolve the annotation, we needed to determine which clade platypus L2 really belong to. Full-length consensus sequences of chicken CR1 elements were extracted from Repbase, and MUSCLE was then used to carry out global alignments between CR1 elements and the 2–4 kb long L2 consensus sequences from tuatara. Alignment output was used to construct a maximum likelihood phylogeny using FastTree. FigTree was used to visualise and annotate the tree using repeat class labels.

##### 4.4.4 Dendrogram construction from LINE nucleotide sequence alignments

In order to determine the evolutionary position of tuatara L2 and CR1 within vertebrates, L2 and CR1 consensus sequences between 2–4 kb long from tuatara identified using our *ab initio* method and tuatara L2 sequences that ranged from 2–4 kb long identified using RMD were extracted. L2 and CR1 consensus sequences (2–4 kb) were also extracted from the Repbase 'Vertebrates' library. We globally aligned the resulting 222 L2 consensus sequences and 231 CR1 consensus sequences individually using MUSCLE. FastTree was used to infer a maximum likelihood phylogeny from the global alignments. Archaeopteryx v0.9901 beta was used to visualise and annotate the trees using repeat class labels.

##### 4.4.5 Phylogenetic analysis of L2 elements using RT domain sequences

All L2 and CR1 sequences were extracted from our *ab initio* consensus sequence libraries, and anole, turtle and crocodile L2 consensus sequences were extracted from the Repbase library. USEARCH[65] was then used to scan for open reading frames in L2 consensus sequences that were at least 60% of the expected length (>1.5 kb for ORF2p). After translation, ORF2p candidates were checked for similarity to known domains using HMM-HMM comparison[66] against the Pfam28.0 database[67] as of May 2015 (includes 16,230 families). ORF2p containing RT domains were extracted using the envelope coordinates from the HMMer domain hits table (–domtblout), with a minimum length of 200 amino acids. Nucleotide sequences that contained RT domains were extracted and assembled into one file (a total of 48 sequences).

Two methods were tested to describe the evolutionary dynamics of potentially active L2 elements. First, 48 RT domain nucleotide sequences within ORF2p were aligned with muscle, and then FastTree was used to infer a maximum likelihood phylogeny (-nt, -gtr). Another RT domain Phylogeny tree was inferred by using MrBayes (lse nst=6; mcmc ngen=30000 samplefreq=200 printfreq=200 diagnfreq=1000)[68] from MUSCLE alignment output. Both of the methods used a GTR model and FigTree was used to visualise and annotate the tree using repeat class labels.

##### 4.4.6 Potential horizontal transfer of L2 elements between tuatara and platypus

CENSOR was used with a custom library of platypus-like and non-platypus-like L2 consensus sequences from tuatara to find similar sequences (both full length and fragments) in five reptile genomes (anole, crocodile, alligator and bearded dragon) and one monotreme genome (platypus). To confirm the validity of hits, each hit was extracted as a nucleotide sequence and aligned with BLASTN (default parameter) against the platypus-like tuatara L2 and non-platypus like tuatara L2 consensus sequences. Hits smaller than 50 bp were discarded.

MUSCLE and PILER were used to build two super consensus sequences, one was generated from platypus-like tuatara L2, the other one was generated from non-platypus like tuatara L2. RepeatMasker was used to align each super consensus against their corresponding tuatara L2 elements, in order to calculate divergence rate using Kimura 2-parameter divergence metric, adjusted for ‘GC’ content.

##### 4.4.7 Divergence rate of CR1 elements in the tuatara genome

In order to plot the relative age profiles of CR1 insertions, tuatara CR1 consensus sequences identified using CARP (3,596 consensus sequences) were aligned and annotated against the tuatara genome by using RepeatMasker version open-4.0 (2) (-pa 32 -a -nolow -xsmall -gccalc -html -excln). Kimura distances were calculated using RepeatLandscape from the RepeatMasker alignment outfiles (-noCpG). Kimura distances are computed based on the rates of transitions and transversions between the tuatara genomic sequences and the CR1 consensus sequence, allowing estimation of the relative ages of inserted CR1 copies.

##### 4.4.8 Identification of novel repeat sequences from the tuatara genome

In order to explore the unclassified consensus sequences from our *ab initio* method, 216,591 unclassified sequences were extracted, and the R package ggplot2[69] was used to visualise their length distribution against copy number.

For high copy number (3,000 copies) families, a coverage plot was used to investigate the positional distribution of genomic sequence fragments with respect to these unclassified sequences. BLASTN and CENSOR were further used to characterise the consensus sub-sequences from the coverage peaks of 12 unclassified sequences found in the coverage plots. We scanned for potential promoters present in coverage peaks using an RNA PolII promoter prediction tool (Promoter 2.0)[70]. This was an initial attempt to find any evidence of a promoter in peak coverage regions. MEME (Multiple Em for Motif Elicitation) suite[71].

The criteria we used to identify novel repeats from unclassified repeat consensus sequences were as follows: the sequence must have copy number greater than 10,000; the sequence should be of a reasonable length for a repeat (200 bp–6 kb); the sequence cannot be identified as a repeat based on similarity to the Repbase library, and it should not be similar to any protein coding sequences (the sequences would then likely to belong to a gene family, or be part of a conserved domain).

##### 4.4.9 Problems with RMD derived consensus sequences:

When we looked at the repeat consensus sequences generated from the RMD method, we noticed that 14 out of 1149 sequences had long stretches of polyN, that parts of these sequences were annotated as unknown sequences, and parts of these sequences were annotated as transposable elements. One example is sequence rnd-3 family-287#LINE/L2 (~3,645 bp), which would appear to be present at high copy number (~16,000) in the tuatara genome.

>rnd-3_family-287#LINE/L2 (Recon Family Size = 42, Final Multiple Alignment Size = 40)

```
NTGGTGAGGTTGCAGCTCGTCTGCATACNNTCCCAGCNNCCTCGGCTGTCATCGCCCGAGGACGCCGCGACCCTCCTGGGGCTTGGCCTCCCTGCCN
CCCCAAGCTCCACCGACCAGCGGGCCGAAAAGCGCGNGCCNAAAGGCGCGCGCCTTANAGGCGCNCGCCTCACANNTACTTTTCTNCTGTACTTCCC
CTCTNCTGTACTTNCCCACTAACNTGTCTNNCCCACTAACATGTCTCGCCCTNTCTTTCCCNCCTCTCCTTCCCCTCCTCTCCTTCTCCTCTTNCCT
CCATCTTTTTCCTCCCCTNCTTTCTCCTCCCAGCCTNTTCCACCCTCTAACCTCCCCCTCTCTGCCTCCCATACATCCTTTGCTCCCTTGGCTACTT
TCTCTCGTTGCTCCCCTTCCGCTCCTGTCTCTAGCCTTCCTCGACGCTCCCCTTCTCCCAATCTTATCCCTATTCCTCCTGTCTCTCCNGCCGACCC
CGTCGCCCTCCTTCCCCTAATCTTATCCCCATTNCTTGCCTCCCCCCTCATCAACTNCCATTCTCCTGCTCCCTCTGGAACTGTCGCTCNTCTGCCA
GTAAAGCCCCCTATANCCATGANCTTTTCCTCTCTCGCTCCCTCCANCNCCTCGCNCTTACTGAAACCTGGGTCACCCCCTCNNNNNNNNNNNNNNN
NNNNNNNNNNNNNNNNNNNNNNNNNNNNNNNNNNNNNNNNNNNNNNNNNNNNNNNNNNNNNNNNNNNNNNNNNNNNNNNNNNNNNNNNNNNNNNNNN
NNNNNNNNNNNNNNNNNNNNNNNNNNNNNNNNNNNNNNNNNNNNNNNNNNNNNNNNNNNNNNNNNNNNNNNNNNNNNNNNNNNNNNNNNNNNNNNNN
NNNNNNNNNNNNNNNNNNNNNNNNNNNNNNNNNNNNNNNNNNNNNNNNNNNNNNNNNNNNNNNNNNNNNNNNNNNNNNNNNNNNNNNNNNNNNNNNN
NNNNNNNNNNNNNNNNNNNNNNNNNNNNNNNNNNNNNNNNNNNNNNNNNNNNNNNNNNNNNNNNNNNNNNNNNNNNNNNNNNNNNNNNNNNNNNNNN
NNNNNNNNNNNNNNNNNNNNNNNNNNNNNNNNNNNNNNNNNNNNNNNNNNNNNNNNNNNNNNNNNNNNNNNNNNNNNNNNNNNNNNNNNNNNNNNNN
NNNNNNNNNNNNNNNNNNNNNNNNNNNNNNNNNNNNNNNNNNNNNNNNNNNNNNNNNNNNNNNNNNNNNNNNNNNNNNNNNNNNNNNNNNNNNNNNN
NNNNNNNNNNNNNNNNNNNNNNNNNNNNNNNNNNNNNNNNNNNNNNNNNNNNNNNNNNNNNNNNNNNNNNNNNNNNNNNNNNNNNNNNNNNNNNNNN
NNNNNNNNNNNNNNNNNNNNNNNNNNNNNNNNNNNNNNNNNNNNNNNNNNNNNNNNNNNNNNNNNNNNNNNNNNNNNNNNNNNNNNNNNNNNNNNNN
NNNNNNNNNNNNNNNNNNNNNNNNNNNNNNNNNNNNNNNNNNNNNNNNNNNNNNNNNNNNNNNNNNNNNNNNNNNNNNNNNNNNNNNNNNNNNNNNN
NNNNNNNNNNNNNNNNNNNNNNNNNNNNNNNNNNNNNNNNNNNNNNNNNNNNNNNNNNNNNNNNNNNNNNNNNNNNNNNNNNNNNNNNNNNNNNNNN
NNNNNNNNNNNNNNNNNNNNNNNNNNNNNNNNNNNNNNNNNNNNNNNNNNNNNNNNNNNNNNNNNNNNNNNNNNNNNNNNNNNNNNNNNNNNNNNNN
NNNNNNNNNNNNNNNNNNNNNNNNNNNNNNNNNNNNNNNNNNNNNNNNNNNNNNNNNNNNTGAGGAAATCTCCTCTCTGCTCTCCTCCTCCAACCCTTC
CACTTGCNCCCTTGATCCCATTCCTTCCCGCCTCCTCACCTCCATTACTCCCACTGTCGCCCCTTCCTTCCTGCACATCATCAACCTCTCCCTCTCT
ACTGGCTCTTTCCCTTCTGCCTTCAAACATGCCACAGTCTCTCCTATTCTCAAAAAGTCCACTCTTGACCCCTCTGACCCTTCTAGCTATCGTCCTG
TCTCACTCCTTCCTTTTGCTTCCAAGCTGCTTGAACGTGTTGTCTACTCCCGCTGCCTTGACTTCCTATCCTCCAACTCCTCCCTTGACCCCCTCCA
```

ATCTGGCTTCCGCCCCCTTCATTCTACTGAAACTGCTCTCGCTAAGATCACCAACGATCTTCTGGTGGCTAAATCTAAACGTCTCTATTCCATTCTT
GTCCTCCTTGATCTCTCTGCTGCCTTCGATACTGTTGACCACTCACTCTTGCTGGATTCCCTTCACTCCCTTGGCTTTTGTGGCTCTGTCCATAACT
GGTTCTCCTCCTACCTCTCTGACCGCTCCTTTAATGTGTCTCTCTCCAACTCCGCCTCCTCCTCCTTCCCCCTCTCAGTTGGTGTTCCTCAAGGCTC
TGTTCTTGGCCCTCTCCTCTTCTCGATCTATACCTCATCCTTAGGCAAACTCATTGCCTCCCATGGCTTCCAATACCACCTCTATGCTGATGACACT
CAACTCTATCTCTCTACTCCCTCTCTCTCTCCCTCTGTTCAAGCTCGTCTCATCAACTGTCTCTCTGACATCTCTACCTGGATGTCTCAACGCCAAC
TCAAACTCAACATGGCTAAAACTGAACTCCTTATCTTCCCCCCTCATCCTTCCTCTCCCTCCTCACTGTCAATAACTATTGATGGAGTCACCATCCT
GCCTGTCCCCCAAGCCCGTAGCCTAGGCTTCATCTTTGACTCACCCCTTTCTTTTAAACCCTACATTGACTCTGTTGCTAAATCCTGTCGCTTCTCC
CTCCATAACATTGCTAAGATTCGCCCTTTTCTCTCTGTCTCGTCTGCCAAAACTCTTGTTCATGCTCTGGTCATCTCTCGCCTCGATTACTGTAACC
TCCTTCTCACTGGCCTTCCCCGGTCTCACCTTTCTCCCCTCATCTCTGTCCAAAACTCTGCTGCCAGGGTCATCCACCTCGCCNGTCGCTTCGACCA
TGTGCAGCCCCTCCTCTCTTCGCTTCACTGGCTCCCCCTTCCTGAACGAATCCTGCACAAGCTCCTAGTCTTCACCTATAAAGCTCTTCACAACCTC
GCTCCCCCCTACCTCTCTGCGCTTATCGCCCTCCGTACTCCAGCCCGTGCACTTCGCTCCTCCTCTGTCCCCTCTCTCGCCCTTCCACGCCTTCCCT
GCTCCCCTAAGCGCCTTCGTCCCTTCTCCCTCGCTGCCCCCCATGCCTGGAACTCCCTCCCTGTCCACATCCGCCTTGCCCCCTCCCTTCGTACCTT
CAAATCCCTCCTCAAAACCCACCTCTTCCGCTCTGCTCTGTCCGACTCTTTATCCTGCTGAGCCACTATTCCTATACCTACGATCCGTTTACCCTTA
TCCTTGCAACATGGACCTGCCTGCTGTCATTCTCATCGTAACCAAGCCAAGTACGTGCACCTGAGACCTCGCATCTTCATGAATGTTAGACCCCTTG
TCTCTCCCTCTGTTCCCCTTCCTNNGTCTCTCCCCCAGCTCCTGTCTCTCAGATTGTAAGCCCTAGGGGCAGGGACCGTGTCTATTTTATTACTTTG
TAAAGCGCCAGGTACATTGCTGGCGCTATATAAATGCTTGTTAATAATAATAATAA

However, when we annotated RMD consensus sequences using CENSOR and the Repbase library, we found some of these sequences had not been accurately annotated using RepeatMasker/RepeatModeler. Figure 4.14 below shows an example. tua2-115#DNA/hAT-AC is annotated with two different repeat classes.

| Name | From | To | Name | From | To | Class | Dir | Sim | Pos/Mm:Ts | Score |
|---|---|---|---|---|---|---|---|---|---|---|
| rnd-1_family-240#DNA/PIF-Harbinger | 292 | 418 | DNA9-53_OS | 91 | 223 | DNA | c | 0.6850 | 1.7368 | 221 |
| rnd-1_family-240#DNA/PIF-Harbinger | 1081 | 1282 | BEL-636_AA-I | 1357 | 1567 | LTR/BEL | c | 0.7310 | 2.7143 | 414 |
| rnd-1_family-240#DNA/PIF-Harbinger | 1431 | 1620 | Copia-103_AA-I | 1990 | 2183 | LTR/Copia | d | 0.7037 | 2.4211 | 400 |

#### Masked Sequence

```
>rnd-1_family-240#DNA/PIF-Harbinger
ACATCANGGAGGGAGCAGGCACCGCCCGCGGCAGCAGGGCGGAGGCGGGAGCCCACAGCGGGGCCAGCAN
CCGGGGCNCGGAAGGCACAGGGGGNCTCACAGCCAGCTGCAGGGGGCCACGAAGGGGCCCGGGGCCGGGT
GGCCCGCCAGGTGAACGAAGTGGGGCCANTGAAGCATGCCCCGGCGGTACCCCAGCACGGACATGTGGTG
GGCGACCAGGTAGTANGCCTCGAAGGGATCCTGCACGGCGAGCCCCANCTGCACATCGGAACGGGACCAC
ACACCNGACAGXXXXXXXXXXXXXXXXXXXXXXXXXXXXXXXXXXXXXXXXXXXXXXXXXXXXXXXXXXX
XXXXXXXXXXXXXXXXXXXXXXXXXXXXXXXXXXXXXXXXXXXXXXXXXXXXXXXXXXXXXXXXXXXXTG
AACGCAGCCAAAGTAGGACACGGCCAGGGCAGGGCCCCCCTCCGCCAGACAGGCCGTGTAGGCGGCCAGG
CTGCACACAGACAGGGCAGACACGGTCTCCGCCGCGGACCACGCCACATGCGAGCACCTCATCTCCGGGA
TGCGCCCATCGGGAAAGGGANNCAGGCCCAGCGTCTCCGCATGCAGCCGGCAATCGCGGGCGCTCGCCTC
ATAGCCAAAGTAGCGNAGNCCGGAGCTCAGGCTCGCATAGGCACCCGACAGGCTGTCCGGGCAGCTCAGG
AGCAATTGCACGGCCGGGGAATGCCACAGGTCACACAGGGCCATGGTCTCGCGGAAGGTCCAGCTATGGG
GGGCCCCATGGGGGCCGGGAGCCCNGNGGAAGCGCAGCCAGCCATGGTCCAGCAAGCGANGGAGACTAAC
NGCAAGGACAGGGGGGCCCCACAATCCCTGGAGGGACGGGAGGGGANGCAGGGGGGCCCCAAGGCCAGGC
GATAAGGCCACAGGCCGCAACATNGCACGGCAAACGAGCGAGGCGTGGGGTGCGGACCCAGCACAGCTGC
NGGAACNGAGCGCACAGACAGGACAGCCTACACACTAATCAGGGCGGCGAGGACAACAGGCAGCCAACAC
NGAGAGGTGGAGGTGATGACAGCAGGAATXXXXXXXXXXXXXXXXXXXXXXXXXXXXXXXXXXXXXXXXX
XXXXXXXXXXXXXXXXXXXXXXXXXXXXXXXXXXXXXXXXXXXXXXXXXXXXXXXXXXXXXXXXXXXXXX
XXXXXXXXXXXXXXXXXXXXXXXXXXXXXXXXXXXXXXXXXXXXXXXXXXXXXXXXXXXXXXXXXXXXXX
[illegible]
```

**Figure 4.14: Example of bad annotation of consensus sequence by RMD (tua2-115#DNA/hAT-AC).**

### 5 REPEAT ANNOTATION: SINES AND DNA TRANSPOSONS

Valentina Peona, Claire R. Peart, Vera M. Warmuth, Alexander Suh*

#### 5.1 Methods

Raw consensus sequences of SINE retrotransposons and DNA transposons were initially predicted de novo by RepeatModeler (version 1.0.8[12]). First, the names of all raw consensus sequences were shortened to ease downstream analyses (e.g., "rnd-1_family-1#Unknown" was renamed into "tua1-1#Unknown"). Subsequently, consensus sequences classified as SINE retrotransposons (or with sequence similarity to SINEs or small RNAs; see *Supplementary Materials 7 Tuatara genome: non-coding RNA annotation and analysis* for further details on the latter) and DNA transposons were subject to manual curation using standard procedures[72,73]. Briefly, each consensus sequence was queried against the entire tuatara draft using BLASTn (version 2.2.28+[34]), and up to 20 of the best hits were extracted with 1 kb flanks and automatically aligned to each consensus using MAFFT (version 6[74]). From these alignments, majority-rule consensus sequences were generated, carefully inspected by eye, and considered 'complete' if their 5'- and 3'-termini were flanked by unique, single-copy sequence in the alignments. The information on the termini was further used to manually estimate the size of target site duplications and to reclassify consensus sequences into TE families. This yielded manually curated consensus sequences as a proxy for each novel subfamily of 64 DNA transposons (Table 5.1) and 18 SINE retrotransposons (Table 5.2). These were grouped into families of consensus sequences (subfamilies) with sequence similarities to each other. Following the nomenclature of similar efforts in birds[75], subfamilies with sequence similarity to vertebrate repeats from Repbase[76] in CENSOR (http://www.girinst.org/censor/index.php) were named according to the name of the best hit + suffix "_tua", with the exception of MIR SINEs which were named according to their phylogeny (see below). Subfamilies with only a partial hit in CENSOR were named with the suffix "-L_tua" ("L" indicating "like"), and subfamilies with no hit in CENSOR (and newly defined tuatara MIR families, see below) were defined as novel repeat families indicated by the name prefix "tua" before the TE superfamily name (e.g., tuaMIR-1a and tuaDNA1). Subsequently, a custom library of these classified tuatara SINE and DNA subfamilies, combined with all repeat subfamilies from birds, crocodilians, turtles, and squamates available in Repbase, was used to annotate SINEs and DNA transposons in the tuatara genome assembly via RepeatMasker (version 4.0.7[77]). Final downstream analyses comprised the generation of TE landscapes (Figures 5.1–4) and a phylogenetic tree of MIR SINEs (Figure 5.5).

Both analyses followed procedures described in detail elsewhere[78]. Briefly, using the RepeatMasker ".align" output file, the Kimura 2-parameter distance between each repeat copy and its consensus sequence was calculated using the calcDivergenceFromAlign.pl script from the RepeatMasker program package under exclusion of hypermutable CpG sites. TE landscapes plots were then generated using the total amount of annotated sequence per TE consensus in 1%-bins of divergence. Furthermore, tuatara MIR SINEs were aligned with all MIR elements available in Repbase using MAFFT (E-INS-i, version 7). After manual re-alignment, maximum likelihood sequence analysis was performed in RAxML (8.0.0[79], GTRCAT model, 1,000 bootstrap replicates) on the CIPRES Science Gateway[80]. The resulting phylogenetic tree topology was used to define tuatara MIR families (Figure 5.3).

#### 5.2 Results and discussion: DNA transposons

Our manual curation of DNA elements yielded 64 novel subfamilies that are present in the tuatara genome (Table 5.1). These include 24 families (groups of subfamilies with sequence similarity to each other) which have no sequence similarity to previously known transposable elements in Repbase[76] and were thus named tuaDNA* to imply that they are tuatara-specific families. Although most of these novel families have target site duplications with 8 bp as typical for the hAT superfamily[81], most elements were non-autonomous, i.e., lacking the transposase open reading frame needed for further classification. We thus conservatively classified these families as DNA elements of an unknown superfamily (Table 5.1). RepeatMasker analysis of the tuatara genome assembly detected copies of a total of 492 DNA transposon subfamilies and the relative age distributions of the most common ones are shown as a DNA transposon landscape (Figure 5.2). At least 30 of these DNA transposon subfamilies were recently active, as each of them has copies totalling over 0.01 Mb with distances to consensus $\leq 5\%$ (Figure 5.1a). Notably, these belong to a diverse range of DNA transposon superfamilies, namely Charlie, Harbinger, hAT, Mariner, piggyBac, and Polinton/Maverick (Figure 5.1a). There even seems to be potentially ongoing DNA transposon activity, as suggested by Harbinger, hAT, piggyBac, and tuaDNA families with dozens or hundreds of copies with 0% distance to consensus (Figure 5.1b). The tuatara genome is thus not only characterized by a very high diversity of ancient and lineage-specific DNA transposon families, but also a high diversity of recently active families, which is unique among amniotes.

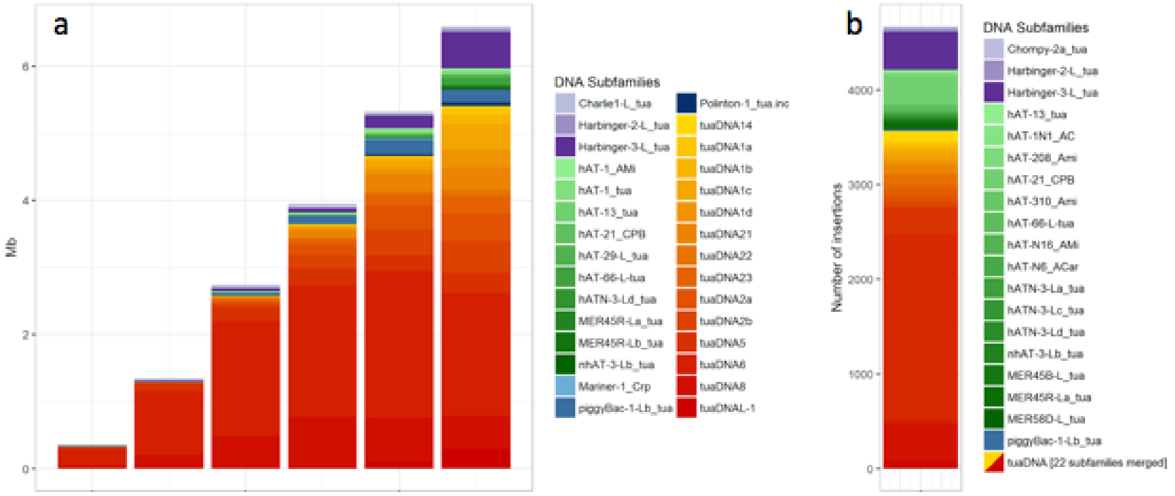

**Figure 5.1: DNA transposon landscape of the tuatara genome.** (**a**) Only DNA subfamilies that occupy more than 0.01 Mb with distances to consensus ≤ 5% are shown. (**b**) Subfamily composition of potentially active DNA transposons, i.e., copies/fragments with 0% divergence to consensus. Only the 41 DNA subfamilies with at least 10 copies in this divergence bin are shown.

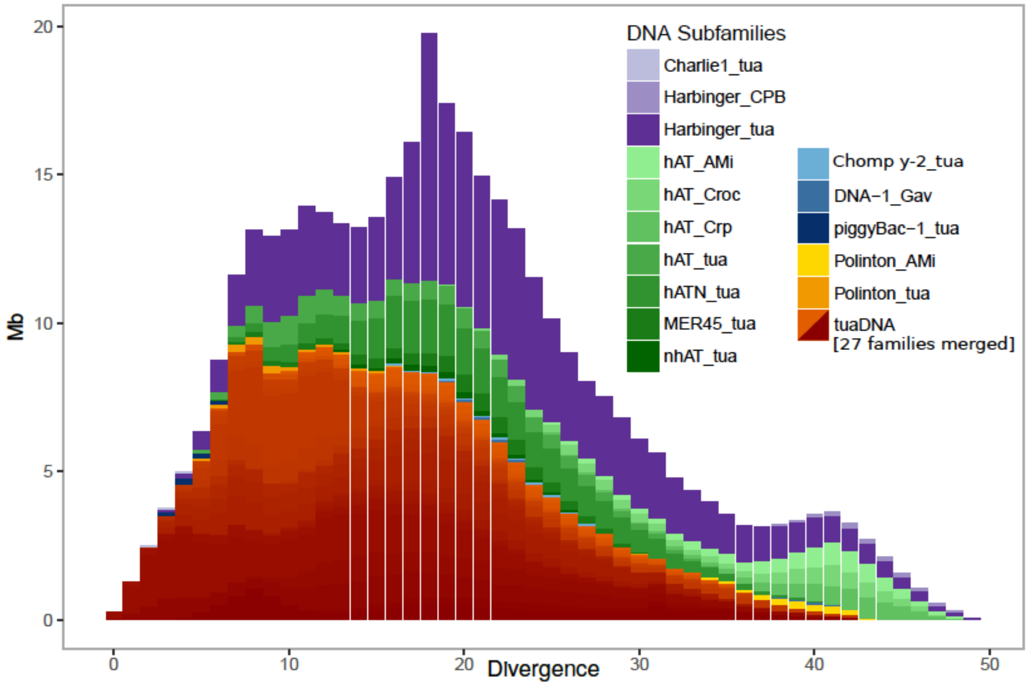

**Figure 5.2: Landscape plot of DNA transposons including 24 novel DNA transposon families from tuatara.**

#### 5.3 Results and discussion: SINE retrotransposons

Our manual curation of SINE elements yielded 18 novel subfamilies that are present in the tuatara genome (Table 5.2). Of these, 15 have L2-like tails and are therefore recognized and mobilized by the enzymatic machinery of L2 LINEs[82], and two of these have CR1-like tails and are thus mobilized by CR1 LINEs[83,84]. RepeatMasker analysis of the tuatara genome assembly detected copies of a total of 60 SINE subfamilies and the relative age distributions of the most common ones are shown as a SINE retrotransposon landscape (Figure 5.3, 5.4). At least 17 of these SINE subfamilies were recently active, as each of them has a total of over 1,000 bp of copies with distances to consensus ≤ 5% (Figure 5.3a). While ancient amniote CORE-SINE subfamilies (i.e., LFSINE[85], AmnSINE[86,87], MIR[88,89]) are present in tuatara, likely since the amniote ancestor, most SINEs are of more recent origin. Half of the SINE-derived base pairs below with ≤ 5% distance to consensus belong to a family of CR1-mobilized SINEs (tuaCR1-SINE1) and the other half falls into three families of tuatara-specific MIR elements (thus also belonging to CORE-SINEs); namely tuaMIR-a, tuaMIR-b, and tuaMIR-c (Figure 5.3, 5.4). These three MIR families consist of multiple closely related subfamilies, suggesting that the tuatara lineage underwent continuous activity and diversification of MIR elements (Figure 5.5). There even seems to be potentially ongoing MIR activity, as implied by the presence of dozens or hundreds of tuaMIR-b and tuaMIR-c copies with 0% distance to consensus (Figure 5.2b). Notably, the MIR subfamily diversity of tuatara surpasses that of all previously analyzed genomes, even that of marsupial and monotreme mammals, with millions of copies derived from relatively few MIR subfamilies[90,91].

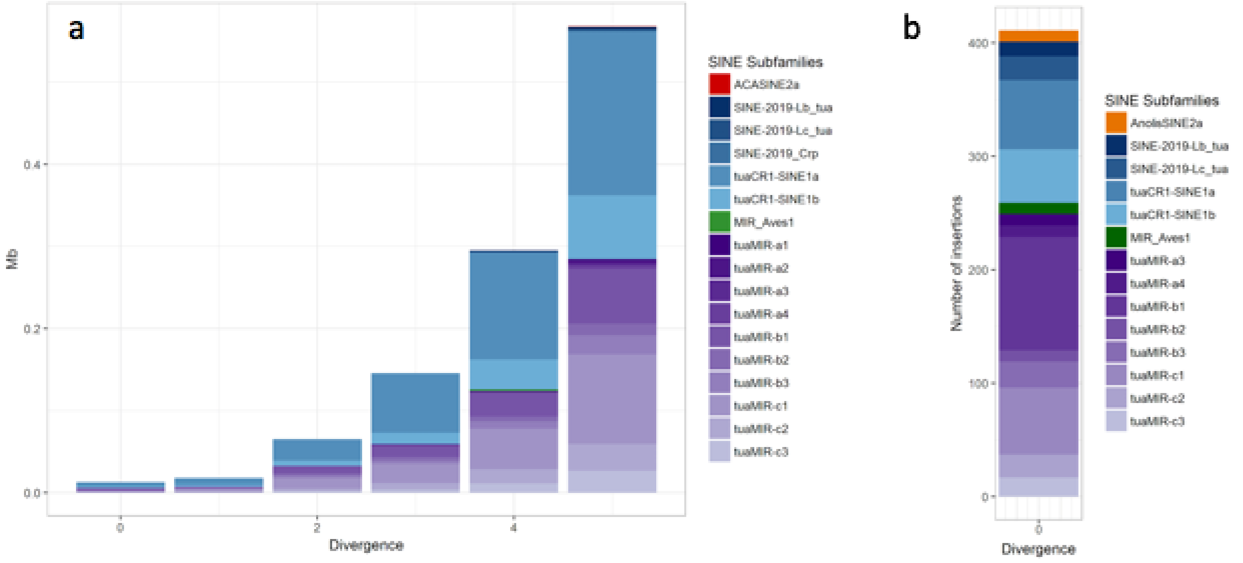

**Figure 5.3: Recent SINE retrotransposon landscape of the tuatara genome.** (**a**) Only SINE subfamilies that occupy more than 1,000 bp with distances to consensus ≤ 5% are shown. (**b**) Subfamily composition of potentially active SINE retrotransposons, i.e., copies/fragments with 0% divergence to consensus. Only the 14 SINE subfamilies with at least 10 copies in this divergence bin are shown.

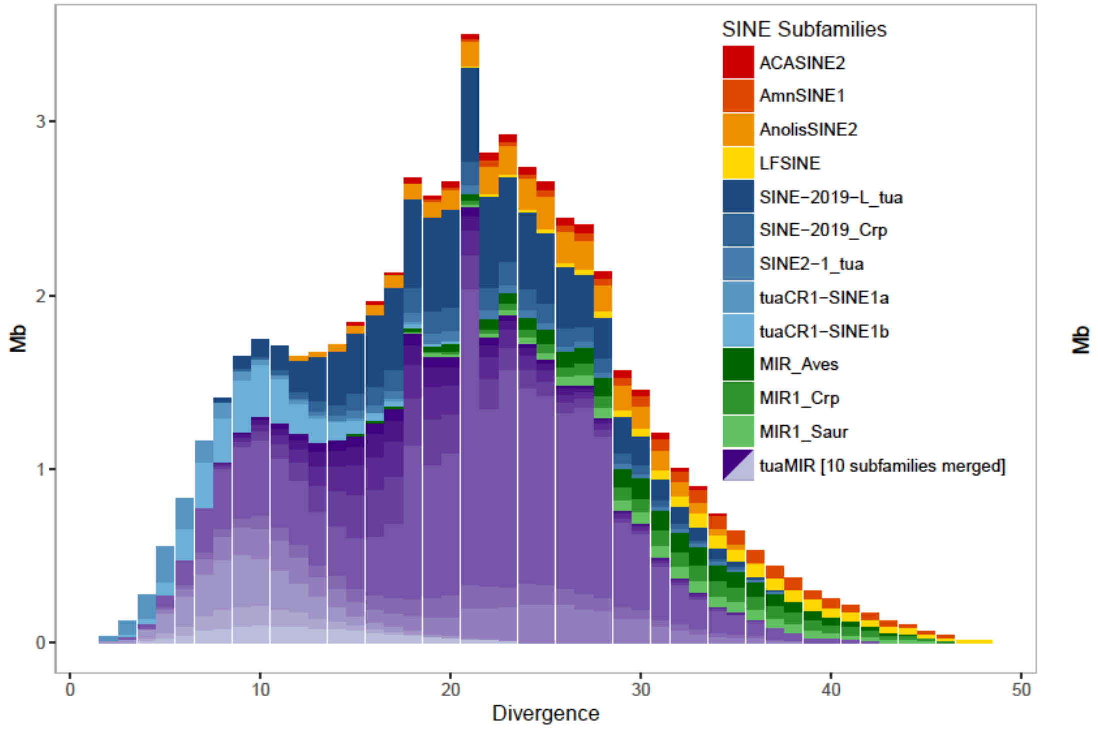

**Figure 5.4: Landscape plot of SINE retrotransposons.** The tuatara genome is dominated by MIR sequences most typically associated with mammals, with tuatara now the amniote genome in which the greatest MIR diversity has been observed.

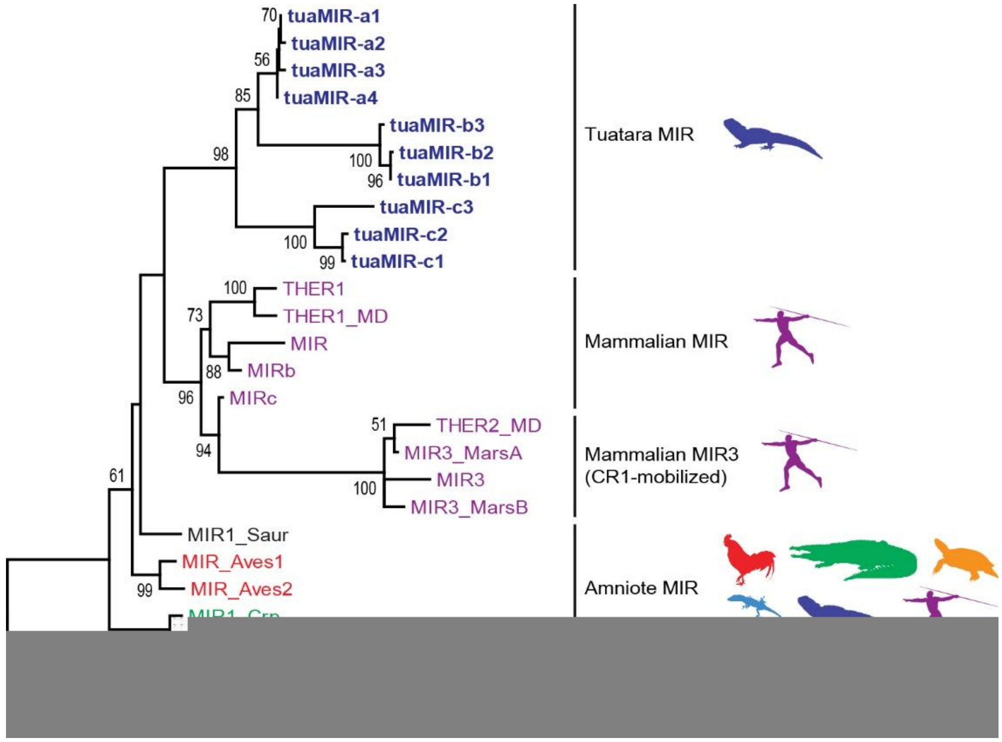

**Figure 5.5: Diversification of MIR SINEs in the tuatara lineage.** Maximum likelihood phylogram with consensus sequences colour-coded according to the genome of first description. MIR1_Saur was first described in the American alligator genome and was probably active in the common ancestor of Sauropsida [73]. The presence of low copy numbers of MIR_Aves1 and MIR_Aves2 in mammalian genomes (Alexander Suh, unpublished data) suggests that all these early-branching MIR lineages resulted from ancient sauropsid or amniote activity. Note that all MIR families have L2-like tails with the exception of mammalian MIR3 which exhibits CR1-like tails, therefore parasitizing a different superfamily of LINEs.

### Table 5.1: Characteristics of novel tuatara DNA transposons.

| Class | Subclass | Superfamily | Family | Subfamily | Similarity to Repbase repeats | Comment | Consensus status | Consensus length | TSD | TSD motif | TIR |
|---|---|---|---|---|---|---|---|---|---|---|---|
| DNA transposon | DNA | Harbinger | Chompy-2 | Chompy-2a_tua | Chompy-2_Croc (79%; Crocodylia) | Non-autonomous | Complete | 79 | 3 | | 7 |
| DNA transposon | DNA | Harbinger | Chompy-2 | Chompy-2b-tua | Chompy-2_Croc (81%; Crocodylia) | Non-autonomous | Complete | 70 | 3 | | 17 |
| DNA transposon | DNA | Harbinger | Chompy-4 | Chompy-4-L_tua | Partially Chompy-4_Croc (86%; Crocodylia) | Non-autonomous | Complete | 90 | 3 | | 43 |
| DNA transposon | DNA | Harbinger | Harbinger-2 | Harbinger-2-L_tua | Partially Harbinger-3_AMi (65%; Alligator mississippiensis) + Harbinger-2_AMi (69%; Alligator mississippiensis) | | Complete | 6627 | 3 | | 20 |
| DNA transposon | DNA | Harbinger | Harbinger-3 | Harbinger-3-L_tua | Partially Mariner-N13_HSal (83%; Harpegnathus saltator) + LTR-12_AMi (67%; Alligator mississippiensis) + Harbinger-3_AMi (64%; Alligator mississippiensis) + hAT-N11_AMi (69%; Alligator mississippiensis) + L1MA9_5 (73%; Mammalia) | Harbinger with lots of other TE fragments nested within, potentially non-autonomous | Complete | 6639 | 3 | | 20 |
| DNA transposon | DNA | hAT | hAT-1 | hAT-1_tua | hAT-1_AMi (96%; Alligator mississippiensis) | | Complete | 3504 | 8 | | 9 |
| DNA transposon | DNA | hAT | hAT-13 | hAT-13_tua | hAT-13_Ami (89%; Alligator mississippiensis) | Non-autonomous | Complete | 339 | 8 | | 12 |
| DNA transposon | DNA | hAT | hAT-39 | hAT-39_tua | hAT-39_LCh (84%; Latimeria chalumnae) except for middle part | | Complete | 1171 | 8 | | 22 |
| DNA transposon | DNA | hAT | hAT-4 | hAT-4_tua | hAT-4_Crp (85%; Crocodylus porosus) | Non-autonomous | Complete | 401 | 8 | | 14 |
| DNA transposon | DNA | hAT | hAT-N18 | hAT-N18_tua | hAT-N18_Crp (81%; Crocodylus porosus) + hAT-N18_SSa (75%; Salmo salar) | Non-autonomous | Complete | 177 | 8 | | 16 |

| | | | | | | | | | | | |
|---|---|---|---|---|---|---|---|---|---|---|---|
| DNA transposon | DNA | hAT | Charlie1 | Charlie1-L_tua | Partially CHARLIE1 (84%; Mammalia) | Contains CR1 fragment (72% similarity to CR1-C; Gallus gallus) | Complet e | 1308 | 8 | NNNTAN NN | 2 |
| DNA transposon | DNA | hAT | hAT-29 | hAT-29-L_tua | Partially hAT-29_CPB (78%; Chrysemys picta) | Non-autonomo us | Complet e | 418 | 8 | | 5 |
| DNA transposon | DNA | hAT | hAT-66 | hAT-66-L-tua | Partially hAT-66_HM (83%; Hydra magnipapillata ) | Non-autonomo us | Complet e | 920 | 8 | | 15 |
| DNA transposon | DNA | hAT | hAT-7 | hAT-7-L-tua | Partially hAT-7_CPB (76%; Chrysemys picta) + hAT-5_CPB (74%; Chrysemys picta) | Non-autonomo us | Complet e | 307 | 8 | NNNTAN NN | 14 |
| DNA transposon | DNA | hAT | hAT-N1 | hAT-N1-L_tua | Partially hAT-N1_XT (82%; Xenopus tropicalis) | Non-autonomo us | Complet e | 470 | 8 | NNNTAN NN | 12 |
| DNA transposon | DNA | hAT | hAT-N11 | hAT-N11-L_tua | Partially hAT-N11_AMi (81%; Alligator mississippiens is) + hAT-N4_Gav (77%; Gavialis gangeticus) | Non-autonomo us | Complet e | 592 | 8 | | 10 |
| DNA transposon | DNA | hAT | hAT-N18 | hAT-N18-La_tua | Partially hAT-N18_Crp (79%; Crocodylus porosus) + hAT-N27_Ssa (84%; Salmo salar) | Non-autonomo us | Complet e | 175 | 8 | NNNTAN NN | 16 |
| DNA transposon | DNA | hAT | hAT-N18 | hAT-N18-Lb_tua | Partially hAT-N18_Crp (87%; Crocodylus porosus) | Non-autonomo us | Complet e | 177 | 8 | NNNTAN NN | 16 |
| DNA transposon | DNA | hAT | hAT-1598 | hAT-1598-L_tua | Partially hAT-N22_Ami (79%; Alligator mississippiens is) + hAT-1598_Ga v (80%; Gavialis gangeticus) | Non-autonomo us | Complet e | 507 | 8 | | 11 |
| DNA transposon | DNA | hAT | hATN-3 | hATN-3-La_tua | Partially hATN-3_SM (79%; Schmidtea mediterranea) | Non-autonomo us | Complet e | 320 | 8 | NNNTAN NN | 16 |
| DNA transposon | DNA | hAT | hATN-3 | hATN-3-Lb_tua | Partially hATN-3_SM (80%; Schmidtea mediterranea) | Non-autonomo us | Complet e | 321 | 8 | NNNTAN NN | 16 |
| DNA transposon | DNA | hAT | hATN-3 | hATN-3-Lc_tua | Partially hATN-3_SM (82%; Schmidtea mediterranea) | Non-autonomo us | Complet e | 314 | 8 | NNNTAN NN | 16 |

| | | | | | | | | | | | |
|---|---|---|---|---|---|---|---|---|---|---|---|
| DNA transposon | DNA | hAT | hATN-3 | hATN-3-Ld_tua | Partially hATN-3_SM (83%; Schmidtea mediterranea) | Non-autonomous | Complete | 320 | 8 | NNNTANNN | 16 |
| DNA transposon | DNA | hAT | nhAT-3 | nhAT-3-La_tua | Partially nhAT-3_EF (81%; Eptesicus fuscus) | Non-autonomous | Complete | 386 | 8 | NNNTANNN | 14 |
| DNA transposon | DNA | hAT | nhAT-3 | nhAT-3-Lb_tua | Partially nhAT-3_EF (86%; Eptesicus fuscus) | Non-autonomous | Complete | 102 | 8 | NNNTANNN | 3 |
| DNA transposon | DNA | Maverick | Polinton-1 | Polinton-1_tua.inc | Polinton-1_CPB (73%; Chrysemys picta) | | Incomplete 5' and 3' ends | 6211 | ? | | ? |
| DNA transposon | DNA | piggyBac | piggyBac-1 | piggyBac-1-La_tua | Partially piggyBac-1_Ami (68%; Alligator mississippiensis) | Non-autonomous | Complete | 370 | 4 | TTAA | 4 |
| DNA transposon | DNA | piggyBac | piggyBac-1 | piggyBac-1-Lb_tua | Partially piggyBac-1_AMi (69%; Alligator mississippiensis) | | Complete | 1196 | 4 | TTAA | 11 |
| DNA transposon | DNA | Zator | Zator-2 | Zator-2_tua | Zator-2_HM (73%; Hydra magnipapillata) | Non-autonomous | Complete | 363 | 3 | | 26 |
| DNA transposon | DNA | Mariner | tuaMar | tuaMar1 | None | Non-autonomous | Complete | 192 | 2 | TA | 22 |
| DNA transposon | DNA | | tuaDNA1 | tuaDNA1a | None | | Complete | 1737 | 8 | | 3 |
| DNA transposon | DNA | | tuaDNA1 | tuaDNA1b | None | | Complete | 1741 | 8 | | 3 |
| DNA transposon | DNA | | tuaDNA1 | tuaDNA1c | Partially hAT-N13_Crp (70%; Crocodylus porosus) | | Complete | 1756 | 8 | | 12 |
| DNA transposon | DNA | | tuaDNA1 | tuaDNA1d | None | | Complete | 1741 | 8 | | 12 |
| DNA transposon | DNA | | tuaDNA2 | tuaDNA2a | None | | Complete | 1056 | 8 | NTAAATAG | 11 |
| DNA transposon | DNA | | tuaDNA2 | tuaDNA2b | None | | Complete | 1034 | 8 | | 12 |
| DNA transposon | DNA | | tuaDNA3 | tuaDNA3a | Partially Chompy-1_Croc (91%; Crocodylia) | Not a Harbinger due to different TSDs | Complete | 1365 | 8 | | 11 |

| | | | | | | | | | | | |
|---|---|---|---|---|---|---|---|---|---|---|---|
| DNA transposon | DNA | | tuaDNA3 | tuaDNA3b | Partially Chompy-3_Croc (87%; Crocodylia) | Not a Harbinger due to different TSDs; contains LTR fragment (75% similarity to Gypsy-50_GA-LTR; Gasterosteus aculeatus) | Complete | 1519 | 8 | | 13 |
| DNA transposon | DNA | | tuaDNA4 | tuaDNA4 | Partially Chompy-4_Crp (78%; Crocodylus porosus) | Non-autonomous, not clear if Harbinger or not due to unknown TSDs | Complete | 725 | ? | | 9 |
| DNA transposon | DNA | | tuaDNA5 | tuaDNA5 | Partially Harbinger-1B_Crp (83%; Crocodylus porosus) | Not a Harbinger due to different TSDs | Complete | 1532 | 8 | | 4 |
| DNA transposon | DNA | | tuaDNA6 | tuaDNA6 | None | Contains Mariner fragment (81% similarity to Mariner-31_HM; Hydra magnipapillata) | Complete | 1176 | 8 | | 15 |
| DNA transposon | DNA | | tuaDNA7 | tuaDNA7 | None | Non-autonomous | Complete | 563 | 3 | | 13 |
| DNA transposon | DNA | | tuaDNA8 | tuaDNA8 | None | Non-autonomous | Complete | 336 | 8 | | 15 |
| DNA transposon | DNA | | tuaDNA9 | tuaDNA9 | None | Non-autonomous | Complete | 382 | 8 | | 11 |
| DNA transposon | DNA | | tuaDNA10 | tuaDNA10 | None | Non-autonomous | Complete | 303 | ? | | 12 |
| DNA transposon | DNA | | tuaDNA11 | tuaDNA11 | None | Non-autonomous | Complete | 654 | 8 | | 10 |
| DNA transposon | DNA | | tuaDNA12 | tuaDNA12 | None | Non-autonomous | Complete | 827 | 8 | | 3 |
| DNA transposon | DNA | | tuaDNA13 | tuaDNA13 | None | Non-autonomous | Complete | 860 | 8 | | 10 |
| DNA transposon | DNA | | tuaDNA14 | tuaDNA14 | None | Non-autonomous | Complete | 738 | 8 | | 18 |
| DNA transposon | DNA | | tuaDNA15 | tuaDNA15 | None | Non-autonomous | Complete | 368 | 8 | | 15 |
| DNA transposon | DNA | | tuaDNA16 | tuaDNA16 | None | Non-autonomous | Complete | 499 | ? | | 19 |
| DNA transposon | DNA | | tuaDNA17 | tuaDNA17 | None | Non-autonomous | Complete | 468 | 8 | | 7 |
| DNA transposon | DNA | | tuaDNA18 | tuaDNA18 | None | Non-autonomous | Complete | 689 | 8 | | 16 |

| | | | | | | | | | | | |
|---|---|---|---|---|---|---|---|---|---|---|---|
| DNA transposon | DNA | | tuaDNA19 | tuaDNA19 | None | Non-autonomous, contains Mariner fragment (75% similarity to MARINERNA8_MD; Monodelphis domestica) | Complete | 264 | 8 | | 8 |
| DNA transposon | DNA | | tuaDNA20 | tuaDNA20 | None | Non-autonomous, contains SINE head (78% similarity to MIR1_Crp; Crocodylus porosus) | Complete | 423 | 8 | | 15 |
| DNA transposon | DNA | | tuaDNA21 | tuaDNA21 | None | | Complete | 1382 | 8 | | 14 |
| DNA transposon | DNA | | tuaDNA22 | tuaDNA22 | None | | Complete | 1185 | 8 | | 4 |
| DNA transposon | DNA | | tuaDNA23 | tuaDNA23 | None | | Complete | 1031 | 8 | | 9 |
| DNA transposon | DNA? | | tuaDNAL | tuaDNAL-1 | None | Contains Mariner fragment (67% similarity to Mariner-7N1_CPB; Chrysemys picta) | Complete | 2641 | ? | | 3 |
| DNA transposon | DNA | | MER45 | MER45B-L_tua | Partially MER45B (80%; Homo sapiens) | Non-autonomous | Complete | 889 | 8 | | 15 |
| DNA transposon | DNA | | MER45 | MER45R-La_tua | Partially MER45R (81%; Homo sapiens) | | Complete | 1655 | ? | | 14 |
| DNA transposon | DNA | | MER45 | MER45R-Lb_tua | Partially MER45R (83%; Homo sapiens) | Non-autonomous | Complete | 242 | 8 | | 14 |
| DNA transposon | DNA | | MER45 | MER45R-Lc_tua | Partially MER45R (85%; Homo sapiens) | Non-autonomous | Complete | 251 | 8 | | 14 |
| DNA transposon | DNA | | MER58 | MER58D-L_tua | Partially MER58D (84%; Mammalia) | Non-autonomous | Complete | 668 | 8 | | 16 |
| Retrotransposon | LTR | ERVK? | tuaLTR1 | tuaLTR1 | None | Solo-LTR | Complete | 511 | 5 | | 3 |
| Retrotransposon | LTR | Gypsy | LTR-58 | LTR-58-L_tua | Partially LTR-58_Gav (71%; Gavialis gangeticus) | Solo-LTR | Complete | 1008 | 5 | | 6 |

### Table 5.2: Characteristics of novel tuatara SINE retrotransposons.

| Class | Subclass | Superfamily | Family | Subfamily | Similarity to Repbase repeats | Comment | Consensus status | Consensus length | TSD | TSD motif | TIR |
|---|---|---|---|---|---|---|---|---|---|---|---|
| Retrotransposon | SINE | MIR | MIR1 | MIR1_tua | MIR1_Crp (96%; Crocodylus porosus) | Very short element, core region only | Complete | 55 | var | | N/A |
| Retrotransposon | SINE | MIR | tuaMIR | tuaMIR-a1 | THER1 (80%; Mammalia) | | Complete | 235 | var | | N/A |
| Retrotransposon | SINE | MIR | tuaMIR | tuaMIR-a2 | THER1 (80%; Mammalia) | | Complete | 238 | var | | N/A |
| Retrotransposon | SINE | MIR | tuaMIR | tuaMIR-a3 | THER1 (79%; Mammalia) | | Complete | 245 | var | | N/A |
| Retrotransposon | SINE | MIR | tuaMIR | tuaMIR-a4 | THER1 (79%; Mammalia) | | Complete | 244 | var | | N/A |
| Retrotransposon | SINE | MIR | tuaMIR | tuaMIR-b1 | Partially SINE2-1_XT (78%; Xenopus tropicalis) | Different SINE head (82% similarity to SINE2-2_ACar ; Anolis carolinensis) | Complete | 295 | var | | N/A |
| Retrotransposon | SINE | MIR | tuaMIR | tuaMIR-b2 | Partially SINE2-1_XT (78%; Xenopus tropicalis) | Different SINE head (79% similarity to MIR3_MarsB; Marsupialia) | Complete | 274 | var | | N/A |
| Retrotransposon | SINE | MIR | tuaMIR | tuaMIR-b3 | Partially SINE2-1_XT (75%; Xenopus tropicalis) | Different SINE head (79% similarity to MIR1_AMi; Alligator mississippiensis) | Complete | 296 | var | | N/A |
| Retrotransposon | SINE | MIR | tuaMIR | tuaMIR-c1 | MIR_Aves1 (76%; Aves) | Potentially different SINE head | Complete | 249 | var | | N/A |
| Retrotransposon | SINE | MIR | tuaMIR | tuaMIR-c2 | MIR_Aves1 (75%; Aves) | Potentially different SINE head | Complete | 241 | var | | N/A |
| Retrotransposon | SINE | MIR | tuaMIR | tuaMIR-c3 | Partially MIR (93%; Mammalia) | Different SINE head (79% similarity to SINE2-2_ACar ; Anolis carolinensis) | Complete | 269 | var | | N/A |
| Retrotransposon | SINE | tRNA | SINE2-1 | SINE2-1_tua | SINE2-1_Croc (79%; Crocodylia) | | Complete | 266 | var | | N/A |
| Retrotransposon | SINE | tRNA | tuaCR1-SINE | tuaCR1-SINE1a | Partially Squam2 (81%; Squamata) CR1-15_Ami (81%; Alligator mississippiensis) | tRNA-Ala head, CR1 tail | Complete | 284 | var | | N/A |

| | | | | | | | | | | | |
|---|---|---|---|---|---|---|---|---|---|---|---|
| Retrotransposon | SINE | tRNA | tuaCR1-SINE | tuaCR1-SINE1b | Squam2 (83%; Squamata) | tRNA-Ala head, CR1 tail | Complet e | 270 | var | | N/A |
| Retrotransposon | SINE? | L2 | SINE-2019 | SINE-2019-La_t ua | Partially SINE-2019_Cr p (69%) | Different SINE head | Complet e | 457 | var | | N/A |
| Retrotransposon | SINE? | L2 | SINE-2019 | SINE-2019-Lb_t ua | Partially SINE-2019_Cr p (69%; Crocodylus porosus) | L2 tail (82% similarity to L2-2_Croc; Crocodylia) | Complet e | 885 | var | | N/A |
| Retrotransposon | SINE? | L2 | SINE-2019 | SINE-2019-Lc_t ua | Partially SINE-2019_Cr p (70%) | Different SINE head | Complet e | 738 | var | | N/A |
| Retrotransposon | SINE? | L2 | SINE-2019 | SINE-2019-Ld_t ua | Partially SINE-2019_Cr p (76%; Crocodylus porosus) | Different SINE head | Complet e | 315 | var | | N/A |

### 6 REPORT ON THE DIVERSITY OF LTR RETROELEMENTS IN THE GENOME OF *SPHENODON PUNCTATUS*

Jose Horacio Grau*

#### 6.1 Introduction

Transposable elements are generally classified into Class I elements (called retrotransposons or retroelements), which use an RNA intermediate for transposition in a copy paste fashion; and Class II elements, which replicate without an RNA intermediate, either by a cut-and-paste mechanism (DNA transposons), by rolling circle DNA replication (helitrons), or by so far unknown mechanisms (politrons/mavericks). Among the Class I elements two major subclasses are recognized: (1) retroelements (REs) with long terminal repeats (LTRs) and (2) elements without LTRs (non-LTR REs)[92,93]. LTR REs, can be classified into four major families, namely Bel/Pao, Ty1/Copia, Ty3/Gypsy, and retroviruses[94,95]. A common LTR retrotransposon typically encodes two polyproteins, termed GAG and POL. The capsid protein (GAG) usually contains matrix, capsid, and nucleocapsid domains; POL consists of aspartic proteinase (AP), reverse transcriptase (RT), ribonuclease (RN), and integrase (INT) domains, the latter three (RT, RN, INT) are responsible for retrotranscribing cDNA from RNA intermediates and inserting it into the host genome. Endogenous retroviruses constitute a specific class of LTR REs that sometimes additionally contain an open reading frame (ORF) for an envelope protein (ENV), which enables ERVs to move from one cell to another. In contrast, all other LTR REs either lack or contain a remnant of an ENV gene and can only reinsert into their own host genome[96–98]. There are, however, ERVs that secondarily lost their ENV gene and thus their infectious ability. Such ERVs are retrotransposing instead of infecting other cells like typical retroviruses[99].

As a precondition for understanding the role of LTR retroelements in shaping animal genomes the diversity of these elements has to be systematized[100–102]. Several computer programs have been developed to automatically detect LTR REs[103]. Some of these computing methods have made it possible to detect and identify previously unknown elements[104] and there is an increasing body of work that suggests many genomes still host remnants of inactive retrotransposons corresponding to ancient retrotransposition events. Here we extend these analyses to explore LTR RE diversity in the tuatara genome.

#### 6.2 Results

We identified approximately 7,500 full-length long terminal repeat (LTR) retroelements including endogenous retroviruses (ERVs), which we classified into at least 12 groups using phylogenetic reconstructions of their retrotranscriptase domains (Figure 6.1). We identified nearly 450 ERVs from five major retroviral clades and some 5,500 Ty3/Gypsy elements representing five major clades. Notably, a Ty1/Copia element (Mtanga-like) is abundant in the tuatara genome with more than 1,140 full-length elements. The general spectrum of tuatara LTR retroelements is comparable to that of other sauropsids, but lacks Bel/Pao LTR retroelements which are present in anole[105]. An unexpected result is the presence of at least 37 complete spumaretroviruses in the tuatara genome (Figure 6.2).

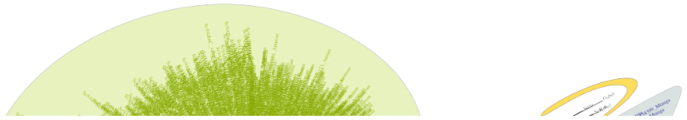

**Figure 6.1 The tuatara genome contains ~7500 full-length long terminal repeat (LTR) retroelements including nearly 450 endogenous retroviruses (ERVs).** The general spectrum of tuatara LTR retroelements is comparable to that of other sauropsids, but lacks Bel/Pao LTR retroelements which are present in anole, and has an abundance of Ty3/Gypsy retroelements.

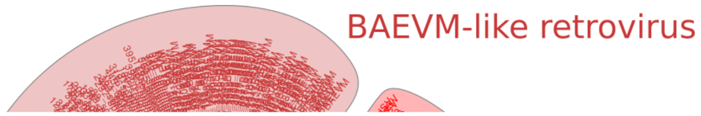

**Figure 6.2 Maximum likelihood phylogram of endogenous retroviruses found in the tuatara genome.** Colour-coded according to the retrovirus family detailed in the phylogram. No Betaretrovirus ERVs were found in the tuatara genome.

#### 6.3 Discussion and conclusions

The general spectrum of LTR-Retroelements with the genome of the tuatara is comparable to that of other Sauropsids, in that it contains abundant LTR-retroelements typically expanded in both Lepidosaurs and Archeosauronmorphs. An unexpected result is the presence of an old spumaretrovirus clade within the tuatara genome. Spumaretroviruses are considered among the most ancient ERVs[106,107] and were previously found only in coelacanth[108], sloth[109], aye-aye[110] and Cape golden mole[111]. Thus, the occurrence of several full-length endogenous spumaretroviruses in tuatara, some encoding a complete envelope gene, is the first in a sauropsid genome.

#### 6.4 Methods

LTR RE searches used a reference collection of retroelement domains and alignments obtained from the publicly available Gypsy Database 2.0 (GyDB)[102]. For the detection of LTR REs the retro-transcriptase (RT) domain was used because it is the best conserved through evolutionary time[112]. In order to obtain a custom representation of the LTR RE diversity the program suffixerator, which is part of GenomeTools[113] was applied with default parameters and created an enhanced suffix file which was later scanned with LTR harvest[14] with default parameters to predict putative LTR REs through the detection of their LTRs. To leave out the false LTR RE predictions made by LTR harvest, we then searched each LTR harvest predicted sequence against a database of RT domains of GyDB using blastx. Matches with an e-value of 1e-30 and alignments for only the best match were reported. Since blastx has the ability to produce full-length amino acid alignments from nucleotide sequences with little penalty for frameshifts, a bioperl script was used to parse out putative protein alignments of the RT domain from blastx report. Because this approach yielded thousands of RT alignments for a single genome, we clustered the LTR RE dataset using the program CD-HIT[9,114] with an identity threshold of 80%, and discarded sequences shorter than 120 aa to reduce the high number of similar and identical copies of each retroelement.

The non-redundant dataset was fused with the complete RT domains of GyDB and aligned and used to infer a series of maximum-likelihood (ML) and neighbour-joiningNJ) trees in order to accurately place the retroelements in a phylogenetic context. All alignments were conducted with the program Mafft[115] using local alignment and a Blosum 30 aa substitution matrix as parameters. Final alignment files were prepared by removing columns with more than 70% of gaps. ML trees were inferred with RaxML 8.2[116] using the PROTCATRTREV protein model of evolution, NJ trees were inferred with NINJA 1.2.2[117]. Both ML and NJ trees essentially yielded the same results.

### 7 TUATARA GENOME: NON-CODING RNA ANNOTATION AND ANALYSIS

Paul P. Gardner* and James M. Paterson

#### 7.1 Introduction

Non-coding RNAs (ncRNAs) perform vital roles in translation (e.g. ribosomal RNAs, transfer RNAs), messenger RNA maturation (e.g. spliceosomal RNAs) and gene regulation (e.g. microRNAs)[118,119]. Yet the annotation of ncRNAs is complicated by the fact that they tend to conserve complex secondary structures more than primary sequence. To address this problem we use covariance models (CMs) that account for both covarying base-paired sites and sequence conservation[120,121]. These have proven to have a major accuracy advantage over alternative methods for ncRNA gene annotation[122].

#### 7.2 Methods

To annotate ncRNAs in the tuatara genome (version 30Sep2015_rUdWx) we used CMs from the RNA families (Rfam) database (version 13.0)[20] and tRNAscan-SE (version 1.3.1)[22,123]. Overlapping annotations resulting from deep homology were resolved using clan annotations from Rfam[124] as previously described[125]. In brief, in cases of overlapping tRNA and tRNAscan-SE annotations, the more specific tRNAscan-SE results were used, if families from the same clan overlap, the family with the highest is used in a winner-takes-all approach.

The Anolis genome (version AnoCar2.0) was annotated using the same methodology. Comparisons between ncRNA copy numbers encoded in the Anolis and tuatara genomes give an indication of expansions and contractions of different families since these reptiles last shared a common ancestor (~250 million years ago).

High genomic copy-number families were assessed for likelihood of incorporation into repeat-generating machinery (e.g. retrotransposons)[82]. See *Supplementary Materials 5 Repeat annotation: SINES and DNA transposons* for further details.

### 7.3 Figures

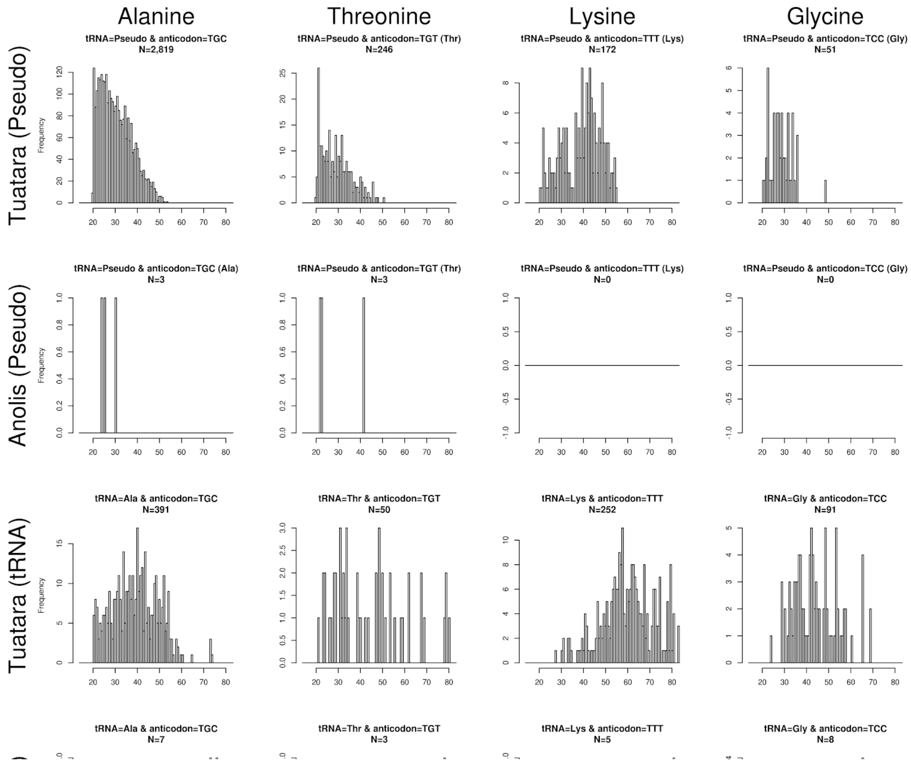

**Figure 7.1: Major expansions of alanine, threonine, lysine and glycine tRNA isotypes.** The bitscore distributions for functional and pseudogenised genomic copies are shown for both tuatara and Anolis paralogues. Bitscores give an indication of similarity to an idealised functional tRNAs.

### 7.4 Tables

**Table 7.1: Genomic copy numbers of RNA families and broad RNA types for annotated tuatara and Anolis genomes.** Only the high copy-number (N>10) ncRNA families have been included.

| ncRNA type | ncRNA family | Anolis genomic copy-number | Tuatara genomic copy-number |
|---|---|---|---|
| **Cis-regulatory elements** | | 110 | 196 |
| | Histone 3\` UTR stem-loop | 67 | 144 |
| **ncRNA genes** | | 362 | 66 |
| | 7SK RNA | 307 | 18 |
| **lncRNAs** | | 14 | 14 |
| **miRNAs** | | 290 | 319 |
| | mir-598 | 65 | 18 |
| **rRNAs** | | 1640 | 381 |
| | 5S rRNA | 1596 | 85 |
| | LSU rRNA | 29 | 229 |
| | SSU rRNA | 15 | 67 |
| **C/D box snoRNAs** | | 108 | 140 |
| **H/ACA box snoRNAs** | | 73 | 90 |
| **scaRNA** | | 13 | 16 |
| **Spliceosomal RNAs** | | 264 | 162 |
| | U1 | 38 | 22 |
| | U2 | 128 | 23 |
| | U6 | 67 | 50 |
| **tRNA** | | 375 | 3341 |

**Table 7.2: Genomic copy numbers of selected tRNA isotypes (functional and pseudogenised copies, as predicted by tRNAscan-SE).** Only high copy-number (∑N>100) isotypes have been reported.

| | | Anolis | | Tuatara | |
|---|---|---|---|---|---|
| **Isotype** | **Anticodon** | **#tRNAs** | **#Pseudogenes** | **#tRNAs** | **#Pseudogenes** |
| Thr | AGT | 4 | 5 | 90 | 23 |
| Met | CAT | 13 | 0 | 90 | 7 |
| Lys | CTT | 7 | 2 | 148 | 117 |
| Ser | GCT | 6 | 15 | 66 | 42 |
| Ser | GGA | 1 | 207 | 1 | 7 |
| His | GTG | 5 | 20 | 75 | 28 |
| Val | TAC | 2 | 1 | 18 | 112 |
| SeC/SeC(e) | TCA | 7 | 0 | 22 | 0 |
| Gly | TCC | 8 | 0 | 91 | 51 |
| Ala | TGC | 7 | 3 | 391 | 2819 |
| Pro | TGG | 4 | 6 | 61 | 142 |
| Thr | TGT | 3 | 3 | 50 | 246 |
| Glu | TTC | 9 | 2 | 84 | 57 |
| Lys | TTT | 5 | 0 | 252 | 172 |

### 8 THE nCpG DISTRIBUTION OF THE TUATARA GENOME PERMITS REVISITING THE PATTERNS AND EVOLUTION OF CpG CONTENT IN VERTEBRATES

Valeria Velásquez Zapata, Zhiqiang Wu, Nicole Valenzuela*

#### 8.1 Introduction

DNA methylation is an epigenetic modification common in animals where methyl groups are added to cytosine nucleotides[126], thus altering gene expression by preventing transcription factor binding or favouring repressor binding without changing the DNA sequence[127–129]. Because DNA methylation participates in development, disease, and mediates responses to environmental inputs[130–133], increasing efforts are devoted to understand its function and evolution. Vertebrate DNA methylation targets CpG dinucleotides across genic and intergenic regions[134], such that the distribution of CpG within genomes can impact DNA methylation and its regulatory consequences[135]. The normalized CpG content (or nCpG) is used as an *in silico* proxy of DNA methylation. This nCpG measures the ratio of the observed to expected abundance of CpG dinucleotides in a particular DNA sequence given the frequency of cytosines (C) and guanines (G) present in the genome [CpG(O/E) = CpG observed/expected][136]. 5-methylcytosine tends to deaminate spontaneously, thus mutating to thymine and reducing the nCpG content overtime, rendering nCpG an index of historic methylation[137–140].

The genomic distribution of nCpG content has been studied in several vertebrates, including human and other primates, opossum, platypus, chicken, turtle, lizard, frog, fish, and in tunicates (a chordate)[135,141,142]. These comparative studies of nCpG revealed commonalities and differences across vertebrate lineages, particularly between painted turtles and other chordates[142]. However, because painted turtle is the only vertebrate with temperature-dependent sex determination (TSD) whose nCpG has been studied thus far while all others have genotypic sex determination (GSD), it is difficult to discern if the differences observed in turtle are due to its sex-determining mechanism or to lineage specific effects. Here we analyze the nCpG distribution of the newly sequenced tuatara genome (a TSD reptile) and other TSD and GSD reptiles (some not previously studied) plus chicken and human (Figure 8.1), to illuminate the evolution of vertebrate nCpG content.

### 8.2 Methods

#### 8.2.1 Data source

The annotated genomes of the tuatara (this study) and several TSD and GSD reptiles, chicken and human (both GSD) were obtained from public databases (Table 8.1). Genomic regions were identified from the available annotation files. Two databases were used: the Ensembl database (data source for the *Homo sapiens* and *Pelodiscus sinensis* genomes) and the University of California-Santa Cruz database –UCSC- (data source for the *Gallus gallus*, *Alligator mississippiensis*, *Anolis carolinensis*, and *Chrysemys picta bellii* genomes). For the UCSC genomes, gtf files were obtained in the table browser of the UCSC database, downloading format for genes and gene predictions, using the genescan annotation.

#### 8.2.2 Data processing

We followed the protocols described in Radhakrishnan et al 2017[142] with some modifications. Briefly, CDS sequences were extracted to generate fasta files for gene, exons, and intron regions using Bedtools[143]. Promoter and intergenic sequences were then extracted from the upstream regions of these genes at various scales (100 bp, 300 bp, 500 bp, 1000 bp and 3000 bp) using the size chromosome data and gene direction information as a reference. Sequences with undefined nucleotides were eliminated prior to further analysis, which reduced the final dataset for some genomic regions in some taxa more substantially, particularly in reptiles other than tuatara (Figure 8.2).

The nCpG content was calculated for each genic and intergenic region as

$$nCpG = \frac{(\frac{cg}{l})}{(\frac{c}{l}) \times (\frac{g}{l})}$$

where l = sequence length, c = number of cytosines, g = number of guanines, and cg = number of cytosines bordered by a guanine linked by a phosphate group (CpG)[136].

nCpG distributions for each genomic region were plotted using R software version 3.3.1[144]. The likelihood of a mixture model with one (G=1) and two (G=2) components was assessed using the R package Mclust[145] and the better fit model was chosen by a likelihood ratio test.

### 8.3 Results and discussion

We present the first examination of nCpG distribution in tuatara, the only representative of the reptilian order Rhynchocephalia, and shed new light on the commonalities and diversity of nCpG distribution across vertebrates, a trait that is linked to DNA methylation and may consequently impact gene expression. First, our results revealed that the tuatara exhibited bimodal nCpG distribution in all genomic regions examined (gene promoters, exons, introns, and intergenic sequences), similar to recent reports in painted turtles, the only other TSD reptile previously studied[142], a pattern also shared by alligator (TSD) (Figure 8.1).

Bimodal nCpG distributions at promoters characterize all vertebrates examined previously and in this study (human, non-human primates, opossum, platypus, chicken, alligator, *Chrysemys* turtle, *Pelodiscus* turtle, *Anolis* lizard, *Xenopus* frog, *Danio* fish) with the exception of platypus and *Ciona* tunicates (a representative of a chordate lineage basal to vertebrates)[135,141,142], suggesting that this bimodal pattern in promoters may have arisen with the emergence of vertebrates but was lost in monotremes. It should be noted that the previous assessment of unimodality in platypus promoters was qualitative[135] and it is likely that explicit statistical tests such as mixed models [[142] and this study] would have uncovered a better fit for a bimodal model, albeit much more subtle than the bimodality previously reported in birds and other mammals.

Second, the bimodal nCpG pattern seen in introns of TSD reptiles (tuatara, painted turtle and alligator) would appear at first glance to mark a divide between TSD taxa and the GSD vertebrates previously reported to display a unimodal nCpG distribution in introns[135,141,142]. Furthermore, a bimodal intronic nCpG distribution was also reported in tunicates, which would suggest that such bimodal pattern may be ancestral to vertebrates and may have been lost in GSD vertebrate lineages, at least in those examined to date. However, here we revisited the genomes of several of these previously studied GSD vertebrates (human, chicken, anole) and investigated a GSD turtle for the first time (*Pelodiscus*). We tested explicitly for the existence of bimodality in all genomic regions examined using mixed models. Our results challenge the notion of intronic unimodality in nCpG distribution among vertebrates in general, supporting instead a bimodal pattern that appears to be ancestral and conserved across all vertebrates irrespective of their sex-determining mechanism. This finding agrees with the recent detection of bimodal DNA methylation in gene bodies in human, chicken, Danio fish, and Ciona tunicates[146].

#### 8.4 Figures

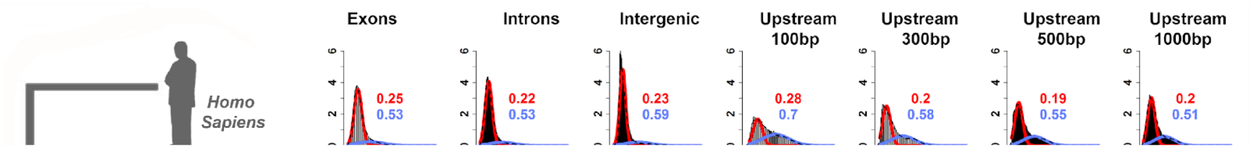

**Figure 8.1: Normalized CpG distributions (nCpG) for tuatara and other vertebrates.**

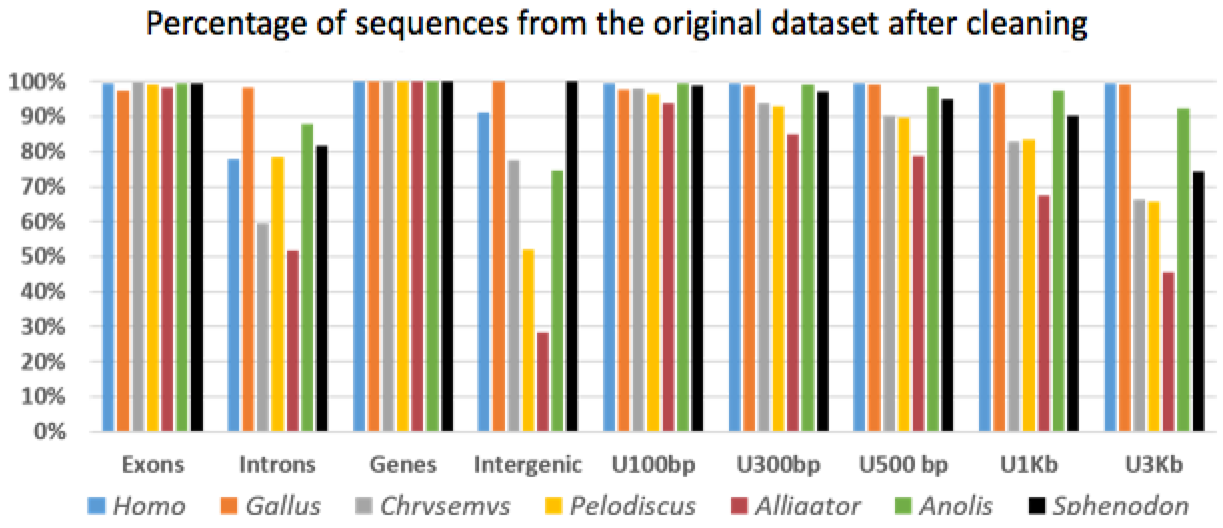

**Figure 8.2: Effect of removal of sequences with undefined nucleotides (Ns) from the database used in this study.**

#### 8.5 Tables

**Table 8.1: Vertebrate genomes examined in this study.** SDM = Sex-determining mechanism. TSD = Temperature-dependent sex determination. GSD = Genotypic sex determination.

| **Group** | **Species** | **SDM** | **Source** | **Genome_ID** |
|---|---|---|---|---|
| Tuatara | *Sphenodon punctatus* | TSD | UCSC | This study |
| Lizard | *Anolis carolinensis* | GSD | UCSC | AnoCar2.0 (GCA_000090745.1) |
| Turtle | *Chrysemys picta bellii* | TSD | UCSC | ChrPic1 (GCA_000241765.1) |
| Turtle | *Pelodiscus sinensis* | GSD | Ensembl | PelSin_1.0 (GCA_000230535.1) |
| Crocodilian | *Alligator mississippiensis* | TSD | UCSC | AllMis (GCA_000281125.1) |
| Bird | *Gallus gallus* | GSD | UCSC | Galgal5 (GCF_000002315.4) |
| Mammal | *Homo sapiens* | GSD | Ensembl | GRCh38.p7 (GCA_000001405.22) |

### 9 EVOLUTION OF GENOMIC ORGANIZATION OF THE MHC

Yuanyuan Cheng*, Hilary Miller

Genes of the major histocompatibility complex (MHC) play an important role in disease resistance and kin recognition and are among the most polymorphic in the vertebrate genome. Prior work[147] located the tuatara core MHC region to chromosome 13q, although a number of class I and class II genes were mapped to several other autosomes. Here, annotation of tuatara MHC regions and comparisons of gene organization to seven other representative species (Figure 9.1) identified 56 genes spanning 13 scaffolds, including six class I genes, six class II genes, 19 class III genes, 18 framework genes, and seven extended class II genes. Four class I and four class II genes which are not linked to other MHC related genes were excluded from the comparative analysis, as they are likely located outside the core MHC region[147]. As shown in Fig. 9.1, the genomic organization of tuatara MHC is most similar to that of the anole, which we interpret as being typical of the Lepidosauria. Unlike the chicken MHC, which was proposed to represent the minimal essential MHC gene cluster[148], MHC of the tuatara, anole, alligator, and turtle all show a high gene content and complexity that is more similar to the MHC regions of amphibians and mammals. While the majority of genes annotated in the tuatara MHC are well conserved as one-to-one orthologs, extensive genomic rearrangements were observed among these distant lineages (Figure 9.1). The most noteworthy rearrangement lies in the relative locations of genes in the class I and class III regions (as defined by their locations in the human MHC), with these two regions inverted between the mammalian and frog MHC, and the genes interspersed in the tuatara and anole genomes. This finding raises questions over the traditional categorization of these genes as class I or class III region genes, as evolutionarily there is no explicable difference between the two sets of genes in terms of gene clustering, organization, or function.

Prior work identified strong population differentiation at tuatara MHC class I genes, with population bottlenecks and isolation considered a larger influence in shaping this diversity than selection[149]. Further work using the expanded MHC gene set identified here might identify further population-level features important for the conservation of this species.

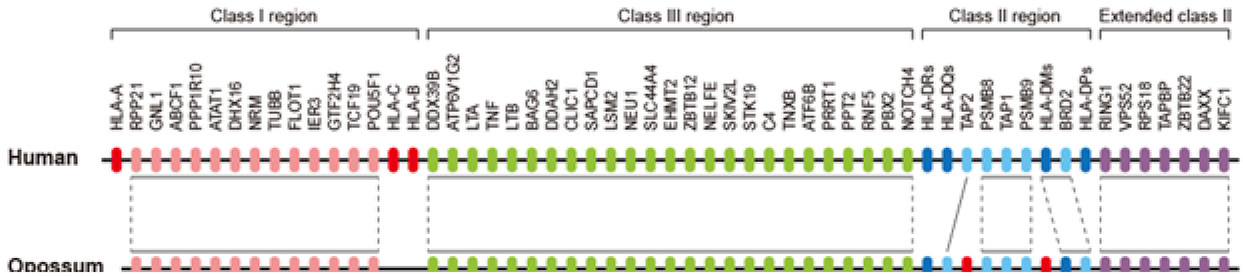

**Figure 9.1 Comparative analysis of the MHC core region.** Species examined include human (assembly version GRCh38.p11), gray short-tailed opossum *Monodelphis domestica* (MonDom5), western painted turtle *Chrysemys picta bellii* (Chrysemys_picta_bellii-3.0.3), Chinese alligator *Alligator sinensis* (ASM45574v1), green anole *Anolis carolinensis* (AnoCar2.0), chicken *Gallus gallus* (Gallus_gallus-5.0), and western clawed frog *Xenopus tropicalis* (Xenopus_tropicalis_v9.1). Only genes that were annotated in the tuatara genome were included in the analysis. Orthologs between species are connected by a solid line; the grey bars above/below genes indicate syntenic blocks and are linked by dashed lines between species. Anolis class I/II and extended class II regions are not shown due to the high degree of genome assembly fragmentation in these regions. Colour legend: red – class I genes, pink – class I region framework genes, green – class III genes, dark blue – class II genes, light blue – class II region framework genes, purple – extended class II genes.

### 10 MOLECULAR EVOLUTION OF TUATARA VISUAL SYSTEM

Ryan K Schott*

#### 10.1 Introduction

The tuatara is primarily a nocturnal predator that is able to capture prey under extremely dim light conditions[150]. Vision appears to be the primary sensory system for pretty capture: auditory and olfactory cues alone were insufficient for pretty capture, but under even very dim light (down to 0.00615 lux), in the absence of other cues, tuatara were able to capture prey[150,151]. Despite this extreme nocturnal visual adaptation, the tuatara eye has an interesting mix of diurnal and nocturnal features including a fovea typical of diurnal species and a slit pupil found in many nocturnal species, as well as a distinct complement of photoreceptors[152].

Previous morphological analyses of the tuatara retina revealed four types of photoreceptors: large single rods, small single rods, double rods, and small single cones[152,153]. The small single cone did not have an oil droplet and was restricted to the periphery of the retina. Unusually, the rods were found to have colourless oil droplets, which are normally found in cones. Most vertebrates also only have a single type of rod, however, some geckos and snakes have multiple rod types, which are evolutionary derived from cones through the process of photoreceptor transmutation[152,154,155]. Based on detailed ultrastructural analysis using electron microscopy, Meyer-Rochow and Ahnelt[156] found that the 'rods' of tuatara were also derived from cones. Unfortunately, the ultrastructure of the small single cone reported by Walls[152] was not analyzed. Furthermore, there have been no molecular studies of the tuatara visual system.

The tuatara genome provides a unique opportunity to uncover the molecular basis of these interesting morphological adaptations. Utilizing this resource, we analyzed the loss of visual genes in tuatara, evaluated visual pigment and photoreceptor complement, and tested for shifts in selective pressure in visual genes that may reflect the distinct visual adaptations in this enigmatic species.

#### 10.2 Methods

To explore the molecular evolution of the tuatara visual system a set of 119 visual genes were targeted for analysis[157,158]. These were extracted from the *Sphenodon punctatus* genome assembly and CDS annotation using BLAST searches (blastn, discontiguous megablast, tblastn) using *Anolis* and *Chrysemys* sequence queries. Gene identifications were confirmed with reciprocal BLAST and phylogenetic analysis, as necessary. Additionally, sequences were manually curated to ensure open reading frames.

Visual gene loss in tuatara was compared to that in other amniotes following the analysis of Perry et al.[158], which was updated to account for recent genome publications (most notably an updated platypus genome assembly[159]). Genes were initially searched for in the NCBI nucleotide database. If a sequence could not be found, BLAST was used as described above to search genome assemblies directly. Gene loss was inferred if a sequence could not be found in any of the available genomes for a particular lineage. Partial sequences were counted as present if they had an open reading frame.

Phototransduction genes were targeted for additional analyses to determine how selection has shaped the molecular evolution of the tuatara visual system. The phototransduction gene datasets of Schott et al.[160,161] were used with the addition of the tuatara sequences. Additionally, new datasets were built for SWS2 and SLC24A1 since these genes were not analyzed previously because they were lost in squamates (but not tuatara) following the methodology outlined in Schott et al.[160,161] .

To estimate the strength and form of selection acting on these genes we utilized the codon-based likelihood model implemented in PAML[162]. Specifically, we used the random sites models to estimate alignment-wide selection and Clade Model C (CmC [163]) to compare selective pressure in tuatara with other reptiles. Since geckos and snakes have previously been shown to be under different selective pressures, we evaluated whether tuatara had similar selective pressures as these groups by including tuatara in partitions with geckos, snakes, and both geckos and snakes, as outlined in Figure 10.1. All analyses were run with varying starting values to avoid potential local optima. To determine significance, model pairs were compared using a likelihood ratio test (LRT) with a $\chi 2$ distribution, while non-nested models where evaluated using Akaike Information Criterion (AIC).

#### 10.3 Results and discussion

##### 10.3.1 Visual gene loss in tuatara and amniotes

Loss of visual genes in nocturnal animals is expected due to relaxed selective pressures, most often on genes involved in bright light vision. Indeed, previous analyses showed high levels of gene loss in groups that are hypothesized to have undergone strong nocturnal bottlenecks (mammals, snakes, geckos), while primarily nocturnal predators (such as crocodilians) showed moderate gene loss[158]. Surprisingly, we found that visual gene loss in tuatara was remarkably low compared to other major amniote lineages (Fig. 10.2). Tuatara specifically lost only a single gene, TMT3, a nonvisual opsin, that was lost independently in mammals, snakes, and archosaurs (birds and crocodilians). Additionally, two genes (GNGT1 and PDE6A) were also absent in tuatara due to their loss in the reptilian ancestor. The total loss of only three visual genes is the lowest among amniote lineages tied with diurnal squamates (e.g., anguids and iguanids).

##### 10.3.2 Visual pigments and photoreceptors of tuatara

Morphological analyses of tuatara photoreceptors suggest that the three types of 'rods' (large, single, double) are actually evolutionarily derived from cones through photoreceptor transmutation[152,153,156]. However, this could not previously be confirmed from a molecular perspective. We found that tuatara possess the full complement of visual opsins (and therefore visual pigments) that were present in the vertebrate ancestor (RH1, RH2, LWS, SWS1, SWS2), as well as both the rod and cone phototransduction genes present in other reptiles. These genes are evolutionary conserved (see below) suggesting functioning rod and cone phototransduction pathways. This provides strong molecular support for photoreceptor transmutation in tuatara.

The multiple types of rod-like cone photoreceptors, which contain oil droplets, likely express cone visual pigments. If tuatara follows the typical vertebrate pattern this would place the long-wavelength sensitive (LWS) visual pigment in the large single 'rods' and double 'rods' and the other cone visual pigments (RH2, SWS2, SWS1) in subtypes of the small single 'rod' (Figure 10.3). The small single (dwarf) cone is interesting as it lacks an oil droplet and is restricted to the periphery of the retina, features typical of rods. Most diurnal lizards lack morphological rods, but still express rod visual pigments and other rod phototransduction proteins[160], presumably in transmuted cone photoreceptors. If cone-like rods are the ancestral condition for squamates, this raises the possibility that the dwarf cone in tuatara is also a rod-like cone, homologous with those in squamates, and therefore would be the rod opsin (RH1)-bearing photoreceptor (Figure 10.3). It is also possible that RH1 is instead expressed in one of the droplet-bearing rod-like cones. If this is the case then the dwarf cone likely instead contains the UV-sensitive SWS1 pigment, which is often the rarest photoreceptor type in other vertebrates. RH1 expression in a droplet-bearing cell would be unusual, but may be the case for the diurnal iguanid *Polychrus marmoratus*[164]. Unfortunately, the opsins underlying the photoreceptor types in squamates have not been confirmed, which makes inferring such patterns in tuatara difficult. Despite this, these results point to interesting patterns of evolution in the photoreceptors of tuatara and other lepidosaurs that warrant further study.

The five main classes of visual pigment (encoded by the visual opsins) each absorb light at a characteristic wavelength. Differences in the absorption between classes are due to differences in protein sequence at spectral tuning sites[165]. We analyzed the opsin sequences of tuatara at known spectral tuning sites to roughly estimate the wavelengths of maximal absorbance ($\lambda_{max}$) of the visual pigments. Overall, we found little variation at known sites, although it should be noted that spectral tuning is not well understood in non-avian reptiles and there are likely to be other sites that affect spectral tuning in tuatara that have not been accounted for here. In rod opsin (RH1) we found N83 in tuatara. This substitution causes a small blue-shift in many vertebrate RH1s and has also been implicated as a dim-light adaptation in several lineages that inhabit light-limited environments[166–168]. Thus, tuatara RH1 may be further dim-light adapted and likely has a slightly blue-shifted $\lambda_{max}$ around 490–495 nm. In the LWS opsin, we found that tuatara has A269, which results in a ~15 nm blue-shift in mammals[165], likely resulting in a tuatara LWS $\lambda_{max}$ around 535–540 nm. The tuatara RH2, SWS2, and SWS1 pigments do not show any differences at known spectral tuning sites and these pigments thus likely had $\lambda_{max}$s around 500, 440, and 360 nm, respectively.

###### 10.3.2.1 Selective pressures of tuatara phototransduction genes

Tuatara has a highly sensitive visual system and is able to capture prey under extremely low light conditions (down to 0.00615 lux[150]). This is achieved using rod-like photoreceptors that are evolutionarily derived, through photoreceptor transmutation, from cones. This process is expected to impose distinct selective pressures on the visual system. Previous work has shown that the phototransduction genes of caenophidian snakes and geckos underwent a long-term shift in selective pressure associated with this process of photoreceptor transmutation[160]. We tested for a similar shift in tuatara phototransduction genes.

Overall, tuatara phototransduction genes do not show a significant long-term shift in selective pressures. In only five of the 33 genes was tuatara included in the best-fitting partition (Table 10.1; Supplementary Data 10.5281/zenodo.2597599). This included two rod genes (*CNGB1*, *PDE6B*) and three cone genes (*CNGA3*, *GNGT2*, and *PDE6H*). Interestingly there does seem to be a focus in both the rod and cone genes on the cyclic nucleotide gated channels (CNGs) and phosphodiesterase (PDE), although it is not clear if this could point to a functional adaptation towards dim-light vision. The low number of phototransduction genes under distinct selective pressures in tuatara is in contrast to caenophidians being in the best fitting partition for 25 of the 29 genes and geckos for 20 of the 25 genes present in these lineages. These results suggest that photoreceptor transmutation did not impose similar selective pressures in tuatara as in snakes and geckos.

###### 10.3.2.2 Evolution of the tuatara visual system

The results presented here provide a novel molecular perspective on the evolution of the tuatara visual system that corroborates the morphological data that tuatara possess a unique mixture of diurnal and nocturnal adaptations. Despite substantial dim-light adaptation and rod-like cone photoreceptors, the tuatara visual system is highly conserved with the tuatara lineage having one of the lowest rates of gene loss of any amniote and highly conserved phototransduction genes.

The maintenance of five visual opsins suggests that tuatara has a robust colour vision system that, due to the presence of rod-like cones, may function in dim-light conditions, similar to nocturnal geckos, which also have rod-like cones[169]. Geckos only have 3 visual pigments, however, implying that tuatara nocturnal colour vision could have superior colour discrimination. Both geckos and snakes showed concerted, long-term shifts in selective pressure on phototransduction genes associated with photoreceptor transmutation. Although tuatara also has strong evidence for photoreceptor transmutation we did not recover a similar selective pattern in in this lineage. This suggests a distinct path to nocturnal adaptation in tuatara that is perhaps constrained by its unusual life history: juvenile tuatara are often diurnal and arboreal to avoid predation by cannibalistic adults, while adults hunt primarily at night, but also bask during the day[170]. This dichotomous selective pressure likely contributed strongly to the unique visual system of the tuatara.

#### 10.4 Figures

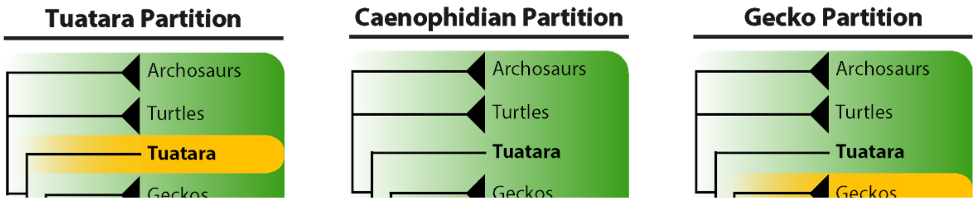

**Figure 10.1. Clade partitioning schemes used to test for long-term shifts in selective pressures in tuatara phototransduction genes.** The clade/lineage highlighted in yellow was placed into a separate partition from the rest of the tree allowing independent estimates of $d_N/d_S$ in the divergent site class for the two partitions.

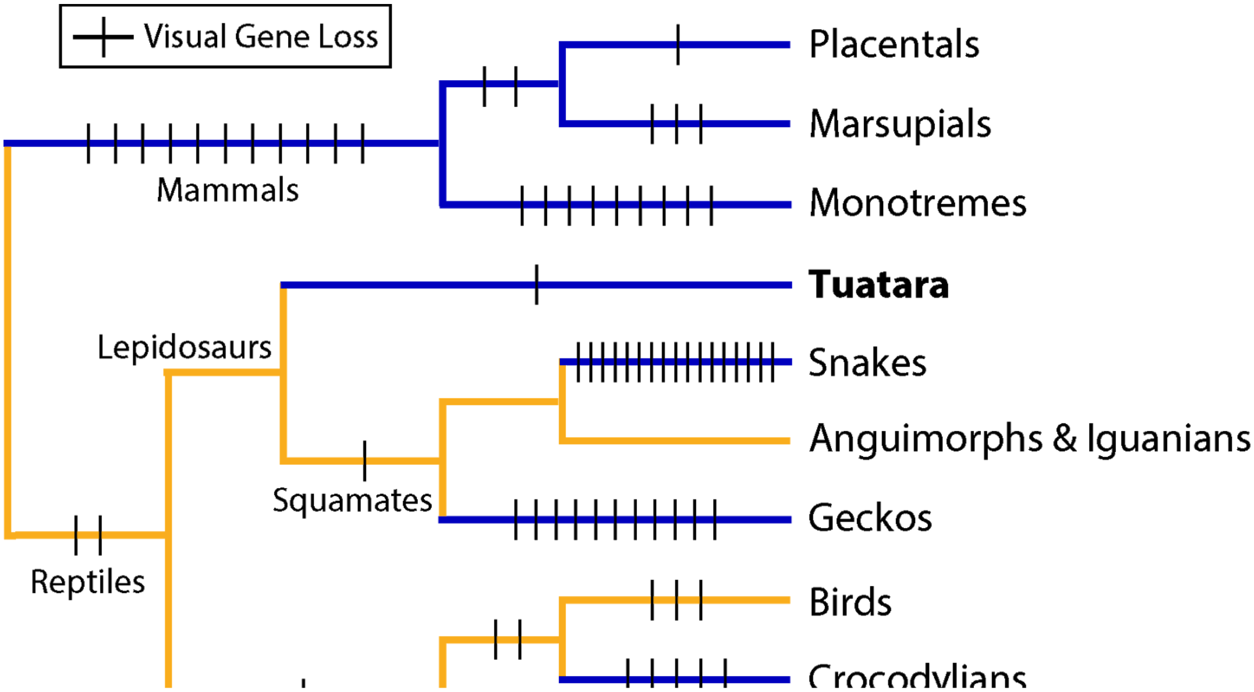

**Figure 10.2. Phylogenetic tree of reptiles depicting inferred visual gene losses.** Lineages are coloured based on a rough approximation of their ancestral activity pattern (blue, nocturnal; yellow, diurnal). Note that that tuatara lineage has experienced some of the lowest rates of gene loss despite a nocturnal ancestry, which in other lineages is associated with increased gene loss.

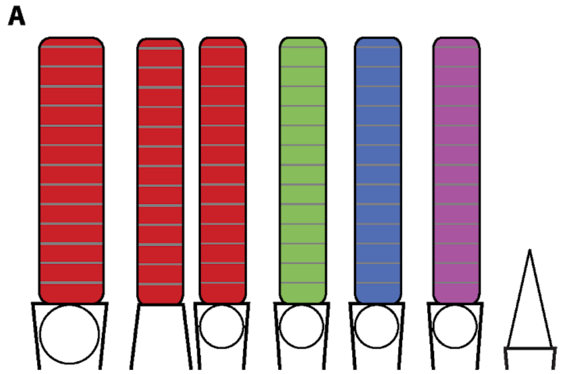

**Figure 10.3. Schematic of photoreceptors types of tuatara showing two potential distributions of visual pigments.** Tuatara has three morphological types of rod-like cones and one dwarf cone [153], and has four cone opsins (LWS, RH2, SWS2, and SWS1) and one rod opsin (RH1). The dwarf cone lacks an oil droplet and is absent from the fovea and central retina making it a strong candidate for the rod opsin (RH1)-bearing photoreceptor, perhaps homologous with the rod opsin-bearing cones of diurnal lizards (A). Alternatively RH1 may be present in a subtype of the rod-like cone, making the dwarf cone most likely to contain a UV-sensitive SWS1, which is often the rarest photoreceptor type.

#### 10.5 Tables

**Table 10.1. Summary of analyses comparing selective pressures in tuatara with those in reptiles, geckos and caenophidian snakes.** The best fitting partition (Figure 10.1) is shown for each gene (when significant vs the null model, M2a_rel). Full results tables are presented in the Supplementary Data (DOI: 10.5281/zenodo.2597599). Abbreviations: **C**, caenophidian partition; **G**, gecko partition; **T**, tuatara partition; **GC**, gecko plus caenophidian partitions; **TC**, tuatara plus caenophidian partition; **TG**, tuatara plus gecko partition; **TGC**, tuatara plus gecko plus caenophidian partition.

| Type | Gene | Best-fitting Partition |
|---|---|---|
| Rod | *CNGA1* | - |
| Rod | *CNGB1* | **TGC** |
| Rod | *GNAT1* | GC |
| Rod | *GNB1* | - |
| Rod | *GRK1* | G |
| Rod | *PDE6B* | **TC** |
| Rod | *PDE6G* | C |
| Rod | *RH1* | C |
| Rod | *SAG* | GC |
| Rod | *SLC24A1* | - |
| Cone | *ARR3* | GC |
| Cone | *CNGA3* | **TGC** |
| Cone | *CNGB3* | GC |
| Cone | *GNAT2* | GC |
| Cone | *GNB3* | GC |
| Cone | *GNGT2* | **TGC** |

| | | |
|---|---|---|
| Cone | *GRK7* | GC |
| Cone | *GUCA1C* | G |
| Cone | *LWS* | C |
| Cone | *PDE6C* | GC |
| Cone | *PDE6H* | **TG** |
| Cone | *RH2* | G |
| Cone | *SLC24A2* | GC |
| Cone | *SWS1* | C |
| Cone | *SWS2* | - |
| Both | *GNB5* | GC |
| Both | *GUCA1A* | C |
| Both | *GUCA1B* | C |
| Both | *GUCY2D* | GC |
| Both | *GUCY2F* | C |
| Both | *RCVRN* | C |
| Both | *RGS9* | GC |
| Both | *RGS9BP* | GC |

### 11 ODORANT RECEPTORS

Melissa Jordan and Richard Newcomb

#### 11.1 Methods

Odorant receptor sequences from the green anole (*Anolis carolinensis*) and Burmese python were used as query sequences for tBLASTn[34] searches of the tuatara genome using Geneious 10.0.3 (https://www.geneious.com). Iterative searches were performed using newly identified tuatara ORs to ensure a comprehensive set of genes was attained. Genes were classified in the following way; intact genes consisted of an uninterrupted open reading frame >300 amino acids with an appropriate start and stop codon, ORs were considered pseudogenes if they contained interrupting stop codons or frameshifts. Translated amino acid sequences for phylogenetic analysis were obtained from manually annotated published data for *Gallus gallus* and *Taeniopygia guttata*[171] or from automatic annotation via NCBI for *Anolis carolinesis*, *Gekko japonicas*, *Notechis scutatus*, *Ophiophagus hannah*, *Pogona vitticeps*, *Protobothrops mucrosquamatus*, *Pseudonaja textilis*, *Python bivittatus* and *Thamnophis sirtalis*. Only sequences greater than 280 amino acids in length were used in the analysis. Multiple sequence alignment was performed using FFT-NS-1 parameters in MAFFT (v7.338)[115]. The alignment was trimmed such that any columns containing more than 75% gaps were removed, which resulted in an alignment consisting of 322 characters. Phylogenetic relationships were estimated using the Neighbor-Joining method with Poisson substitution model and 1000 bootstrap replicates within MEGA 7.0.21[172] and the tree drawn and edited using Figtree v1.4.4 (https://tree.bio.ed.ac.uk/software/figtree).

#### 11.2 Results

Vertebrate ORs are G protein-coupled receptors that contain seven transmembrane domains[173]. They are expressed in the membrane of olfactory neurons within the olfactory epithelium where odorants are detected by extracellular regions of the receptor. Binding of an odorant to its corresponding receptor initiates a signal transduction pathway which ultimately leads to transmission of the action potential to the olfactory bulb[174]. Four hundred and seventy two genes encoding predicted odorant receptors (ORs) were identified from the genome of tuatara (10.5281/zenodo.2592798). Of these, 62 genes were identified as pseudogenes, containing one to multiple stop codons or frameshifts within their sequence. The percentage of intact genes was therefore 72%, which is perhaps a slight underestimate as some of the 60 partial sequences are likely to represent intact genes.

Vertebrate ORs have a complex classification but can generally be split into two types, Type I and Type II. Type I can be subdivided into further groupings (alpha-zeta)[175,176] and also encompasses the mammalian OR classification groupings of Class II (gamma) and Class I (alpha and beta)[177]. Class I ORs are thought to be more specialised towards detecting water soluble odorants and correspondingly all fish ORs fall into Class I. Class II ORs recognise airborne odorants.

Phylogenetic analyses were undertaken to compare ORs from tuatara with those from other terrestrial Sauropsids, including snakes (Burmese python, brown snake, garter snake, king cobra, pit viper and tiger snake), lizards (bearded dragon, green anole and Japanese gecko) and birds (chicken and zebra finch). Tuatara ORs are predominantly members of the Class II gamma group, with a small number of Class I ORs corresponding to the alpha group (Figure 11.1). Within the gamma group tuatara have an extensively radiated clade containing 65 ORs that is sister to a much larger equivalent set of radiated ORs in the birds (gamma-c group[171]). The presence of tuatara ORs across the tree indicates that the origin of the major OR lineages predates the split of birds, lizards and tuatara and likely evolved rapidly via duplication and divergence from a small number of Class II ORs in fish (one in zebrafish) and then amphibians[175,176]. The tuatara genome demonstrates that this rapid expansion of ORs that recognise airborne odorants occurred across all the major lineages within the Sauropsida.

**Figure 11.1 The evolutionary history of terrestrial Sauropsid odorant receptors inferred using the neighbor-joining method.** The unrooted tree contains 3213 amino acid sequences. Branches are coloured according to the following categories: Green – tuatara, Blue – birds (*Gallus gallus*, *Taeniopygia guttata*), Red – snakes (*Notechis scutatus*, *Ophiophagus Hannah, Protobothrops mucrosquamatus*, *Pseudonaja textilis*, *Python bivittatus* and *Thamnophis sirtalis*), Orange – lizards (*Anolis carolinesis, Pogona vitticeps*) and Purple – gecko (*Gekko japonicas*). Bootstrap support values above 75% (1000 replicates) are indicated for major branch splits relating to the different OR groups and branches leading to the species-specific OR expansions in birds (group γ-c) and tuatara (*).

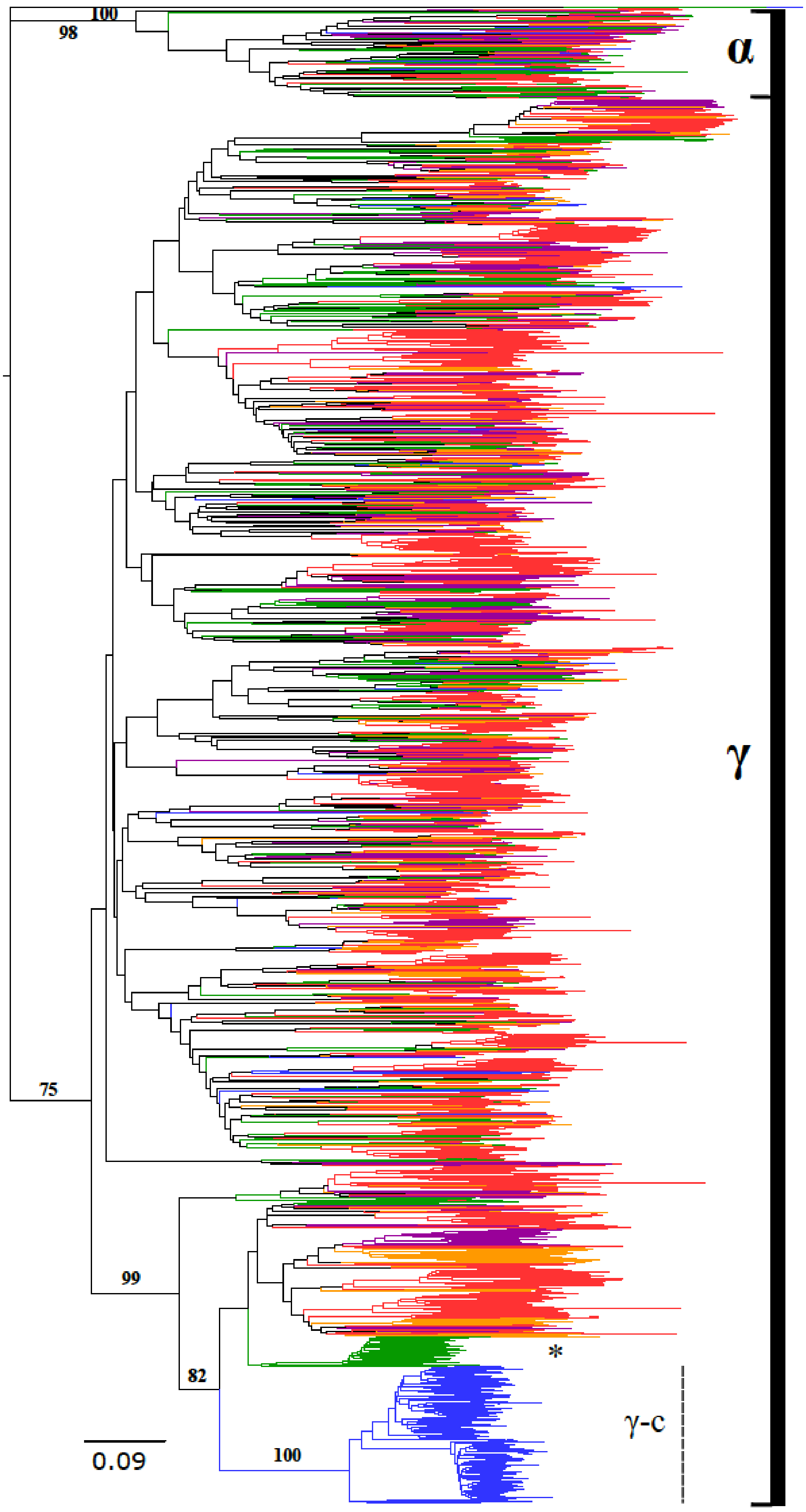
100
98
α
γ
75
99
82
100
*
γ-c
0.09

### 12 TRANSIENT RECEPTOR POTENTIAL (TRP) ION CHANNELS GENES IN SPHENODON PUNCTATUS

José Ignacio Arroyo

#### 12.1 Background

Tuatara is a behavioural thermoregulator. The Transient Receptor Potential (TRP) ion channels genes play an important role in thermoregulation as some of these genes participate in thermosensation and cardiovascular physiology. To search for the genomic adaptations related to thermoregulation in tuatara we performed a comparative genomic analysis of TRP genes in tuatara and six other species of amniotes. Duplicated thermo- and non-thermosensitive TRP genes were identified in tuatara that are differentially retained in this species, including members of most of the TRP subfamilies. A signature of positive selection in heat-sensitive TRP genes was observed when comparing orthologous genes as well as when comparing recently duplicated paralogous genes within Sphenodon.

Gene duplication is a fundamental mechanism of genomic innovation. After duplication paralogous genes can diverge under the action of positive selection and be differentially retained among descending lineages[178]. Both these processes, gene divergence and differential retention, can be potentially related to changes in the traits of species including their reproduction and survival. One of the fundamental traits of species is their ability to maintain their body temperature within certain ranges, a task that they can perform physiologically (endotherms) or behaviourally as is the case of ectotherms[179].

Tuatara (*Sphenodon punctatus*) are behavioural thermoregulators[180–182], meaning that in order to regulate their body temperature they move through their environment to find optimal temperature habitats. Temperature is a crucial environmental variable that affects physiological processes, life history traits and fitness[183]. In behavioural thermoregulation, thermosensation of the environment is a critical step. From a molecular standpoint, thermosensation is controlled by Transient Receptor Potential (TRP) ion channels[184]. TRPs are cold/heat-gated ion channels that have a long length, all of them including at least six putative transmembrane domains. For instance, human TRP genes are diverse in length and range between 11.4 and about 911 kb, with the number of exons varying from 11 to 39[185]. The TRP family includes seven well-recognized subfamilies (TRPA1, TRPV, TRPML, TRPM, TRPC, TRPP-also called PKD-, TRPN[186,187]. Most of these genes are dispersed in vertebrate genomes despite some of them being tandemly arranged, likely due to being derived from small-scale duplication events. For instance, in the human genome

TRPM3 and -6 are in chromosome 9, TRPV1-3 are in tandem in chromosome 17, TRPV5-6 are in chromosome 7 and TRPML2-3 are in chromosome 1, whereas all the remaining members are dispersed in different chromosomes in the genome[185]. The repertoire of these genes varies among species. This variation has been relatively well studied in the model species in Bilateria. For instance, in Drosophila there are 17 TRP genes and in human 27[185]. Within chordates there is lower variation but it remains not well studied and mostly limited to model species. Among TRPs, TRPA1, TRPM and TRPV are thermosensitive (thermoTRPs)[188–190], which detect from cold (i.e. TRPV1) to hot (i.e. TRPM8) stimuli[185]. TRP genes participate in thermoregulation, which is critical for reproduction and survival in ectotherms such as reptiles, thus it seems probable that these genes could have been a target of natural selection whether on specific sites or on the divergence of duplicated genes. For instance, it has been shown that the inhibition of cold- and heat-sensitive TRP channels in the saltwater crocodile produces alterations in thermoregulation[191].

#### 12.2 Methods

Here, we performed a comparative genomic analysis of TRPs in tuatara and six representative species of amniotes in order to characterize the evolutionary history of this gene family in this species, including the variation in the repertoire of genes due to duplication and subsequent differential retention and rates of molecular evolution on orthologous and paralogous (divergence). To accomplish this goal, protein sequences for the seven TRP subfamilies were obtained from the Ensembl database for four representative amniote species covering the main lineages: human (*Homo sapiens*), chicken (*Gallus gallus*), lizard (*Anolis carolinensis*) and turtle (*Pelodiscus sinensis*). To annotate TRP genes from draft genomes of viper (*Vipera berus*), alligator (*Alligator mississippiensis*) and tuatara (*Sphenodon punctatus*) we used as a reference human TRP protein sequences that were aligned against draft genomes using tblastn[34]. We only included sequence matches with a cover and identity >30%. New identified genes were named in agreement to their phylogenetic relationships. Alignment was performed using MAFFT (strategy L-INS-i[115]). Phylogenetic relationships were estimated using the approximately maximum likelihood approach of FastTree2[63] as implemented in the CIPRES portal[80]. The tree was rooted with VDAC which are also voltage-gated ion channels (VGIC), which yields a topology consistent with previous studies[186]. All data are detailed in TreeBASE submission ID 23409.

#### 12.3 Results and discussion

We found that the repertoire of TRP genes varied among the species surveyed between 28–37 genes (Table 12.1). Tuatara has the largest repertoire of TRP genes among the species surveyed.

We identified 37 TRP-like sequences in the tuatara genome draft, but because TRPs are long genes, most identified genes corresponded to partial sequences (coverage of recovered alignments with human orthologous varied from 40–100%). Nonetheless for five genes (TRPV1-3, TRPM1-2) we recovered complete or (five cases of) almost complete genes (>90% of coverage to human orthologous). Most genes were recovered in separate scaffolds, except for some cases where it was possible to recover tandemly arranged TRP sequences (Table 12.1). These genes included partial sequences of TRPN-like genes (ScrUdWx_1447), TRPML2-3 (ScrUdWx_1600), TRPV1-3 (contig ScrUdWx_1875), TRPN and TRPC (ScrUdWx_55) and a pair of PKD1L3s (ScrUdWx_67_1, ScrUdWx_67_2) (Table 12.1).

The phylogenetic reconstruction recovered well-supported monophylies for each of the seven subfamilies (Figure 12.1; most of them >0.8) and well supported monophylies of each of the respective paralogous within each subfamily (Figure 12.1; supports not shown). The TRPP and TRPML clades were recovered in one clade and the -V, -C, -N, -A and -M in a separate clade (Figure 12.1). Relationships among subfamilies were consistent with previous studies[185].

A detailed examination of the topologies of the TRP subfamilies revealed repeated gene duplications followed by extensive losses and differential retentions of genes within amniotes (Figure 12.2). These duplicates include new (not previously reported) genes in tuatara (and other amniotes), including TRPV1/2/4, TRPC3/5/7, TRPN-2, TRPN-4, TRPN-5, PKD1-2, PKDREJ, PKDL13-2 and TRPML3 (asterisks in Figure 12.1, Figure 12.2). Some duplicates appear to be old as they were recovered as sisters of other subfamilies as is the case of TRPV1/2/4, TRPC3/5/7, while other duplications seem more recent originating in the common ancestor of reptiles or being specific to a single species. This analysis shows that the evolutionary history of these genes has been more dynamic than previously shown for some model species.

We explored variation in $\omega$ (dN/dS), the ratio of the rate of non-synonymous substitutions (dN) to the rate of synonymous substitutions (dS), in a maximum likelihood framework using the codeml program from PAML v4.4[162]. In brief, $\omega \approx 1$ indicates neutrality, $\omega < 1$ indicates negative or purifying selection and $\omega > 1$ indicates positive selection. Branch-site models were implemented to explore changes in $\omega$ across codons in a specific branch of the tree[192]. It only included sequences with a coverage >90%. In this case the branch of the tuatara was labelled as the foreground branch and non-tuatara branches were labelled as background. The Bayes Empirical Bayes (BEB) method was used to identify sites under positive selection[193,194]. The modified model A[195,196] with a $\omega$ free to vary was compared with a model with a $\omega$ fixed to 1, representing neutrality. The model estimates the proportion of four types of site classes ($p_0$, $p_1$, $p_{2a}$ and $p_{2b}$) and their respective omega values ($\omega_0$, $\omega_1$, $\omega_{2a}$ and $\omega_{2b}$). Site class 0 includes codons that are conserved throughout the tree, with $0<\omega0<1$ estimated. Site class 1 includes codons that are evolving neutrally throughout the tree with $\omega1=1$. Site classes 2a and 2b include codons that are conserved or neutral on the background branches, but become under positive selection on the foreground branches with $\omega_2>1$.

A branch-site analysis of molecular evolutionary rates among thermosensitive genes (testing only the five sequences with a coverage >90%; TRPV1-3, TRPM1-2) found significant positive selection in the heat-sensitive genes TRPM2 ($2\Delta lnL=3.18$, p-value=0.07, $p_2=0.014$, $\omega_2=4.66$) and TRPV2 ($2\Delta lnL=12.04$, p-value=0.0005, $p_2=0.071$, $\omega_2=9.29$).

In addition, to explore selection in genes where it was not possible to obtain complete coverage we used a pairwise comparative approach to estimate dN/dS rates as implemented in the program yn00[194]. In this approach we compared *Sphenodon* genes with TRP genes of *Anolis*, any other species that shared differentially retained genes with *Sphenodon* and also compared duplicated paralogous genes within *Sphenodon*. Among the 21 pairwise comparisons, many of them have an $\omega>1$ but in three cases there were moderately accelerated rates of dN>dS (~0.5), particularly among comparisons of partial TRPN sequences (Figure 12.3).

In general these results show a high rate of gene differential retention and positive selection in genes for which a function in heat sensation is well established[190,197]. Given the role of these genes, it seems probable that these genomic changes in TRP genes are associated with the evolution of thermoregulation in this species. Duplications in non-thermosensitive TRP genes that participate in cardiovascular physiology also appear to play a role in thermoregulation (see[184,186]). While the function of these genes have not been specifically studied in tuatara, function has been studied in other species. For instance PKD1 is an integral membrane protein involved in cell-cell/matrix interactions[198], PKD1L3 participates in taste reception[199], TRPN has roles in proprioreception and hearing[200,201].

#### 12.4 Figures

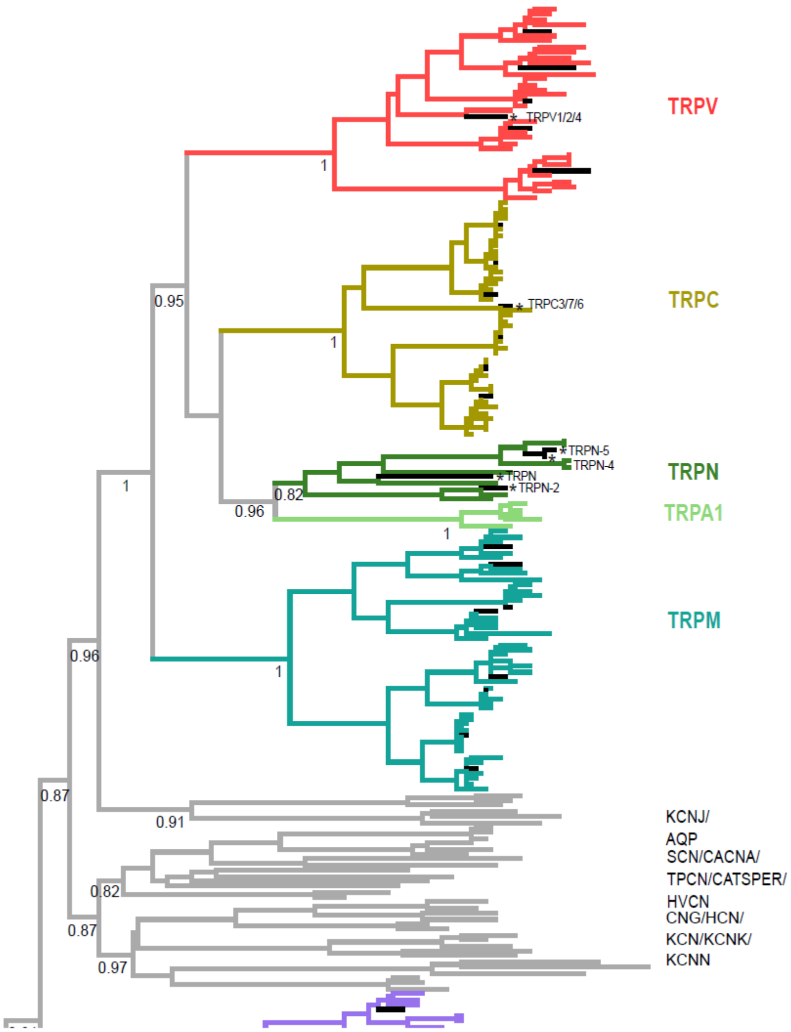
TRPV
TRPV1/2/4
TRPC
TRPC3/7/6
TRPN-5
TRPN-4
TRPN
TRPN-2
TRPN
TRPA1
TRPM
KCNJ/
AQP
SCN/CACNA/
TPCN/CATSPER/
HVCN
CNG/HCN/
KCN/KCNK/
KCNN
1
0.95
1
0.82
0.96
1
1
1
0.96
0.87
0.91
0.82
0.87
0.97

**Figure 12.1. Phylogenetic relationships among TRP ion channels genes in amniotes.** Blackened terminal branches denote tuatara branches. Node supports correspond to local bootstrap support values derived from 1,000 pseudoreplicates, as implemented in FastTree2 (only relevant nodes are showed). Asterisks indicate TRPs differentially retained in tuatara. The differential retention of genes is characterized in a phylogenetic tree because after their evolutionary origin they are lost in most species but are retained only in a few, as is the case of the new genes characterized in tuatara. These new genes (not previously reported) include: (from top to bottom) TRPV1/2/4, TRPC3/5/7, TRPN-2, TRPN-4, TRPN-5, PKD1-2 and PKDL3-2. These new genes were named according to their phylogenetic affiliations. For instance, the gene TRPV1/2/4 retained only in tuatara and turtle is sister to the genes TRPV1, TRPV2 and TRPV4. Other voltage-gated ion channels and voltage-gated ion channels (e.g. CATSPER, AQP) were also included in the phylogenetic reconstruction. The tree was rooted with the voltage-gated non-TRP human VDAC genes.

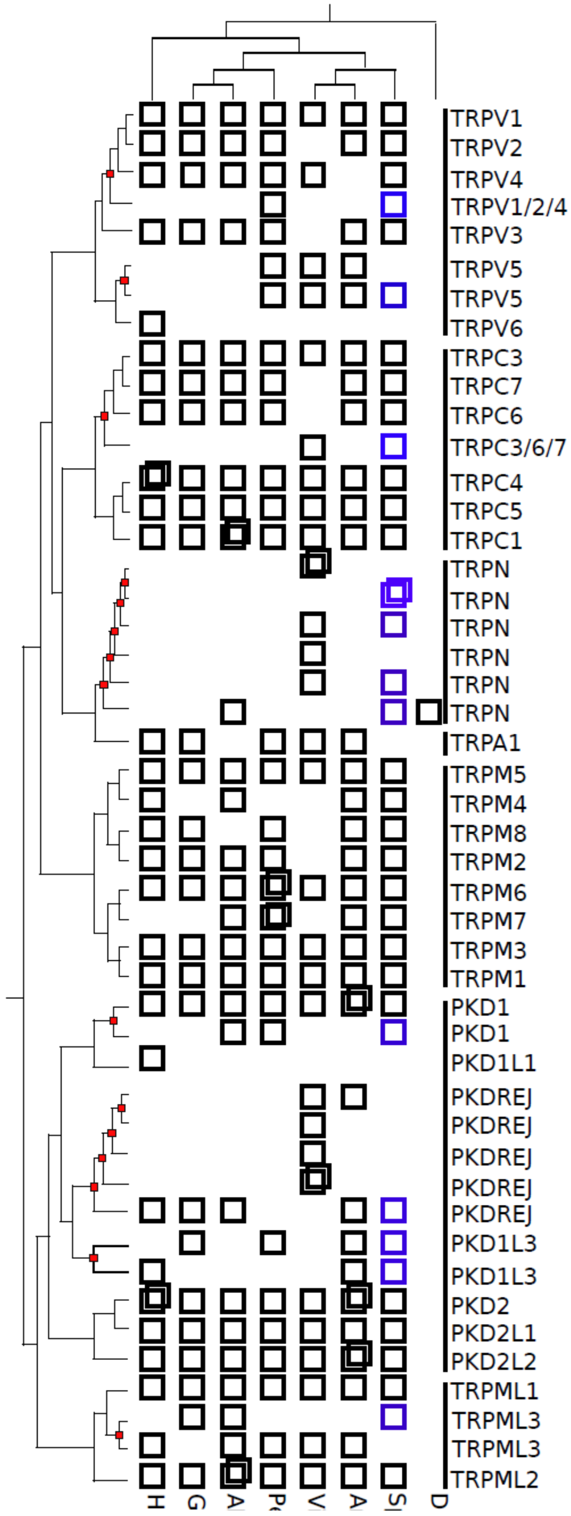
TRPV1
TRPV2
TRPV4
TRPV1/2/4
TRPV3
TRPV5
TRPV5
TRPV6
TRPC3
TRPC7
TRPC6
TRPC3/6/7
TRPC4
TRPC5
TRPC1
TRPN
TRPN
TRPN
TRPN
TRPN
TRPN
TRPA1
TRPM5
TRPM4
TRPM8
TRPM2
TRPM6
TRPM7
TRPM3
TRPM1
PKD1
PKD1
PKD1L1
PKDREJ
PKDREJ
PKDREJ
PKDREJ
PKDREJ
PKD1L3
PKD1L3
PKD2
PKD2L1
PKD2L2
TRPML1
TRPML3
TRPML3
TRPML2

**Figure 12.2. Repertoire of TRP genes among S*phenodon punctatus* and other six vertebrate species.** Small red squares on nodes indicate gene duplications, blue boxes indicate gene differential retentions and duplicated boxes indicate species-specific recent duplicates. Empty spaces are differential losses.

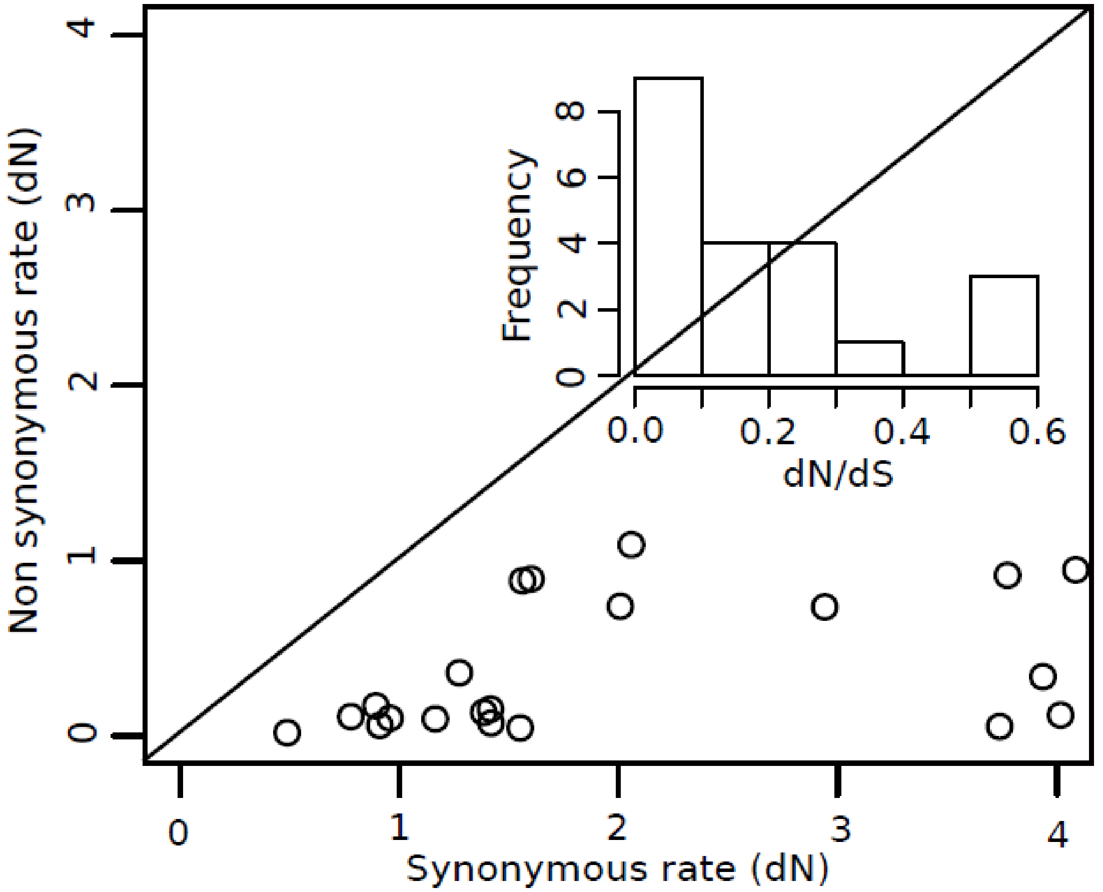

**Figure 12.3. Pairwise comparisons of dN-dS rates of TRP genes between *Sphenodon* and *Anolis* and between *Sphenodon* duplicates.** (Right upper corner) distribution of Omega values.

### 12.5 Tables

**Table 12.1. Genomic composition of TRP genes identified in *Sphenodon punctatus* and other six species of vertebrates.**

| Species | ML | P(PKD) | M | A | N | C | V |
|---|---|---|---|---|---|---|---|
| *Anolis* | 3 | 8 | 8 | 1 | NF | 6 | 6 |
| *Vipera* | 4 | 10 | 3 | 1 | 6 | 3 | 4 |
| *Sphenodon* | 3 | 8 | 8 | NF | 5 | 7 | 6 |
| *Alligator* | 5 | 6 | 8 | NF | 1 | 9 | 4 |
| *Gallus* | 3 | 6 | 7 | 1 | NF | 6 | 5 |
| *Pelodiscus* | 3 | 7 | 9 | 1 | NF | 7 | 8 |
| *Homo* | 3 | 7 | 7 | 1 | NF | 6 | 6 |
| Contig (ScrUdWx)* | | | | | | | |
| 1447 | | | | | p-p | | |
| 1600 | 2-3 | | | | | | |
| 1875 | | | | | | | 1-2-3 |
| 550 | | | | | p-3 | | |
| 67_1, 67_2 | | 1L3-1L3-2 | | | | | |

NF: Not Found. *Here we show the TRP genes of Sphenodon that were found in tandem in different contigs. For instance, in contig ScrUdWx_1600 there were found in tandem genes TRPML2-TRPML3.

### 13 SELENOPROTEINS

Didac Santesmasses, Marco Mariotti and Roderic Guigó*

#### 13.1 The tuatara selenoproteome

Selenoproteins incorporate selenium in the form of selenocysteine (Sec), the 21st amino acid. The human genome encodes 25 selenoproteins whose roles include antioxidation, redox regulation, thyroid hormone synthesis, calcium signal transduction and others[202]. Sec is inserted in response to a UGA codon, normally a stop codon, within selenoprotein mRNAs. UGA recoding is directed by an RNA secondary structure termed SECIS located in the 3'UTR of eukaryotic selenoprotein mRNAs, and requires a dedicated machinery[202]. We identified 26 selenoprotein genes in the tuatara genome (Table 13.1), along with the machinery genes for their synthesis (Table 13.2), using Selenoprofiles[203]. The number of selenoproteins, and the protein families they belong to, broadly fit with our expectations according to other vertebrate genomes previously analyzed[204]. One unexpected finding was the presence of two selenoproteins from the SELENOW (previously SelW) family. SELENOW is a short protein with unknown function that has a thioredoxin domain (Rdx family, PF10262). Two SELENOW selenoproteins, SELENOW1 and SELENOW2, were predicted to be present in the last common ancestor of vertebrates, though most genomes have only one of the two genes[204]. SELENOW2 was predicted to have been lost in the amniote stem (sauropsids and mammals). The presence of SELENOW2 in tuatara, and in other Lepidosauria genomes analyzed here, suggests that the loss of SELENOW2 must have occurred later in evolution, after the split of sauria and mammals. We could identify the SECIS element only in the SELENOW1 gene, but we detected the two genes in the transcriptome assembly, as evidence for their expression.

#### 13.2 Multiple tRNASec gene copies

The selenocysteine specific tRNA ($tRNA^{Sec}$) plays a central role in both biosynthesis and incorporation of Sec[202]. We identified four tRNA-Sec genes in the tuatara genome using Secmarker[205]. Other Lepidosauria genomes analyzed here had, instead, only a single $tRNA^{Sec}$. Therefore, the three extra copies, which were found in the same scaffold within a ~5 kb region, may be specific of tuatara. There is generally a single functional copy of $tRNA^{Sec}$ in genomes[202], and when additional copies are present they show characteristics of non-functional pseudo-genes[205]. In tuatara, all extra copies conserved the same predicted secondary structure, but possed a compensatory mutation in the acceptor stem and other mutations that were not expected to disrupt the pairing potential. Additionally, the upstream promoter elements necessary for tRNA$^{Sec}$ transcription[206] were present in all four tRNA$^{Sec}$ (Figure 13.1). These external promoter elements are not transcribed[207], hence none of the extra tRNA$^{Sec}$ copies appeared by retrotransposition. These observations suggest all four genes might be functional.

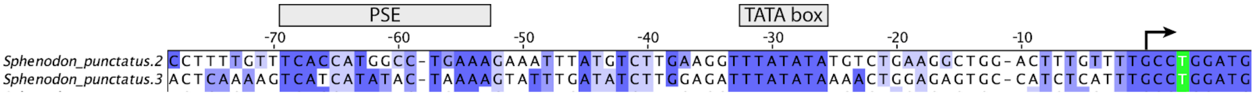

**Figure 13.1. Multiple sequence alignment of tRNA$^{Sec}$ from *Lepidosauria* and human.** The top four sequences, numbered from 1 to 4, were identified in the tuatara genome. The prediction number 1 is the direct ortholog of the tRNA$^{Sec}$ gene in other species, while the extra copies 2–4 are specific to this genome. The upstream promoter elements TATA box and the proximal sequence element PSE[206] are conserved in all tRNA$^{Sec}$, with the exception of the two bottom sequences. These correspond to the known pseudogene in human (Homo_sapiens.2)[208], and an additional prediction in the king cobra genome (Ophiophagus_hannah.2), which is also probably a pseudo-gene. Mutations in the tRNA are coloured according to the secondary structure (represented by arrows at the bottom); green corresponds to compensatory changes (mutations in both residues in a pair, maintaining the pairing potential), orange corresponds to single-sided variation (mutation in one of two paired residues, with pairing maintained), and red corresponds to disrupting changes (mutations that break pairings). The blue coloured scale corresponds to sequence conservation.

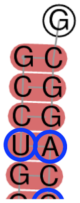

**Figure 13.2. Predicted secondary structure of one of the extra copies of tRNA-Sec in tuatara.** All three extra copies had the same predicted structure. The sequence corresponds to Sphenodon_punctatus.2 in Figure 13.1. Circled in blue are the different residues compared to the ancestor gene, Sphenodon_punctatus.1 in Figure 13.1.

**Table 13.1. The 26 Selenoprotein genes in the tuatara genome.** The protein family, the scaffold of the genome assembly, and the corresponding transcript identifiers are indicated, when available. Transcripts marked with a star contain the in-frame UGA-Sec codon. Otherwise their sequence might be incomplete.

| Family | Genome | Transcriptome |
|---|---|---|
| DI | scaffold_2907.3419 | |
| DI | scaffold_7697.7712 | c171299_g1_i1 |
| DI | scaffold_8323.8272 | |
| GPX | scaffold_3968.4331 | |

| | | |
|---|---|---|
| GPX | scaffold_452.452 | c196142_g1_i1* |
| GPX | scaffold_629.629 | c60729_g1_i1* |
| GPX | scaffold_6905.6973 | c145789_g1_i1,c28440_g1_i1* |
| MSRB1 | scaffold_26505.25541 | c154665_g2_i1* |
| SELENOF | scaffold_129.129 | c149446_g1_i1* |
| SELENOH | scaffold_3111.3595 | c195195_g1_i1 |
| SELENOI | scaffold_270.270 | c153700_g1_i2*,c153700_g1_i1*,c153700_g1_i3* |
| SELENOK | scaffold_100.100 | c166617_g1_i1 |
| SELENOM | scaffold_10583.10360 | c139842_g1_i1* |
| SELENON | scaffold_89.89 | c165428_g1_i1,c129907_g1_i3,c52835_g1_i1,c129907_g1_i1,c129907_g1_i2 |
| SELENOO | scaffold_53.888 | c147028_g1_i1* |
| SELENOP | scaffold_4793.5110 | c128991_g1_i1* |
| SELENOS | scaffold_239.1054 | c110652_g1_i2,c110652_g1_i1 |
| SELENOT | scaffold_19238.18510 | |
| SELENOT | scaffold_3522.3949 | c140842_g1_i1* |
| SELENOU | scaffold_1074.1784 | c129718_g1_i2,c129718_g1_i3*,c129718_g1_i1* |
| SELENOW | scaffold_4241.4607 | c153750_g4_i2*,c153750_g4_i1* |
| SELENOW | scaffold_4981.5251 | c48917_g1_i2*,c48917_g1_i1* |
| SPS2 | scaffold_8800.8726 | c2589_g1_i2*,c144616_g1_i2,c144616_g1_i1,c2589_g1_i1* |
| TR | scaffold_121.954 | c130940_g1_i2,c118043_g1_i2*,c118043_g1_i1,c130940_g1_i1 |
| TR | scaffold_2078.2679 | c143772_g2_i1* |
| TR | scaffold_7984.7975 | c148842_g1_i1*,c148842_g1_i2 |

**Table 13.2. Proteins from the selenoprotein synthesis machinery.** The corresponding scaffold and transcript identifiers are indicated.

| Family | Genome | Transcriptome |
|---|---|---|
| eEFsec | scaffold_434.434 | c155634_g2_i1 |
| pstk | scaffold_814.1566 | c144447_g1_i1,c126277_g1_i1,c126277_g1_i2,c126277_g1_i3 |
| SBP2 | scaffold_2348.2913 | c154056_g1_i1,c154056_g1_i2 |
| secp43 | scaffold_1648.2293 | c150861_g1_i2,c150861_g1_i3,c150861_g1_i1 |
| SecS | scaffold_3344.3788 | c144130_g1_i1,c144130_g1_i2 |

### 14 THE TUATARA GENOME AND A COMPARISON OF DNA SUBSTITUTION RATES ACROSS AMNIOTES

Marc Tollis* and David Winter

#### 14.1 Introduction

The tuatara (*Sphenodon punctatus*) is a well known example of a "living fossil": the sole representative of an ancient reptilian group that has existed virtually unchanged since the Mesozoic Era. As a member of the lepidosaurian order Rhyncocephalia - the sister taxon to all living squamates - the tuatara has a conserved tetrapod phenotype, and its fossil record shows that the modern species does not differ significantly from more ancient relatives. In addition, the tuatara is a long-lived species with a relatively late onset of sexual maturity (12–22 years) and reported lifespans up to 120 years. The combination of extreme morphological conservation and long generation times in the tuatara suggests strong conservation at the genomic level, as DNA substitution rates, generation times and phenotypic divergence are often correlated in reptiles[209]. Meanwhile, the rate of DNA substitution along the tuatara branch with respect to other amniotes is a controversial subject. For instance, one study used ancient DNA specimens and a demographic model to estimate that the tuatara has the fastest recorded substitution rate among studied amniotes[210]. These results suggested a decoupling of phenotypic divergence and the rate of DNA substitution. However, these conclusions remain under considerable debate[211,212]. Clearly, more data in a larger comparative context will shed light on whether or not the tuatara embodies a paradox of molecular evolution. Here, we leverage the tuatara genome assembly, as well as those of 26 additional tetrapods, in a comparative genomic approach to estimate substitution rates across amniotes. We aim to test the hypothesis that slow life history and phenotypic divergence in tuataras are underpinned by a similarly slow divergence at the genomic level.

#### 14.2 Whole genome alignments

We constructed 26 pairwise whole genome alignments using the green anole lizard genome (*Anolis carolinensis*, anoCar2.0) as a reference. The selection of query species was meant to represent an even phylogenetic sampling of the major groups of amniotes (lepidosaurs, birds, crocodilians, turtles and mammals), including *Sphenodon*, with *Xenopus tropicalis* as an outgroup (Table 1). Each query species' genome assembly was aligned to the reference using

LASTZ v1.02[213] to create the initial alignments, followed by "chaining" to construct gapless blocks in the alignment and "netting" to rank and select the highest scoring chains, utilizing the HOXD55 substitution score matrix[214]. Final pairwise alignments were combined into a multiple alignment using MULTIZ v11.2[215], using the following guide tree: (Xenopus, ((Ornithorhynchus, (Monodelphis, ((Loxodonta, Dasypus), ((Homo, Mus), (Canis, Bos))))), ((((Struthio, ((Gallus, Anas), (Melopsittacus, (Taenopygia, Geospiza)))), (Alligator, (Crocodylus, Gavialis))), (Chrysemys, Pelodiscus)), (Sphenodon, (((Anolis, Pogona), ((Python, Boa), (Crotalus, Ophiophagus))), Gekko))))) which is uncontroversial based on the literature[73,216–218]. The multiple genome alignment was filtered to contain blocks with data for 24 out of 27 species.

#### 14.3 Data extraction

To extract genetic features from the whole genome alignment, we downloaded the coding sequence annotations for the reference species *A. carolinensis* (anoCar2.0, Ensembl v86, last accessed August 2016). We then extracted sequence information for fourfold degenerate sites based upon the codon positions in the multiple genome alignment using msa_view in PHylogenetic Analysis with Space/Time Models (PHAST[219]).

#### 14.4 Phylogenetic analysis

We reconstructed the phylogeny of the 27 tetrapods included in our analysis using the fourfold degenerate site data as a single data partition in RAxML v8.2.3[116]. We generated 20 maximum likelihood (ML) trees under the GTRCAT substitution model and conducted 500 bootstrap replicates to assess statistical support of the best ML tree. Given that this phylogeny was constructed using fourfold degenerate site data, which should evolve at close to the neutral rate, we used the topology of the best ML tree to obtain a nonconserved model of evolution for the substitution rate analysis using phyloFit in PHAST, by fitting the tree model to the data using a time-reversible substitution model (REV).

#### 14.5 Substitution rate estimation

Using the topology and branch lengths obtained from the best ML phylogeny, we estimated absolute rates of molecular evolution in terms of substitution per site per million years and estimated the divergence times of amniotes via the semiparametric penalized likelihood (PL) method[220] with the program r8s v1.8[221]. We ran two separate PL analyses based on different node constraint criteria. The first set of criteria utilized fossil constraints: the minimum age of Diapsida (i.e., the time to most recent common ancestor or TMRCA of chicken and Anolis) was set at 255.9 My, the minimum age of Archosauria was set at 247.1 My, the minimum and maximum ages for Mammalia were set at 164.9–201.5 My, minimum and maximum ages for

Neognathae (i.e., the TMRCA of chicken and zebra finch) were set at 66–86.8 My, and the minimum and maximum ages for Serpentes (i.e., the TMRCA of python and cobra) were set at 98.32–113 My[222,223]. The second set of criteria utilized the median estimated ages obtained from www.timetree.org[224]: the TMRCA of sauropsids was fixed at 283 My, the TMRCA of lepidosaurs was fixed at 252 My, the TMRCA of Archosauria was fixed at 237 My, the TMRCA for Mammalia was fixed at 177 My, the TMRCA for Eutheria was fixed at 105 My. Across both analyses, we fixed the root of the tree at 352 My, the TMRCA of amniotes at 312 My, and the TMRCA of crocodilians at 80 My[73]. The PL method estimates a different substitution rate on each branch and relies on a "roughness" penalty when these rates differ greatly, quantified by a smoothing parameter for which larger values indicate a clock-like model[220]. For each set of node constraint criteria, we used cross-validation to optimize the smoothing parameter, allowing values to range on a log10 scale starting from 100 with the exponent increasing 0.3 for a total of ten steps, and reran the analysis with the optimal value. We also used the gradient check implemented in r8s v1.8 to ensure the signs of any active constraints were correct (i.e. negative if a minimum constraint was used) and estimated divergence times for all unconstrained nodes.

#### 14.6 Results

The filtered whole genome alignment was 790,566,929 bases in length, with 17.2% gaps and an average of 25.7 species represented in each alignment block, and we extracted 818,968 fourfold degenerate sites. The ML phylogeny included a topology consistent with expectations: most notably that Sphenodon was the sister taxon to all included squamates, with 100% bootstrap support for all branches (Figure 14.1). Node constraints utilizing both criteria passed all gradient checks, and the TMRCA for Sphenodon and squamates (i.e. Lepidosauria) was highly similar across the two methods (Figures 14.2 and 14.3) at 251.2 and 252 My, respectively. The optimal value for the smoothing parameter was one, indicating a significant deviation from the molecular clock. The analyses using different constraint criteria differed slightly in their absolute estimated substitution rates, but not in relative rates. Across analyses, the tuatara resulted in the slowest estimated substitution rate among lepidosaurs (Figure 14.4).

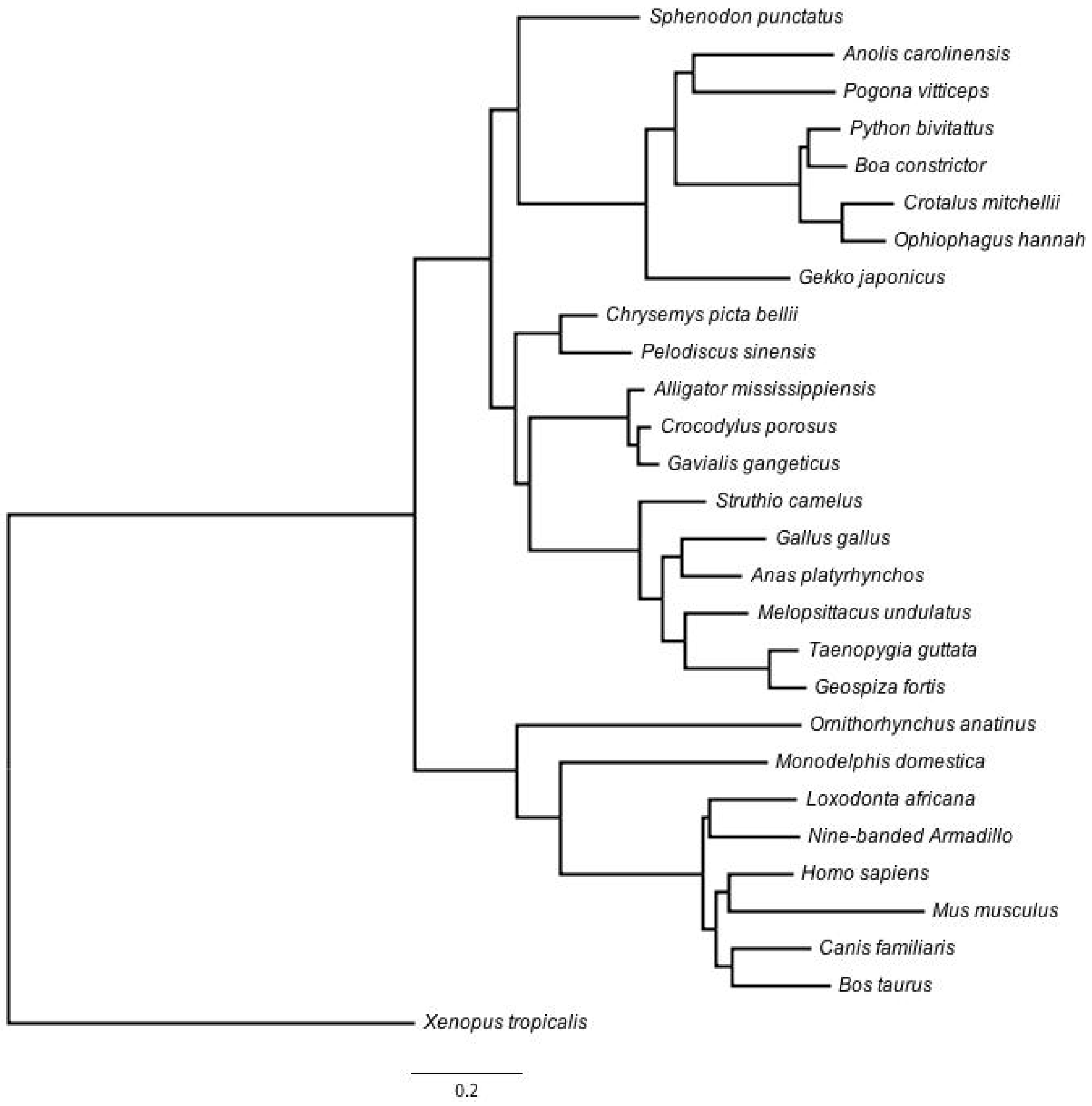

**Figure 14.1. Maximum likelihood phylogenetic reconstruction of 27 tetrapods using 818,968 fourfold degenerate sites.** We obtained full support for all branches using 500 bootstrap replicates.

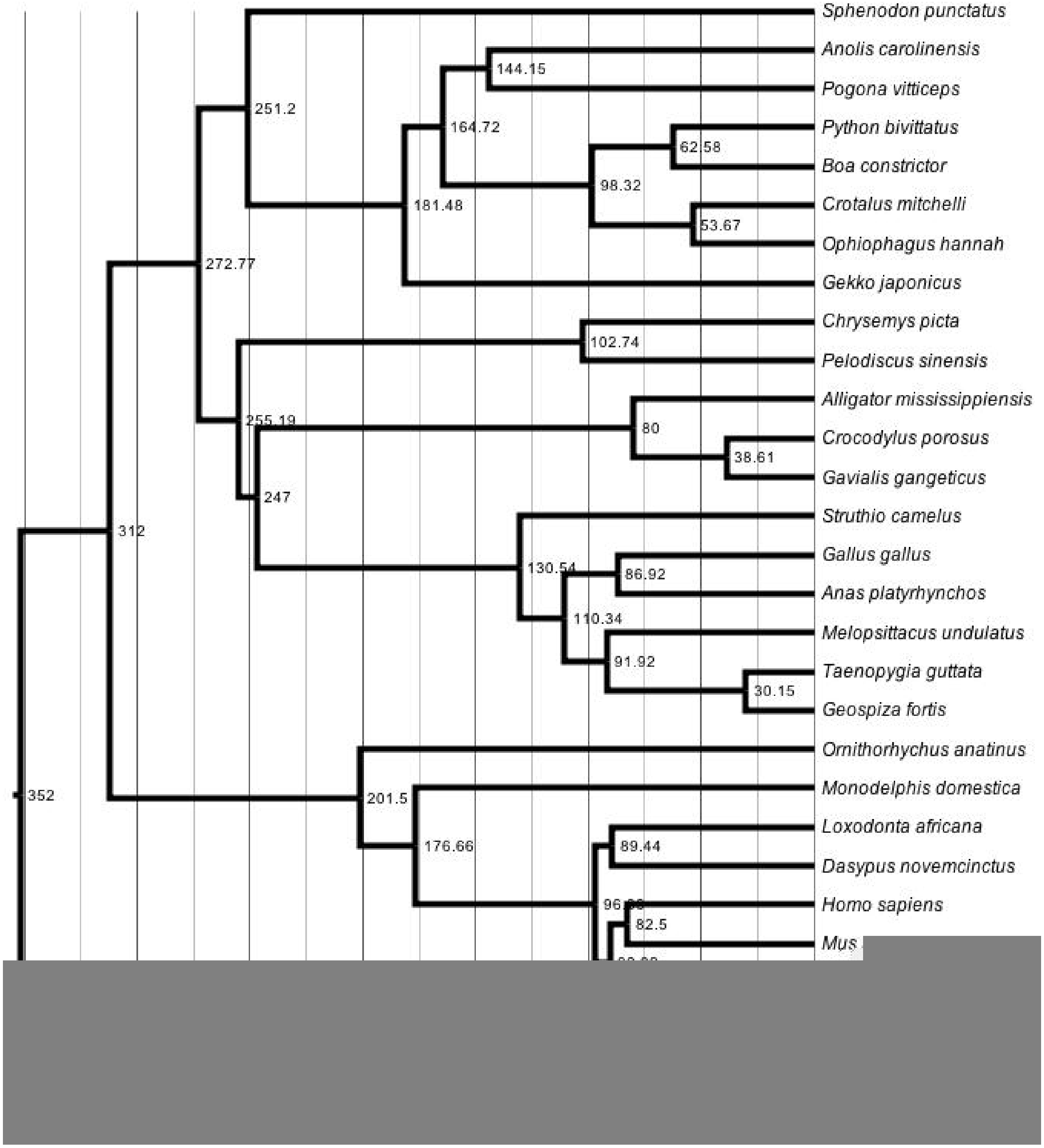

**Figure 14.2. Chronogram of amniote evolution obtained from penalized likelihood estimation of the phylogeny derived from fourfold degenerate sites and fossil calibrations.** Node labels indicating ages and x-axis are given in millions of years.

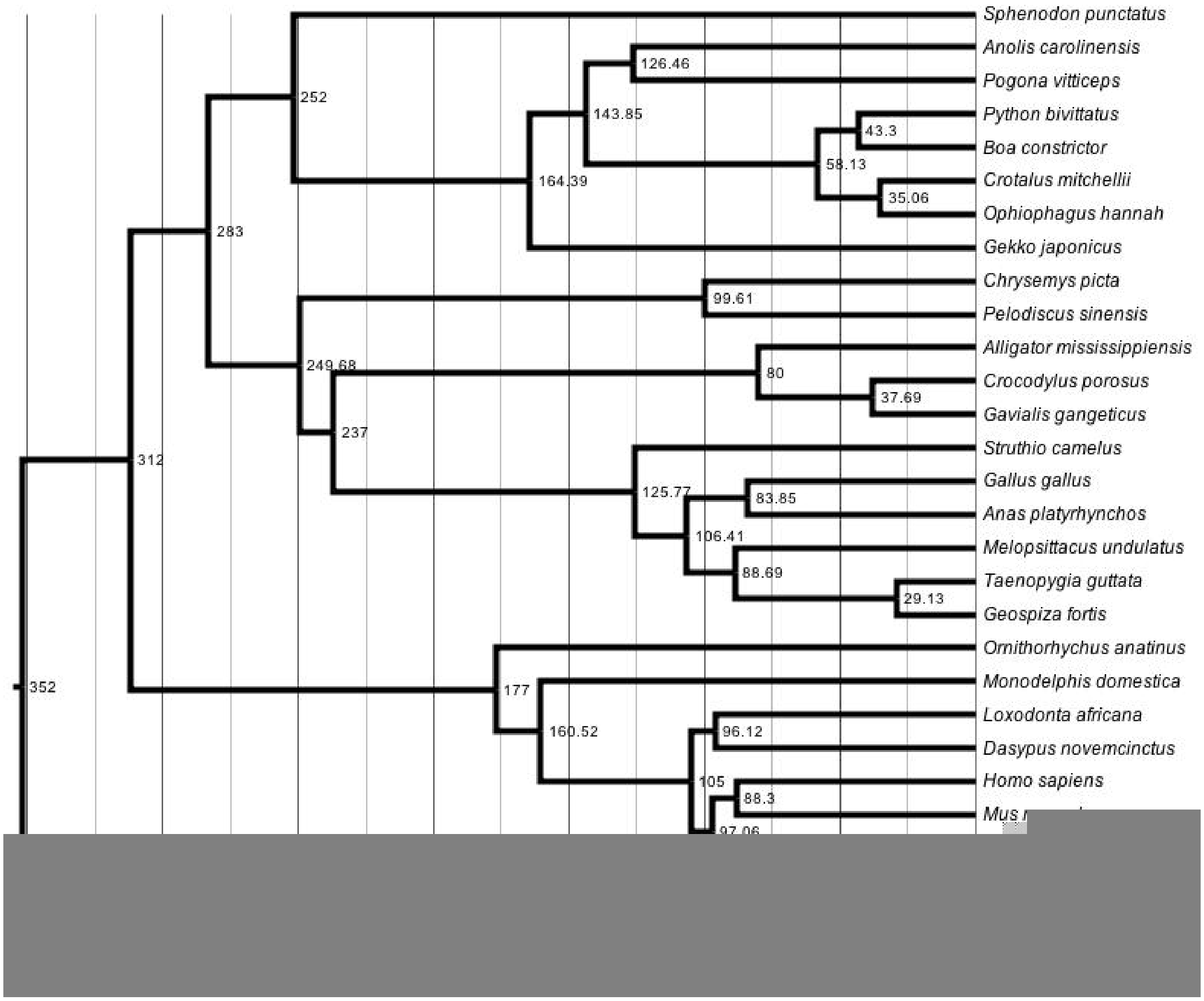

**Figure 14.3. Chronogram of amniote evolution obtained from penalized likelihood estimation of the phylogeny derived from fourfold degenerate sites and timetree calibrations.** Node labels indicating ages and x-axis are given in millions of years.

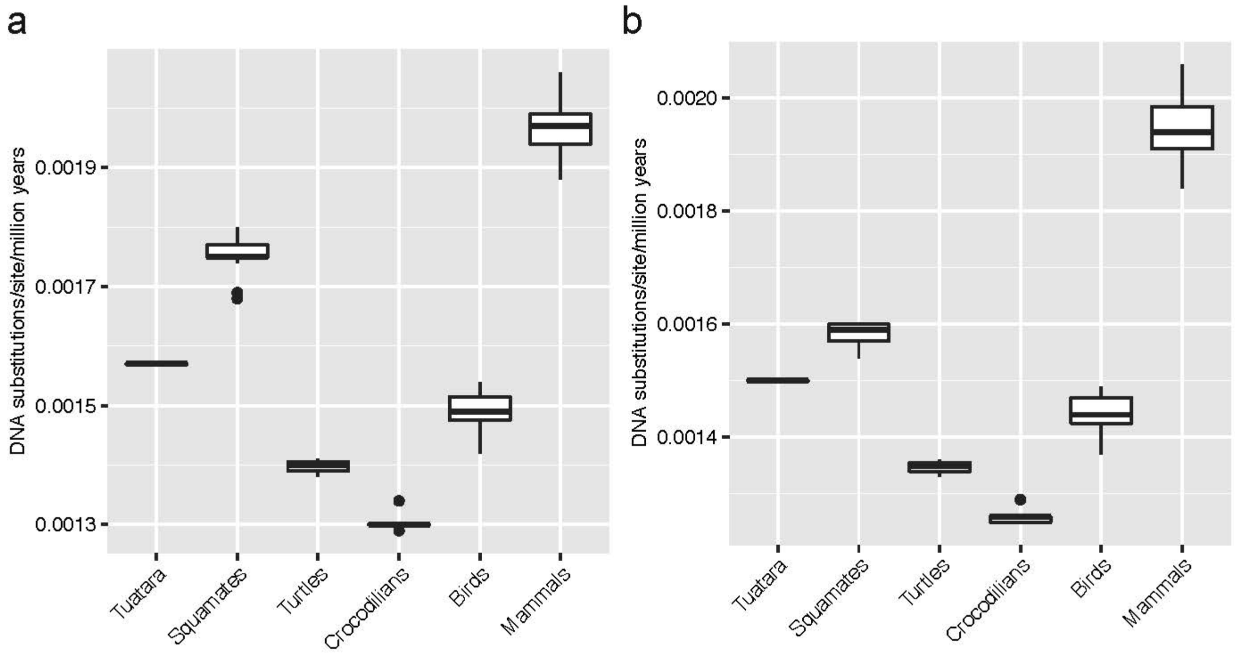

**Figure 14.4. Estimated DNA substitution rates of amniote clades based on fourfold degenerate sites.** Boxplots showing distribution of estimated substitution rates by clade using semiparametric penalized likelihood in r8s with (a) fossil constraints from Benton et al. [222] and Head [223] and (b) median TMRCA estimates from timetree.org.

**Table 14.1. Major clades, species, genome assemblies and sources used for the whole genome alignments in this study.** *reference sequence; UCSC, University of California Santa Cruz Genome Browser; NCBI, National Center for Biotechnology Information

| Clade | Common Name | Scientific Name | Genome Assembly Version | Repository |
|---|---|---|---|---|
| Lepidosaurs | Tuatara | *Sphenodon punctatus* | tuatara_30Sep2015_rUdWx | Current |
| | Green anole | *Anolis carolinensis* | AnoCar2.0* | UCSC |
| | Burmese python | *Python molurus bivittatus* | pitBiv5.0.2 | NCBI |
| | Speckled rattlesnake | *Crotalus mitchelli* | CroMitch1.0 | NCBI |
| | King cobra | *Ophiophagus hanna* | OphHan1.0 | NCBI |
| | Schlegel's Japanese gecko | *Gekko japonicus* | Gekko_japonicus_V1.1 | NCBI |
| | Boa constrictor | *Boa constrictor* | snake_7C_scaffolds | DOI: 10.1186/2047-217X-2-10 |
| Birds | Common ostrich | *Struthio camelus* | STRCAM | GigaDB |
| | Chicken | *Gallus gallus* | galGal3 | UCSC |
| | Duck | *Anas platyrhyncos* | anaPla1 | UCSC |
| | Zebra finch | *Taenopygia gutatta* | taeGut3 | UCSC |
| | Medium ground finch | *Geospiza fortis* | geoFor1 | UCSC |
| | Budgerigar | *Melopsittacus* | melUnd1 | UCSC |

| | | | | |
|---|---|---|---|---|
| Crocodilians | Saltwater crocodile | *Crocodylus porosus* | croc_sub2 | www.crocgenomes.org |
| | Indian gharial | *Gavialis gangeticus* | ggan.v0.2 | www.crocgenomes.org |
| | American alligator | *Alligator mississippiensis* | allMis1 | UCSC |
| Turtles | Chinese softshell turtle | *Pelodiscus sinensis* | PelSin_1.0 | NCBI |
| | Western painted turtle | *Chrysemys picta bellii* | Chrysemys_picta_bellii-3.0.3 | NCBI |
| Mammals | Gray short-tailed opossum | *Monodelphis domestica* | monDom5 | UCSC |
| | Platypus | *Ornithorhynchus anatinus* | ornAna1 | UCSC |
| | House mouse | *Mus musculus* | mm10 | UCSC |
| | Domestic dog | *Canis lupus familiaris* | canFam3 | UCSC |
| | Cow | *Bos taurus* | bosTau8 | UCSC |
| | African savannah elephant | *Loxodonta africana* | loxAfr3 | UCSC |
| | Nine-banded armadillo | *Dasypus novemcinctus* | dasNov3 | UCSC |
| | Human | *Homo sapiens* | hg19 | UCSC |
| Amphibian | Western clawed frog | *Xenopus tropicalis* | xenTro3 | UCSC |

### 15 PHYLOGENETIC ANALYSIS OF VERTEBRATE SINGLE-COPY ORTHOLOGS

Marc Tollis* and Stefan Prost

We analyzed 245 single copy orthologs present in 26 vertebrates, including tuatara, which were aligned by the codon-based PRANK algorithm[225] and concatenated and partitioned with FASconCAT-G v1.02[226]. Using RAxML v8.2.3[116], we generated 20 maximum likelihood trees from the 245 partitions under the GTRGAMMA substitution model, and selected the tree with the highest likelihood (Figure 15.1). We assessed branch support on the best tree using 1,000 bootstrap replicates.

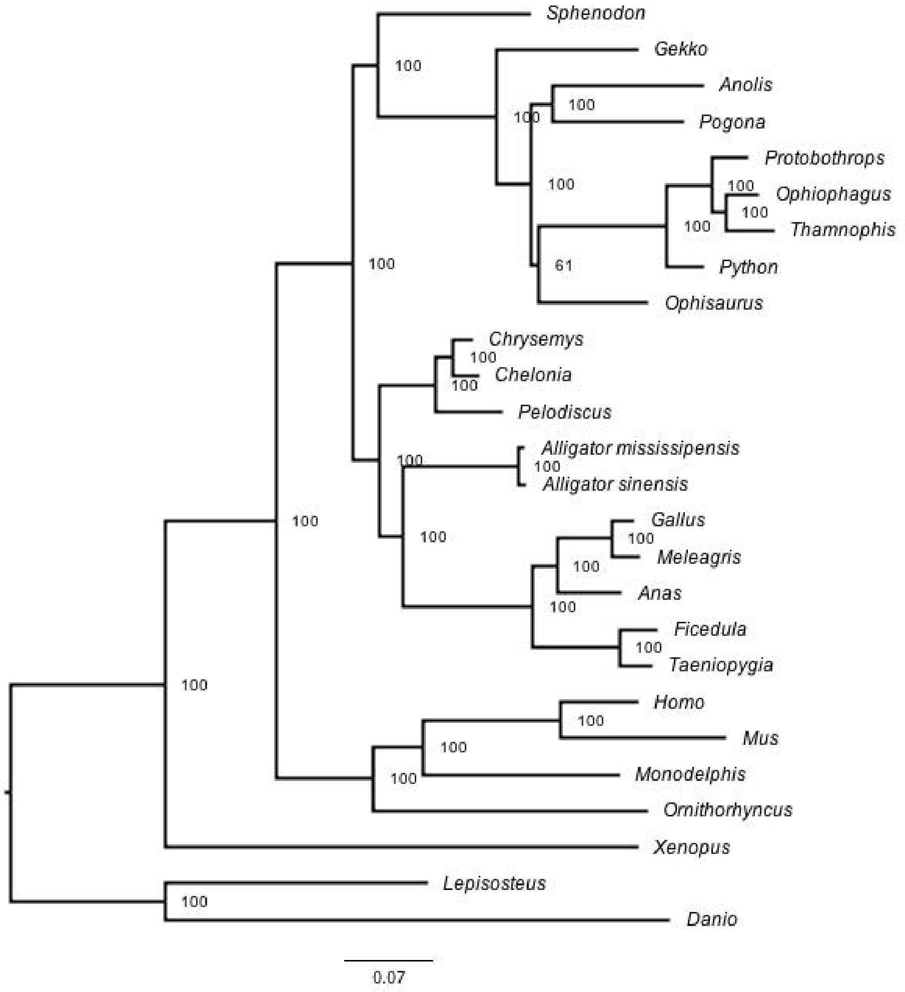

**Figure 15.1. Maximum likelihood phylogeny estimated from a concatenated supermatrix of 245 single copy ortholog partitions.** Branch lengths are in terms of substitutions per site. Branch support from 1,000 bootstrap replicates is shown.

We also generated gene trees based on the 245 one-to-one orthologs found between all species. To do so, each aligned gene sequence was first analyzed phylogenetically using RaxML[116]; using the GTRGAMMA model). Subsequently, we binned all gene trees to reconstruct a species tree (Figure 15.2) using Astral[227], assessing branch support using local posterior probabilities[228]. Astral applies binning of gene trees with similar topologies, based on an incompatibility graph between gene trees. Subsequently, it chooses the most likely species tree under the multi-species coalescent model.

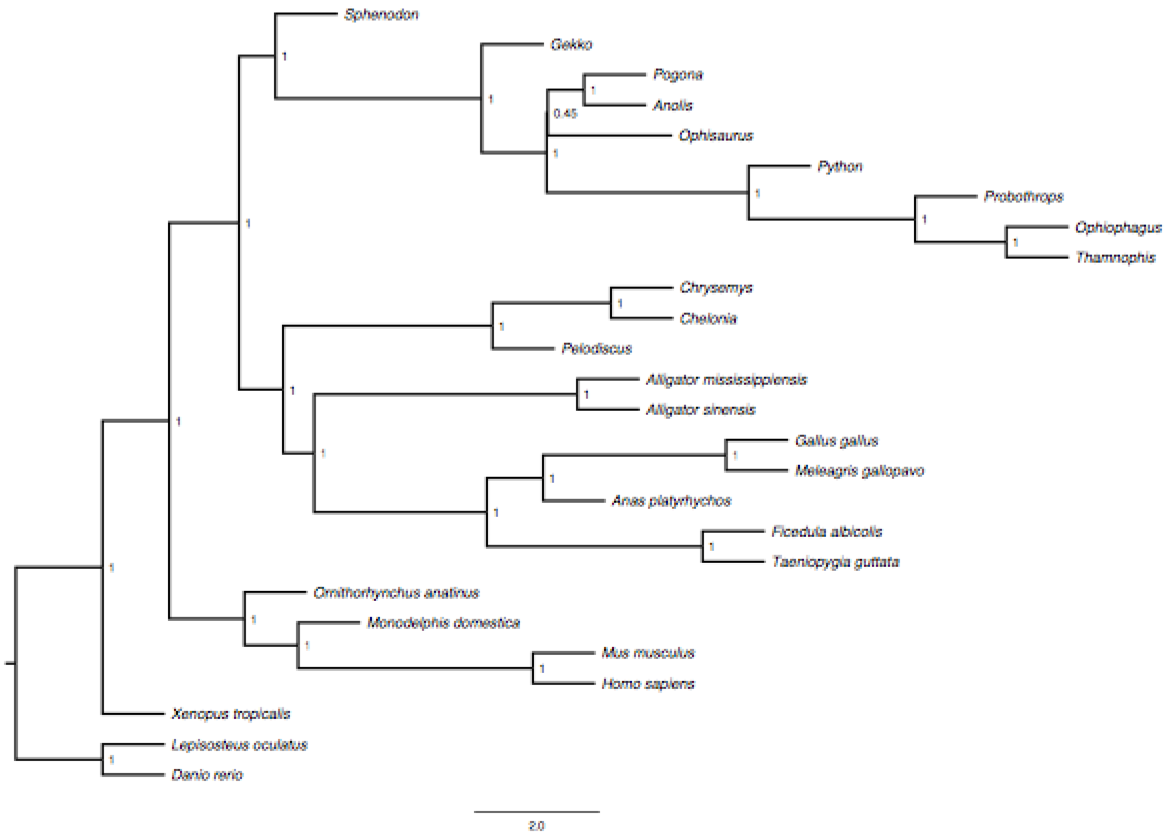

**Figure 15.2. Species tree reconstruction using 245 single copy orthologs.** Branch lengths are given in terms of coalescent units. Local posterior probabilities are given for each branch.

### 16 AN ANALYSIS OF DIVERGENCE TIMES AND TEST FOR PUNCTUATED EVOLUTION

Chris L. Organ

#### 16.1 Divergence times

We inferred time-calibrated phylogenies with BEAST v2.4.8[229] using the CIPRES Science Gateway[80]. We randomly sampled 50 of the 245 one-to-one ortholog alignments generated for the gene tree reconstruction (Supplementary Materials 15). This alignment had 166,307 sites, which were partitioned using a GRT model with a four-category gamma distribution of rate variation. The proportion of invariant sites was set to 0.1 and the nucleotide state frequencies was set to empirical. We used a calibrated Yule prior for the tree with a gamma prior (alpha=0.001, beta=1,000) on the birth rate. We also implemented a relaxed log normal clock – the prior for the mean and standard deviation of the branch rates were drawn from gamma distributions (alpha=0.001, beta=1,000; and alpha=0.5, beta=1). Given the general uncertainty surrounding the ancestry of fossil lineages, we chose to be conservative and used normal priors on node calibrations drawn from established resources[222,223] as follows: Archosauria (m=254, s=2.0), Diapsida (m=276, s=7.0), Mammalia (m=183.2, s=7.0), Neornithes (m=76.5, s=4.0), Root/Vertebrata (m=444.5, s=8.0), Serpentes (m=105.5, s=3.0), and Tetrapoda (m=344, s=3.0). The MCMC chain ran for 20,000,000 generations with a sampling frequency of 1,000. The analysis was repeated using a new random gene set four times, which, along with the ESS parameter reported in BEAST, was used to assess convergence.

#### 16.2 Punctuated evolution

Evolution is a temporally heterogeneous process. The rate of evolution is expected to vary through time and across taxonomic groups, especially if change is associated with speciation events, a process known as punctuated evolution[230]. The process of punctuated genome evolution is expected to produce a pattern of correlation between the amount of genome evolution and the net number of speciation events along a lineage. We used Bayesian phylogenetic generalized least squares to regress the total phylogenetic path length (of four-fold degenerate sites) on the net number of speciation events (nodes in a phylogenetic tree)[231]. We find strong evidence for punctuated evolution at these sites (pMCMC $\approx$ 0; 100% of the posterior probability for the line's slope was greater than 0, the null value). Thirty-nine percent (95% credible interval = 0.15 to

0.62) of branch length variation is attributable to punctuated evolution. Furthermore, punctuated evolution accounts for 33.5% ($r^2$; 95% credible interval = 0.34 to 0.38) of deviation from the molecular clock at four-fold degenerate sites (Figure 16.1).

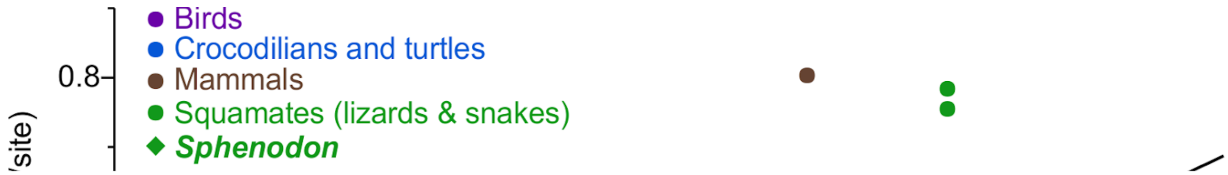

**Figure 16.1. Punctuated genome evolution.** Note that the *Sphenodon* genome has undergone less evolution than expected given its history of net speciation.

As a "living fossil", we might expect Sphenodon to be an outlier in this analysis, with rates below the phylogenetic average. To test this, we removed Sphenodon and inferred another punctuated regression model (as described above) to predict its path length using its phylogenetic position and its net number of speciation events[232]. We predict a path length of 0.44 substitutions/site (95% credible prediction interval = 0.29 to 0.58) for *Sphenodon*, which does not differ substantially from its inferred length of 0.38 substitutions/site. The rate of *Sphenodon* evolution is 0.0008 substitutions/site/millions of years. The only more slowly evolving genomes are those of basal Archelosauria (turtles and crocodilians), which have an average rate of 0.00056 substitutions/site/millions of years. Previous analyses of the rate at which genomes evolve have not accounted for punctuated evolution. An analysis similar to previous studies (of path lengths but not including nodes) supports *Sphenodon* as having undergone less evolution than other species (posterior predictive median = 0.52, 95% credible prediction interval = 0.36 to

0.68, pMCMC = 0.045). Together, these models support the hypothesis that the slowly evolving genome of *Sphenodon* is attributable to its 240-plus million-year history without substantial diversification.

These results could be explained by underestimated branch lengths in regions of a phylogeny that have few species (either because of taxon sampling or extinction, or both). Multiple substitutions won't be detected along these paths[233], but this artifact can be detected[233,234]. We tested for this artifact (http://www.evolution.reading.ac.uk/pe/index.html) but found no evidence for its presence.

### 17 PATTERNS OF SELECTION ON TUATARA ORTHOLOGS

Shawn M. Rupp, Victoria G. Twort, Thomas R. Buckley*, Melissa A. Wilson*,

We performed two analyses on ortholog sets from squamate genomes. These analyses were focused on investigating patterns of selection among squamate genomes and comparing patterns of molecular evolution at sex determining genes relative to other orthologs.

#### 17.1 Methods

##### 17.1.1 Positive selection on the tuatara lineage

The first analysis was targeted at detecting positive selection along the branch leading to tuatara. One-to-one orthologs were extracted from the ortholog file and those absent in the tuatara were excluded. The resulting ortholog sets were realigned using a codon-based method. First, nucleotides were translated to amino acids using Translatorx[235], then aligned using MAFFT[115,236], using the auto option to select the most appropriate alignment strategy. The corresponding codon alignment was constructed from the amino acid alignment using PAL2NAL[237]. Alignments were then screened using a set of custom scripts and TrimAl[238] to check the frame, replace stop codons and remove gapped regions. Only alignments with 12 or more of the 17 species were included. Gene trees reconstructed in Garli under the GTR+I model.

Alignments were then analysed in CodeML from the PAML[162] package using the tuatara branch as the foreground branch. Gapped sites were included in these analyses. The branch model was employed. The resulting p-values were corrected with FDR of 5% with the R package Qvalue[239]. Gene ontology (GO) and KEGG enrichment tests analyses were carried out with Kobas[240]. A Fisher's exact test was used to assess significantly overrepresented GO and KEGG categories in the test set of tuatara orthologs. The Anolis, mouse and human annotations were used as references with the tuatara orthologs used as the test data set. Transcripts were identified using Blastx against the nr database.

##### 17.1.2 Patterns of molecular evolution at sex determining genes

The second analysis compared patterns of molecular evolution at sex determining genes relative to other orthologs. We used multiz[213] to create a multiple alignment of six of the species previously aligned to the green anole (anoCar2). These species included the tuatara (sphPun), gecko (gekJap1), bearded dragon (pogVit1), Burmese python (pytBiv1), speckled rattlesnake (croMit1), and king cobra (ophHan1). This resulted in an alignment of seven species (including the anole) comparing the tuatara and six squamate species.

To obtain aligned coding sequences, we submitted the multiple alignment to Galaxy's Stitch Gene Clocks Tool[241] using the BED12 genome annotation for the green anole (anoCar2) imported from the UCSC Table Browser[242]. We submitted the resulting CDS fasta alignment to AlignmentProcessor0.12 (https://github.com/WilsonSayresLab/AlignmentProcessor) on Arizona State University's Ocotillo cluster to perform quality control and run CodeML[162] on each gene alignment. CodeML[162] was run twice for each gene: once for the null model in which dN/dS was assumed to be zero, and again for the alternative model in which the tuatara was specified as the forward branch and dN/dS was allowed to vary for each branch. Next, we subset the CodeML[162] output to include only 1 to 1 orthologs using the database generated previously for this project.

We compiled CodeML[162] output for each model and conducted a Likelihood Ratio Test in R3.3.1[144] using the log likelihood values generated by CodeML[162]. We then compared overall dN, dS, dN/dS, and tree lengths between sex-determining genes and non-sex-determining genes for gene alignments with a minimum of three species which contained the tuatara. We calculated mean and median values in R3.3.1[144] and determined 95% confidence interval using R boot package[243] using 1,000 bootstrap replicates. We calculated p values both using Wilcoxon Rank Sum Tests and permutation analyses with 10,000 replicates using in lab scripts.

Lastly, we calculated the branch lengths of the tuatara branch and the species with the longest branch length (other than the tuatara) and merged this data with the likelihood ratio test output to compare the tuatara's branch length to the overall tree length excluding the tuatara. We examined significant genes to see whether the tuatara showed a pattern of faster evolution when compared to the rest of the phylogeny. Genes were considered "faster" than the rest of the tree if the tuatara had a greater branch length than the squamate species with the longest branch length and were considered "slower" if the tuatara's branch length was less than the longest squamate branch. We calculated the percentage of significantly faster, significantly slower, and not significantly different genes for sex-determining genes, non-sex-determining genes, and all genes.

#### 17.2 Results

##### 17.2.1 Positive selection on the tuatara lineage

Extended analyses through an independent pipeline yielded a total of 4284 ortholog sets which were analysed in CodeML. Of these 659 orthologs had corrected p-values $<0.05$ from the likelihood ratio test of the equal $\omega$ across branches versus a model where the tuatara (foreground) branch had a different $\omega$ value to shared $\omega$ across all other (background) branches. None of the tests with significant p-values have a foreground ω>1. Supplementary Data 1 .https://zenodo.org/record/2583274

Using the *Anolis*, human and mouse annotations a total of 14,435 and 11 KEGG pathways were significantly overrepresented, respectively. With human and mouse annotations a total of 1,910 and 73 GO categories were overrepresented. The KEGG pathways from the comparison to *Anolis* include a number of RNA and mRNA regulation and metabolic pathways as well as general metabolism. Supplementary Data 2. https://zenodo.org/record/2583274

##### 17.2.2 Patterns of molecular evolution at sex determining genes

We find that most genes exhibit a pattern of molecular evolution that suggests the tuatara branch evolves at a different rate than the rest of the tree (Table 17.1). When comparing genes involved in the sex-determination pathways (Table 17.2), we find that both the entire length of the tree is shorter for these genes (Table 17.3) and that dN and dN/dS values are significantly lower for genes involved in sex determination than typical genes in the genome (Table 17.4; Figure 17.1).

**Table 17.1. Tuatara–squamate gene tree evolution compared between all genes and sex determining genes.** A log-likelihood test comparing the likelihood of a model where the rate of evolution was the same on all branches compared to the likelihood of a model where there was a different rate of evolution on the tuatara branch was conducted for all genes present in both the tuatara and anole (the reference for the alignment) and were present in a minimum of three species. The log likelihood was calculated for each gene for both the null and alternative model in CodeML[162] and a likelihood ratio test conducted in R3.3.1[144] using alpha of 0.05 and 1 degree of freedom.

| | | **Genes which Are Significantly Different from the Null Model** | **Total Number of Genes** |
|---|---|---|---|
| **All Genes** | **Percentage** | 75.91% | 9,586 |
| | **Number of Genes** | 7,277 | |
| **Sex Determining Genes** | **Percentage** | 84.0% | 25 |
| | **Number of Genes** | 21 | |
| **Non-Sex-Determining Genes** | **Percentage** | 75.89% | 9,561 |
| | **Number of Genes** | 7,256 | |

**Table 17.2. Sex determining gene substitution rates.** We computed substitution rates, $d_N$, $d_S$, and $d_N/d_S$, using PAML[162] for genes previously identified as playing a primary role in sex determination. For eight sex determining genes we did not identify homologous sequence in the tuatara in the multiple species alignment (highlighted in gray); which does not mean that they don't exist in the tuatara genome, but that they weren't in the current version of the multiple alignment. This could have been due to high divergence.

| TranscriptID | GeneSymbol | dN | dS | dNdS | Tree Length | Tuatara branch length |
|---|---|---|---|---|---|---|
| ENSACAT00000009605 | AR (DHTR) | 0.0900 | 2.0497 | 0.0439 | 1.9847 | 0.0571163 |
| ENSACAT00000008263 | ARX | 0.0749 | 1.4746 | 0.0508 | 1.3473 | 0.0348331 |
| ENSACAT00000000495 | CIRBP | 0.0461 | 2.4627 | 0.0187 | 2.1876 | 0.0636031 |
| ENSACAT00000004647 | CSDC2 | 0.0933 | 1.4834 | 0.0629 | 1.5226 | 0.106651 |
| ENSACAT00000003256 | CSDE1 | 0.0464 | 0.8424 | 0.0551 | 0.8258 | 0.0575427 |
| ENSACAT00000015874 | DHCR7 | 0.3298 | 1.6549 | 0.1993 | 2.1308 | 0.134717 |
| ENSACAT00000010360 | EMX2 | 0.1142 | 1.8239 | 0.0626 | 1.7465 | 0.0348258 |
| ENSACAT00000006243 | ESR1 | 0.1894 | 1.4322 | 0.1322 | 1.6464 | 0.113636 |
| ENSACAT00000001292 | FEM1B | 0.0809 | 2.3919 | 0.0338 | 2.2297 | 0.0435242 |
| ENSACAT00000015028 | JAKMIP1 | 0.3526 | 2.9505 | 0.1195 | 3.2637 | 0.056045 |
| ENSACAT00000014908 | JAKMIP2 | 0.0494 | 1.5791 | 0.0313 | 1.4699 | 0.0361767 |
| ENSACAT00000008030 | JAKMIP3 | 0.0968 | 1.2884 | 0.0751 | 1.3204 | 0.095763 |
| ENSACAT00000002517 | LHX1/LIM1 | 0.1131 | 2.3564 | 0.0480 | 2.2445 | 0.0276626 |
| ENSACAT00000014613 | LHX9 | 0.0500 | 0.9320 | 0.0536 | 0.8877 | 0.0480989 |
| ENSACAT00000030595 | M33/TUSC3 | 0.0155 | 1.4471 | 0.0107 | 1.2941 | 0.0125199 |
| ENSACAT00000014130 | PAX2 | 0.0271 | 0.8004 | 0.0339 | 0.7732 | 0.0621367 |
| ENSACAT00000030504 | PDGFB | 0.3974 | 1.2046 | 0.3299 | 1.9014 | 0.280557 |
| ENSACAT00000017280 | RSPO1 | 0.2540 | 1.0624 | 0.2391 | 1.4557 | 0.192456 |
| ENSACAT00000016485 | SF1 | 0.0415 | 2.0755 | 0.0200 | 1.8859 | 0.00712568 |
| ENSACAT00000008756 | SOX9 | 0.1188 | 1.8735 | 0.0634 | 1.8038 | 0.129618 |
| ENSACAT00000018021 | STAT3 | 0.0357 | 2.1405 | 0.0167 | 1.9370 | 0.0282673 |
| ENSACAT00000005401 | UBA1 | 0.1646 | 1.9622 | 0.0839 | 2.0987 | 0.11493 |
| ENSACAT00000012577 | WNT4 | 0.4560 | 2.9148 | 0.1564 | 3.2907 | 0.0161123 |
| ENSACAT00000002516 | WT1 | 0.0986 | 1.2456 | 0.0792 | 1.3415 | 0.110817 |
| ENSACAT00000009898 | ZFPM2/FOG 2 | 0.2839 | 1.3392 | 0.2120 | 1.7684 | 0.227515 |
| ENSACAT00000015597 | AMH | 0.5398 | 1.9654 | 0.2747 | 2.8312 | NA |
| ENSACAT00000002340 | DMRT1 | 0.0895 | 0.7924 | 0.1129 | 0.8935 | NA |
| ENSACAT00000001907 | GATA4 | 0.1677 | 1.5525 | 0.1080 | 1.6969 | NA |
| ENSACAT00000007239 | MSL2 | 0.0344 | 0.5761 | 0.0597 | 0.5901 | NA |
| ENSACAT00000010492 | SDC1 | 0.7469 | 1.5435 | 0.4839 | 2.9324 | NA |
| ENSACAT00000006298 | STAT1 | 0.1039 | 0.8107 | 0.1282 | 0.9141 | NA |
| ENSACAT00000025425 | STAT2 | 0.3812 | 1.1877 | 0.3210 | 1.8272 | NA |
| ENSACAT00000009792 | STAT6 | 0.2880 | 1.1722 | 0.2457 | 1.6266 | NA |

**Table 17.3. Tuatara–squamate tree lengths for sex determining genes.** Here we compare tree lengths from the alternative models. All genes examined were present in both the tuatara and anole and were present in minimum of three species. For table A, 95% confidence intervals calculated with 1000 bootstrap replicates. P values were calculated using a Wilcoxon rank sum test.

| | | Sex Determining Genes | All Other Genes | P Wilcoxon Test |
|---|---|---|---|---|
| **dN** | **mean** | 0.1448 (0.1295, 0.1571) | 0.2495 (0.2426, 0.2864) | 0.0014 |
| | **median** | 0.0968 (0.0909, 0.1029) | 0.2051 (0.1487, 0.1993) | |
| **dS** | **mean** | 1.712 (1.733, 1.868) | 1.876 (1.857, 2.273) | 0.3976 |
| | **median** | 1.579 (1.455, 1.712) | 1.735 (1.662, 1.803) | |
| **dN/dS** | **mean** | 0.0893 (0.0799, 0.0978) | 0.2670 (0.000, 1.391) | 0.0024 |
| | **median** | 1.774 (1.695, 1.831) | 0.1175 (0.1019, 0.1304) | |
| **Tree Length** | **mean** | 1.774 (1.738, 1.870) | 2.147 (1.886, 2.279) | 0.0298 |
| | **median** | 1.768 (1.665, 1.779) | 1.986 (1.801, 2.000) | |
| **Number of Genes** | | 25 | 9561 | |

**Table 17.4. Log-likelihood of tuatara branch length comparisons across gene trees.** All genes examined were present in the tuatara, anole, and at least one additional species and had log likelihood values and tree lengths for both the null and alternative models. Genes were identified as significant or not using a likelihood ratio test comparison between a null (equal rate) and alternative (unequal rate) model of the substitution rate on the tuatara branch length.

| | | Tuatara Branch is Significantly Shorter | Tuatara Branch is Significantly Longer | Tuatara Branch is not Significantly Different | Total Number of Genes |
|---|---|---|---|---|---|
| **Sex Determining Genes** | **Percentage** | 44.0% | 40.0% | 16.0% | 25 |
| | **Number of Genes** | 11 | 10 | 4 | |
| **Non-Sex Determining Genes** | **Percentage** | 36.44% | 39.34% | 24.11% | 9,561 |
| | **Number of Genes** | 3,484 | 3,761 | 2,305 | |
| **All Genes** | **Percentage** | 49.83% | 39.34% | 24.09% | 9,586 |
| | **Number of Genes** | 3,495 | 3,771 | 2,309 | |

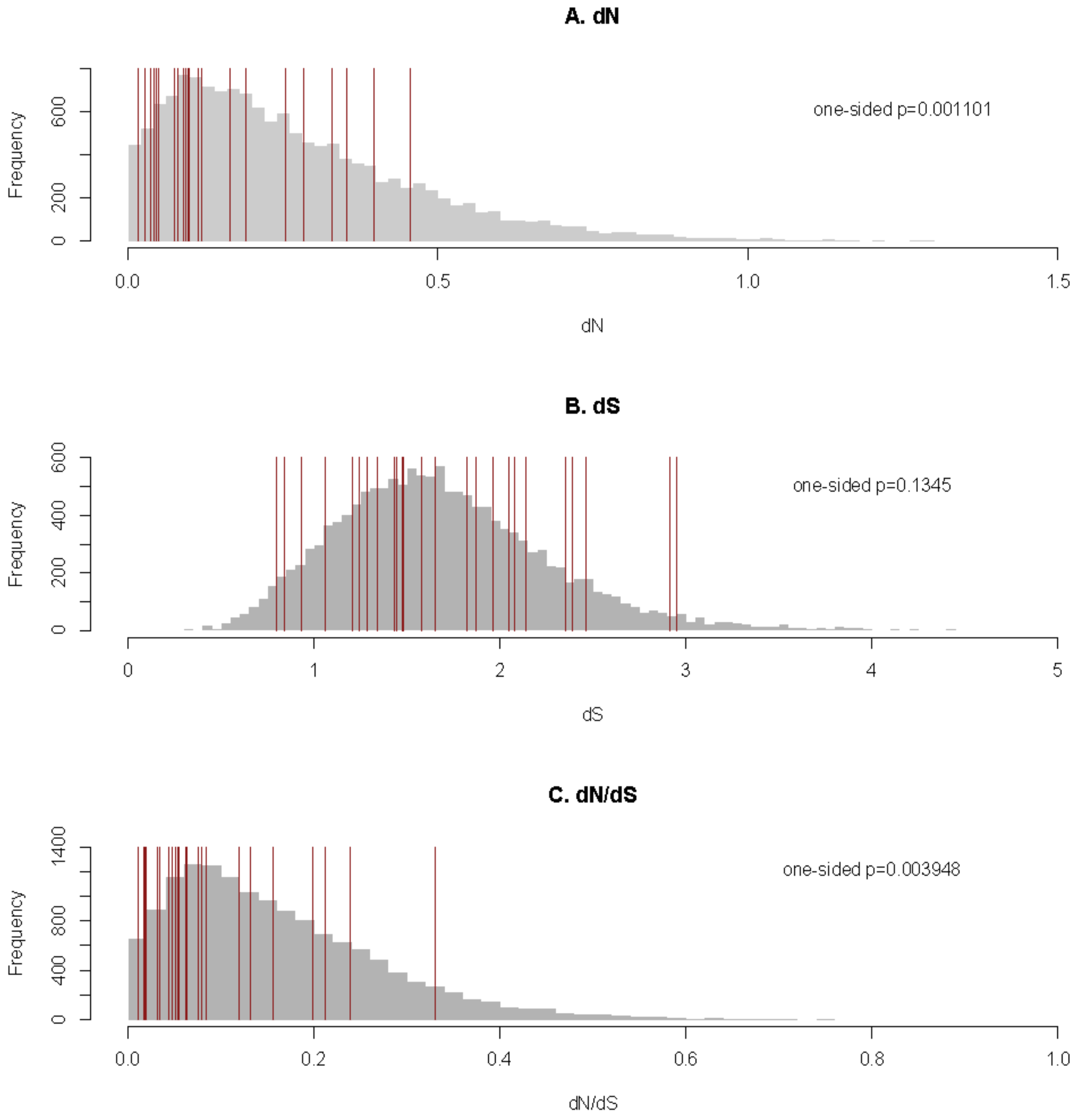

**Figure 17.1. Distribution of dN, dS, and dN/dS values for sex-determining genes versus all genes in the seven-way species alignment.** The seven-species multiple species alignment includes tuatara (sphPun), gecko (gekJap1), bearded dragon (pogVit1), Burmese python (pytBiv1), speckled rattlesnake (croMit1), and king cobra (ophHan1) aligned to green anole (anoCar2). We also show the p-value from a one-sided Wilcoxon Rank Sum testing the hypothesis that the value of dN, dS, or dN/dS for the sex-determining genes (marked in red) is less than the rest of the genome (in gray).

#### 17.3 Discussion

Our analyses show significant deviation between patterns of molecular evolution at the codon level between tuatara and other reptiles and birds. Approximately 15% of the orthologs we tested had significantly different $\omega$ values on the tuatara branch relative to other birds and reptiles tested. Although none of these orthologs had $\omega$ values >1, suggestive of strong positive selection, the results do indicate that shifts in patterns of selection are affecting many genes and functional categories of genes across the tuatara genome.

Further, our analysis of sex-determining genes is consistent with an overall pattern of extreme conservation of genes involved in sex determination. Specifically, even with a biased subset of genes that align, and are conserved across species, including tuatara, sex-determining genes are especially conserved in terms of overall substution rate, as well as in dN/dS. This suggests that they may be more likely to also be conserved in terms of function.

### 18 RECONSTRUCTION OF THE DEMOGRAPHIC HISTORY OF THE TUATARA

Stefan Prost*, Jose Grau* and Neil Gemmell

#### 18.1 Methods

Demographic history was inferred from the diploid sequence of our tuatara genome using a pairwise sequential Markovian coalescent (PSMC) method[244]. To do so we generated a diploid genome by mapping the raw read data back to the tuatara de novo assembly (QEPC00000000.1) using BWA mem[32] and called the diploid consensus sequence using Samtools mpileup[245]. We then conducted a range of preliminary analyses and found that PSMC plots were not sensitive to the values chosen for the maximum number of iterations (N), the number of free atomic time intervals (p), the maximum time to the most recent common ancestor (t), and the initial value of ρ. Based on these investigations, our final PSMC analyses used values of $N = 25$, $t = 5$, $\rho = 1$ and $p = 4 + 25 \times 2 + 4 + 6$ (data not shown). We determined the variance in estimates of Ne using 100 bootstrap replicates.

Plots of demographic history were scaled using a generation time of 30 years, slightly shorter than the 32-44 years estimated by[246], and estimates of mutation rates of 1.4 and $2x10^{-8}$ based on the divergence time between Anolis and tuatara and substitutions per site per generation obtained from our analysis of 4-fold degenerate sites (Supplementary Materials section 14).

#### 18.2 Results

The reconstructed demographic histories did not vary substantially between the different values of p, t and ρ parameters. Using the 'optimal' settings we recovered a fluctuating population trend of repeated population increases and decreases. More specifically, we found an increase in effective population size ($N_e$) dating back to about 10 Mya ago, a drastic decrease in $N_e$ about 1–3 Mya, and a subsequent rapid increase in $N_e$ between 500 ka and 1 Mya.

#### 18.3 Discussion

Our analyses show a significant fluctuations in effective population size, with increases detectable around 10 Mya, a decreases about 1–3 Mya, and a further rapid increase occurring between 500 kya and 1 Mya. These events correlate well with the known geological history of New Zealand (Zealandia), which experienced a pronounced marine transgression during the

Oligocene (ca. 30–21 Mya)[247]. At its peak, ~23 Mya, Zealandia likely consisted of a series of small island masses, each separated by significant marine water bodies. Endemic terrestrial biota incapable of dispersing across these major marine barriers, like the tuatara, would have been geographically isolated during this period, with populations only reconnecting as the landmasses subsequently enlarged and land bridges formed. The increase in effective population size at around 10 Mya may be reflective of the larger landmass and thus habitat available for tuatara during that period. The subsequent reduction in effective population size 1–3 Mya is consistent with a period of climatic cooling , which may have limited tuatara population size, while the rapid increase in effective population size 500 ka to 1 Mya seems strongly concordant with the formation of land bridges that connected the two main island land masses 1.5–2 Mya [248].

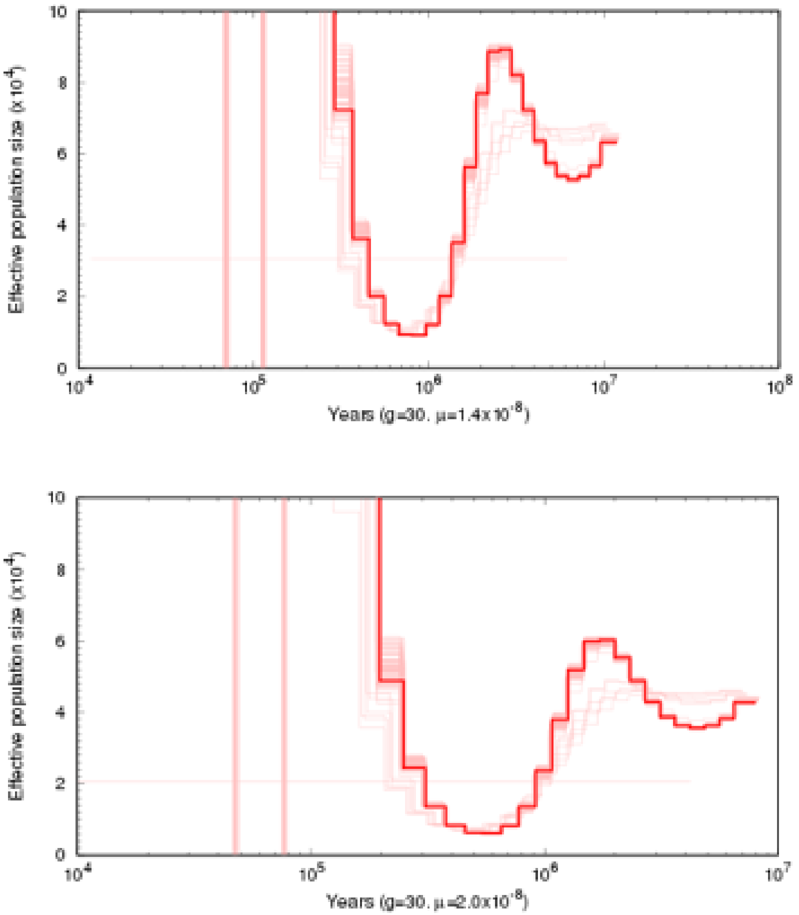

**Figure 18.1. PSMC plots of the demographic history of tuatara.** Top: using a mutation rate of 1.4X10-8 substitutions per site per generation. Bottom: using a mutation rate of 2X10-8 substitutions per site per generation.

### 19 POPULATION GENOMICS ANALYSES

David Winter*, Nicolas Dussex, Helen Taylor, Neil Gemmell*

#### 19.1 Methods

##### 19.1.1 Sample collection and library preparation

We obtained 30 blood samples previously collected for animals from Hauturu (Little Barrier Island), Takapourewa (Stephens Island) and Nga Whatu Kai Ponu (North Brother Island) (Table 19.1). Samples were collected under Victoria University of Wellington Animal Ethics approvals 2006R12; 2009R12; 2012R33; 22347 and held and used under permit 32037-RES issued by the New Zealand Department of Conservation. We extracted DNA using NucleoSpin Blood QuickPure columns (Macherey-Nagel) and evaluated DNA quality and concentration using both a Nanodrop 2000c spectrophotometer (Thermo Fisher Scientific) and a Fluorometric Quantitation Qubit system (Thermo Fisher Scientific). We undertook genotyping-by-sequencing (GBS)[249] on 28 samples where DNA quality and quantity were sufficient, using the restriction enzymes ApeKI and MspI. GBS libraries were prepared as described in[250], but further purified the amplified library utilising a Pippin Prep (SAGE Science, Beverly, Massachusetts, United States) to select the DNA sequencing library in the size range of 150–500 bp. We performed single-end sequencing (1x100 bp) on an Illumina HiSeq2500 utilising v4 chemistry at AgResearch, Invermay, New Zealand, which yielded approximately 25Gb of raw sequence data per lane.

##### 19.1.2 Read mapping and genotyping

Sequencing reads were de-multiplexed using GBSX[251] before being mapped to the tuatara genome with bwa mem[252]. Only reads with a mapping quality score > 30 that did not produce split or secondary alignments where used for downstream analysis. Because the results of GBS analysis can be sensitive to the particular analysis methods used, we took two separate approaches to calling single nucleotide variants (SNVs) from our data. We first used the reference mapping pipeline from Stacks 1.4.4[253], a software package designed for reduced-representation sequencing data. We also used the gatk haplotypecaller[254,255], a general-purpose variant caller that uses a more sophisticated genotyping model than stacks, to call SNVs. We filtered the gatk data to remove loci that were multiallelic, those with evidence for strand bias (odds ratio > 4) and those for which there was a significant difference in the quality scores of reads containing reference and alternative alleles (z-score > 5). For both datasets we minimized the effects of allelic dropout by retaining only those loci that could be genotyped in at least four individuals in all three populations.

##### 19.1.3 Functional annotation of SNVs

We used snpEFF[256] along with the functional annotation produced in this work to generate information of each SNV in the gatk dataset. We used pyVCF[257] to isolate SNVs falling in protein coding genes or close to putative mRNA splice sites. This subset of the data was then used to calculate alternative allele frequency spectra for both synonymous and non-synonymous variants in protein coding genes. We also examined the allele frequency of those SNVs that introduce and in-frame stop codon.

##### 19.1.4 Population genomic analysis

We used vcftools[258] and the "population" program included in stacks to generate summary statistics for both the gatk and STACKS SNV dataset. Population differentiation was measured using Weir and Cockerham's estimator of $F_{ST}$[259], having first thinned each dataset to contain at most one SNV per 10kb of the reference genome. We calculated $F_{ST}$ for the complete dataset and, additionally, for each pairwise combination of population-samples.

Our GBS experiment produced relatively low coverage alignments. For this reason, we incorporated the uncertainty associated with calling genotypes from low-coverage data[260] in our population genomic analysis by using genotype likelihood methods as implemented in angsd[261] and related software. We estimated a covariance matrix from genotype likelihoods using pcangsd[262], then used the R programming language[144] to perform a principal component analysis from this matrix. Admixture proportions for each individual were estimated using NGSadmix[261] with the value of K (the number of hypothetical populations from which individuals could inherit their genome) set between 2 and 4. Nucleotide diversity was calculated for each population using angsd.

##### 19.1.5 Differentiation with respect to sex

We tested our SNV data for genetic differentiation with respect to the sex using the "population" program in STACKS. $F_{ST}$ was calculated for each SNV while treating male and female tuatara as separate populations. The p-values obtained from the STACKS pipeline were transformed to q-values (i.e. controlled for multiple comparisons using a false-discovery rate approach) using the R function "p.adjust".

#### 19.2 Results

##### 19.2.1 Both SNV datasets support strong population structure

We generated a total of 22 gigabases of sequencing from our 28 individuals (Table 19.1).

The gatk haplotypecaller produced considerably more SNVs from this data than the stacks pipeline (Table 19.2). This result arises from the higher depth-threshold required to call a site in stacks, which is also reflected in a much greater depth of coverage for called SNVs in the stacks dataset. The stacks dataset also has a much higher among-site variance in sequencing depth and more evidence for reference bias when compared to the gatk data. Both of these results suggest the stacks data may contain artifactual variant calls[263], possibly because the coverage thresholds used by this software can be expected to enrich the called-variants for multicopy and paralogous sequences when applied to low coverage datasets. For this reason we report only analyses from the gatk data in the main paper. Nevertheless, the evidence for strong population structure in modern tuatara populations is supported by both datasets ($F_{ST} = 0.42$ in the stacks data, 0.45 in the gatk data).

Pairwise $F_{ST}$ estimates support the distinctiveness of the North Brother Island population, as reported in the main paper. The smallest pairwise $F_{ST}$ was recorded between the Stephens and Little Barrier populations ($F_{ST} = 0.29$), while comparisons with the North Brother Island produced higher estimates ($F_{ST} = 0.51$ when combined with Little Barrier and 0.43 for Stephens).

##### 19.2.2 No loci are differentiated by sex

After corrected for multiple comparison no SNV was significantly differentiated with regard to sex (Figure 19.1), the lowest q-value obtained for any locus was 0.41.

##### 19.2.3 No evidence for excess genetic load in any population

The results presented in the body of this work demonstrate that modern tuatara populations have relatively low genetic diversity and have been isolated to small islands for thousands of years This history of isolation may have implications for the management of remaining tuatara populations, as small isolated populations are prone to accumulating deleterious mutations. The population on Nga Whatu Kai Ponu / North Brother Island is of particular concern in this regard, given its low genetic diversity and the fact it is currently managed as a separate conservation unit. The generally low coverage of our GBS data prevents us from estimating inbreeding coefficients, even using genotype likelihood approaches. We can, however, directly test for an excess of potentially damaging alleles in any population.

Non-synonymous mutations in protein coding genes are predominantly deleterious, so we started by considering the occurrence and frequency of synonymous and non-synonymous variants in coding regions across all populations. In total, 1101 (1.2%) of our called SNVs fall into a coding sequence. There are slightly fewer non-synonymous variants (518) than synonymous ones (572). Given these classes of variant arise in an approximately 3:1 ratio, this result suggests natural selection is at least partially preventing the accumulation of protein-altering mutations. When we consider the frequency of variants of each class in each population (Figure 19.2), we find this result holds for alleles that are fixed within a population and for segregating variants.

SNVs that truncate a protein by introducing a premature stop codon in a protein coding gene are particularly likely to be deleterious. These SNVs are rare in our data, with only five such variants detected. Hauturu / Little barrier is the only population to have more than two such alleles, and all four truncating alleles that occur in this population occur at a low frequency (Figure 19.3)**.**

Although our GBS data only allows to consider a small number of potentially damaging variants, we find no evidence for an excess of such variants in any population. The population from Nga Whatu Kai Ponu / North Brother Island has less genetic diversity than other populations we consider, but we find no evidence that it has accumulated protein-altering mutations at a greater rate than either of the more diverse populations.

**Table 19.1. Tuatara samples used for population genetic analysis via genotype-by sequencing and epigenetics.** Samples in **BOLD** were not used for genotype-by-sequencing , while those in *Italics* sample not used for post-bisulfite adapter tagging (PBAT).

| Sample Name | ID | FT Number | Population | Sex | Status | Date Collected |
|---|---|---|---|---|---|---|
| *SI IM* | 486 | CD0516 | Stephens Island | Male | adult | 1/27/1984 |
| SI 2M | 487 | CD0517 | Stephens Island | Male | adult | 1/28/1984 |
| SI 3F | 488 | CD0518 | Stephens Island | Female | adult | 1/28/1984 |
| SI 4F | 490 | CD0520 | Stephens Island | Female | adult | 1/28/1984 |
| ***SI 5M*** | 491 | CD0521 | Stephens Island | Male | adult | 1/28/1984 |
| SI 6F | 492 | CD0522 | Stephens Island | Female | adult | 1/28/1984 |
| *SI 7F* | 493 | CD0523 | Stephens Island | Female | adult | 1/28/1984 |
| SI 8M | 494 | CD0524 | Stephens Island | Male | adult | 1/29/1984 |

| | | | | | | |
|---|---|---|---|---|---|---|
| SI 9F | 495 | CD0525 | Stephens Island | Female | adult | 1/29/1984 |
| SI 10M | 496 | CD0526 | Stephens Island | Male | adult | 1/29/1983 |
| LBI 1M | 2133 | FT2925 | Little Barrier Island | Male | adult | 2/1/1991 |
| LBI 2F | 2134 | FT2926 | Little Barrier Island | Female | adult | 2/1/1991 |
| LBI 3M | 2135 | FT2927 | Little Barrier Island | Male | adult | 2/1/1991 |
| LBI 4F | 2136 | FT2928 | Little Barrier Island | Female | adult | 2/1/1991 |
| LBI 5M | 2188 | FT2980 | Little Barrier Island | Male | adult | 3/23/1991 |
| LBI 6F | 2189 | FT2981 | Little Barrier Island | Female | adult | 3/23/1991 |
| LBI 7M | 2190 | FT2982 | Little Barrier Island | Male | adult | 3/23/1991 |
| LBI 8F | 2191 | FT2983 | Little Barrier Island | Female | adult | 3/23/1991 |
| LBI 9F | 2200 | FT2992 | Little Barrier Island | Female | adult | 3/23/1991 |
| ***LBI 10M*** | 2201 | FT2993 | Little Barrier Island | Male | adult | 3/23/1991 |
| NBRO 1M | 1025 | FT0198 | North Brother Island | Male | adult | 1/21/1988 |
| NBRO 2M | 1026 | FT0199 | North Brother Island | Male | adult | 8/21/1988 |
| *NBRO 3F* | 1033 | FT0206 | North Brother Island | Female | adult | 3/21/1989 |
| NBRO 4M | 1034 | FT0207 | North Brother Island | Male | adult | 10/21/1989 |
| NBRO 5M | 1035 | FT0208 | North Brother Island | Male | adult | 5/21/1990 |
| NBRO 6M | 1036 | FT0209 | North Brother Island | Male | adult | 12/21/1990 |
| NBRO 7F | 1039 | FT0212 | North Brother Island | Female | juvenile | 7/21/1991 |
| NBRO 8F | 1042 | FT0215 | North Brother Island | Female | adult | 2/21/1992 |
| NBRO 9F | 1051 | FT0224 | North Brother Island | Female | adult | 9/21/1992 |
| NBRO 10F | 1053 | FT0226 | North Brother Island | Female | adult | 4/21/1993 |

**Table 19.2 Summary statistics from SNV calling.** "N" is the number of variable sites. "Ts:Tv" is the ratio of transitions to transversions in the dataset, "Depth" provides the mean total depth SNVs in each dataset, with variance across sites given in parentheses. "Reference freq." is the mean frequency of reads containing the reference allele at each site. "$F_{ST}$" is the mean value of Weir and Cockerham's estimator of this measure across all sites.

| Daset | N | Ts:Tv | Depth | Reference freq. | $F_{ST}$ |
|---|---|---|---|---|---|
| STACKS | 19379 | 2.82 | 83.7 (1771) | 0.524 | 0.42 |
| gatk | 89256 | 2.79 | 47.6 (279) | 0.502 | 0.45 |

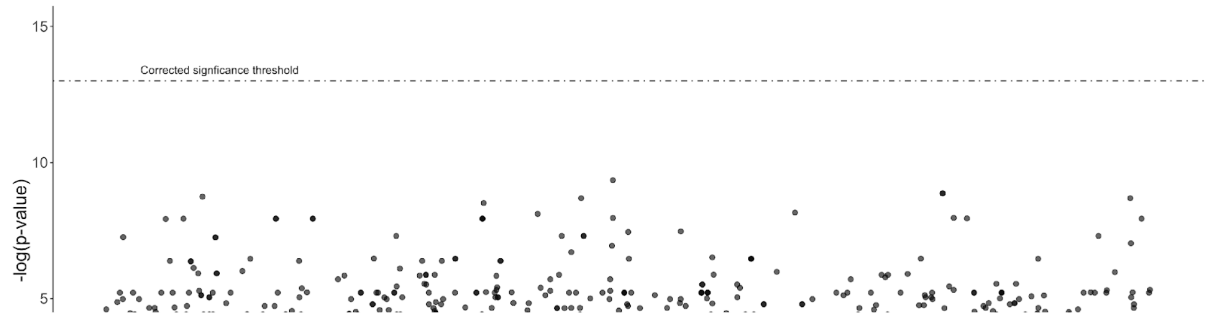

**Figure 19.1 No SNV is significantly differentiated with respect to sex.** Each point represents a p-value from a test of population differentiation for a single SNV. In this case, the sex of individual tuatara was used in place of a population. The dashed line represents the threshold for statistical significance after accounting for multiple testing. Note the y-axis is log-transformed.

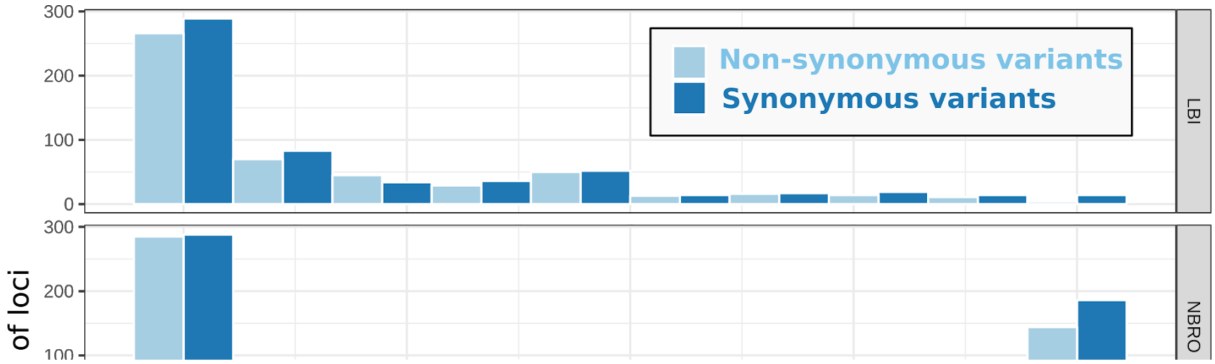

**Figure 19.2 No evidence for an excess of non-synonymous variants in any population.** Each sub-plot displays the allele frequency spectrum of SNVs in protein coding regions of the tuatara genome from one population. LBI = Little Barrier Island, NBRO = North Brother, SI = Stephens Is. In each case the spectrum is plotted separately for synonymous (dark blue) and non-synonymous variants (light blue). Note that Hauturu/Little Barrier Island population is from Northern New Zealand, and thus likely more closely related to the reference individual, leading to more sites fixed for the reference allele and fewer fixed for an alternative allele.

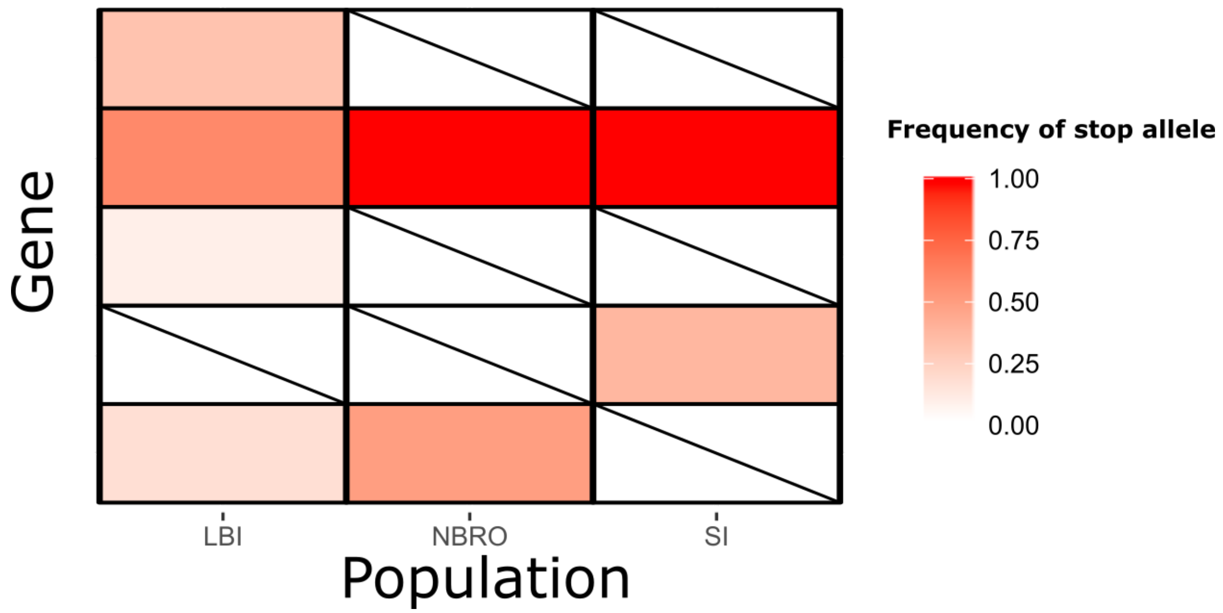

**Figure 19.3 Frequency of alleles that introduce a premature stop codon.** Each row represents a distinct gene and each column a population. Cells are shaded to reflect frequency of the stop-codon allele in each population, cells with a diagonal line have an allele frequency of zero.

### 20 GENOME RESEARCH AGREEMENT TEMPLATE

Between XXXXX and ZZZZZ

#### 1.0 Preamble

1.1 XXXXX and ZZZZZ (“the Researcher”), are undertaking a collaborative research project to sequence the genome of _______, that will increase our knowledge and understanding of _______ (“the Project”). The expected outcome of this collaboration is the publication and release of the ______ genome sequence to the international scientific community.

1.2 _______ are located on lands or waters to which XXXXX holds traditional title or has cultural significance to XXXXX.

1.3 ______ are species of special significant to XXXXX and must be respected by all people granted access to them. They must be protected at all times for the present and future i generations.

1.4 It is acknowledged that ______________are providing funding or services in kind for the Project.

#### 2.0 Purpose of Agreement

2.1 This Agreement sets out the terms and conditions for the Researcher participating in the Project. This Agreement will clearly identify the obligations of each of the parties towards meeting the Project outcomes.

#### 3.0 Obligations of the Parties

3.1 XXXXXX will ensure that the following obligations are met:

a) Appoint individuals to oversee the coordination and management of the project. Their responsibilities will include interacting with, and instructing the research team of decisions of XXXXXX concerning the study.

b) Ensure that appropriate Communication Protocols will be followed.

c) Participate in and support the project (workshop discussions, feedback on the project and the information gathered)

d) Approve unpublished reports, media releases, peer-reviewed publications, presentations, photographs, and all results and information related to the research for distribution to XXXXXX and the general public

e) Provide guidance and advice and support to the Researcher on all cultural matters where assistance is required.

f) Approve the sending of DNA, tissue samples and data to collaborating overseas laboratories for genome sequencing, assembly and analysis. Such approval will not be unreasonably withheld.

3.2 The Researcher will ensure that they comply with the following conditions:

a) Act in good faith and work professionally while carrying out their required work with XXXXX.

b) Ensure that confidentiality is maintained when they conduct their research;

c) Ensure that XXXXX participants are provided with all necessary advice and guidance when interpreting scientific results obtained as part of the project.

d) Discuss all aspects of their proposed work with XXXXX and seek approval before any work is undertaken on the islands, or organic or non-organic material is removed from the islands.

e) Where proposed work has been amended, then the Researcher must notify XXXXX of the change and seek approval before work is undertaken.

f) The Researcher will inform XXXXX when reports are sent to funders, and will provide XXXXX with the full scientific research information contained within these reports.

g) The Researcher will consult with XXXXX prior to the sending of DNA, tissue samples and data to collaborating overseas laboratories for genome sequencing, assembly and analysis.

h) Where other parties may wish to contribute to the project or obtain access to the data prior to its release into the public domain, such arrangements will be developed in consultation with XXXXX and further research agreements may be negotiated where appropriate.

i) Any information relating to XXXXX, and the cultural practices to which that knowledge relates, obtained by the Researcher remains the intellectual property of XXXXX.

j) The Researcher will not engage with or use XXXXX information or resources without seeking written consent to do so.

k) It is agreed by the Researcher that the aim of this research is to sequence the genome of _______ and not commercialisation resulting from this genome sequencing. However, should the researchers believe there may be potential for further research leading to commercialisation they must first consult with XXXXX for approval and for discussion on a benefit sharing agreement.

#### 4.0 Access to Information / Release of Scientific Results

4.1 The first point of contact for any results and interpretations from the study will be the individuals appointed by XXXXX to oversee the coordination and management of the project

4.2 XXXXX will be responsible for reviewing and providing prior written consultation, which may be given by email by the project coordinator appointed by XXXXX, for the release of research results and information, traditional knowledge, meeting minutes, general dialogue, and photographs in peer-reviewed publications, popular articles, interviews, presentations, and any other form of media.

4.3 XXXXX members will provide feedback regarding the release of all research results and information, traditional knowledge, meeting minutes, general dialogue, and photographs in peer-reviewed publications, popular articles, interviews and presentations to the appointed project coordinator.

The project coordinator appointed by XXXXX will notify the Researcher of XXXXX feedback within two weeks.

4.4 The Researcher must be willing to report the results of the research at meetings, annual Hui or any other gathering as directed by XXXXX.

#### 5.0 Media

5.1 No statements are to be made to the media, Department of Conservation or any other agency without the Parties consulting with each other over this first.

5.2 Any information contained in reports, media releases and articles that has previously been approved for release and is therefore already in the public domain, is no longer subject to clause 5.1.

#### 6.0 Confidentiality

6.1 Information disclosed by one party to the other concerning the Project and the arrangements between the Parties that are detailed in this Agreement are confidential.

6.2 Neither Party will disclose any confidential information to any third party, other than funders and approved collaborators without the express agreement of the other Party. This agreement will not be unreasonably withheld.

6.3 The Parties agree that confidential information received under this Agreement will not be used for any purpose other than carrying out the Project and any future subcontract or research agreements relating to the Project.

6.4 This confidentiality provision will survive the expiration or termination of this Agreement and shall be legally enforceable by one Party against the other, until that confidential information enters the public domain.

#### 7.0 Breach of terms and conditions of this Agreement

7.1 Where any terms within this agreement have been breached, then the following process will be followed:

a) XXXXX will be notified of the nature of the breach;

b) The Researcher or nominee will be asked to attend a meeting with XXXXX or its representatives to discuss the incident;

c) XXXXX will determine the severity of the breach in consultation with ________ and then discuss the course of action to be taken;

d) The Researcher will then be notified of the outcome.

#### 8.0 Dispute Resolution

8.1 Any dispute concerning the subject matter of this Agreement will be settled by full and frank discussion and negotiation between Parties. These discussions will take place kanohi ki te kanohi (face to face) and be conducted in a manner that is non-threatening and nurtures resolving of issues.

8.2 Should the dispute not be resolved satisfactorily by these means, the Parties agree that they will engage in mediation conducted in accordance with the terms and conditions of the LEADR New Zealand Inc Standard Mediation Agreement.

**Signed for and on behalf of XXXXX:**

Signature: Date:

Name: Position:

**In the presence of:**

Name of witness:

Signature:

**<u>Signed for and on behalf of ZZZZZ:</u>**

Signature: Date:

Name: Position:

**<u>In the presence of:</u>**

Name of witness:

Signature:

**<u>Read and Understood:</u>**

Signature: Date:

Name: Position:
